# Isostere ^18^F-protein post-translational editing enables dynamic tracking of neurodegeneration biomarkers

**DOI:** 10.1101/2025.01.26.634877

**Authors:** Nan Yang, Adeline W. J. Poh, Andrew M. Giltrap, Patrick G. Isenegger, Jeroen B. I. Sap, Joseph Ford, Tatsiana Auchynnikava, Kel Vin Tan, Sarah Able, Nikita Levin, Matthew Davy, Brian Josephson, Yue Yu, Maxwell W. G. Miner, Jatta S. Helin, Luciana Kovacs, Richard Aarnio, Noora A. Rajala, Matthew Tredwell, Shabaz Mohammed, Andrew J. Baldwin, Alex M. Dickens, Katherine A. Vallis, Jens Kuhle, Daniel R. McGowan, Anu J. Airaksinen, Daniel C. Anthony, David Leppert, Veronique Gouverneur, Benjamin G. Davis

## Abstract

The neurofilament light chain protein (NfL) is a suggested general marker for neuronal loss. Its release from brain parenchyma into cerebral spinal fluid, and presumed detection in blood has seen it established as a first blood-based marker of disease activity and drug efficacy in multiple sclerosis (MS) and in the presymptomatic diagnosis and assessment of disease course for other neurodegenerative disorders.^1^ However, the lack of characterisation of its behaviour in circulation, largely due to its antibody-dependent measurement, have hampered the biological interpretation of these measurements, especially after acute injury such as in MS relapse or head trauma.^2^ Here, we describe a strategy for exploiting positron emission tomography (PET) imaging using isosteric protein mimics following the installation of a fluorine-18 label that is benign enough to provide sensitive, real-time information on the dynamics and trafficking of NfL protein. This circumvents the limits of current methods that integrate ^18^F into proteins through the bio-conjugation of bulky, unnatural groups, which we show perturb NfL’s assembly and functional properties from those in the natural state. We use a visible-light-driven reaction to access radioactive isostere proteins that are unperturbed and so closely resemble their native form. In this way, generation of [^18^F]fluoroalkyl radicals that can be rapidly reacted at pre-defined sites on proteins creates mimics of proteinogenic side chains bearing near-zero-size labels to probe proteins in functionally ‘true’ form. These prosthetic-free, protein radiotracers can be generated in excellent radiochemical yield (up to 67%) via a semi-automated protocol in just 15 mins. High associated molar activities (precursor up to 102 GBq μmol^-1^) allowed high sensitivity dynamic observations in blood, brain and cerebrospinal fluid, enabling even the first unambiguous observations of spinal flow kinetics using proteins. These dynamics, including the high rate of spinal flow (on the order of mm per min) and drainage of NfL from CSF into sacral lymph nodes, now provides evidence that the slow fall rate of antibody-detected markers that is observed after acute neural insults is not due to a long half-life, but rather reflects sustained neuronal loss. This discovery will now help to better correlate clinical and radiological features of disease with NfL blood levels. Our methodology now demonstrates the broad potential of a near-zero-size labelling method for the functional study of proteins in whole organisms without interfering with their biological activity and native assembly.

## Introduction

Dynamic artefact-free tracking of proteins in living organisms in a non-invasive manner remains a holy grail. Its implementation would allow not only the discovery of new transport processes but also the accurate interpretation of putative indicators of physiological dysfunction. Neurofilament light chain (NfL) is a neuron-specific cytoskeletal protein (**Figure 1a,b**) that is an essential component of the highly-organised assembly of neurofilament (Nf) proteins, responsible for neuronal signalling, the preservation of axonal stability and neuron survival.^3^ Elevated levels of NfL in cerebrospinal fluid (CSF) and associated markers in blood are therefore a pathological feature of axonal injury and are associated with loss of neuronal function in multiple sclerosis (MS), traumatic brain injury (TBI) and primarily neurodegenerative diseases such as Alzheimer’s disease and other forms of dementing illnesses.^4^ Recent studies have established that NfL levels in cerebrospinal fluid and markers in blood are highly correlated, allowing the use of serum or plasma NfL-markers for detecting neurodegeneration.^5–7^ The dynamics of NfL in the blood are, however, not completely understood^8^ as blood concentrations (estimated in the 10 – 100 pg/mL range^6^) are below the threshold required for reliable detection by current non-invasive methods. Current *ex vivo* measurement techniques, such as single molecule array (‘Simoa’) assays, are based on antibody-mediated capture of NfL from blood and amplified signal;^9–12^ these are inherently blind to the types and quantity of NfL isoforms or fragments.^13,14^ Importantly, therefore this method does not allow the biological interpretation of the slow fall rate of NfL-marker levels^2,15,16^ after acute injury or of persistently elevated levels for months after a MS relapse^15^ or even minimal brain injury.^16^ Moreover, such static timepoint monitoring of NfL via capture-amplification does not allow real-time tracking and possible consequent associated statistical and analytical benefits that this might bring to the clinic.^17–19^ Direct fluorescent labelling of neurofilament subunits allows only visualisation at a cellular level (instead of whole-organism) and so is not directly relevant to clinical interpretation;^20^ associated fusing of fluorescent protein (FP) modules can also drive artefacts.^21^

**Figure 1.**
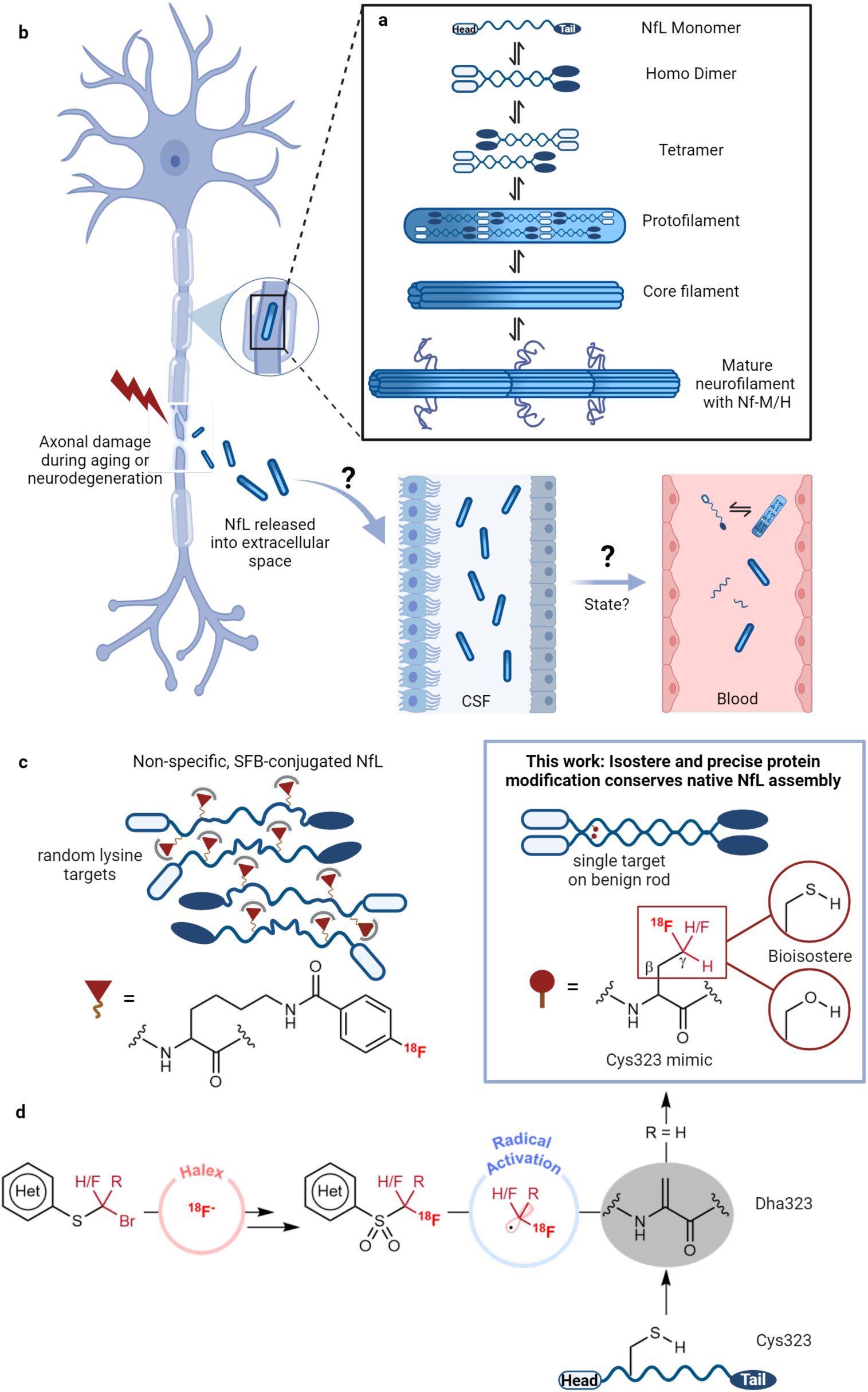
Isostere ^18^F-protein labelling for NfL for dynamic *in vivo* tracking. **a.** Presumed structure and assembly of NfL. NfL is composed of three domains, a central conserved α-helical rod domain sandwiched between N-terminal head and C-terminal tail domains containing sites for phosphorylation and *O*-linked glycosylation.^4,98^ NfL can exist in multiple assembly forms.^56,99^ In particular, the head and rod domains, essential for interactions between NfL subunits, drive dimerisation of NfL monomers. These dimers can further polymerise into tetramer and eventually form a core filament for Nf-M/H (medium/heavy) association.^100^ **b.** Pathophysiology of NfL into cerebrospinal fluid (CSF) and blood. In the event of an axonal injury, NfL is released from the neurons and elevated levels of NfL can be detected in the CSF. The underlying mechanism governing NfL release and trafficking between brain, CSF and blood has not been characterised. **c.** The installation of bioisosteric label at a pre-determined site on proteins (right) designed to overcome current methodologies conjugating bulky, prosthetic labels (non-specifically) (left) to access ^18^F-NfL without perturbation of its native state thereby enabling precise functional studies. Following this design an [^18^F]NfL ‘radioprotein’ can be generated as a precise functional mimic via post-translational editing Cys323 → Dha323 → ^18^F-Cys323 mimic using overall –CH_2_SH → CH_2_[^18^F]CF_n_H sidechain conversion. **d.** A designed two-step process – halogen exchange (halex) to generate ^18^F radical reagent and subsequent visible-light driven initiation to facilitate fluoroalkyl radical addition onto Dha for the installation of isosteric ^18^F-labels and ^18^F-amino acid (aa) side chain mimics in a rapid and efficient manner.

Fluorine-mediated detection offers a potentially sensitive approach to study proteins such as NfL even in complex biological matrix, such as the circulatory system, with minimal background interference even at low concentrations (in the micromolar to picomolar range) given its near complete absence in living systems.^22,23^ In particular, the radionuclide fluorine-18 can potentially be used to study fundamental processes in biology and disease by tracking the fate of (bio)molecules *in vivo* and in real time, dynamically at even picomolar levels using positron emission tomography (PET).^24–26^ Notably, however, most licensed PET tracers are small molecules.^27,28^ Many of the current methods suggested to install ^18^F onto proteins involve the conjugation of a bulky prosthetic group or linker, and also often lack control thereby leading to modification of multiple residues on a given protein.^29–31^ The resulting population of heterogeneously labelled proteins (**Figure 1c, left**) makes it challenging to correlate the relationship of an observed function to a particular molecular state or modification.^32,33^ Moreover, such labelling may modify or even ablate function.^34,35^

Here, through the development of a highly efficient protein ^18^F-labelling reagent with excellent molar activity (up to 102 GBq μmol^-1^), we describe a rapid and robust protocol for producing structurally-defined ^18^F-fluorinated proteins with near-zero-size radiolabels in excellent yield (RCY up to 67%), using reagents in low stoichiometry thereby minimising potential side-reactions. We demonstrate that the desired ^18^F-modified protein can produce unrivalled signal-to-noise in serial, dynamic imaging scans *in vivo*.^31,36^ This not only reveals NfL’s real time biodistribution by PET but also demonstrates the importance of imposing minimal structural changes (‘zero-size’) when studying protein function in order to preserve native behaviour.

## Results

### Design of fluoro-proteins for PET

Minimal size fluorinated motifs can be considered as (i) bioisosteres of native residues and/or (ii) ^18^F-mimics of amino acid side chains (**Figure 1c, right**). We reasoned that fluoroalkyl radicals, especially carbon-centred (C•) radicals, could be usefully exploited to create near-native protein constructs under conditions which complement radiofluorination (**Figure 1d**). Such C• radicals would exploit the stabilising and activating^37^ effects of adjacent C–F bonds allowing for potential generation *in situ* from visible light activation and single electron transfer (SET) processes in a benign and controlled fashion.^38^ More importantly, they can enable fast C–C bond forming reactions in water^37^ on timescales compatible with the half-life of fluorine-18 (109.8 minutes). Employed site-selectively, they could therefore create radio-bioisostere proteins (**Figure 1c**).

Previous elegant, radical radiofluorination methods to, for example, introduce small ^18^F-groups or transform C─H to C─^18^F bonds directly have been restricted to only simple biological molecules, i.e. peptides, and with low molar activity (1.3–5.3 MBq μmol^−1^) and sometimes modest radiochemical yield (RCY, 11–56%). They can also lead to mixtures of products as a result of the formation of multiple regioisomers.^39,40^ Furthermore, the required use of organic co-solvent^39,41^ critically limits the usefulness of certain protocols for protein labelling, especially for those with sensitive aggregation properties that bear challenging, intrinsically disordered regions (IDRs) such as NfL.

By taking advantage of the exquisite polarity match between nucleophilic C-centred C• radicals bearing alkyl substituents and electrophilic radical-acceptor dehydroalanine (Dha) residues, selective single-site modification dominates^42–44^ over any undesired side-reaction with endogenous amino acid side chains, such as tyrosine and tryptophan (**Figure 1d**).^45–49^ Dha can be readily introduced into native proteins^50^ using various methods from low-abundance amino acids such as natural cysteine^51,52^, modified phosphoserine^53^ and unnatural selenocysteine^54,55^ variants. We therefore thought to explore its potent SOMOphilicity with C-centred [^18^F]fluoroalkyl radicals [^18^F]CF_n_• to generate high-yielding, homogenous radioproteins.

Notably, NfL contains a single cysteine (Cys323) present in the intrinsically disordered rod domain that does not participate in disulfide bond formation.^56,57^ Therefore, we reasoned, this would allow for benign Dha generation (Dha323) without interfering with NfL core assembly (**Figure 1d**). The installation of a radiolabelled side chain mimic at this Dha site via radical chemistry would allow the recapitulation of a ‘near-wild-type’ protein sequence. The CF_2_H motif is a known bioisostere of the cysteine thiol^58^, as like the S–H group^59^, the C–H bond in CF_2_H can serve as a hydrogen bond donor (atypical for C–H bonds).^60^ The use of a carbon-centred [^18^F]•CF_2_H radical^44,49,61^ or similar could therefore yield a Cys-mimicking side chain through overall SH ⇉ CF_2_H transformation and hence allow the potential construction of a minimally perturbed radio-protein equivalent.

### Scoping for optimal ^18^F-fluorinated radical precursor reagents

We identified two synthetic requirements for our ^18^F-protein design: first, generation of [^18^F]•CF_n_H radicals from any putative precursor should be rapid, protein-compatible and efficient not only for productive ^18^F-protein labelling but also for minimising unwanted ^18^F-side products; and second, fluorine-18 incorporation into the radical precursor should be rapid and high yielding (**Figure 1d**). To satisfy these requirements, we envisaged the optimised combination of (1) *in situ* visible light activation of *N*-heteroaromatic sulfone reagents^61^ with (2) an ^18^F-halogen exchange (‘halex’) reaction^62^.

*N*-heteroaromatic sulfone reagents are known fluoroalkyl radical sources,^49,58,61^ capable of initiation via photostimulated outer-sphere SET metal complexes (‘photoredox catalysts’). To ensure that this could occur within a redox window that would allow for protein-compatible radical activation (and avoid unwanted side reaction of native redox-active residues^63,64^), we designed precursor sulfones bearing systematically altered N-heteroaromatic motifs: pyridine (**S1**: pySOOF), pyrimidine (**S2**: pymSOOF) and benzothiazole (**S3**: btSOOF). In this way (**Figure 2a**) controlled variation of the potential needed for radical initiation (**S1**: E_red_ = -1.50 V, **S2**: E_red_ = -1.35 V, **S3**: E_red_ = -1.17 V *vs* saturated calomel electrode, SCE^61^) would allow scoping for benign generation and addition of [^18^F]•CF_2_H radicals at Dha residues to create the cysteine radiobioisostere [^18^F]*bis*-fluorohomoalanine (*bis*[^18^F]FhAla or [^18^F]*b*Fha) through overall SH ⇉ CF_2_H chemical mutation (**Figure 2a**). Since [^18^F]F^-^ is produced from the cyclotron in the low nano-to picomole range,^65^ ^18^F-labelled species are typically the limiting reagents in radiochemical reactions. Therefore, radical generation from our chosen sulfone precursors needed to be highly efficient to drive sufficient ^18^F-incorporation into proteins.

**Figure 2.**
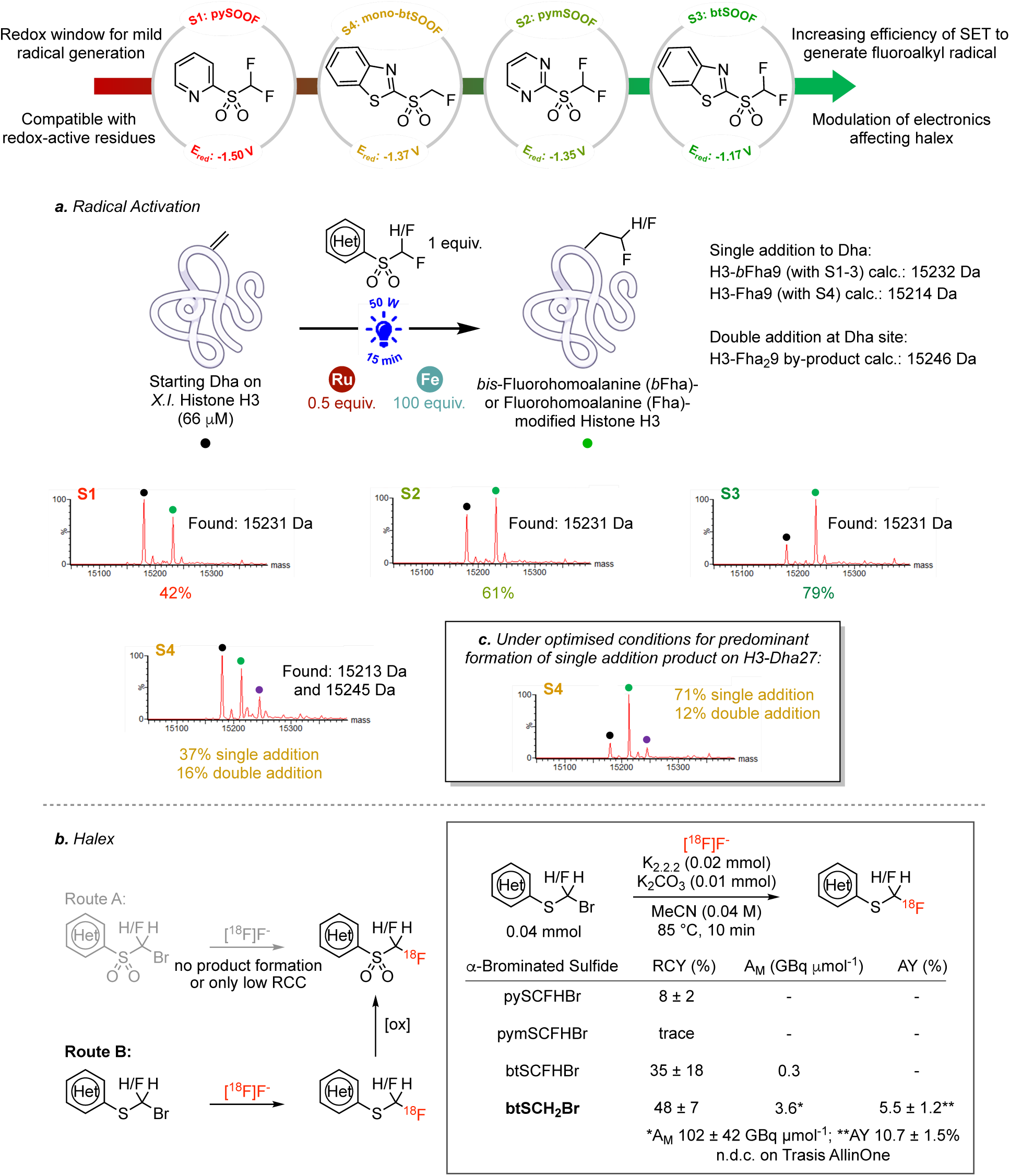
Use and generation of C•-centred radical precursors for the synthesis of bio-isosteric radioproteins. The heteroaryl moieties of the halex-initation route (Figure 1d) modulate both steps. **a.** Heteroaromatic sulfones as fluoroalkyl radical precursors could allow installation of ‘near-zero-size’ ^18^F side chains into proteins. Reduction potentials of sulfones with pyridine, pyrimidine and benzothiazole backbones are within the redox window for residue-specific and protein compatible radical reaction. Under degassed buffered conditions (500 mM NaOAc, pH 6, 3 M Gdn•HCl, <10 ppm O_2_) using sulfones as radical-generating reagents (1 equiv.), FeSO_4_•7H_2_O as the reductant (100 equiv.) and sub-stoichiometric amount of Ru(bpy)_3_Cl_2_•6H_2_O as the photocatalyst (0.5 equiv.), btSOOF (**S3**) efficiently delivered the desired single addition product with the introduction of isosteric CF_2_H group onto the Dha site (>79% conversion to histone protein H3-*b*Fha9; protein 1 mg mL^-1^, 450 nm blue light, 50 W, 15 min at rt) (Supplementary Information). Increasing formation of desired ‘edited’ product is observed as reduction potentials of the fluoroalkyl sulfone radical precursors decrease. Unoptimized mono-btSOOF (**S4**) as a source of •CH_2_F radical also generated some double addition by-product (addition of a second CH_2_F group on the α-carbon backbone, 16% double addition; single:double ratio is 2.3:1.0). **b.** Direct nucleophilic substitution alpha to sulfone (S(VI)) fails but is successful alpha to sulfide (S(II)). The heteroaryl moiety of the substrate also modulates halex reactivity. Sulfides with benzothiazole as the aryl group proved necessary for efficient ^18^F incorporation (reflected by radiochemical yield (RCY) values of the ^18^F/Br exchange (halex) reaction). The starting fluorine-containing sulfide (btSCHFBr) resulted in an order of magnitude decrease in molar activity of the halex product compared to btSCH_2_Br, attributed to a competing non-radioactive fluorine-releasing decomposition pathway (Supplementary Information). **c.** Mono-btSOOF (**S4**) reaction optimisation to favour single addition product over unwanted double addition. Reducing light flux and increasing reaction times (10 W, 30 min) successfully produced the desired single addition product with excellent yield and importantly, minimised the generation of unwanted double addition (for experimental details, see Supplementary Information).

To demonstrate broad relevance to IDRs in proteins, initial scoping used the N-terminal IDR^66^ found in intact histone proteins as a model. Our initial studies with the three non-radioactive •C^19^F_2_H radical precursors (**S1, S2, S3**) revealed comparably efficient conversions (83-93% by mass spectrometry) to –CH_2_CF_2_H side chains to generate *bis*-fluorohomoalanine (*b*Fha)-containing histone model protein H3-*b*Fha9 after 15 minutes of blue light exposure (450 nm, 50 W) under degassed conditions with excess Ru(bpy)_3_Cl_2_•6H_2_O as the photocatalyst (2 equiv.; E_ox_ [Ru(II)/Ru(I)] = -1.33 V *vs* SCE) (Supplementary Table 1). BtSOOF (**S3**) also gave low levels of double addition product (12%)^44^ arising from quenching of an alpha C• intermediate with a second •CF_2_H. To minimise this side reaction the catalyst loading was lowered to sub-stoichiometric levels (0.5 equiv. of photocatalyst relative to 1 mg/mL protein concentration) and efficiently produced desired H3-*b*Fha9 protein (conversion ∼80%) when btSOOF (**S3**) was employed as the radical precursor; pySOOF (**S1)** and pymSOOF (**S2)** under these conditions gave the product with lower 42% and 61% conversion, respectively, consistent with their graduated E_red_ values (**Figure 2a**).

Having established efficient radical reactivity based on tuned precursors, our second requirement was efficient installation of fluorine-18 into the radical precursor. ^18^F-Halex reaction of an appropriate heteroaromatic precursor would enable a direct telescoped, even one-pot, approach as a viable route to access equivalent ^18^F-labelled radical precursor reagents [^18^F]**S1-S3**. Initial halex reaction screens on the α-brominated fluorosulfones pym- and bt-SOO–CFHBr under typical halex conditions were unsuccessful, yielding instead only S_N_Ar products from substitution at the C-2 aryl site (Supplementary Table 2), despite the precedent for non-metal catalysed α-halosubstitution.^67^ Similarly, whilst attempted halex on py-SOO–CFHBr gave some desired [^18^F]pySOOF reagent **S1**, the yield was low (RCY = 7%).

These poor results were consistent with Bordwell’s long-standing suggestion^68^ that whilst nucleophilic substitution alpha to sulfur (i.e. as sulfonyl S(VI)) may be considered electronically analogous to those alpha to carbon in carbonyls, any such activating effects are over-shadowed by steric effects; in essence sulfonyl S(VI) can be considered to show formal reactive similarity to neopentyl. Therefore, despite establishing initial low-level access to [^18^F]**S1**, this inefficiency combined with **S1**’s lower protein labelling efficiency in comparison to other heteroaryls (i.e., **S2** and **S3**), prompted us to reconsider the design of the halex precursor. We therefore chose other oxidation states of sulfur. We dismissed sulfinyl S(IV), as the additional sulfoxide stereocentre would show only partly reduced steric burden^68^ and also generate diastereomeric mixtures of ^18^F-labelled products thereby further complicating downstream analysis. We instead focussed our efforts on the potential of exploiting substitution at carbon alpha to the least hindered sulfenyl S(II) oxidation state^69^ (i.e. het-S-CFHBr).

Het-S-CFHBr halex precursors were readily generated by direct alkylation of a corresponding thiol precursor (Het-SH) using a corresponding halide (i.e., dibromofluoromethane or equivalent). Interestingly, these again displayed a reactivity trend that varied with the nature of the heteroaryl motif. Halex reactions using pyrimidyl (pym) and pyridyl (py) heteroaromatic systems (to yield [^18^F]pymSCF_2_H and [^18^F]pySCF_2_H, precursors to [^18^F]**S2 and** [^18^F]**S1**, respectively) were poor and broader screening conditions saw no further improvement to the RCY (**Figure 2a**, Supplementary Tables 3 and 4). Pleasingly, α-brominated fluorosulfide btS–CFHBr underwent rapid and efficient halex reaction with [^18^F]F^-^ to give [^18^F]btSCF_2_H. When followed by subsequent high yielding (“quantitative”) ‘on-cartridge’ oxidation with *in situ* generated RuO_4_ (using RuCl_3_•xH_2_O and NaIO_4_) this effectively delivered desired [^18^F]btSOOF ([^18^F]**S3**) (**Figure 2b**, Supplementary Table 5).^44^ The radiochemical yield (RCY, as measured by HPLC and TLC analysis of the crude reaction mixture) of the halex reaction could be improved to 35 ± 18% (n = 3) (*vs* 6 ± 1 % (*n* = 2) RCY), through the addition of excess Kryptofix 2.2.2 and K_2_CO_3_ highlighting their importance in facilitating [^18^F]F^-^ solubilisation into organic solvents and enhancing [^18^F]F^-^ reactivity for nucleophilic substitution^31^.

As well as radiochemical efficiency, good molar activity^70^ is vital for the sensitivity needed for dynamic use of a radiotracer. Surprisingly, despite the good RCY obtained for the production of [^18^F]btSOOF (**S3**), it could only be isolated with a low molar activity of 0.30 GBq µmol^-1^ (**Figure 2b**). We reasoned that this low final molar activity might be attributed to competing decomposition of the starting reagent btSCHFBr under the halex conditions, resulting in the release of fluorine-19 from the starting reagent. The resulting competing [^19^F]F^-^ would then lead to eventual accumulation, via dilution, of non-radioactive btSCF_2_H as a by-product, even in the absence of an external fluoride source (Supplementary Note 3).^71,72^

### Prevention of isotopic dilution allows efficient and site-selective creation of radioproteins

With the aim of increasing molar activity, we next tested this dilution hypothesis with a non-fluorinated starting halex precursor, btSCH_2_Br, that would preclude such a putative isotopic dilution event. Gratifyingly, application of the halex protocol to btSCH_2_Br delivered [^18^F]btSCH_2_F in an excellent yield (48 ± 7% (n = 2) RCY) and an isolated activity yield of 5.5 ± 1.2% (n = 4, non-decay corrected starting from ∼10-15 GBq of dried [^18^F]fluoride on a manual radiosynthesiser) after RuO_4_-mediated oxidation to [^18^F]**S4** (Supplementary Information). In this way, circumvention of competing dilution generated [^18^F]**S4** with a molar activity that was an order of magnitude greater than that obtained for [^18^F]**S3** (3.55 vs 0.3 GBq μmol^-1^, respectively) (Supplementary Information).

Consistent with altered reduction potential (**Figure 2a**, E_red_ = -1.37 V vs. SCE^61^) and altered fluorination state^37^ the reactivity of **S4** was slightly reduced and initially gave lower conversions (**Figure 2b** 37% vs **S1**: 42%; **S2**: 61%; **S3**: 79% with sub-stoichiometric photocatalyst) to a corresponding fluorohomoalanine (Fha)-modified histone protein H3-Fha9. Notably, however, previously suggested requirements for additional electron-withdrawing groups on the heteroaryl moiety^61^ or on the C-centred radical^44^ proved not to be necessary for efficient radical formation. Ready reoptimization, which was guided by characterization and minimization of competing side-reactions (**Figure 2b**, Supplementary Notes 1, 2), revealed that slightly prolonged reaction times (by a further 15 – 30 min) and reduction of light flux (from a 50 W source to 10 W) coupled with use of low photocatalyst loading (1-2 equiv.) allowed good production (> 70%, **Figure 2c**) of fluoro-protein.

### Efficient automation-assisted generation of mimic radioproteins with ‘near-zero-size’ protein ^18^F-labelling using [^18^F]-carbon-centred radical sidechains

Having identified [^18^F]mono-btSOOF ([^18^F]**S4**) as a potentially powerful reagent for radioprotein generation, we next implemented its automated production and utilization. Thus, a cassette-based platform (Trasis All-In-One) allowed radiosynthesis of [^18^F]**S4** in high molar activity (*A*_m_ = 102 ± 42 GBq μmol^-1^) and an activity yield of 10.7 ± 1.5% (non-decay corrected using ∼25 GBq of cyclotron-produced [^18^F]F^-^) in a manner that was found to be readily programmed by others.

Using this workflow, we successfully demonstrated ^18^F-labelling at varied representative sites to probe potential location-dependent variation in different histone IDR proteins; all were converted in excellent yields (up to 67% RCY) to create ^18^F-labelled human histone H3-[^18^F]Fha4, ^18^F-labelled *Xenopus laevis* histone H3-[^18^F]Fha27 and ^18^F-labelled histone H3.NTEV-[^18^F]Fha2 using [^18^F]**S4** as the radical precursor and photocatalyst loading as low as 0.5 equivalents relative to protein concentration (Supplementary Information, 125 μM protein concentration, 0.5 – 2 equiv. Ru(bpy)_3_Cl_2_•6H_2_O, 100 – 250 equiv. FeSO_4_•7H_2_O, NH_4_OAc (500 mM, pH 6, 3 M Gdn•HCl) or HEPES (100 mM, pH 7.4) buffer with <10% DMSO, 10 – 50 W blue LED, RT, 15 – 30 min and <10 ppm O_2_ levels). Consistent with designed rapid reactivity and the required short reaction times needed for use of fluorine-18, increasing reaction times did not result in an appreciable increase in RCY (56% RCY in 30 min vs 62% RCY in 45 min) (Supplementary Information). Size-exclusion chromatography (PD SpinTrap G-25) purified the radiolabelled protein from the reaction components, achieving excellent radiochemical purity (>99%) and UV-determined purity (>99%) in under 5 min for downstream *in vivo* applications (**Figure 3b**). In this way, the entire protocol – from radio-precursor synthesis to accessing ^18^F-labelled protein – was completed within a single half-life (t_1/2_ ∼110 min) of fluorine-18.

**Figure 3.**
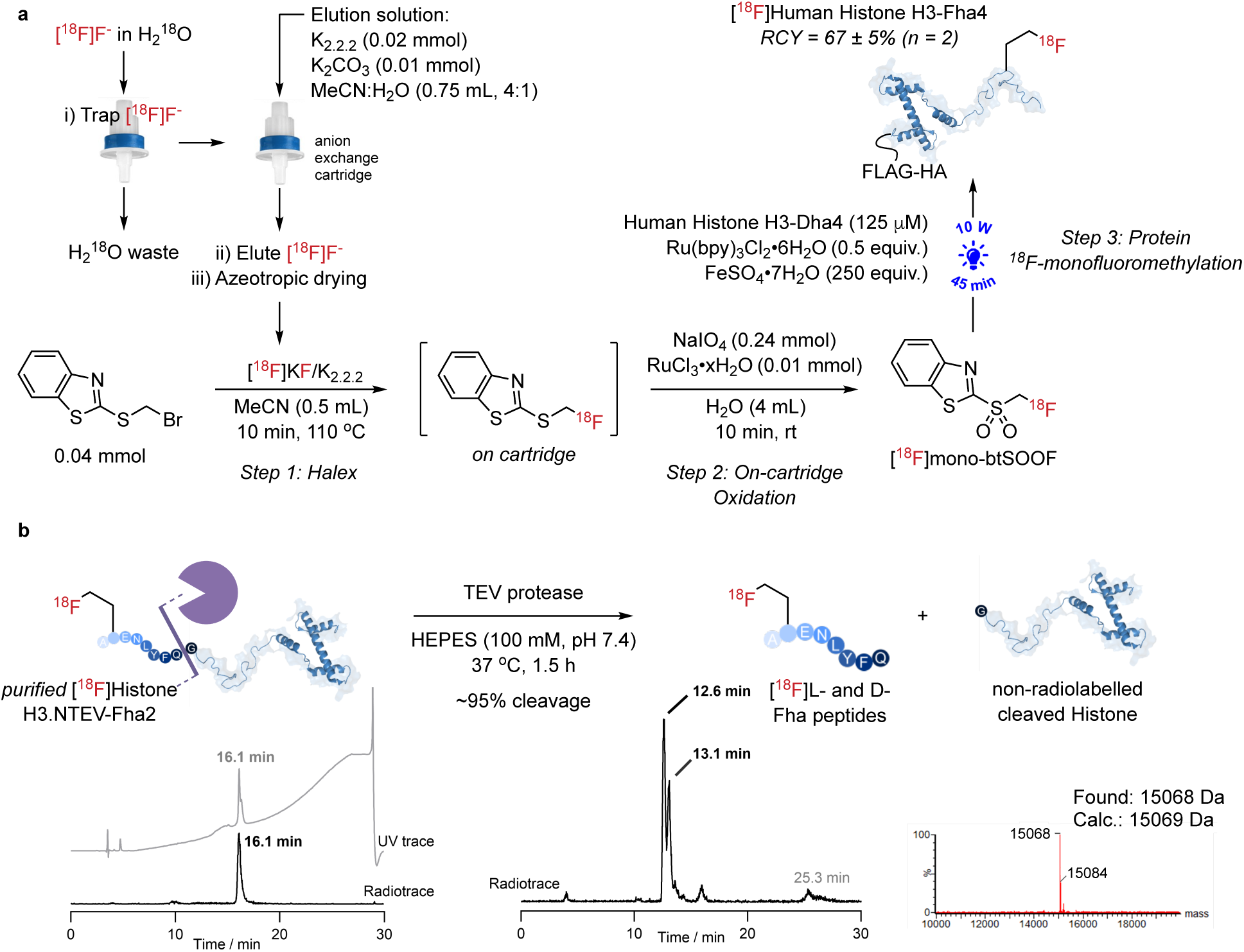
Efficient generation of mimic radioproteins with ‘near-zero-size’ protein ^18^F-labelling using ^18^F-carbon-centred radical sidechains and characterization of its site selectivity. **a.** The workflow for generating a protein-compatible, ^18^F-fluorinating reagent, [^18^F]mono-btSOOF (**[^18^F]S4**, involved a fast two-step process – halogen exchange (halex) at the α-carbon of an S(II) intermediate (10 min) followed by on-cartridge oxidation to S(VI) (10 min) – and semi-preparative HPLC purification to give an isolated activity yield of 5.5 ± 1.2% (n = 4) (n.d.c. on a manual radiosynthesiser) in under 90 min (i.e., within the half-life of fluorine-18). Protein ^18^F-labelling in the presence of a Ru(bpy)_3_(II) photocatalyst and an Fe(II) reductant was conducted in a blue light photoreactor (450 nm) in degassed aqueous buffer (Supplementary Information). For human histone protein H3-Dha4, radiolabelling was achieved in excellent yields (67 ± 5% (n = 2)) even using a sub-stoichiometric amount of photocatalyst (0.5 equiv.) and low micromolar protein concentration (125 μM). **b.** ^18^F-labelled histone protein with a Dha residue situated in its N-terminal region [^18^F]Histone H3.NTEV-Fha2 could also be generated in an excellent yield (67% RCY) and isolated with up to 98% radio- and UV purity (bottom left traces). Reaction conditions: 125 μM protein solution, Ru(bpy)_3_Cl_2_•6H_2_O (2 equiv.), FeSO_4_•7H_2_O (250 equiv.) 500 mM NaOAc, pH 6, 3 M Gdn•HCl, <10% DMSO, <10 ppm O_2_, 450 nm blue light, 50 W, 15 min at rt. Using TEV protease the N-terminal peptide could be cleaved for direct analysis of site-selectivity. The radio-HPLC chromatogram showed the ^18^F-labelled N-terminal epimeric peptide product as the dominant radioactive species (>95% cleavage; full-length [^18^F]Histone H3.NTEV-Fha2 elutes at 16.1 min on a 30 min HPLC run (bottom left) or at 25.3 min on a 50 min HPLC run (bottom right)), thereby demonstrating site-selective labelling essentially exclusively at the targeted site.

Finally, to further ease translation of ^18^F-radioprotein generation into general radiochemistry practice, the methodology was additionally adapted to remove any need for specialist equipment in a typical radiopharmacy. In particular, control of oxygen levels within the reaction buffer (<10 ppm) reduces unwanted oxidative damage to proteins under photochemical reaction conditions^44^ and can be achieved through buffer equilibration in a glovebox. We demonstrated that simply purging of the reaction mixture with nitrogen for 20 min also proved sufficient to ensure anaerobic conditions, which allowed for excellent conversion (>85% with **S3**) with negligible oxidative damage. This in turn enabled more general implementation in different radiochemistry laboratories (see Supplementary Information for further details).

### Determining site-selectivity in ^18^F-radioprotein synthesis

Characterization of radiochemical products necessarily dictates additional corroborative methods beyond those that may be readily applied to ‘cold’ products. We therefore complemented our full characterization of ‘cold’ [^19^F]-histone-H3 by devising an additional method to assess site selectivity in [^18^F]-histone-H3. For example, tryptic LC-MS/MS is not readily available for radioactive materials. Moreover, unlike ^18^F-labelling on small molecules (in which installation of a F atom can sufficiently change structure and polarity to aid characterization), the conversion of a sidechain –CH_2_**SH →** –CH_2_**[^18^F]CH_2_F** in a protein represents a minimal structural change in line with our design of minimal protein alteration. Such bioisostere proteins are, therefore, unlikely to show significant differences in retention times from their blueprint wild-type counterparts during chromatography^44^ (and so perhaps from some other undesired protein by-products).

To evaluate site-selectivity, and again with an emphasis on translatable potential, we therefore designed a cleavable protein system capable of releasing an appropriate ‘reporter fragment’ that might be readily analyzed by traditional radiochemical means. To test our hypothesis, fluorine-18 was first successfully installed using our standard methods into a histone protein variant to generate an N-terminal sequence A-**[^18^F]Fha**-ENLYFQ…. using [^18^F]**S4**, again in excellent yield (67% RCY). This protein construct is, by virtue of its N-terminal sequence, susceptible to specific cleavage at its Gln8 residue by tobacco etch virus (TEV) protease. Strikingly, incubation with TEV-protease (**Figure 3b**, Supplementary Information) showed essentially complete (>95%) cleavage by radio-HPLC of the radioemission-associated protein peak at t_R_ = 25.3 min, consistent with site-selective location of ^18^F exclusively at site 2 in the N-terminus of the protein. This concomitantly yielded two radioemission-associated peaks at t_R_ = 12.6 and 13.1 min (**Figure 3b**) corresponding to peptides A-L-**[^18^F]Fha**-ENLYFQ and A-D-**[^18^F]Fha**-ENLYFQ (consistent with the known^50^ generation of D/L-epimers by reaction of C• centred radicals with Dha). Their identities were further, unequivocally confirmed by chromatographic comparison using corresponding authentic ^19^F-labelled synthetic peptide standards. Together these data confirmed exclusive ^18^F-labelling at the intended protein residue site.

### Bioisoteric NfL generated using zero-size methods mimics WT behaviour in contrast to classical radioprotein labels

To characterise structural and hence, functional consequences of modifications on NfL, we compared the aggregation kinetics of bioisosteric NfL to wild-type NfL (NfL-WT). The assembly of NfL is a key mode-of-action, with likely additional consequences for its circulation dynamics as a putative biomarker; spectroscopic observation offers an effective method of capturing the global evolution of transient and heterogenous oligomeric states over time under near physiological conditions.^73,74^ Moreover, unlike widely used fluorescence techniques, such methods do not require an exogenous probe molecule that may perturb, or even promote, aggregation.^75^ For current ^18^F-protein radiolabelling methods, which exploit addition of prosthetics bearing fluorine-18, the use of [^18^F]SFB ([^18^F]-*N*-succinimidyl 4-fluorobenzoate) is often regarded as the ‘gold standard’^76^ ^18^F-reagent for protein radiolabelling. We therefore made a direct comparison of the functional consequences of using this method.

NfL self-assembly kinetics were determined spectroscopically in two ways: through light scattering experiments measuring turbidity at 350 nm (**Figure 5a**) and by NMR spectroscopy (**Extended Data Figure 1**) monitoring changes in aggregate and monomer concentrations, respectively. Observed oligomerisation, initiated upon dilution of chaotropic agent urea on samples rigourously purified by analytical ultracentrifugation, revealed assembly kinetics of isosterically labelled NfL-Fha323 closely matched those of NfL-WT under the same conditions, as observed by both scattering (**Figure 5a**) and NMR spectroscopy (**Extended Data Figure 1**). By contrast, heterogeneously-SFB-modified NfL showed greater deviation, with deviation differences increasing in line with total SFB conjugation level over a range of solution conditions and total protein concentrations (**Figure 5a** and **Extended Data Figure 1b**). Together these data suggested that our editing of Fha323 into NfL can be considered as a ‘minimal label’, here preserving the native assembly behaviour of NfL and consistent with fully retained functional state.

### Dynamic tracking of [^18^F]NfL reveals short peripheral circulation half-life and delineates protein drainage modes from the brain

After successfully demonstrating the reactivity and Dha-selectivity of radical reactivity generated from [^18^F]**S4** on varied model protein sites, we next tested it in the generation of our target radiobioisostere protein NfL-[^18^F]Fha323 via direct ^18^F-labelling of NfL through the creation of an isosteric Cβ–[^18^F]CγH_2_F bond from the precursor Cβ– SH bond at residue 323 in wild-type NfL (**Figure 1d, 4**). Recombinant mouse NfL (mNfL) was generated in good yield through mammalian cell expression (HEK293T, Supplementary Information). This also ensured display of native post-translational phosphorylation (Supplementary Information), the identity of which was demonstrated by fluorescence-based phosphoprotein-specific detection (so-called ‘Pro-Q diamond’ methods^77,78^) and further confirmed through enzymatic dephosphorylation (λ-phosphatase) (Supplementary Note 4).

**Figure 4.**
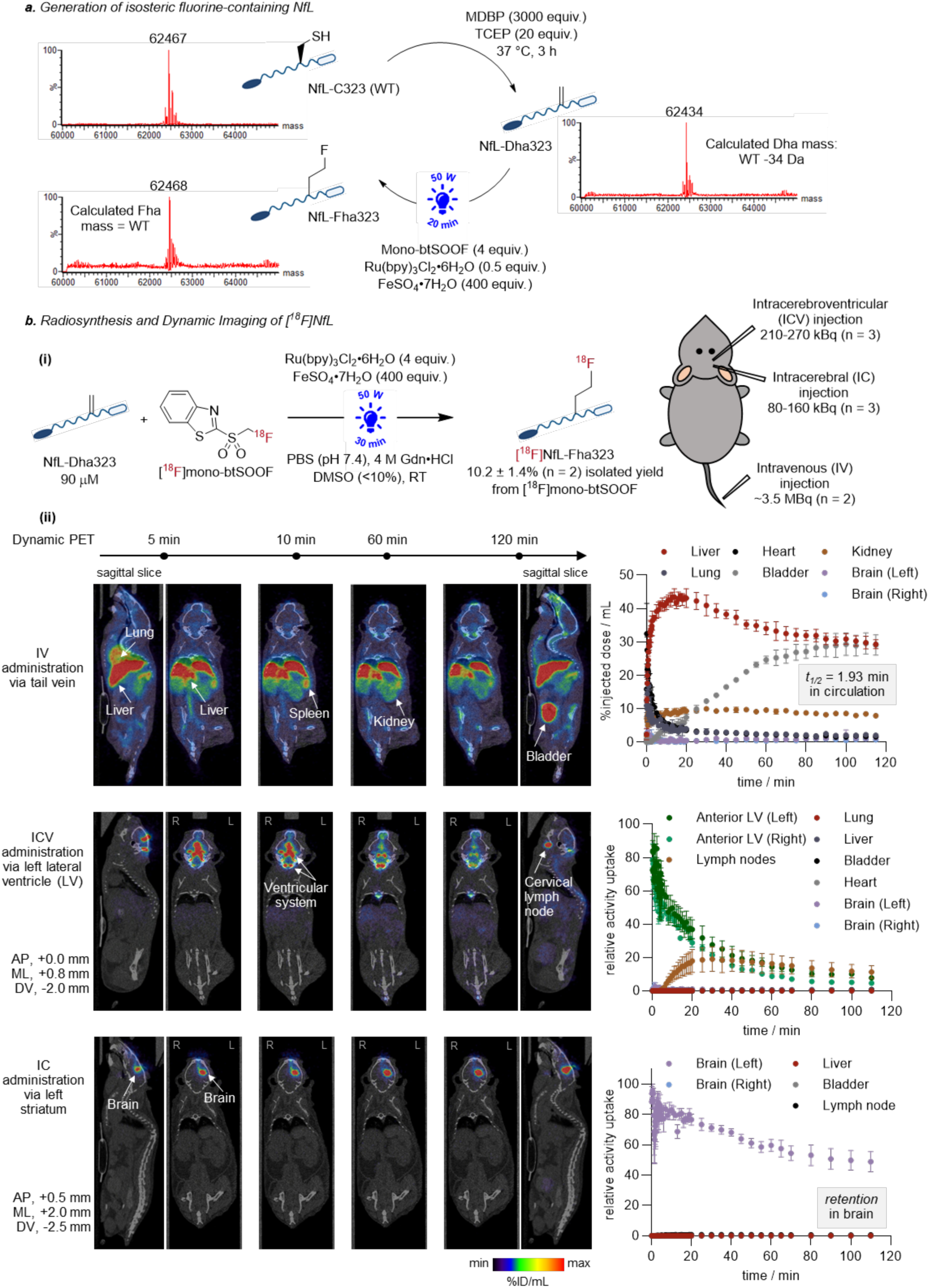
Generation of [^18^F]NfL reveals its utility as a tracer. **a.** WT NfL (and all its phosphoforms) containing a single Cys was chemically converted to the Dha-containing protein and systemically modified via visible light-mediated SET-initiated radical reaction to introduce the fluorine-containing isosteric label, Fha. Conversions were confirmed by LC-MS. **b. (i)** Efficient generation of [^18^F]NfL (10.2 ± 1.4% (n = 2) isolated activity from 2.5 – 3.0 GBq of [^18^F]mono-btSOOF (**[^18^F]S4**) produced from the Trasis AllinOne automated platform) allowed for intravenous, intracerebral and intracerebroventricular monitoring in dynamic *in vivo* PET (see also **Supplementary Movies**). Reaction conditions: 90 μM protein solution, Ru(bpy)_3_Cl_2_•6H_2_O (4 equiv.), FeSO_4_•7H_2_O (400 equiv.), PBS, pH 7.4, 4 M Gdn•HCl, <10% DMSO, degassed conditions, 450 nm blue light, 50 W, 20 min at rt. **(ii)** In naïve mice intravenous administration revealed the short half-life of [^18^F]NfL in circulation and its rapid uptake primarily by the liver. [^18^F]NfL injected intracerebroventricularly distributed throughout the ventricular system, drained into the cervical lymph nodes and was transported along the spinal cord. [^18^F]NfL is largely retained in the brain after intracerebral injection. Time-activity curves (TAC) were reported as %injected dose per mL (ID mL^-1^) for IV experiments and relative activity uptake for ICV- and IC-related studies (Supplementary Information).

To install the Cβ–[^18^F]CγH_2_F side chain from precursor Cβ–SH, *bis*-alkylation/elimination with methyl 2,5-dibromopentanoate (MDBP) under standard conditions^79,80^ converted the free cysteine at site 323 in NfL-WT to Dha to give NfL-Dha323 (**Figure 4a**, Supplementary Information). Reaction of **S4** (2 equiv.) with NfL-Dha323 was efficient and effective with, again, low stoichiometry of the photocatalyst (0.5 equiv. Ru(bpy)_3_Cl_2_•6H_2_O, >95% conversion, **Figure 4a**, Supplementary Information). Moreover, protein concentrations even as low as 10 μM could be modified with efficiencies of up to 70% (100 equiv. **S4**, 5 equiv. Ru(bpy)_3_Cl_2_•6H_2_O, 400 equiv. FeSO_4_•7H_2_O in NH_4_OAc 500 mM, pH 6, 3 M Gdn•HCl buffer, 50 W blue LED, RT, 15 min, <10 ppm O_2_). This efficient use of low concentrations of protein substrates is notable for those such as NfL that are prone to aggregation at higher concentrations – in this way, NfL represents an excellent and stringent test. Notably, while the highly reactive benzothiazole-based sulfonyl radical precursors developed here could modify NfL protein, negligible modification of NfL was detected with the corresponding pySOOF radical reagent under these conditions. Finally, reaction of low concentration (70.0 μM) NfL-Dha323 with [^18^F]**S4** to yield radioprotein NfL-[^18^F]Fha323 proved successful even despite the only low, limiting (nano-to picomole) quantities of [^18^F]**S4** that are necessarily generated (39 ± 7% (n = 2) RCY based on HPLC, radiochemical purity >99%; UV-determined purity >98%) (Supplementary Information). Importantly, our automated protocol allowed efficient access to an [^18^F]NfL radiobioisostere with sufficient isolated activity (10.2 ± 1.4% (n = 2) from 2.5 – 3.0 GBq of [^18^F]mono-btSOOF) for imaging of not only endpoint biodistribution but also dynamic observations in blood and the CSF via PET (**Figure 4bi**).

First, following intravenous injection (IV) of NfL-[^18^F]Fha323 (∼3.5 MBq (n = 2 animals)) into naive mice, no adverse effects were observed. Time-course PET revealed rapid multi-pass circulation from the site of injection in the tail to the heart and lungs. The calculated half-life in the blood, measured by a volume of interest (VOI) placed in the left ventricle of the heart, revealed a very short half-life of 1.93 min (Supplementary Information). [^18^F]NfL was rapidly cleared from blood into the heart and lungs, concomitant with redistribution into the liver, plateauing at approximately 90 min, and to a lesser extent, the spleen (see **Supplementary Movie 1** and representative timepoints in **Figure 4bii, top** and **Extended Data Figure 2a** for post-PET, organ-specific *ex vivo* analyses of accumulated dose). In the kidneys, a steady state was reached quickly after 10 min, whereas in the bladder, continuous accumulation was observed throughout the dynamic scan duration (**Figure 4bii, top**). No significant accumulation was noted in the brain, CSF, or lymph nodes (**Figure 4bii, top** and Supplementary Information). The rapid clearance of radiolabelled [^18^F]NfL from blood (as seen also through gamma-counting analyses) was further supported by antibody (anti-NfL)-mediated (Western) analyses that showed apparent complete absence of antibody-detectable NfL-WT in blood plasma as soon as 20 min after an intravenous administration (40 μg injected). Indeed, NfL-WT was only successfully detected in samples shortly after administration (Supplementary Information).

Second, efficient labelling also enabled the administration of NfL-[^18^F]Fha323 intracerebroventricularly (ICV) via the left lateral ventricle for serial PET imaging in the CSF (210-270 kBq μL^-1^, n = 3 animals). 1 μL injections were performed using a capillary needle with a 50 μm lumen to ensure precise stereotaxic delivery while minimising damage and leakage from the injection site. ICV injection of radiolabelled [^18^F]-NfL contrasted the intravenous result; the radiotracer was initially distributed solely throughout the ventricular system (clearly visualised in the early frames of imaging) before then progressing down the spinal cord (see **Supplementary Movie 2** and **Figure 4bii, middle**). Notably, no significant uptake was detected in the brain parenchyma, either ipsilateral or contralateral to the site of injection in the striatum adjacent to the ventricles. Rapid drainage of NfL into the deep cervical lymph nodes was also observed, with an initial signal appearing after 5 min and a plateau reached at 20 min (**Figure 4bii, middle**). Strikingly, drainage from the lumbar cord to the sacral lymph nodes was also observed after 40 min (**Extended Data Figure 3**). Low-level tracer signal was detectable in the liver after 5 min in 1 out of 3 animals, while very low levels were detected in the kidney and bladder (**Extended Data Figure 2**). The potential drainage of [^18^F]NfL from the CSF to the bone marrow of the skull was explored, but no evidence of such a route was found. Draining via the cribriform plate was also not observed. Instead, distribution of radiolabelled NfL in one of the animals suggested possible drainage along the auditory tract.

Third, following intracerebral (IC) injection of NfL-[^18^F]Fha323 into the left striatum (60-80 kBq μL^-1^), there was no apparent drainage to the blood or bone, contrasting even more sharply with the ICV injection (n = 3 animals). [^18^F]NfL remained localised within the ipsilateral brain parenchyma and did not cross the midline via the corpus callosum even after 2 h post-injection (see **Supplementary Movie 3** and **Figure 4bii, bottom**). In 1 out of 3 animals, radiolabelled NfL was detected in the CSF, with subsequent accumulation in the cervical lymph nodes after approximately 15 min, similar to but delayed compared to ICV injection spleen (see **Supplementary Movie 3** and Supplementary Information). Again, no clear drainage pathway from the brain to the periphery via the skull or cribriform plate was observed; low tracer signals were detected in the kidney and bladder (**Extended Data Figure 2**).

Finally, such was the activity and sensitivity possible from isosteric protein ^18^F-labelling and serial dynamic PET that real-time observations even enabled use of NfL-[^18^F]Fha323 to estimate rates of spinal-cord transport. Triplicate measurements based on VOI placement in the thoracic and sacral spinal cord segments allowed a protein migration rate of approximately 0.8 – 1.5 mm min^-1^ to be determined (**Figure 5b**). To our knowledge these represent the first known transport measurements for endogenous proteins in mammalian spine.

**Figure 5.**
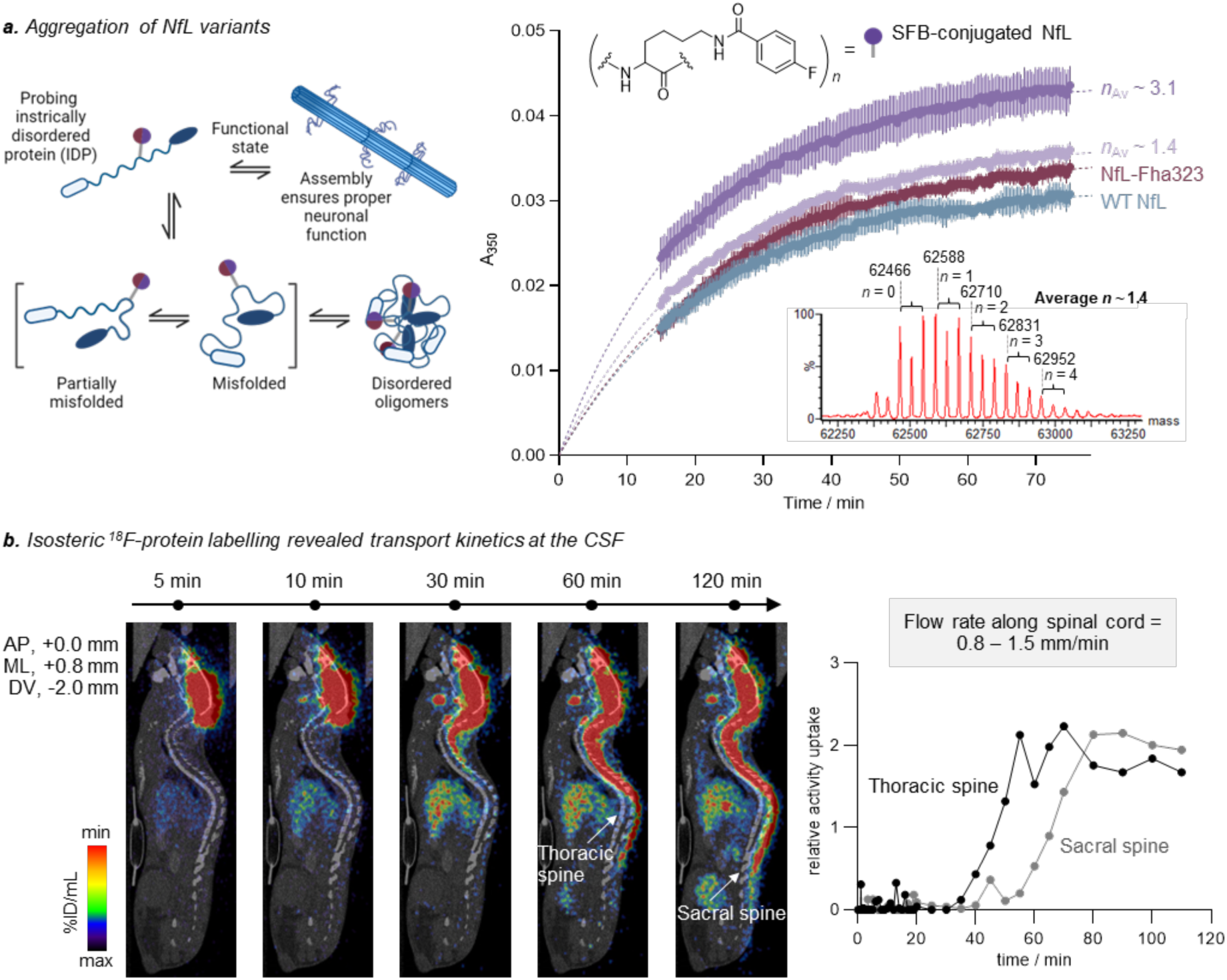
Functional and dynamic *in vivo* observations of NfL confirm mimicry and reveal associated physiological parameters enabled by precise isosteric protein ^18^F-labelling. **a.** Assembly of NfL was followed by light scattering at 350 nm (5.6 μM NfL (WT or modified), 0.23 M urea, 20 mM PIPES, 200 mM NaCl, 2 mM MgCl2, 1 mM DTT, pH 8.0, n = 5). Data were fitted to a simple aggregation model (dotted line, Supplementary Information) that accounted for dead time in measurement. NfL-Fha323 bearing bioisostere Fha (in red) demonstrated similar aggregation profiles to WT NfL (in blue) while SFB-labelled NfL heterogeneously conjugated at various stoichiometry levels (∼1.4 in light purple and ∼3.4 in dark purple) aggregated to a greater extent in the same time period, with aggregation tendency increasing in line with greater SFB loading. This altered SFB-induced behaviour confirmed the importance of Fha as a designed, site-specific edit that allows the creation a ‘near-zero-size’ label for observing proteins in as close to a native functional state as possible. This is likely to be particularly important in cases, such as here, where oligomerization states may prove critical. **b.** Intracerebroventricular administration of [^18^F]NfL enabled kinetic measurements of transport rate of protein along the spinal cord, from the thoracic spine to the sacrum.

## Discussion

Methods for the detection and characterisation of proteins that may be turned over or redistributed upon pathology into different body fluid compartments are an unmet need in biomarker development in a range of neurologic diseases. For NfL, the current standard measurement method is based on matched-pair antibodies that are necessarily indirect and that limit interpretation due to missed isoforms or fragments that cannot be captured comprehensively by this assay type.^81^ The detection of NfL in blood via these methods, therefore raises the question of the identity of the recognised analyte, its turnover and how this relates to brain pathology.

The method for direct tracking of NfL we describe here exploits the inherent sensitivity of radioemission in combination with the potential of ^18^F to act as a bioisosteric label. The precise positioning of ^18^F in proteins is rare and to our knowledge has not been previously accomplished in a bio-isosteric manner. The generation and use here now of ‘off-protein’ carbon-centred radicals bearing ^18^F has allowed the precise editing of amino acid sidechains as near-perfect radiolabels for *in vivo* PET imaging studies. Optimization of radical generation allowed use of low photocatalyst loadings, even down to sub-stoichiometric levels, and small amounts of radiolabelled precursor in a rapid, highly efficient manner with radiochemical yields of up to 67%. Automated generation of isosteric ^18^F-side-chain precursor with high molar activity allows distributable creation of ^18^F-proteins, with application not only for circulation studies (100 μL) but even for the very small volumes required for effective intracranial use (< 1 μL in small mammals). This, in turn, has enabled, to the best of our knowledge, the first dynamic tracking of NfL in the circulatory system of a whole organism in real time and of an endogenous protein in the CSF.

The relatively short half-life of intact NfL in circulation that we now reveal here, seemingly contrasts with the sustained elevation of NfL observed using other methods^8,15,16^ after acute insults to the brain parenchyma. Blood levels of NfL-associated marker can take days to weeks after injury^16,82^ to reach peak concentrations in clinical conditions such as multiple sclerosis relapses^83^ or neurosurgical intervention traumatic injury^16^ and it was not known until now whether this results from long half-life or continuous neuronal loss. The results we have obtained here now point to a dynamic equilibrium of rapid clearance that is supplied by continued transit from CSF to serum, as opposed to a long half-life of NfL. The central biological interpretation of this finding is that neuronal degeneration is therefore a continuous process in subacute phases after injury, a finding that in turn reveals that neuroprotective treatment has a clear therapeutic target. Moreover, our observed selective removal of NfL into certain organs, could suggest that the gradual increase of detectable levels of NfL with increasing age^84^ might also in part reflect reducing liver and renal function for clearance in this equilibrium.

The fast clearance of intact protein depicted by our *in vivo* results is particularly noteworthy, as this supports the concept that immunological assays designed to detect circulating levels of NfL, as biomarkers of active neurodegenerative processes, are more likely to reflect the presence of truncated NfL products, whose half-life in CSF and blood are presently unknown. A recent study has shown that the Uman-type antibodies bind to a very short epitope that in part only becomes unmasked during the course of a neuronal injury likely as a consequence of degeneration-induced proteolysis.^81^

The absence of radiolabelled NfL in the brain parenchyma when administered intracerebroventricularly or intravenously is in keeping with the known challenges large proteins face in traversing the intact blood-CSF barrier following ICV injection, and blood-brain barrier (BBB) with IV injection^85^. Our results indicate that once full-length NfL is in the CSF, it does not easily diffuse into and within the brain parenchyma. Instead, our observations suggest that physiological drainage occurs predominantly from the CSF to the deep cervical lymph nodes and to the sacral lymph nodes after a delay via the spinal cord, rather than across the blood-CSF barrier or through the cribriform plate or, as recently suggested, via the marrow in the overlying bone connected by dural channels^86^.

The drainage we observe, in some animals, of radioprotein from the brain parenchyma after IC injection strikingly contrasts with the general assumption that most large proteins leave the CNS through transcytosis across endothelial cells. It instead supports an emerging view^87^ of the glymphatic system in playing a critical role in clearing macromolecules from the CNS to peripheral lymphatics.^88,89^ Older studies, using larger injection volumes, have reported that antigen drainage from the brain parenchyma to the lymph nodes is distinct from CSF drainage, which preferentially flows into the blood^90^. By contrast, we observed the outflow of [^18^F]NfL from both the CSF and from the brain parenchyma, albeit after a delay in the latter, into the deep cervical lymph nodes. Our data, which reveal a direct path to these lymph nodes, therefore now provide clear support for suggested models in which tolerance to CNS antigens may be maintained through their drainage to and processing within the lymph nodes^91,92^ (rather than the traditional view of export to the spleen or other lymphoid structures).

Most notably, we have now also made the first estimates of transport rates of proteins within the ventricular system. It was striking that the NfL signal persisted for longer than expected, indicating a slower rate of clearance from the CSF than would be conventionally inferred based on proteins as solutes. The transport of NfL along the spinal cord, observed now through our methods, reveals a steady directional movement within the CSF, rather than within the parenchyma itself. The rate of transport – around 0.8-1.5 mm.min^-1^ – is much faster than the known active transport mechanisms for NfL traveling within neuronal axons (estimated to be around 10 mm.day^-1^).^93^ As proteins in the CSF can originate from both intrathecal synthesis and blood transudation, distinguishing their source in the past has proven problematic.^94^ This has now been made possible here by the precise creation and administration of a near-identical radioprotein variant. Our findings represent, to our knowledge, the first real-time, *in vivo* observation of protein transport along the spinal cord in mammals enabled by this ‘near-zero-size’ protein ^18^F-labelling, offering a new avenue for investigating the dynamics of CSF-based transport in health and disease.

The apparent lack of drainage from the brain parenchyma to either the overlying bone marrow of the skull or systemic circulation following IC injection also suggests a compartmentalisation of fluid dynamics within the CNS, depending on the deposition of the protein. In the case of IC injection, [^18^F]NfL remained predominantly ipsilateral, indicating limited diffusion within the parenchyma, potentially due to a combination of low interstitial fluid pressure and a lack of active clearance mechanisms. Previous studies have indicated that interstitial solutes in the brain parenchyma can be cleared via both convective and glymphatic mechanisms, but there is little previous evidence of any movement of proteins into the CSF from the parenchyma or more widespread distribution across the midline of the brain^95^. Our discovery that very low levels of radioactivity reaches peripheral organs such as the kidney and bladder also hints at much slower elimination than might have been expected, and, given the rapid half-life of full-length NfL in the blood measured here, raises questions about the fate of neurofilament proteins and/or their fragments before or after they leave the CNS. A better understanding of the nature and behaviour of the degradation products of NfL in CSF and blood therefore appears essential for correlation to each other and to clinical and radiological measures of neuronal injury, as it becomes increasingly clear that full-length NfL is unlikely to be the dominant marker contributing to the NfL-associated signals being determined by the standard antibody-based assays in clinics.^96^

This radioprotein platform therefore now sets the stage for the unambiguous mapping of the behaviour of NfL (and its metabolites) in neurodegeneration without the need for interpolation of analyte identity. This would allow, for example, the elucidation of the mechanism of protein entry and clearance from the CSF and blood and identify the significance, if any, of different NfL fragments or proteoforms in various neurodegenerative diseases. More generally, the precise trafficking of NfL mapped here *in vivo* through bioisosteric mimicry of a Cys residue by [^18^F]Fha insertion suggest a broader path to accessing fully-snative side chain radio-mimics (such as the near ubiquitous Lys) into proteins to create ‘radio-equivalent’ proteins^97^. Such proteins that would be essentially structurally and functionally indistinguishable from endogenous and that can be detected through highly sensitive methods such as PET, achieved using ‘near-zero-size’ protein ^18^F-labelling, could have wide-ranging and powerful applications within imaging and beyond.

## Author Contributions

N.Y., A.W.J.P., P.G.I., D.C.A., V.G., D.L., and B.G.D. conceptualized the project, conceived and / or designed the experiments.

A.W.J.P. scoped sulfone reagents and explored protein reactions.

A.W.J.P., J.B.I.S., J.F. and P.G.I. screened and optimised synthesis of ^18^F-monoBtSOOF and derivatives.

A.W.J.P., J.F., T.A., N.A.R. and A.J.A. automated synthesis of ^18^F-monobtSOOF on Trasis AllinOne.

A.W.J.P. designed and performed all ^18^F-protein experiments.

N.Y. designed and constructed NfL plasmid, and N.Y. and A.M.G. performed NfL expression and purification.

N.Y., A.W.J.P. and A.M.G. optimised NfL-Dha reaction with (mono)-btSOOF.

K.V.T., S.A., R.A., J.S.H., L.K., M.W.G.M., A.M.D., K.A.V. and D.C.A. performed *in vivo* mice and associated biodistribution and radiometabolite experiments. A.M.D. and D.C.A conducted i.c. and i.c.v. administration.

N.Y. performed recombinant NfL organ re-extraction experiments after IV injections. A.W.J.P., D.C.A and D.R.M analysed PET data.

A.W.J.P. and A.M.G. designed in vitro absorbance assay, and A.W.J.P., N.Y., M.D. and A.J.B. designed NMR experiments to measure aggregation kinetics.

N.L. performed LC-MS/MS experiments.

B.J. and Y.Y. constructed and expressed Histone H3 NTEV protein.

A.P. and B.G.D. wrote the paper. All authors read and commented.

## Acknowledgements

Funding: EPSRC supports the Chemistry theme at the Franklin Institute. University of Brunei Darussalam Chancellor Scholarship to A.W.J.P. Progressive MS Alliance to D.L., D.C.A. and J.K. (PMSA PA-2002-36227). Swiss National Science Foundation to J.K., D.L. We thank Dr Yibo Zeng for technical assistance in protein purification. We thank Sebastiano Ortalli for their assistance and expertise of the Trasis AllinOne platform. We thank Johan Rajander (Åbo Akademi) for production of fluorine-18 and Mira Eisala, Aake Honkaniemi and Marko Vehkanen for technical assistance during *in vivo* mouse experiments and PET imaging at Turku PET Centre, Finland.

## Extended Data Figures

**Extended Data Figure 1.**
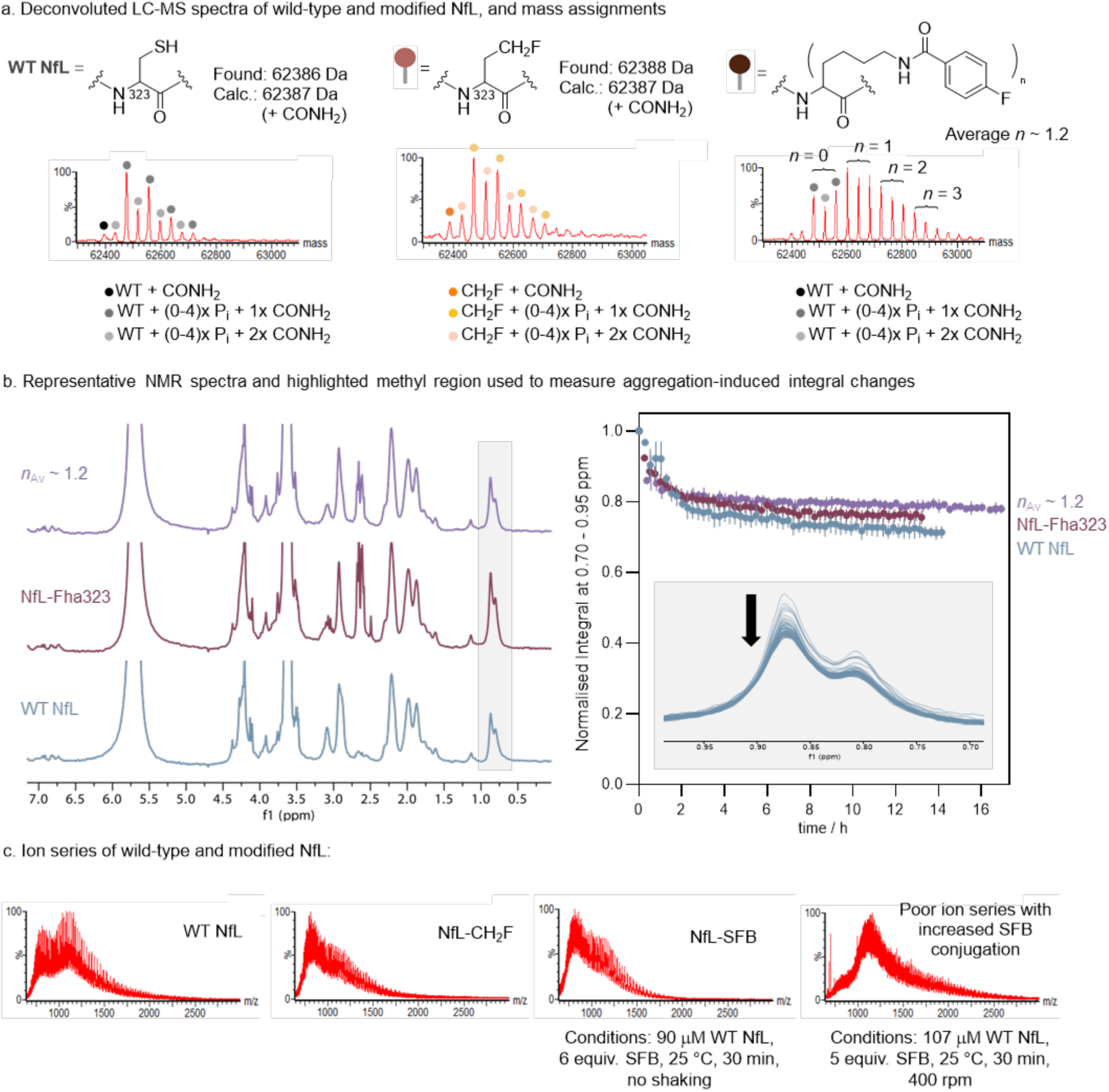
Real-time solution phase NMR spectroscopy revealed that aggregation kinetics of bioisosteric Fha-modified NfL are highly similar to wild-type. **a.** Wild-type and modified NfL variants (Fha323 and SFB-conjugated) used for NMR-based aggregation studies were characterised by MS. SFB-modified NfL was generated with an average conjugation stoichiometry of ∼1 (here n_av_ = 1.2). Mammalian cell-expressed NfL bears varying levels of phosphorylation; N-terminal carbamoylation is also observed from urea-dependent experimental conditions (see below). **b.** Aggregation of NfL and variants was followed by NMR. Integrals from all forms of NfL was detected and found to decrease with time (black arrow) reflecting the formation of higher molecular weight oligomers with relaxation rates that render them effectively undetectable. The relative behaviour of the NfL variants was compared by integrating the methyl region (0.70 to 0.95 ppm highlighted in grey). Final conditions were 46 μM NfL (WT or modified), 162 mM NaCl, 2.7 mM KCl, 8.1 mM Na_2_HPO4 and 1.5 mM KH_2_PO_4_, 0.89 mM urea, 0.44 M DTT in D_2_O, pH 7.8. Aggregation was initiated by diluting the chaotropic agent urea of a 206 μM protein sample from 4 M to 0.89 M (Supplementary Information). Similarly to the A_350_ measurements (Figure 5a), the behaviour of WT NfL (n = 2) and Fha-modified (n = 1) were similar while ∼mono-labelled SFB-conjugated NfL (n = 2) exhibited greater differences in characteristic behaviour. **c.** Notably, heterogeneously-SFB-labeled NfL proved problematic under a range of conditions, especially as conjugation stoichiometry increased. Representative intact protein MS spectra for a series of NfL variants shows that attempts to increase the extent of SFB conjugation leads to increasing deterioration in the protein-associated ion series.

**Extended Data Figure 2.**
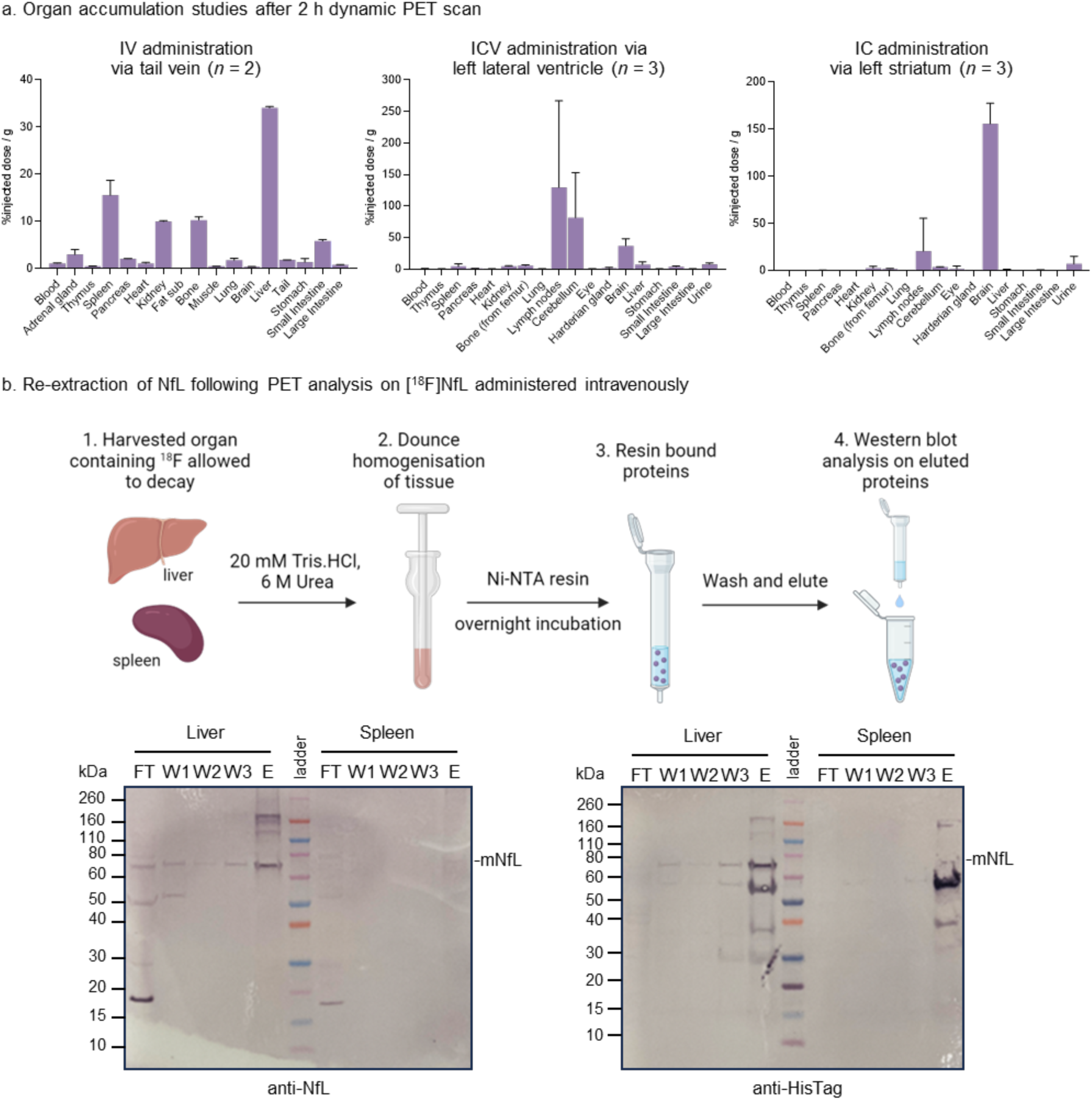
Post-PET biodistribution analysis of [^18^F]NfL. **a.** *Ex vivo* organ analysis supported observations from serial PET imaging. Specifically, the highest dose accumulation was found in the liver and spleen following direct injection into the circulatory system (IV) while [^18^F]NfL administered intracerebroventricularly (ICV) predominantly accumulated in the lymph nodes. In contrast [^18^F]NfL administered intracerebrally (IC) remained predominantly in the left striatum i.e. the site of injection. **b.** Following organ harvest and homogenisation, NfL administered intravenously via the tail vein for a 2 h PET scan could be detected in the liver and spleen via Western blot analysis.

**Extended Data Figure 3.**
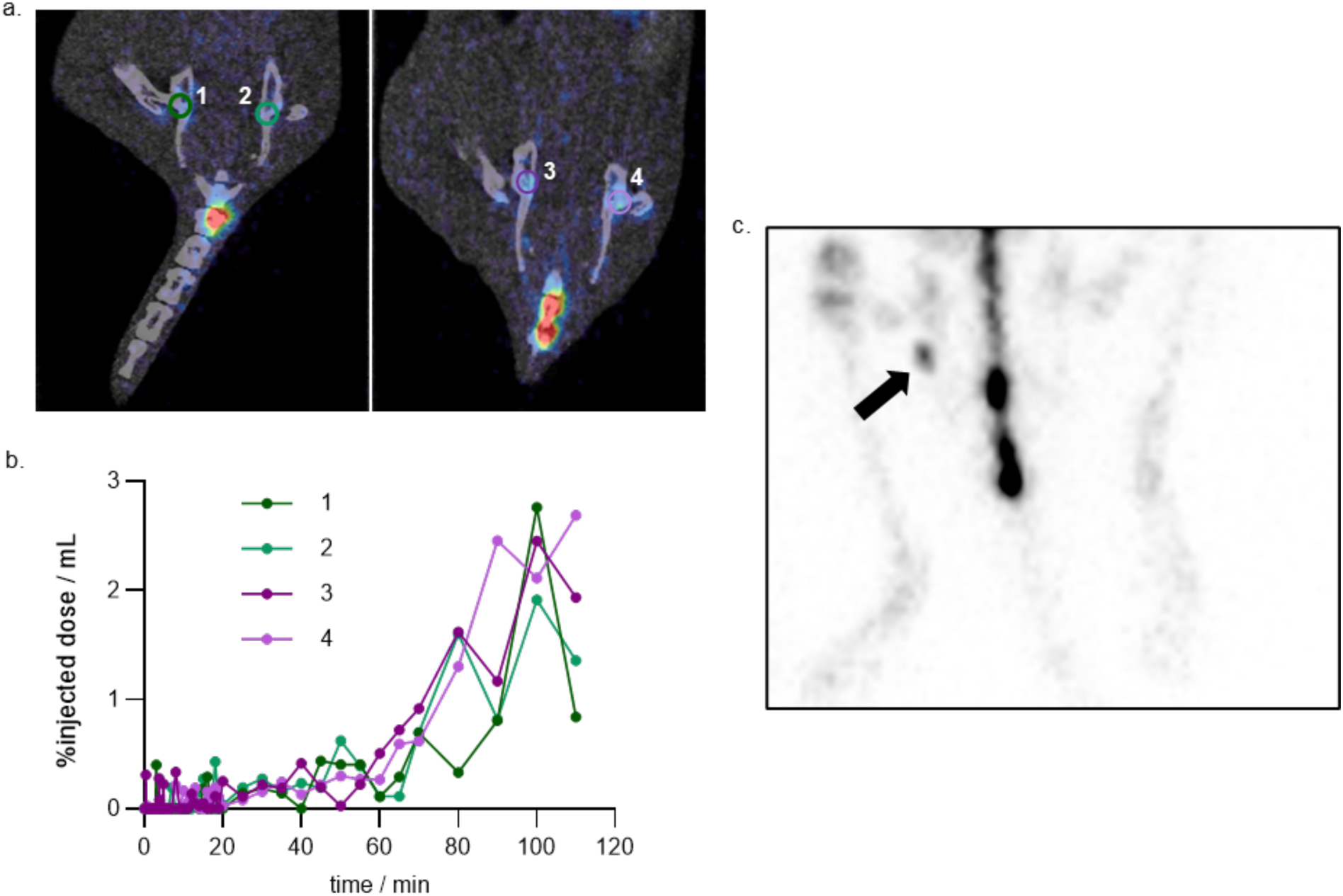
Evidence of [^18^F]NfL accumulation in the sacral lymph nodes following intracerebroventricular injection. **a.** Coronal PET/CT image (0-120 min) demonstrating mapping of [^18^F]NfL to the sacral lymph nodes. The radiotracer is transported from site of injection (lateral ventricle) in the CSF and drains into the lymphatic system. **b.** The time activity curves reveal that 40 min after injection the tracer has moved caudally along the cord and then to the sacral lymph nodes. Here, regions of interest were drawn around the lymph nodes to assess lymphatic uptake. **c.** Maximal intensity projection of the PET scan after 2 h clearly delineating the accumulation of the tracer in a lymph node. Arrows indicate the position of the lymph nodes.

