## Supplementary Information for "Isostere ^18^F-protein post-translational editing enables dynamic tracking of neurodegeneration biomarkers"

**This PDF file includes:**

Supplementary Tables

Supplementary Notes

Supplementary Methods

Supplementary References

### Table of Contents

#### Table of Contents


Supplementary Tables ---------------------------------------------------------------------------------- 7

Supplementary Table 1: Comparing the reactivity of various N-heteroaromatic sulfone radical precursors ---------------------------------------------------------------------------------- 7

Supplementary Table 2: Halex screenings on *N*-heteroaromatic sulfone precursors - 8

Supplementary Table 3: Halex screenings on 2-py–SCFHX-------------------------------- 9

Supplementary Table 4: Halex screenings on 2-pym–SCFHX ----------------------------10

Supplementary Table 5: Halex screenings on 2-bt–SCFHX -------------------------------11

Supplementary Table 6: Halex screenings on 2-bt–$SCH_2X$ -------------------------------13

Supplementary Notes ---------------------------------------------------------------------------------14

1. Double Addition: Origin and Reaction Optimisation---------------------------------------14

2. Mono-btSOOF vs btSOOF competition experiments -----------------------------------19

3. Origin of isotopic dilution of [$^{18}$F]btSOOF ------------------------------------------------20

4. NfL Phosphorylation--------------------------------------------------------------------------22

5. A comparison of commercially available antibodies against various recombinant neurofilament light chain proteins ------------------------------------------------------27

Supplementary Methods ----------------------------------------------------------------------------28

1. Small Molecule Synthesis --------------------------------------------------------------------28

1.1 General Experimental Procedures------------------------------------------------------28

1.2 Reagent synthesis ---------------------------------------------------------------------------30

2-((difluoromethyl)thio)pyridine -------------------------------------------------------------33

2-((difluoromethyl)thio)pyrimidine -----------------------------------------------------------34

2-((difluoromethyl)thio)benzothiazole -------------------------------------------------------35

2-((fluoromethyl)thio)benzothiazole----------------------------------------------------------36

2-((bromofluoromethyl)thio)pyridine --------------------------------------------------------37

2-((bromofluoromethyl)thio)pyrimidine ---------------------------------------------------38

2-((bromofluoromethyl)thio)benzothiazole ---------------------------------------------39

2-((bromomethyl)thio)benzothiazole---------------------------------------------------40

2-((difluoromethyl)sulfonyl)pyridine (S1 – pySOOF) ---------------------------------41

2-((difluoromethyl)sulfonyl)pyrimidine (S2 – pymSOOF)-----------------------------41

2-((difluoromethyl)sulfonyl)benzothiazole (S3 – btSOOF) ---------------------------42

2-(bromofluoromethyl)sulfonyl)pyridine ----------------------------------------------43

2-(bromofluoromethyl)sulfonyl)pyrimidine--------------------------------------------44

2-(bromofluoromethyl)sulfonyl)benzothiazole----------------------------------------45

Ethyl 2-(benzothiazole-2-ylthio)-2-fluoroacetate--------------------------------------46

Ethyl 2-(benzothiazole-2-ylsulfonyl)-2-fluoroacetate ---------------------------------47

2-((fluoromethyl)sulfonyl)benzothiazole (S4 – mono-btSOOF)-----------------------48

*tert*-Butyl N-(*tert*-butoxycarbonyl)-L-homoserinate (Boc-L-hSe-O*t*Bu) ----------------49

*tert*-Butyl N-(*tert*-butoxycarbonyl)-O-tosyl-L-homoserinate (Boc-L-hSe(OTs)-O*t*Bu) ---------------------------------------------------------------------------------------------50

*tert*-Butyl (S)-2-(*tert*-butoxycarbonyl)amino)-4-fluorobutanoate (Boc-L-Fha-O*t*Bu)-51

(S)-2-amino-4-fluorobutanoic acid trifluoroacetate (L-Fha•$CF_3CO_2H$) ----------------52

(S)-2-((((9H-fluoren-9-yl)methoxy)carbonyl)amino)-4-fluorobutanoic acid (Fmoc-L-Fha)-------------------------------------------------------------------------------------53

*tert*-Butyl N-(*tert*-butoxycarbonyl)-D-homoserinate (Boc-D-hSe-O*t*Bu)----------------54

*tert*-Butyl N-(*tert*-butoxycarbonyl)-O-tosyl-D-homoserinate (Boc-D-hSe(OTs)-O*t*Bu) ---------------------------------------------------------------------------------------------55

*tert*-Butyl (R)-2-(*tert*-butoxycarbonyl)amino)-4-fluorobutanoate (Boc-D-Fha-O*t*Bu) 56

(R)-2-amino-4-fluorobutanoic acid trifluoroacetate (D-Fha•$CF_3CO_2H$)----------------57

(R)-2-((((9H-fluoren-9-yl)methoxy)carbonyl)amino)-4-fluorobutanoic acid (Fmoc-D-Fha)-------------------------------------------------------------------------------------58

Diethyl 2-acetamido-2-(2,2-difluoroethyl)malonate ---------------------------------------59

2-amino-4,4-difluorobutanoic acid hydrochloride (L/D-*b*Fha•HCl) ----------------------60

2-((((9H-fluoren-9-yl)methoxy)carbonyl)amino)-4,4-difluorobutanoic acid (Fmoc-L/D-*b*Fha) ---------------------------------------------------------------------------------------------61

H-A(L-Fha)ENLYFQ-OH (L-Fha reference peptide) --------------------------------------64

H-A(L/D-*b*Fha)ENLYFQ-OH (L/D-*b*Fha reference peptide)--------------------------------65

2. Protein Production, Modification and Characterisation----------------------------------67

2.1 Intact Protein Mass Spectrometry General Methods and Data Analysis---------67

2.2 Tandem Mass Spectrometry -----------------------------------------------------------69

Histones: In-solution proteolytic digest -----------------------------------------------69

Neurofilament light chain: On-beads proteolytic digest--------------------------------70

2.3 Biological instruments---------------------------------------------------------------------71

2.4 Protein expression and purification------------------------------------------------------72

Histone H3 -----------------------------------------------------------------------------------72

Mouse Neurofilament Light Chain --------------------------------------------------------76

Human Neurofilament Light Chain--------------------------------------------------------85

Commercially available antibodies for NfL detection-----------------------------------93

LC-MS analysis of different NfL samples -----------------------------------------------94

2.5 Dha Formation -------------------------------------------------------------------------------95

Histones H3-----------------------------------------------------------------------------------95

Mouse Neurofilament Light Chain --------------------------------------------------------97

2.6 ‘Cold’ Protein Chemistry--------------------------------------------------------------------98

General protein reaction protocol --------------------------------------------------------98

Formation of *X.l.* Histone H3.NTEV-*b*Fha2 with btSOOF------------------------------99

Formation of *X.l.* Histone H3-*b*Fha9 with btSOOF------------------------------------100

Formation of *X.l.* Histone H3.NTEV-Fha2 with mono-btSOOF--------------------101

Formation of *X.l.* Histone H3-Fha27 with mono-btSOOF ---------------------------102

Formation of Human Histone eH3.1-Fha4 with mono-btSOOF -------------------103

Formation of NfL-Fha323 with mono-btSOOF -----------------------------------------104

2.7 Dephosphorylation of mNfL by λ-protein phosphatase ----------------------------106

3. Radiochemistry ---------------------------------------------------------------------------------- 107

3.1 General Methods for Radiochemistry ------------------------------------------------ 107

3.2 Manual radiosynthesis of [18F]mono-btSOOF and [18F]btSOOF using an Advion Nanotek® radiosynthesizer ------------------------------------------------------------------------ 110

3.3 Automated radiosynthesis of [18F]mono-btSOOF using a Trasis AllinOne radiosynthesizer --------------------------------------------------------------------------------- 113

3.4 Protein Radiochemistry ------------------------------------------------------------------- 117

General protein reaction protocol -------------------------------------------------------- 117

Formation of *X.l.* Histone H3.NTEV-[18F]Fha2 with [18F]mono-btSOOF --------- 118

TEV cleavage on Histone H3.NTEV-[18F]Fha2 ----------------------------------------- 120

Imine formation on Histone H3.NTEV-[18F]Fha2 in the absence of iron --------- 121

Formation of *X.l.* Histone H3.NTEV-[18F]*b*Fha2 with [18F]btSOOF ---------------- 122

TEV cleavage on Histone H3.NTEV-[18F]*b*Fha2 --------------------------------------- 123

Formation of Human Histone eH3.1-[18F]Fha4 with [18F]mono-btSOOF --------- 124

Formation of purified mNfL-[18F]Fha323 with [18F]mono-btSOOF ----------------- 126

Formation of mNfL-[18F]Fha323 for intravenous, intracerebral and intracerebroventricular administration------------------------------------------------------ 128

3.5 Recombinant mNfL re-extraction from plasma -------------------------------------- 129

mNFL re-extraction from plasma under *in vitro* condition ---------------------------- 129

mNFL re-extraction from plasma under *in vivo* condition ---------------------------- 130

3.6 PET imaging of mNfL --------------------------------------------------------------------- 132

Relative Activity Uptake ---------------------------------------------------------------------- 135

mNfL re-extraction from spleen and liver after PET scanning and 18F decay--- 136

3.7 Measuring aggregation kinetics of aggregation-prone proteins using absorbance measurements ---------------------------------------------------------------------------------- 137

Data analysis ------------------------------------------------------------------------------------ 137

Generation of SFB-conjugated NfL for aggregation studies using absorbance measurements ---------------------------------------------------------------------------------- 139

3.8 Measuring aggregation kinetics of aggregation-prone proteins with NMR Spectroscopy ---------------------------------------------------------------------------------- 141

Generation of Fha-modified NfL --------------------------------------------------------- 141

Generation of SFB-conjugated NfL ---------- 143
NfL sample preparation for NMR studies ---------- 145
Initiating NfL aggregation ---------- 145
NMR set-up and analysis ---------- 145
4. Small molecule NMR spectra ---------- 147

Supplementary References ---------- 191

### Supplementary Tables

#### Supplementary Table 1: Comparing the reactivity of various N-heteroaromatic sulfone radical precursors

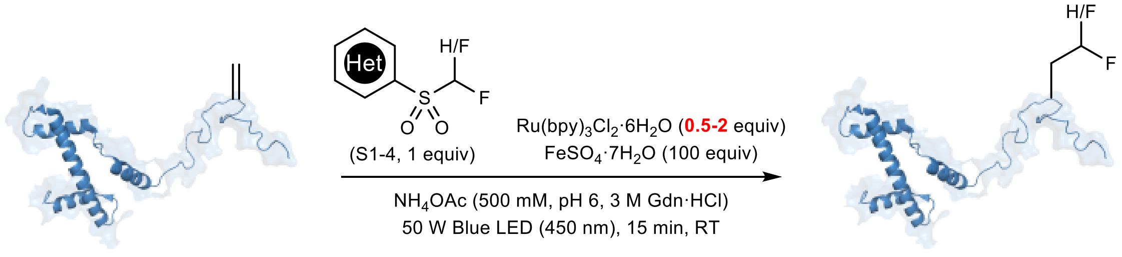


*X.l.* Histone H3-Dha9

*X.l.* Histone H3-Flb9 (S1-3) or Fha9 (S4)

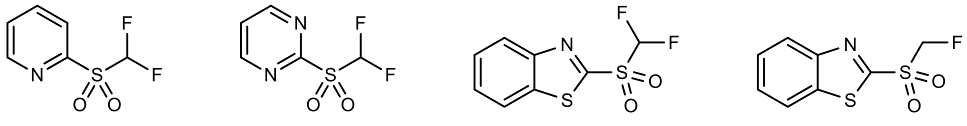

pySOOF **S1**

pymSOOF **S2**

btSOOF **S3**

mono-btSOOF **S4**

| Sulfone | $Ru(bpy)_3^{2+}$ | Starting Dha / % | Single Add. / % | Double Add. / % |
|---|---|---|---|---|
| S1 | 2 equiv | 14 | 86 | 0 |
| S2 | 2 equiv | 7 | 93 | 0 |
| S3 | 2 equiv | 17 | 71 | 12 |
| S4 | 2 equiv | 9 | 44 | 47 |
| S1 | 0.5 equiv | 58 | 42 | 0 |
| S2 | 0.5 equiv | 39 | 61 | 0 |
| S3 | 0.5 equiv | 21 | 79 | 0 |
| S4 | 0.5 equiv | 47 | 37 | 16 |

Standard reaction conditions: Samples were prepared in the glovebox using degassed solvents, 1 mg $mL^{-1}$ of *X.l.* Histone H3 Dha9, fluoroalkyl sulfones in DMSO, $Ru(bpy)_3Cl_2$•$6H_2O$ and $FeSO_4$•$7H_2O$ in $H_2O$, 100 µL total reaction volume. The samples were irradiated with blue LED (450 nm, 50 W) at room temperature and conversions were determined after 15 min.

Supplementary Table 2: Halex screenings on *N*-heteroaromatic sulfone precursors

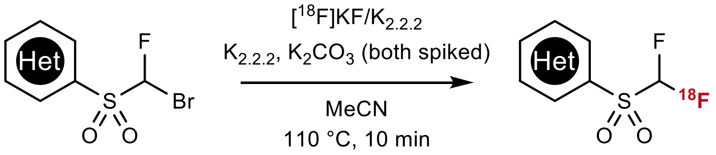


| Heterocycle | Solvent | RCY | Comments |
|---|---|---|---|
| benzothiazole | MeCN | 0% (*n* = 1) | 79% to the $S_NAr$ product. |
| pyrimidine | MeCN | 0% (*n* = 1) | 86% to $S_NAr$ product. |
| pyridine | MeCN | 7% (*n* = 1) | - |
| pyridine[a] | MeCN | 1% (*n* = 1) | - |
| pyridine | DMSO | trace (*n* = 1) | - |

[a]0.08 mmol of sulfone precursor was used.

General procedure: $[^{18}F]KF/K_{2.2.2}$ in MeCN (5 – 20 MBq, <100 µL) was dispensed into a V-vial containing Kryptofix® (0.02 mmol) and $K_2CO_3$ (0.01 mmol). The solvent was removed upon heating at 105 °C under a stream of nitrogen and left to cool. The sulfone precursor (0.04 mmol) dissolved in dry MeCN unless stated otherwise (0.5 mL) was then added and the solution was left to stir at 110 °C for 10 minutes before it was quenched with 300 µL $H_2O$. An aliquot was removed for radio-HPLC analysis for radiochemical conversion and product identity.

#### Supplementary Table 3: Halex screenings on 2-py–SCFHX

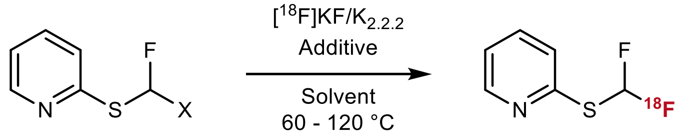


| X | Additive | Solvent | Temp. ; Time | RCY |
|---|---|---|---|---|
| Cl[a] | AgOTf (0.08 mmol) | DCE (0.3 mL) | 60 ºC; 20 min | 19 ± 2% (*n* = 2) |
| Cl | - | Acetone (1.0 mL) | 60 ºC; 20 min | trace (*n* = 2) |
| Cl | DABCO (0.04 mmol) | DMSO (1.0 mL) | 120 ºC; 5 min | trace (*n* = 2) |
| Br[b] | - | MeCN (1.0 mL) | 120 ºC; 5 min | trace (*n* = 2) |
| Br | $K_{2.2.2}$ (0.02 mmol) $K_2CO_3$ (0.01 mmol) | MeCN (1.0 mL) | 85 ºC; 10 min | 8 ± 1% (*n* = 2) |
| Br | $K_{2.2.2}$ (0.04 mmol) $K_2CO_3$ (0.02 mmol) | MeCN (1.0 mL) | 85 ºC; 10 min | 2% (*n* = 1) |
| Br[a] | AgOTf (0.08 mmol) | DCE (0.3 mL) | 60 ºC; 20 min | 3 ± 0% (*n* = 2) |
| Br | AgOTf (0.08 mmol), Pyridine (0.12 mmol) | DCE (0.3 mL) | 60 ºC; 20 min | trace (*n* = 2) |

[a,b]Adapted from previous work on the $^{18}$F-labelling of aryl thioether[1,2].

General procedure: [$^{18}$F]KF/$K_{2.2.2}$ in MeCN (5 – 20 MBq, <100 µL) was dispensed into a V-vial containing the additives stated above. The solvent was removed upon heating at 105 °C under a stream of nitrogen and left to cool. The sulfone precursor (0.04 mmol) dissolved in a given dry solvent was then added and the solution was left to stir at set temperature for a certain period of time before it was quenched with 300 µL $H_2O$ or MeCN/$H_2O$ (9:1). An aliquot was removed for radio-HPLC analysis for radiochemical conversion and product identity.

Supplementary Table 4: Halex screenings on 2-pym–SCFHX

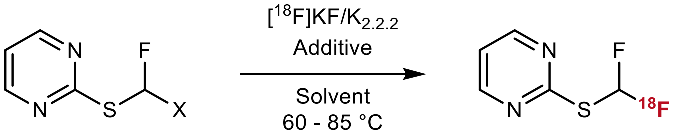

| X | Additive | Solvent | Temp. ; Time | RCY |
|---|---|---|---|---|
| Br | - | MeCN (1.0 mL) | 85 ºC; 10 min | 2 ± 0% (*n* = 2) |
| Br | $K_{2.2.2}$ (0.02 mmol) $K_2CO_3$ (0.01 mmol) | MeCN (1.0 mL) | 85 ºC; 10 min | trace (*n* = 2) |
| Br | AgOTf (0.08 mmol) | DCE (0.3 mL) | 60 ºC; 20 min | 17 ± 2% (*n* = 2) |

General procedure: $[^{18}F]KF/K_{2.2.2}$ in MeCN (5 – 20 MBq, <100 µL) was dispensed into a V-vial containing the additives stated above. The solvent was removed upon heating at 105 °C under a stream of nitrogen and left to cool. The sulfone precursor (0.04 mmol) dissolved in a given dry solvent was then added and the solution was left to stir at set temperature for a certain period of time before it was quenched with 300 µL $H_2O$ or $MeCN/H_2O$ (9:1). An aliquot was removed for radio-HPLC analysis for radiochemical conversion and product identity.

#### Supplementary Table 5: Halex screenings on 2-bt–SCFHX

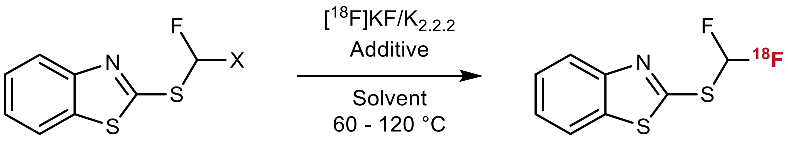

| X | Additive | Solvent | Temp. ; Time | RCY |
|---|---|---|---|---|
| Br | - | MeCN (1.0 mL) | 85 ºC; 10 min | 6 ± 0% ($n$ = 2) |
| Br | $K_{2.2.2}$ (0.02 mmol) $K_2CO_3$ (0.01 mmol) | MeCN (1.0 mL) | 85 ºC; 10 min | 35 ± 18% ($n$ = 2) |
| Br | $K_{2.2.2}$ (0.02 mmol) $K_2CO_3$ (0.01 mmol) | MeCN (0.3 mL) | 85 ºC; 10 min | 38% ($n$ = 1) |
| Br | $K_{2.2.2}$ (0.02 mmol) $K_2C_2O_4$ (0.01 mmol) | MeCN (1.0 mL) | 85 ºC; 10 min | 3 ± 2% ($n$ = 2) |
| Br | $K_2CO_3$ (0.01 mmol) | MeCN (1.0 mL) | 85 ºC; 10 min | 1 ± 1% ($n$ = 3) |
| Br | $K_{2.2.2}$ (0.02 mmol) | MeCN (1.0 mL) | 85 ºC; 10 min | trace ($n$ = 1) |
| Br | $K_2C_2O_4$ (0.01 mmol) | MeCN (1.0 mL) | 85 ºC; 10 min | trace ($n$ = 2) |
| Br | $K_{2.2.2}$ (0.02 mmol) $K_2CO_3$ (0.01 mmol) | PhCN (1.0 mL) | 120 ºC; 10 min | 17% ($n$ = 1) |
| Br | $K_{2.2.2}$ (0.02 mmol) $K_2CO_3$ (0.01 mmol) | *tert*-amyl alcohol (1.0 mL) | 85 ºC; 10 min | trace ($n$ = 1) |
| Br[b] | $K_{2.2.2}$ (0.02 mmol) $K_2CO_3$ (0.01 mmol) | *tert*-amyl alcohol: MeCN (1:1; 1.0 mL) | 85 ºC; 10 min | 42% ($n$ = 1) |
| Br[a] | Tetraglycol (0.02 mmol) $K_2CO_3$ (0.01 mmol) | MeCN (1.0 mL) | 85 ºC; 10 min | 41% ($n$ = 1) |
| Br[a] | Tetraglycol (0.01 mmol) $K_2CO_3$ (0.01 mmol) | MeCN (1.0 mL) | 85 ºC; 10 min | 23% ($n$ = 1) |

| | | | | |
|---|---|---|---|---|
| Br | AgOTf (0.08 mmol) | DCE (0.3 mL) | 60 ºC; 20 min | 11 ± 1% ($n$ = 2) |
| OTs[c] | $K_{2.2.2}$ (0.02 mmol) $K_2CO_3$ (0.01 mmol) | MeCN (1.0 mL) | 85 ºC; 10 min | trace ($n$ = 1) |
| OTs | $K_{2.2.2}$ (0.02 mmol) $K_2CO_3$ (0.01 mmol) | PhCN (1.0 mL) | 120 ºC; 10 min | 0% ($n$ = 1) |
| OTs[c] | $K_{2.2.2}$ (0.02 mmol) $K_2CO_3$ (0.01 mmol) | *tert*-amyl alcohol: MeCN (1:1; 1.0 mL) | 85 ºC; 10 min | trace ($n$ = 1) |

[a]Impurities formed in 30%. [b]Impurities formed in 7%. [c]85-97% RCY to [$^{18}$F]tosyl fluoride.

General procedure: [$^{18}$F]KF/$K_{2.2.2}$ in MeCN (5 – 20 MBq, <100 µL) was dispensed into a V-vial containing the additives stated above. The solvent was removed upon heating at 105 °C under a stream of nitrogen and left to cool. The sulfone precursor (0.04 mmol) dissolved in a given dry solvent was then added and the solution was left to stir at set temperature for a certain period of time before it was quenched with 300 µL $H_2O$ or MeCN/$H_2O$ (9:1). An aliquot was removed for radio-HPLC analysis for radiochemical conversion and product identity.

#### Supplementary Table 6: Halex screenings on 2-bt–$SCH_2X$

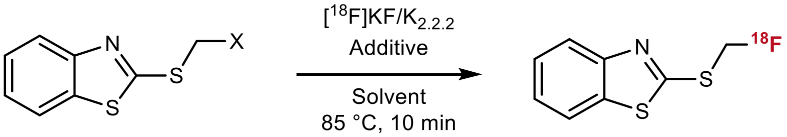


| X | Additive | Solvent | Temp. ; Time | RCY |
|---|---|---|---|---|
| Br | $K_{2.2.2}$ (0.02 mmol) $K_2CO_3$ (0.01 mmol) | MeCN (1.0 mL) | 85 ºC; 10 min | 48 ± 7% (*n* = 2) |
| Br | $K_{2.2.2}$ (0.02 mmol) $K_2CO_3$ (0.01 mmol) | *tert*-amyl alcohol: MeCN (1:1; 1.0 mL) | 85 ºC; 10 min | 38% (*n* = 1) |
| OTs[a] | $K_{2.2.2}$ (0.02 mmol) $K_2CO_3$ (0.01 mmol) | MeCN (1.0 mL) | 85 ºC; 10 min | 62% (*n* = 1) |
| OTs[a] | $K_{2.2.2}$ (0.02 mmol) $K_2CO_3$ (0.01 mmol) | *tert*-amyl alcohol: MeCN (1:1; 1.0 mL) | 85 ºC; 10 min | 51% (*n* = 1) |

[a]Impurities formed in 3%. Subsequent oxidation to form [$^{18}$F]mono-btSOOF proceeded with a RCY of 60% (compared to ~98% oxidation following halex on the brominated precursor). In addition, the impurity from the halex reaction co-eluted with [$^{18}$F]mono-btSOOF.

### Supplementary Notes

#### 1. Double Addition: Origin and Reaction Optimisation

Significant double addition side product was observed when mono-btSOOF was employed as the radical precursor under standard optimised conditions for radical reactions on Dha using N-heteroaromatic sulfones (125 µM protein concentration, 0.5 – 2 equiv. photocatalyst, 100 equiv. Fe(II) additive, 50 W, 15 min) . Notably, the addition of an electron-withdrawing group on the heteroaryl ring[3] or on the C-centred radical[4] was not required for the formation of the desired fluorohomoalanine (Fha)-modified histone (i.e., 37% conversion to H3-Fha9 as demonstrated in Figure 2). The levels of double addition varies depending on protein and Dha site (19% to H3-$Fha_2$9). Proteomic analysis confirmed this competing dialkylation event on the Dha site. To circumvent this side reaction, we reduced light flux (from 50 W to 10 – 15 W) and increased reaction times (by a further 15 to 30 min) to increase conversion to the desired single addition modification. Under re-optimised conditions, low to negligible amount of double addition formation were observed (for H3-Fha27, up to 71% conversion with negligible double addition side product).

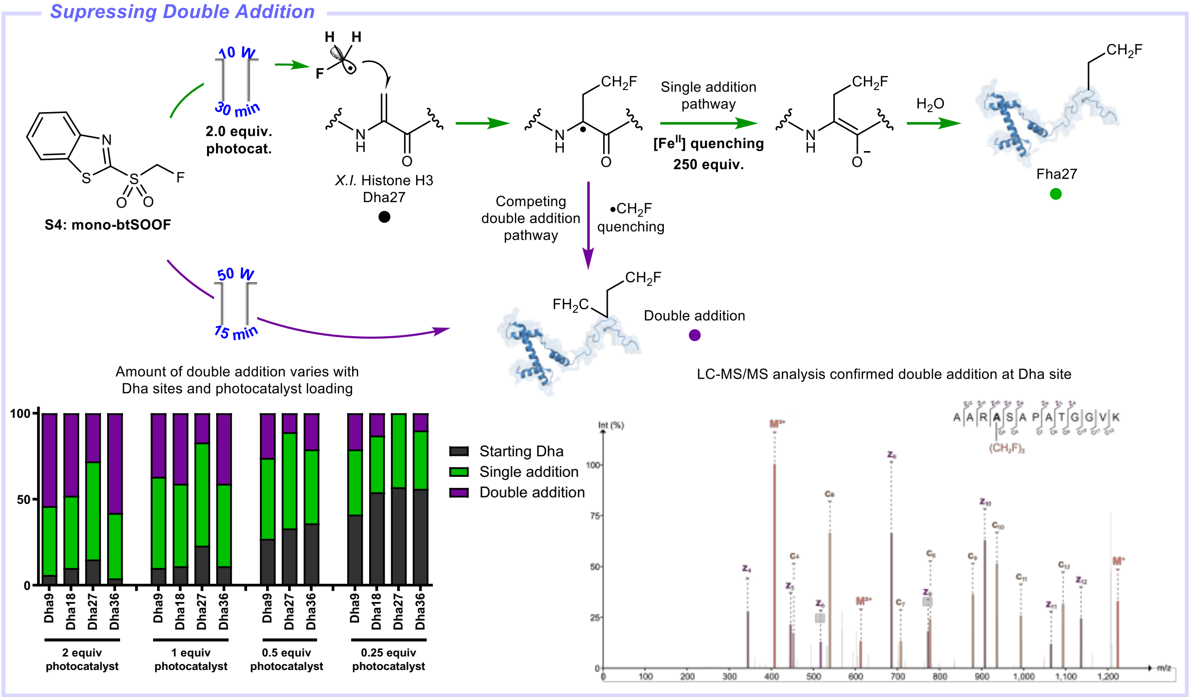

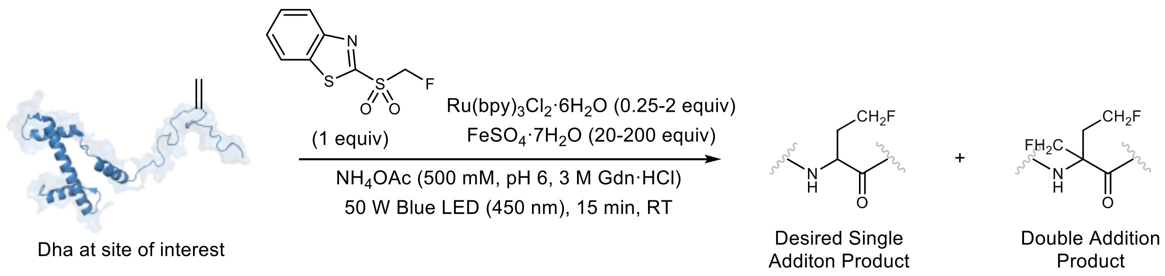


*X.I.* Histone H3:

| Dha site | $Ru(bpy)_3^{2+}$ (equiv.) | $Fe^{2+}$ (equiv.) | Starting Dha / % | Single Add. / % | Double Add. / % |
|---|---|---|---|---|---|
| 9 | 2 | 100 | 6 | 40 | 54 |
| 18 | 2 | 100 | 10 | 42 | 48 |
| 27 | 2 | 100 | 15 | 57 | 28 |
| 36 | 2 | 100 | 4 | 38 | 58 |
| 9 | 1 | 100 | 10 | 53 | 37 |
| 18 | 1 | 100 | 11 | 48 | 41 |
| 27 | 1 | 100 | 23 | 60 | 17 |
| 36 | 1 | 100 | 11 | 48 | 41 |
| 9 | 0.5 | 100 | 27 | 47 | 26 |
| 27 | 0.5 | 100 | 33 | 56 | 11 |
| 36 | 0.5 | 100 | 36 | 43 | 21 |
| 9 | 0.25 | 100 | 41 | 38 | 21 |
| 18 | 0.25 | 100 | 54 | 33 | 13 |
| 27 | 0.25 | 100 | 57 | 43 | 0 |
| 36 | 0.25 | 100 | 56 | 34 | 10 |
| 18 | 2 | 200 | 10 | 45 | 45 |
| 18 | 2 | 100 | 11 | 40 | 49 |
| 18 | 2 | 50 | 13 | 37 | 50 |
| 18 | 2 | 20 | 13 | 34 | 53 |
| 27 | 2 | 200 | 25 | 56 | 19 |
| 27 | 2 | 100 | 29 | 51 | 19 |
| 27 | 2 | 50 | 39 | 44 | 17 |
| 27 | 2 | 20 | 48 | 36 | 16 |

Human Histone eH3:

| Dha site | $Ru(bpy)_3^{2+}$ (equiv.) | $Fe^{2+}$ (equiv.) | Starting Dha / % | Single Add. / % | Double Add. / % |
|---|---|---|---|---|---|
| 4 | 2 equiv | 250 equiv | (<1) | 49 | 51 |
| 4 | 2 equiv | 100 equiv | 6 | 46 | 48 |
| 4 | 2 equiv | 50 equiv | 15 | 57 | 28 |
| 9 | 2 equiv | 250 equiv | 9 | 69 | 22 |
| 9 | 2 equiv | 100 equiv | 9 | 69 | 22 |
| 9 | 2 equiv | 50 equiv | 9 | 63 | 28 |
| 18 | 2 equiv | 250 equiv | (<1) | 44 | 56 |
| 18 | 2 equiv | 100 equiv | (<1) | 38 | 62 |
| 18 | 2 equiv | 50 equiv | (<1) | 37 | 63 |

*X.l.* Histone H3.NTEV:

| Dha site | $Ru(bpy)_3^{2+}$ (equiv.) | $Fe^{2+}$ (equiv.) | Starting Dha / % | Single Add. / % | Double Add. / % |
|---|---|---|---|---|---|
| 2 | 2 equiv | 100 equiv | 10 | 90 | 0 |
| 2 | 1 equiv | 100 equiv | 10 | 90 | 0 |
| 2 | 0.5 equiv | 100 equiv | 9 | 91 | 0 |
| 2 | 0.25 equiv | 100 equiv | 20 | 80 | 0 |

Standard reaction conditions: Samples were prepared in the glovebox using degassed solvents, 1 mg $mL^{-1}$ Dha protein concentration in $NH_4OAc$ (500 mM, pH 6, 3 M Gdn•HCl), fluoroalkyl sulfones in DMSO, $Ru(bpy)_3Cl_2$•$6H_2O$ and $FeSO_4$•$7H_2O$ in $H_2O$, 100 µL total reaction volume. The samples were irradiated with blue LED (450 nm, 50 W) at room temperature and conversions were determined after 15 min.

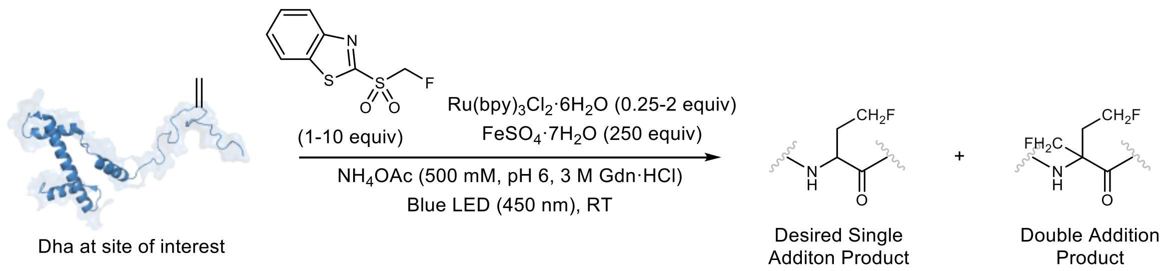


*X.I.* Histone H3:

| Dha site | mono-btSOOF | $Ru(bpy)_3^{2+}$ | Condition | Starting Dha / % | Single Add. / % | Double Add. / % |
|---|---|---|---|---|---|---|
| 36 | 1 equiv. | 1 equiv. | 10 W, 15 min | 43 | 43 | 14 |
| 36 | 1 equiv. | 1 equiv. | 10 W, 30 min | 21 | 58 | 21 |
| 36 | 10 equiv. | 0.5 equiv. | 10 W, 30 min | 47 | 43 | 10 |
| 36 | 10 equiv. | 0.5 equiv. | 10 W, 45 min | 35 | 53 | 12 |
| 36 | 10 equiv. | 1 equiv. | 10 W, 15 min | 44 | 41 | 15 |
| 36 | 10 equiv. | 1 equiv. | 10 W, 30 min | 21 | 59 | 20 |
| 36 | 1 equiv. | 1 equiv. | 7 W, 30 min | 50 | 41 | 9 |
| 36 | 1 equiv. | 2 equiv. | 7 W, 30 min | 38 | 49 | 13 |
| 36 | 1 equiv. | 0.5 equiv. | 8.5 W, 30 min | 63 | 37 | <1 |
| 36 | 1 equiv. | 0.5 equiv. | 8.5 W, 45 min | 57 | 43 | <1 |
| 36 | 1 equiv. | 0.5 equiv. | 15 W, 30 min | 21 | 64 | 15 |
| 36 | 1 equiv. | 0.5 equiv. | 15 W, 45 min | 16 | 69 | 15 |
| 27 | 1 equiv. | 2 equiv. | 10 W, 15 min | 32 | 59 | 9 |

| | | | | | | |
|---|---|---|---|---|---|---|
| 27 | 1 equiv. | 2 equiv. | 10 W, 30 min | 17 | 71 | 12 |
| 27 | 1 equiv. | 1 equiv. | 10 W, 30 min | 30 | 70 | <1 |

Human Histone eH3:

| Dha site | mono-btSOOF | $Ru(bpy)_3^{2+}$ | Condition | Starting Dha / % | Single Add. / % | Double Add. / % |
|---|---|---|---|---|---|---|
| 4 | 1 equiv. | 1 equiv. | 10 W, 30 min | 30 | 56 | 14 |
| 4 | 1 equiv. | 1 equiv. | 10 W, 45 min | 10 | 77 | 13 |
| 4 | 1 equiv. | 0.5 equiv. | 15 W, 30 min | 31 | 56 | 13 |
| 4 | 1 equiv. | 0.5 equiv. | 15 W, 45 min | 16 | 69 | 15 |

Standard reaction conditions: Samples were prepared in the glovebox using degassed solvents, 1 mg $mL^{-1}$ Dha protein concentration in $NH_4OAc$ (500 mM, pH 6, 3 M Gdn•HCl), mono-btSOOF in DMSO, $Ru(bpy)_3Cl_2$•$6H_2O$ and $FeSO_4$•$7H_2O$ in $H_2O$, 100 μL total reaction volume. The samples were irradiated with blue LED (450 nm) and varying light flux at room temperature and conversions were determined after specified times.

#### 2. Mono-btSOOF vs btSOOF competition experiments

Competition experiments suggest that increased fluorine substituents on the γ-carbon side chain facilitates efficient quenching of the α-carbon radical by Fe(II), leading to the predominant formation of *b*Fha-modified histone even when mono-btSOOF was employed in far excess. As Fe(II) is a less effective quencher of the on-protein radical intermediate following the addition of $\bullet CH_2F$, controlled release of this radical (for careful modulation of its effective concentration) appeared to be significant to favour high conversion to the desired single addition product.

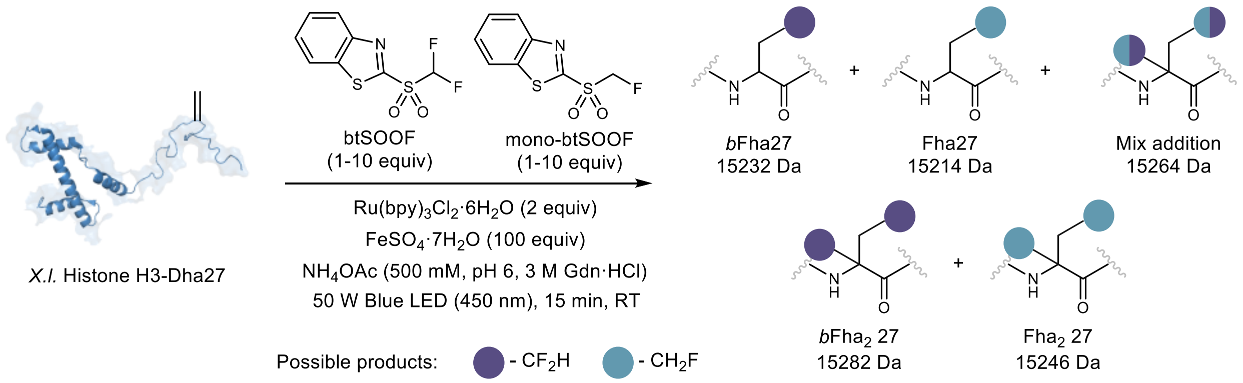


| btSOOF | mono-btSOOF | Starting Dha | *b*Fha | Fha | Mix addition | *b*$Fha_2$ | $Fha_2$ |
|---|---|---|---|---|---|---|---|
| 2 equiv. | 2 equiv. | <1% | 66% | 0% | 9% | 16% | 9% |
| 5 equiv. | 2 equiv. | <1% | 74% | 0% | 0% | 26% | 0% |
| 2 equiv. | 5 equiv. | <1% | 74% | 0% | 13% | 13% | 0% |
| 1 equiv. | 10 equiv. | <1% | 58% | 11% | 13% | 5% | 13% |
| 2 equiv. | 10 equiv. | <1% | 63% | 7% | 12% | 9% | 10% |

Standard reaction conditions: Samples were prepared in the glovebox using degassed solvents, 1 mg $mL^{-1}$ Dha protein concentration in $NH_4OAc$ (500 mM, pH 6, 3 M Gdn•HCl), fluoroalkyl sulfones in DMSO, $Ru(bpy)_3Cl_2{\bullet}6H_2O$ and $FeSO_4{\bullet}7H_2O$ in $H_2O$, 100 μL total reaction volume. The samples were irradiated with blue LED (450 nm, 50 W) at room temperature and conversions were determined after 15 min.

#### 3. Origin of isotopic dilution of [$^{18}$F]btSOOF

Quantitative $^{19}$F-NMR studies revealed the competing decomposition of the halex precursor 2-bt–SCFHBr under the $^{18}$F-halex conditions imposed, resulting in the release of $^{19}$F-fluoride and subsequently, the formation of the non-radioactive 2-btSCF$_2$H, even in the absence of an external fluoride reagent. This isotopic dilution event reduces the molar activity of the $^{18}$F-labelled analogue.

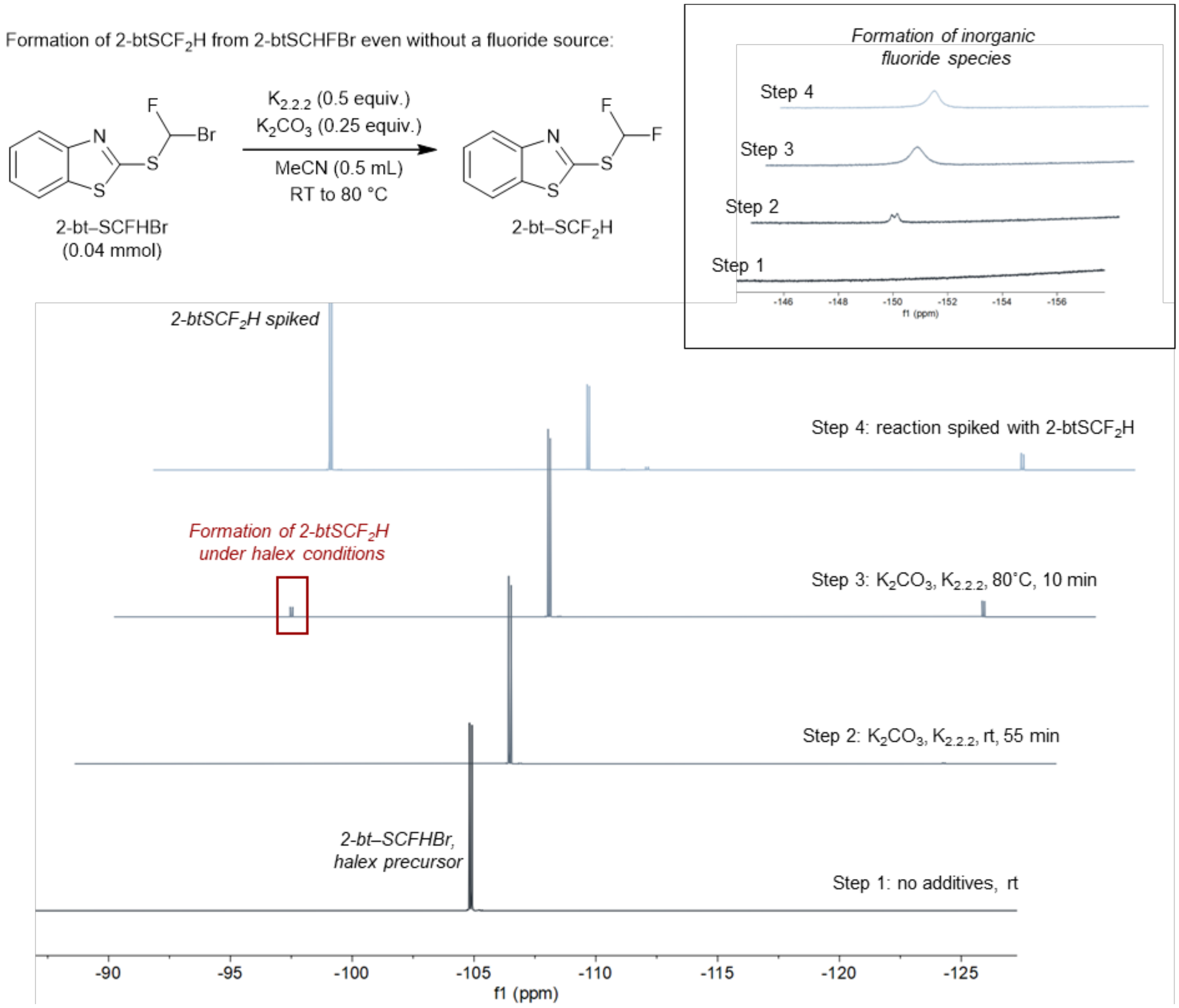


Procedure: To an NMR tube was added 2-bt–SCFHBr (11.1 mg, 0.04 mmol) in MeCN-d$_3$ (0.5 mL). The sample was submitted for NMR analysis (d1 = 30 s, n = 32 scans). $K_2CO_3$ (1.4 mg, 0.01 mmol) and $K_{2.2.2}$ (7.5 mg, 0.01 mmol) were added and the reaction was left at rt for 10 min before $^{19}$F NMR analysis. The mixture was left for a further 45 min (total time 55 min) before heating the sealed tube at 80 °C for 10 min. The reaction was cooled in ice and its $^{19}$F NMR spectrum was taken. Heating was again performed for a further 10 min (total time 20 min) for NMR analysis before finally spiking the reaction mixture with 2-btSCF$_2$H.

As a consequence of this isotopic dilution event resulting in lower molar activity for [$^{18}$F]btSOOF compared to [$^{18}$F]mono-btSOOF, LC-MS analysis of the decayed $^{18}$F-labelled protein when [$^{18}$F]btSOOF was employed as the radical precursor showed formation of the non-radioactive *b*Fha-modified protein. On the other hand, only a single mass corresponding to the Dha starting material was observed when [$^{18}$F]mono-btSOOF was employed as the protein $^{18}$F-labelling reagent.

ESI-MS analysis on Histones H3.NTEV-[$^{18}$F]Fha2 (top) and H3.NTEV-[$^{18}$F]*b*Fha2 (bottom) (after complete $^{18}$F decay):

Calculated H3 Dha mass: 16004 Da

Observed mass: 16003 Da

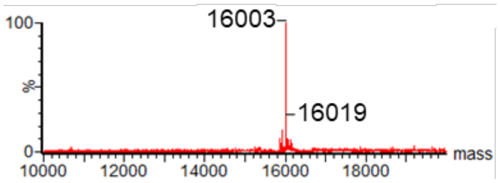


Calculated H3 *b*Fha mass: 16056 Da

Observed mass: 16054 Da

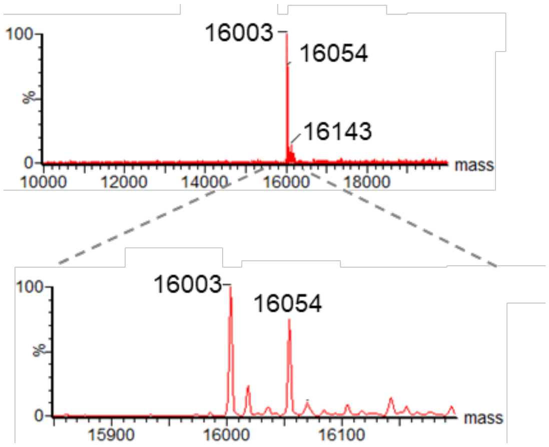

#### 4. NfL Phosphorylation

Multiple peaks observed in the LC-MS spectra of mouse NfL (mNfL) and human NfL (hNfL) expressed in mammalian cells were consistent with sequential phosphorylation. Treatment of the purified NfL proteins with λ-phosphatase removed the phosphoryl groups (-$PO_3$) attached to serine, tyrosine and threonine residues. After treatment, a single peak was observed in the LC-MS spectra for both mNfL and hNfL, confirming phosphorylation. Pro-Q diamond phosphoprotein staining (ThermoFisher) also identified phosphorylation on mNfL and hNfL.

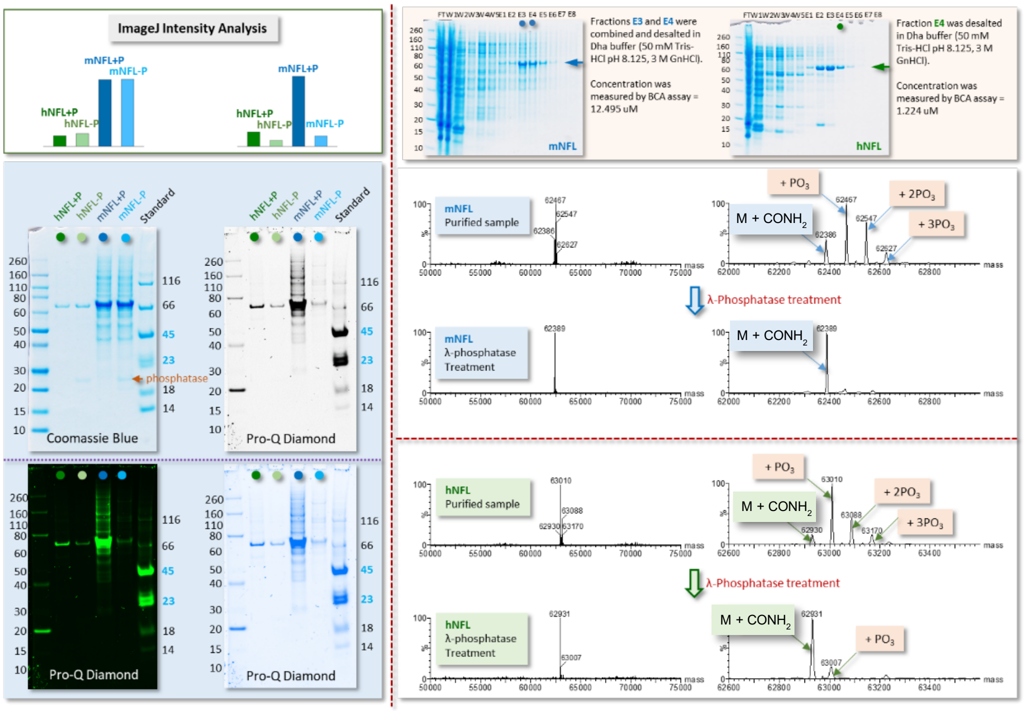

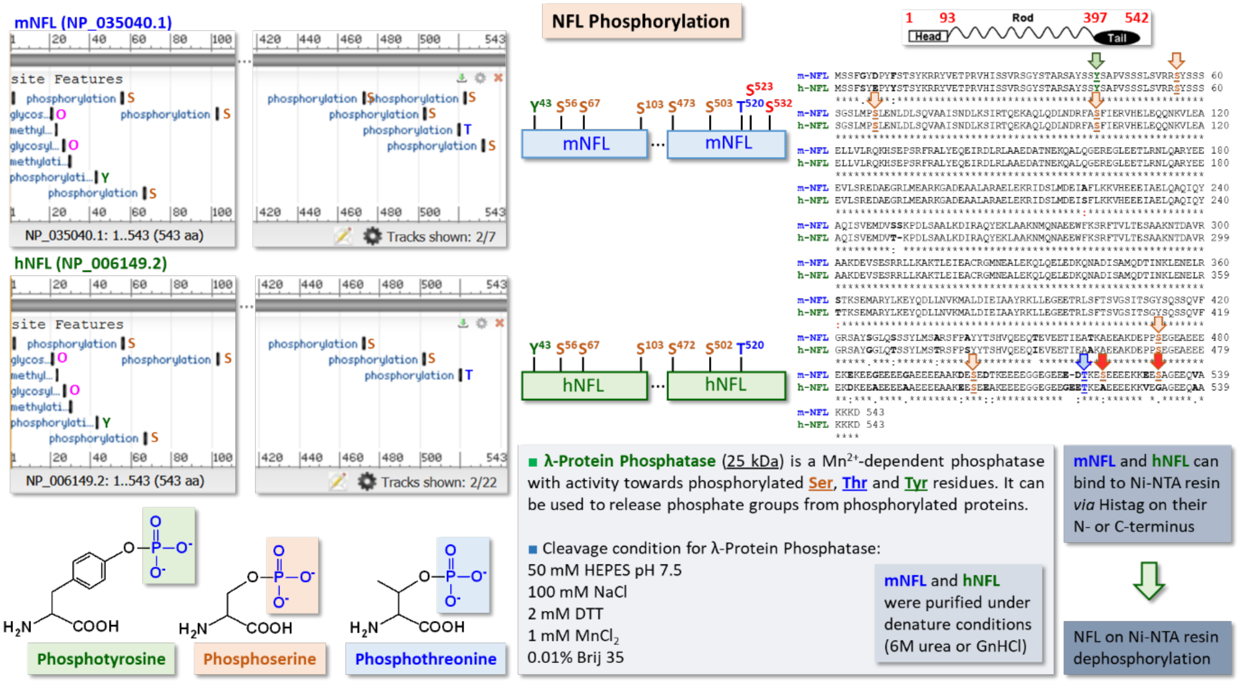


Sequence alignment of mNfL (NCBI: NP_035040.1) and hNfL (NCBI: NP_006149.2) to identify the protein phosphorylation sites. Based on the NCBI database, the phosphorylation sites on mNfL and hNfL are highly conserved except that phosphorylation of S523 and S532 are only observed in mNFL.

Peptide mapping of mNfL (sequence coverage in green) was identified through tandem LC-MS analysis of its tryptic digest. Highlighted in red are sites of phosphorylation found by HCD (high energy collision induced dissociation) and ETciD (electron transfer, collision induced dissociation) on pSer473, pSer532, pSer503 and occasionally, pSer523.

```
        10         20         30         40         50         60
MSSFGYDPYF STSYKRRYVE TPRVHISSVR SGYSTARSAY SSYSAPVSSS LSVRRSYSSS

        70         80         90        100        110        120
SGSLMPSLEN LDLSQVAAIS NDLKSIRTQE KAQLQDLNDR FASFIERVHE LEQQNKVLEA

       130        140        150        160        170        180
ELLVLRQKHS EPSRFRALYE QEIRDLRLAA EDATNEKQAL QGEREGLEET LRNLQARYEE

       190        200        210        220        230        240
EVLSREDAEG RLMEARKGAD EAALARAELE KRIDSLMDEI AFLKKVHEEE IAELQAQIQY

       250        260        270        280        290        300
AQISVEMDVS SKPDLSAALK DIRAQYEKLA AKNMQNAEEW FKSRFTVLTE SAAKNTDAVR
```

```
       310        320        330        340        350        360
AAKDEVSESR RLLKAKTLEI EACRGMNEAL EKQLQELEDK QNADISAMQD TINKLENELR

       370        380        390        400        410        420
STKSEMARYL KEYQDLLNVK MALDIEIAAY RKLLEGEETR LSFTSVGSIT SGYSQSSQVF

       430        440        450        460        470        480
GRSAYSGLQS SSYLMSARSF PAYYTSHVQE EQTEVEETIE ATKAEEAKDE PPSEGEAEEE

       490        500        510        520        530        540
EKEKEEGEEE EGAEEEEAAK DESEDTKEEE EGGEGEEEDT KESEEEEKKE ESAGEEQVAK

       550
KKDGSHHHHH H
```

pSer473:

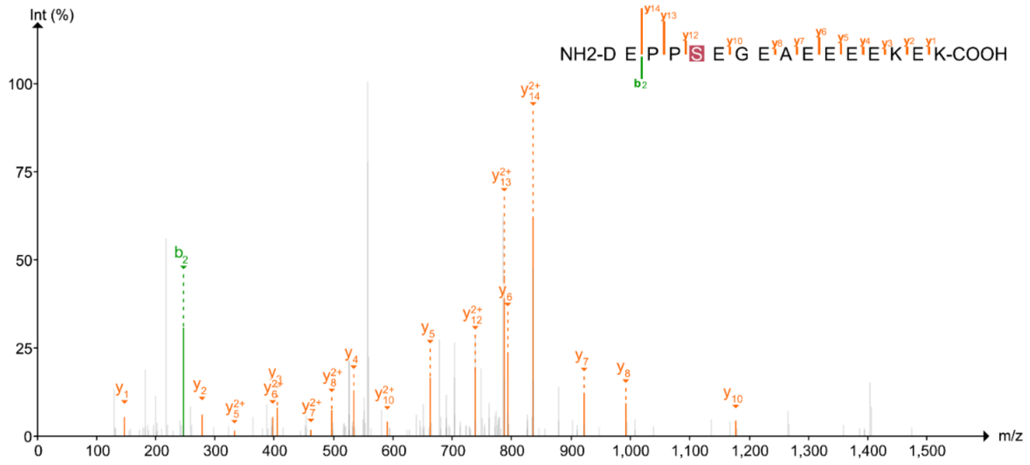


HCD mass spectrum of [DEPPpSEGEAEEEEKEK]$^{3+}$ acquired in the tandem LC-MS analysis of the tryptic digest of mNfL.

pSer503:

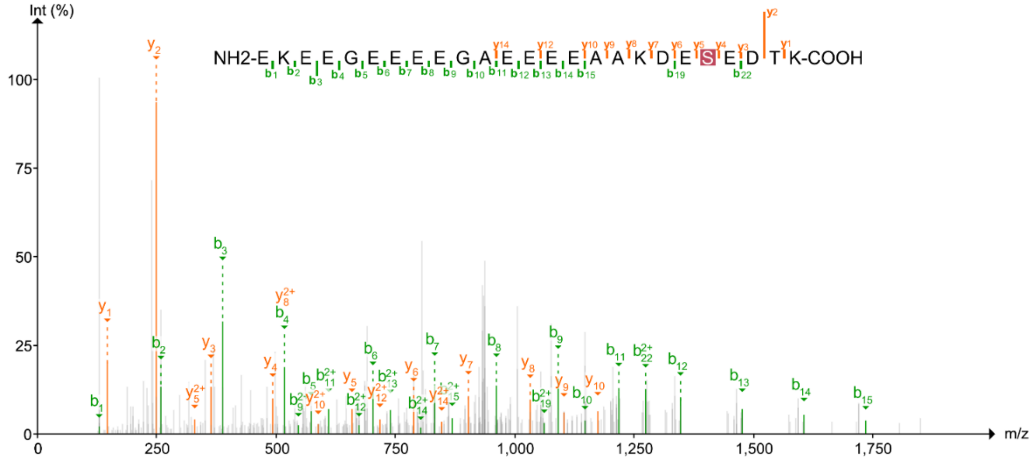


HCD mass spectrum of [EKEEGEEEEGAEEEEAAKDEpSEDTK]$^{3+}$ acquired in the tandem LC-MS analysis of the tryptic digest of *mNfL.*

pSer532:

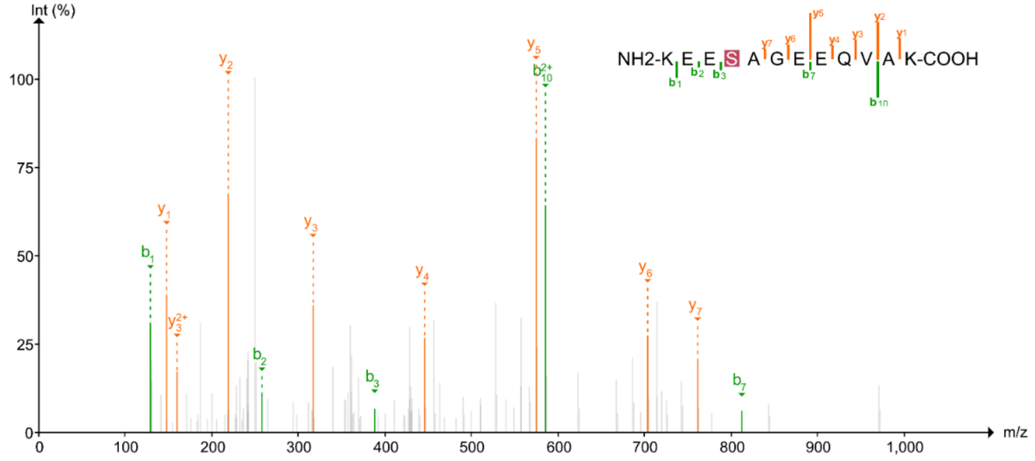


HCD mass spectrum of [KEEpSAGEEQVAK]$^{3+}$ acquired in the tandem LC-MS analysis of the tryptic digest of mNfL.

pSer523:

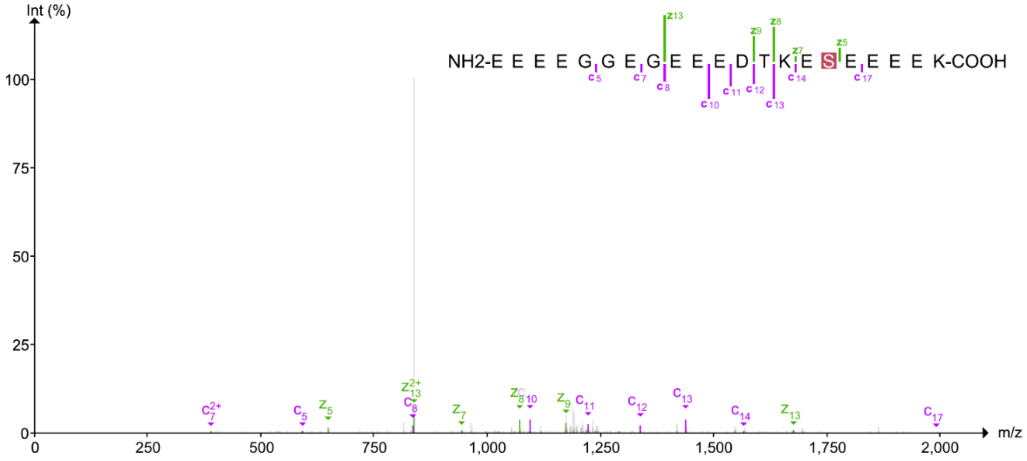


ETciD mass spectrum of $[EEEEGGEGEEEDTKEpSEEEEK]^{3+}$ acquired in the tandem LC-MS analysis of the tryptic digest of mNfL.

5. A comparison of commercially available antibodies against various recombinant neurofilament light chain proteins

Gel electrophoresis and Westerns were performed against various recombinant neurofilament light chain proteins: mouse NfL and human NfL (abbreviated as Ox-mNfL and Ox-hNfL respectively) expressed in-house in mammalian systems, and a commercially available human NfL expressed in an *E. coli* system (abbreviated as En-hNfL; EnCor Biotechnology Inc.; Cat# PROT-r-NF-L). In all cases, the proteins have been purified using affinity columns. For Westerns, various primary antibodies were tested which included UD1 (Uman Diag 27016), UD3 (Uman Diag 27018), DA2 (Novus NB300-132) and *anti*-NFL (GeneTex GTX134085) antibodies. Among these antibodies, the epitope of UD1, UD3 and DA2 is the rod domain of NfL, while the epitope of *anti*-NfL antibody lies within the C-terminus of the protein.

As shown in the Western blot below, binding to NfL varies among the different commercially available antibodies despite using similar protein loadings, as shown by Ponceau S staining. Furthermore, the prominent multiple bands observed for En-hNfL suggest the presence of degradation products containing epitopes that remain reactive to detection by Western.

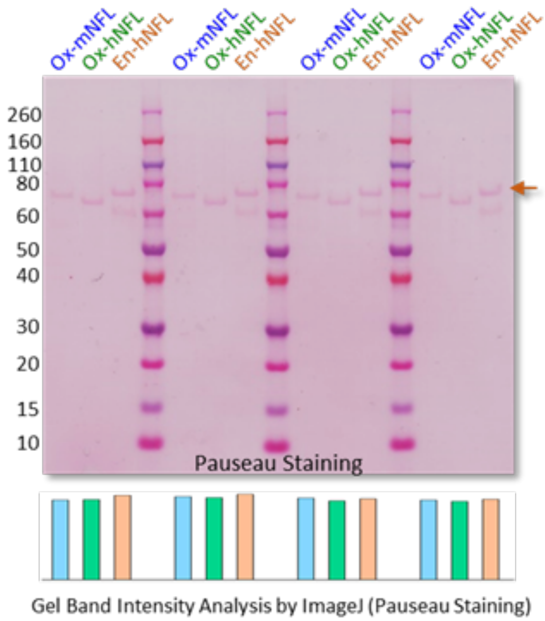


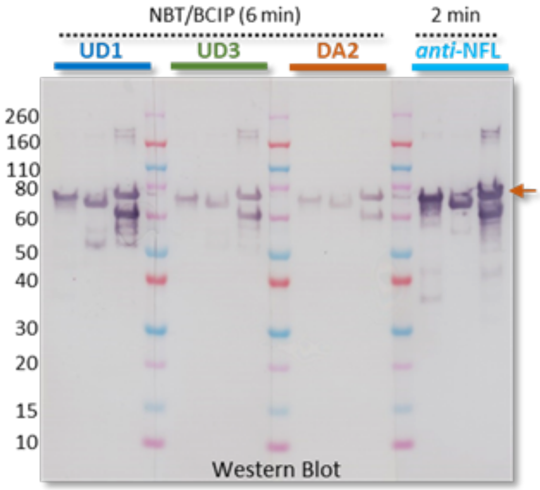

### Supplementary Methods

#### 1. Small Molecule Synthesis

##### 1.1 General Experimental Procedures

Chemicals, solvents, media and *Escherichia coli* cell stocks were purchased from commercial suppliers in the United Kingdom (Acros, Alfa Aesar, Carbosynth, Fisher Scientific, Fluorochem, Sigma Aldrich, VWR) and used as received unless stated otherwise. Unless available as in-house supply, anhydrous solvents were purchased from Sigma Aldrich and stored under argon in a septum-capped bottle. All solvents were either analytical or HPLC grade. Reactions requiring anhydrous conditions were carried out with flame-dried reaction vessels under an inert atmosphere of nitrogen or argon. Thin layer chromatography (TLC) was performed on Merck Millipore aluminium TLC plate, silica gel coated with fluorescent indicator F254 and visualised under UV irradiation at 254 nm, or by staining with aqueous potassium permanganate ($KMnO_4$ (3.00 g), $K_2CO_3$ (20.0 g), 5% NaOH (5 mL), $H_2O$ (300 mL)) with subsequent drying with a heat gun. Flash column chromatography was performed using Geduran® Silica Gel 60 (0.040-0.063 mm) or The Teledyne ISCO CombiFlash NextGen 100 Flash Chromatography System. All NMR spectra were recorded in commercially available deuterated solvents on a Bruker AVIIIHD 400 nanobay ($^1$H at 400, $^{13}$C at 101, $^{19}$F 376) or AVII HD 600 ($^1$H at 600, $^{13}$C at 151, $^{19}$F 565). All NMR data were processed using Mestrenova v.14.1.0. Chemical shifts are quoted in parts per million (ppm) relative to the centre of the solvent peak for $^1$H and $^{13}$C spectral data. All coupling constants are reported in Hz to the nearest 0.1 Hz. $^{13}$C spectra are $^1$H decoupled and $^{19}$F-$^{13}$C heteronuclear coupling on $^{13}$C spectra are reported. $^{19}$F NMR signals are referenced relative to $CFCl_3$. Infrared (IR) spectra of compounds were acquired on a Bruker Tensor 27 Fourier Transform spectrometer as crystals using a diamond attenuated total reflectance (ATR) attachment or as thin films of neat oils. Melting points were measured using a Griffin apparatus on a Leica hotstage microscope and are uncorrected. Low resolution mass spectra were recorded on an Agilent 6120 Quadrupole spectrometer equipped with an electrospray ion source. High resolution mass spectra were obtained on a Thermo Exactive High-Resolution Orbitrap FTMS mass spectrometer with a lock-spray electrospray ion source. Methanol or acetonitrile were used as the carrier solvents. Some compounds were not stable under the MS ionisation methods and therefore, HRMS for these compounds were not obtained.

#### 1.2 Reagent synthesis

General procedure 1: Difluoromethylation of heteroaromatic thiols with sodium chlorodifluoroacetate

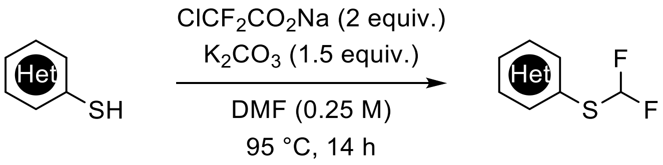


The following procedure was adapted from a known procedure.[5]

For reaction on a 12.0 mmol scale:

Under vacuum, $K_2CO_3$ (1.5 equiv.) was dried using a heat gun before sodium chlorodifluoroacetate (2.0 equiv.) was added and the system was purged 3 times with nitrogen. Dry DMF (0.25 M) was then added followed by the heteroaromatic thiol (1.0 equiv.) slowly with stirring. The reaction was left to stir at 95 ˚C for 14 h before cooling to rt and diluting with EtOAc (100 mL). $H_2O$ (100 mL) was added and the layers were separated. The aqueous layer was extracted with EtOAc (2 x 50 mL) and the combined organic layers were washed with sat. aq. LiCl (50 mL) and brine (50 mL). The organic phase was then dried over $MgSO_4$, filtered and concentrated under reduced pressure. Purification was performed using silica gel column chromatography.

General procedure 2: Fluoroalkylation of heteroaromatic thiols with dibromofluoromethane

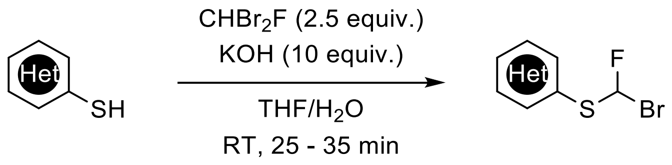


The following procedure was adapted from a known procedure.[2]

For a reaction on 6.03 mmol scale:
KOH (10 equiv.) was dissolved in $H_2O$ (7.5 M) at 0 °C. A solution of the heteroaromatic thiol (1 equiv.) in THF (1.0 M) was added and the resulting solution was allowed to stir at rt for 20 min. Dibromofluoromethane (2.5 equiv.) in THF (7.5 M) was added dropwise slowly over 15 min at rt. The reaction was left to stir at rt for a further 10 – 20 min before diluting with $H_2O$ (40 mL). DCM (60 mL) was added and the layers were separated. The aqueous layer was extracted with DCM (2 x 60 mL). The combined organic phase was then dried over $MgSO_4$, filtered and concentrated under reduced pressure. Purification was performed using silica gel column chromatography.

Notes:
1) Increasing reaction time will lead to further substitution on the desired product by a second alkylation step.
2) Impurity present is the difluoromethylated thiol.

General procedure 3: Oxidation of fluoroalkylated sulfides

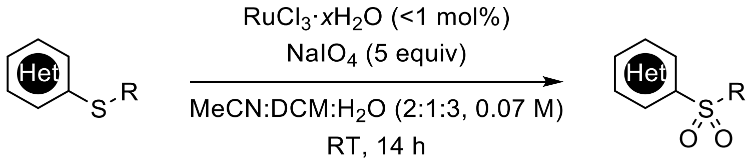


The following procedure was adapted from a known procedure.[6]

For reaction on 6.50 mmol scale:

To a mixture of difluoromethylated sulfide (1.0 equiv.) in $CH_3CN$, DCM and water (2:1:3, 0.07 M) at 0 °C, $NaIO_4$ (5.0 equiv.) followed by ruthenium trichloride hydrate (10 mg) were added. The reaction mixture was then allowed to stir at RT for 14 h before diluting with $H_2O$ (60 mL) while stirring. The crude mixture was extracted with $Et_2O$ (3 x 100 mL) and the combined organic layers were washed with sat. aq. $NaHCO_3$ (100 mL) and brine (100 mL). The organic phase was then dried over $MgSO_4$, filtered and concentrated under reduced pressure. Purification was performed using silica gel column chromatography.

#### 2-((difluoromethyl)thio)pyridine

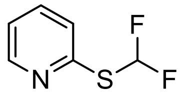

Following general procedure 1, 2-((difluoromethyl)thio)pyridine was prepared from 2-pyridinethiol (600 mg, 5.40 mmol), $K_2CO_3$ (1.12 g, 8.10 mmol) and sodium chlorodifluoroacetate (1.65 g, 10.8 mmol). Purification by column chromatography (5% EtOAc in pet. ether) gave 2-((difluoromethyl)thio)pyridine as a colourless liquid (0.65 g, 74%).

$C_6H_5F_2NS$ (161.2 g/mol):
**$^1$H NMR** (400 MHz, $CDCl_3$) δ 8.42 (ddd, *J* = 5.0, 1.9, 0.9 Hz, 1H), 7.70 (t, *J* = 56 Hz, 1H), 7.54 (td, *J* = 7.8, 1.9 Hz, 1H), 7.20 (dt, *J* = 8.0, 1.0 Hz, 1H), 7.08 (ddd, *J* = 7.5, 4.9, 1.1 Hz, 1H). **$^{19}$F NMR** (376 MHz, $CDCl_3$) δ -96.2 (d, *J* = 56 Hz). **$^{13}$C NMR** (101 MHz, $CDCl_3$) δ 153.4 (t, *J* = 3.6 Hz), 150.2, 137.2, 124.5 (t, *J* = 2.1 Hz), 121.9, 121.4 (t, *J* = 271 Hz). **MS$^+$** 162.0 $[M + H]^+$

Spectroscopic data was consistent with literature reports.[6]

2-((difluoromethyl)thio)pyrimidine

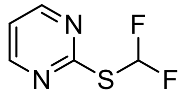

Following general procedure 1, 2-((difluoromethyl)thio)pyrimidine was prepared from 2-pyrimidinethiol (1.40 g, 12.5 mmol), $K_2CO_3$ (2.48 g, 17.9 mmol) and sodium chlorodifluoroacetate (3.65 g, 23.9 mmol). Purification by column chromatography (5% EtOAc in pet. ether) gave 2-((difluoromethyl)thio)pyrimidine as a pale yellow liquid (1.18 g, 58%).

$C_5H_4F_2N_2S$ (162.2 g/mol):
**$^1$H NMR** (400 MHz, $CDCl_3$) δ 8.57 (d, $J$ = 4.9 Hz, 2H), 7.78 (t, $J$ = 56 Hz, 1H), 7.11 (t, $J$ = 4.9 Hz, 1H). **$^{19}$F NMR** (376 MHz, $CDCl_3$) δ -99.2 (d, $J$ = 56 Hz). **$^{13}$C NMR** (101 MHz, $CDCl_3$) δ 168.0 (t, $J$ = 5.8 Hz), 157.9 (2C), 120.8 (t, $J$ = 270 Hz), 118.1. **MS$^+$** 163.0 $[M + H]^+$

Spectroscopic data was consistent with literature reports.[5]

2-((difluoromethyl)thio)benzothiazole

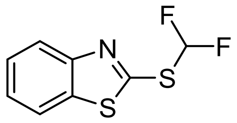

Following general procedure 1, 2-((difluoromethyl)thio)benzothiazole was prepared from 2-benzothiazolethiol (2.00 g, 12.5 mmol), $K_2CO_3$ (2.48 g, 17.9 mmol) and sodium chlorodifluoroacetate (3.65 g, 23.9 mmol). In contrast to the reported procedure[2], $^{19}F$ NMR of the crude reaction mixture revealed a 3:2 ratio in favour of N-difluoromethylation instead of the desired S-difluoromethylated product. The orange mixture was used in the next without purification (1.60 g, 40% purity from NMR).

$C_8H_5F_2NS_2$ (217.3 g/mol):
**$^1$H NMR** (400 MHz, $CDCl_3$) δ 8.00 (ddd, *J* = 8.2, 1.2, 0.6 Hz, 1H), 7.83 (ddd, *J* = 8.0, 1.3, 0.6 Hz, 1H), 7.65 (t, *J* = 56 Hz, 1H), 7.50 (ddd, *J* = 8.3, 7.2, 1.3 Hz, 1H), 7.41 (ddd, *J* = 8.4, 7.3, 1.2 Hz, 1H). **$^{19}$F NMR** (376 MHz, $CDCl_3$) δ -93.2 (d, *J* = 56 Hz). **$^{13}$C NMR** (101 MHz, $CDCl_3$) δ 157.2 (t, *J* = 4.2 Hz), 153.0, 136.1, 126.8, 125.7, 123.0, 121.3, 120.4 (t, *J* = 280 Hz). **MS$^+$** 218.0 $[M + H]^+$

Spectroscopic data was consistent with literature reports.[6]

#### 2-((fluoromethyl)thio)benzothiazole

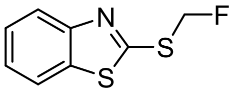

Under argon, to a vial containing 2-benzothiazolethiol (84 mg, 502 μmol), $Cs_2CO_3$ (196 mg, 603 μmol) in anhydrous MeCN (1.0 mL), fluoromethyl tosylate (123 μL, 603 μmol) was added dropwise. The reaction was allowed to stir for 16 h at 70 ˚C before diluting with $H_2O$ (5 mL). DCM (10 mL) was added and the layers were separated. The aqueous layer was extracted with DCM (2 x 10 mL). The combined organic phase was washed with 1 M NaOH (10 mL) then dried over $MgSO_4$, filtered and concentrated under reduced pressure. Combiflash purification by silica gel column chromatography (100% pet. ether to 80% of 45% EtOAc in pet. ether over 10 min) gave 2-((fluoromethyl)thio)benzothiazole as a white solid (74 mg, 74%).

$C_8H_6FNS_2$ (199.3 g/mol):
**$^1$H NMR** (600 MHz, $CDCl_3$) δ 7.97 (ddd, *J* = 8.2, 1.2, 0.6 Hz, 1H), 7.81 (ddd, *J* = 8.1, 1.3, 0.6 Hz, 1H), 7.47 (ddd, *J* = 8.3, 7.3, 1.2 Hz, 1H), 7.37 (ddd, *J* = 8.3, 7.3, 1.2 Hz, 1H), 6.16 (d, *J* = 51.1 Hz, 2H). **$^{19}$F NMR** (565 MHz, $CDCl_3$) δ -186.7 (d, *J* = 50.6 Hz). **$^{13}$C NMR** (151 MHz, $CDCl_3$) δ 162.5 (d, *J* = 3.8 Hz), 153.1, 135.9, 126.6, 125.2, 122.5, 121.3, 85.0 (d, *J* = 222.0 Hz). **MS$^+$** 199.9 $[M + H]^+$

Spectroscopic data was consistent with literature reports.[7]

#### 2-((bromofluoromethyl)thio)pyridine

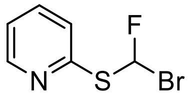

Following general procedure 2, 2-((bromofluoromethyl)thio)pyridine was prepared from 2-pyridinethiol (670 mg, 6.03 mmol), KOH (1.38 g, 60.2 mmol) and dibromofluoromethane (1.20 mL, 15 mmol). Purification by column chromatography (0 to 2% EtOAc in pet. ether) gave 2-((bromofluoromethyl)thio)pyridine as a pale yellow liquid (61 mg, 5%).

$C_6H_5BrFNS$ (220.9 g/mol):
**TLC:** $R_f$ = 0.5 (10% EtOAc/pet. ether) **$^1$H NMR** (600 MHz, $CDCl_3$) δ 8.52 (m, 1H), 8.09 (t, *J* = 54.8 Hz, 1H), 7.62 (td, *J* = 7.7, 1.9 Hz, 1H), 7.22 (dt, *J* = 8.0, 0.9 Hz, 1H), 7.15 (ddd, *J* = 7.5, 4.9, 1.1 Hz, 1H). **$^{19}$F NMR** (565 MHz, $CDCl_3$) δ -103.9 (d, *J* = 55.2 Hz). **$^{13}$C NMR** (151 MHz, $CDCl_3$) δ 155.4 (d, *J* = 2.2 Hz), 150.2, 137.3, 122.9 (d, *J* = 1.8 Hz), 121.9, 89.9 (t, *J* = 292.7 Hz). **HRMS** the compound did not ionize.

2-((bromofluoromethyl)thio)pyrimidine

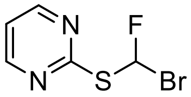


Following general procedure 2, 2-((bromofluoromethyl)thio)pyrimidine was prepared from 2-pyrimidinethiol (675 mg, 6.02 mmol), KOH (1.38 g, 60.2 mmol) and dibromofluoromethane (1.20 mL, 15 mmol). Purification by column chromatography (0 to 2% EtOAc in pet. ether) gave 2-((bromofluoromethyl)thio)pyrimidine as a colourless liquid (227 mg, 17%).

$C_5H_4BrFN_2S$ (221.9 g/mol):
**$^1$H NMR** (600 MHz, $CDCl_3$) δ 8.57 (d, *J* = 4.9 Hz, 2H), 8.07 (d, *J* = 54.2 Hz, 1H), 7.10 (t, *J* = 4.9 Hz, 1H). **$^{19}$F NMR** (376 MHz, $CDCl_3$) δ -107.3 (d, *J* = 54.2 Hz). **$^{13}$C NMR** (101 MHz, $CDCl_3$) δ 168.8 (d, *J* = 2.1 Hz), 157.9 (2C), 118.4, 89.1 (d, *J* = 292.6 Hz). **HRMS** the compound did not ionize.

#### 2-((bromofluoromethyl)thio)benzothiazole

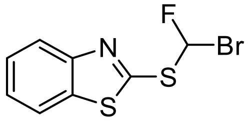


Following general procedure 2, 2-((bromofluoromethyl)thio)benzothiazole was prepared from 2-benzothiazolethiol (1.04 mg, 6.03 mmol), KOH (1.38 g, 60.2 mmol) and dibromofluoromethane (1.20 mL, 15 mmol). Purification by column chromatography (0 to 2% EtOAc in pet. ether) gave 2-((bromofluoromethyl)thio)benzothiazole as a colourless liquid (217 mg, 13%).

$C_8H_5BrFNS_2$ (278.2 g/mol):
**$^1$H NMR** (600 MHz, $CDCl_3$) δ 8.01 (ddd, *J* = 8.2, 1.2, 0.7 Hz, 1H), 7.96 (d, *J* = 54.9 Hz, 1H), 7.84 (ddd, *J* = 8.2, 1.2, 0.6 Hz, 1H), 7.50 (ddd, *J* = 8.3, 7.2, 1.2 Hz, 1H), 7.41 (ddd, *J* = 8.3, 7.3, 1.2 Hz, 1H). **$^{19}$F NMR** (565 MHz, $CDCl_3$) δ -104.1 (d, *J* = 55.2 Hz). **$^{13}$C NMR** (151 MHz, $CDCl_3$) δ 160.1 (d, *J* = 2.5 Hz), 152.8, 135.9, 126.8, 125.7, 123.0, 121.4, 89.3 (d, *J* = 297.8 Hz). **MS$^+$** 277.9 $[M + H]^+$

Spectroscopic data was consistent with literature reports.[2]

2-((bromomethyl)thio)benzothiazole

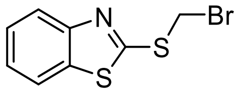

A solution of 2-benzothiazolethiol (168 mg, 1.00 mmol) in THF (1 mL) was added dropwise over 20 minutes to a mixture of $Cs_2CO_3$ (491 mg, 1.51 mmol) in dibromomethane (5 mL) and MeCN (1.5 mL) while stirring at rt. After complete addition, the reaction was monitored by TLC every 10 minutes for complete consumption of 2-mercaptobenzothiazole. The reaction mixture was then filtered and the filtrate was concentrated. Combiflash purification by silica gel column chromatography (0 to 50% DCM in pet. ether over 10 min) gave a white solid (75 mg, 29%).

$C_8H_6BrNS_2$ (260.2 g/mol):

**TLC:** $R_f$ = 0.8 (70% DCM/pet. ether) **$^1$H NMR** (400 MHz, $CDCl_3$) δ 7.98 (ddd, *J* = 8.2, 1.2, 0.6 Hz, 1H), 7.82 (ddd, *J* = 8.0, 1.2, 0.6 Hz, 1H), 7.48 (ddd, *J* = 8.3, 7.2, 1.2 Hz, 1H), 7.37 (ddd, *J* = 8.0, 7.2, 1.2 Hz, 1H), 5.22 (s, 2H). **$^{13}$C NMR** (101 MHz, $CDCl_3$) δ 162.7, 153.0, 135.7, 126.7, 125.1, 122.5, 121.4, 31.3. **IR** (ATR): $\tilde{\upsilon}$/cm$^{-1}$ = 3027, 2921, 2851, 1555, 1454, 1422, 1365, 1309, 1275, 1234, 1192, 1128, 1069, 1018, 999, 941, 807, 758, 728, 178.

##### 2-((difluoromethyl)sulfonyl)pyridine (S1 – pySOOF)

PySOOF (S1) was commercially purchased.

##### 2-((difluoromethyl)sulfonyl)pyrimidine (S2 – pymSOOF)

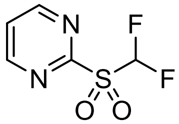


Following general procedure 3, pymSOOF was prepared from a mixture of the starting sulfide (1.05 g, 6.48 mmol), $NaIO_4$ (6.92 g, 32.4 mmol) and ruthenium trichloride hydrate (10 mg). Purification by column chromatography (3% MeOH in DCM) gave pymSOOF as a colourless liquid (0.96 g, 76%).

$C_5H_4F_2N_2O_2S$ (194.2 g/mol):
**$^1$H NMR** (400 MHz, $CDCl_3$) δ 9.01 (d, *J* = 4.9 Hz, 2H), 7.69 (t, *J* = 4.9 Hz, 1H), 6.91 (t, *J* = 53 Hz, 1H). **$^{19}$F NMR** (376 MHz, $CDCl_3$) δ -124.5 (d, *J* = 53 Hz). **$^{13}$C NMR** (101 MHz, $CDCl_3$) δ 163.2, 159.2 (2C), 125.2, 113.9 (t, *J* = 285.7 Hz). **MS$^+$** 195.0 $[M + H]^+$, 217.0 $[M + Na]^+$

Spectroscopic data was consistent with literature reports.[3]

#### 2-((difluoromethyl)sulfonyl)benzothiazole (S3 – btSOOF)

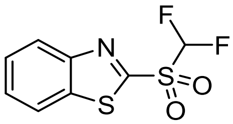


Following general procedure 3, btSOOF was prepared from a mixture of the starting sulfide (1.60 g, 40% purity from NMR), $NaIO_4$ (7.88 g, 36.8 mmol) and ruthenium trichloride hydrate (15 mg) in $CH_3CN$ (40 mL), DCM (20 mL) and water (60 mL). Purification was performed instead by trituration in ether (5 mL) to give btSOOF as a white solid (0.58 g, 79%).

$C_8H_5F_2NO_2S_2$ (249.2 g/mol):
**$^1$H NMR** (400 MHz, DMSO-$d_6$) δ 8.48 – 8.37 (m, 2H), 8.48 – 8.37 (m, 2H), 7.70 (t, $J$ = 52 Hz, 1H). **$^{19}$F NMR** (376 MHz, DMSO-$d_6$) δ -122.9 (d, $J$ = 52 Hz). **$^{13}$C NMR** 159.0, 152.4, 137.5, 129.2, 128.6, 125.6, 123.8, 115.0 (t, $J$ = 285 Hz). **MS$^+$** 250.0 $[M + H]^+$

Spectroscopic data was consistent with literature reports.[3,6]

2-(bromofluoromethyl)sulfonyl)pyridine

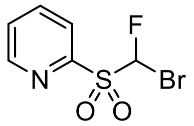

Following general procedure 3, 2-(bromofluoromethyl)sulfonyl)pyridine was prepared from a mixture of the starting sulfide (71 mg, 320 µmol), $NaIO_4$ (324 mg, 1.60 mmol) and ruthenium trichloride hydrate (4 mg). Purification by column chromatography (0 to 80% EtOAc in pet. ether) gave the sulfone product as a white solid (75 mg, 92%).

$C_6H_5BrFNO_2S$ (252.9 g/mol):
**TLC:** $R_f$ = 0.4 (60% EtOAc/pet. ether) **$^1$H NMR** (600 MHz, $CDCl_3$) δ 8.81 (ddd, *J* = 4.7, 1.7, 0.9 Hz, 1H), 8.20 (dt, *J* = 7.8, 1.0 Hz, 1H), 8.05 (td, *J* = 7.8, 1.7 Hz, 1H), 7.67 (ddd, *J* = 7.7, 4.7, 1.1 Hz, 1H), 7.46 (t, *J* = 48.8 Hz, 1H). **$^{19}$F NMR** (565 MHz, $CDCl_3$) δ -142.6 (d, *J* = 49.2 Hz). **$^{13}$C NMR** (151 MHz, $CDCl_3$) δ 152.7, 150.9, 138.6, 128.7, 125.2, 95.4 (d, *J* = 292.6 Hz). **HRMS** (ESI (+), MeOH): (*m/z*) calc. for $C_6H_5BrFNO_2S$: 253.9281 $[M + H]^+$; found: 253.9282

2-(bromofluoromethyl)sulfonyl)pyrimidine

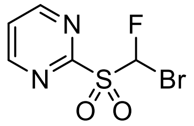


Following general procedure 3, 2-(bromofluoromethyl)sulfonyl)pyrimidine was prepared from a mixture of the starting sulfide (227 mg, 1.02 mmol), $NaIO_4$ (1.09 g, 5.10 mmol) and ruthenium trichloride hydrate (10 mg). Purification by column chromatography (0 to 80% EtOAc in pet. ether) gave the sulfone product as a colourless oil (180 mg, 69%).

$C_5H_4BrFN_2O_2S$ (255.1 g/mol):
**TLC:** $R_f$ = 0.2 (60% EtOAc/pet. ether) **$^1$H NMR** (600 MHz, $CDCl_3$) δ 9.00 (d, *J* = 4.9 Hz, 2H), 7.71 (d, *J* = 48.5 Hz, 1H), 7.69 (t, *J* = 4.9 Hz, 1H). **$^{19}$F NMR** (565 MHz, $CDCl_3$) δ -141.9 (d, *J* = 48.1 Hz). **$^{13}$C NMR** (151 MHz, $CDCl_3$) δ 162.6, 159.2 (2C), 125.1, 95.4 (d, *J* = 293.6 Hz). **HRMS** the compound did not ionize.

2-(bromofluoromethyl)sulfonyl)benzothiazole

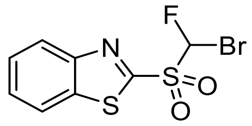


Following general procedure 3, 2-(bromofluoromethyl)sulfonyl)benzothiazole was prepared from a mixture of the starting sulfide (217 mg, 0.781 mmol), $NaIO_4$ (832 mg, 3.91 mmol) and ruthenium trichloride hydrate (10 mg). Purification by column chromatography (0 to 80% EtOAc in pet. ether) gave the sulfone product as a white solid (148 mg, 69%).

$C_8H_5BrFNO_2S_2$ (310.2 g/mol):
**TLC:** $R_f$ = 0.6 (60% EtOAc/pet. ether) **$^1$H NMR** (600 MHz, $CDCl_3$) δ 8.29 (ddd, *J* = 8.3, 1.5, 0.7 Hz, 1H), 8.07 (ddd, *J* = 7.8, 1.5, 0.7 Hz, 1H), 7.71 – 7.66 (m, 2H), 7.37 (d, *J* = 48.7 Hz, 1H). **$^{19}$F NMR** (565 MHz, $CDCl_3$) δ -140.2 (d, *J* = 48.4 Hz). **$^{13}$C NMR** (151 MHz, $CDCl_3$) δ 159.4, 152.9, 137.9, 129.1, 128.3, 126.2, 122.5, 96.0 (d, *J* = 295.0 Hz). **HRMS** (ESI (+), MeOH): (*m/z*) calc. for $C_8H_5BrFNO_2S_2$: 309.9002 $[M + H]^+$; found: 309.9014

#### Ethyl 2-(benzothiazole-2-ylthio)-2-fluoroacetate

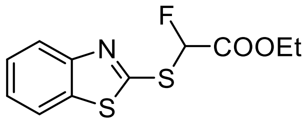


The following procedure was adapted from a known procedure.[8]

Triethylamine (2.08 mL, 15.0 mmol) was added dropwise to a mixture of 2-mercaptobenzothiazole (2.50 g, 15.0 mmol) in EtOH (35 mL) under nitrogen. The solution was allowed to stir at rt for 15 min before adding ethyl bromofluoroacetate (1.95 mL, 16.4 mmol) and leaving to stir for a further 16 h. The crude mixture was quenched with 1 M HCl (20 mL) and extracted with DCM (3 x 50 mL) and the combined organic layers were washed with $H_2O$ (50 mL) and brine (50 mL). The organic phase was then dried over $MgSO_4$, filtered and concentrated under reduced pressure. Purification by silica gel column chromatography (5% EtOAc in pet. ether) gave a colourless oil (3.85 g, 95%).

$C_{11}H_{10}FNO_2S_2$ (271.3 g/mol):
**$^1$H NMR** (400 MHz, $CDCl_3$) δ 7.97 (ddd, *J* = 8.1, 1.2, 0.6 Hz, 1H), 7.81 (ddd, *J* = 8.0, 1.3, 0.6 Hz, 1H), 7.47 (ddd, *J* = 8.3, 7.3, 1.3 Hz, 1H), 7.37 (ddd, *J* = 8.3, 7.2, 1.2 Hz, 1H), 6.94 (d, *J* = 51 Hz, 1H), 4.33 (q, *J* = 7.1 Hz, 2H), 1.32 (t, *J* = 7.2 Hz, 3H). **$^{19}$F NMR** (376 MHz, $CDCl_3$) δ -161.1 (d, *J* = 51 Hz).

Spectroscopic data was consistent with literature reports.[9]

#### Ethyl 2-(benzothiazole-2-ylsulfonyl)-2-fluoroacetate

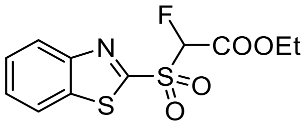


Following general procedure 3, ethyl 2-(benzothiazole-2-ylthio)-2-fluoroacetate was prepared from a mixture of the starting sulfide (400 mg, 1.47 mmol), $NaIO_4$ (1.58 g, 7.37 mmol) and ruthenium trichloride hydrate (3 mg). Purification by column chromatography (0 to 60% EtOAc in pet. ether over 10 min) gave the sulfone product as a yellow solid (272 mg, 61%).

$C_{11}H_{10}FNO_4S_2$ (303.3 g/mol):
**$^1$H NMR** (400 MHz, $CDCl_3$) δ 8.30 – 8.24 (m, 1H), 8.08 – 8.01 (m, 1H), 7.71 – 7.62 (m, 2H), 6.04 (d, $J$ = 47.5 Hz, 1H), 4.47 – 4.34 (m, 2H), 1.34 (t, $J$ = 7.1 Hz, 3H). **$^{19}$F NMR** (376 MHz, $CDCl_3$) δ -181.0 (d, $J$ = 47.6 Hz).

Spectroscopic data was consistent with literature reports.[9]

#### 2-((fluoromethyl)sulfonyl)benzothiazole (S4 – mono-btSOOF)

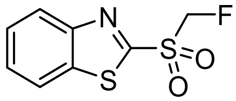

The following procedure was adapted from a known procedure.[10]

To a solution of $BzSO_2CFHCO_2Et$ (245 mg, 808 μmol) in EtOAc (15 mL) was added DBU (170 μL, 1.13 mmol) and $H_2O$ (50 μL). The resulting mixture was allowed to stir at rt for 16 h before quenching with sat. aq. $NH_4Cl$ (5 mL). The organic layer was separated and the aqueous layer was extracted with DCM (2 x 10 mL). The combined organic layers were washed with brine (20 mL) and then dried over $MgSO_4$, filtered and concentrated under reduced pressure. Purification was performed instead by trituration in a cold mixture of ether:DCM (3:1) to a white solid (95 mg, 51%).

$C_8H_6FNO_2S_2$ (231.3 g/mol):
**$^1$H NMR** (400 MHz, DMSO-*d*$_6$) δ 8.44 – 8.36 (m, 1H), 8.37 – 8.29 (m, 1H), 7.82 – 7.72 (m, 2H), 6.14 (d, *J* = 45 Hz, 2H). **$^{19}$F NMR** (376 MHz, DMSO-*d*$_6$) δ -208.3 (t, *J* = 45 Hz). **MS$^+$** 232.0 $[M + H]^+$

Spectroscopic data was consistent with literature reports.[11]

*tert*-Butyl N-(*tert*-butoxycarbonyl)-L-homoserinate (Boc-L-hSe-O*t*Bu)

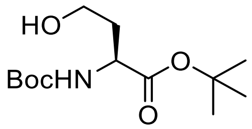


Under argon, Boc-L-Asp-O*t*Bu (5.00 g, 17.3 mmol) was dissolved in dry THF (170 mL) and the solution was cooled to 0 °C. Isobutyl chloroformate (6.72 mL, 51.8 mmol) and DIPEA (4.52 mL, 25.9 mmol) were added and the resulting mixture was allowed to stir at 0 °C for 1 h before warming to RT and stirring for a further 30 min. Sodium borohydride (4.58 g, 121 mmol) was added slowly at 0 °C followed by $H_2O$ (40 mL) under a stream of nitrogen. The mixture was left to stir overnight at RT and then acidified with 1 M HCl until pH 2. The organic later was separated and the aqueous phase was extracted with EtOAc (3 x 100 mL). The combined organic layers was washed with sat. aq. $NaHCO_3$ (2 x 100 mL) and brine (100 mL) then dried over $MgSO_4$, filtered and concentrated under reduced pressure. Combiflash purification by silica gel column chromatography (0 to 80% EtOAc in pet. ether over 18 min) gave Boc-L-Hse(homoserine)-O*t*Bu as a colourless oil (4.36 g, 92%).

$C_{13}H_{25}NO_5$ (275.4 g/mol):
**$^1$H NMR** (400 MHz, $CDCl_3$) δ 5.36 (br s, 1H), 4.38 – 4.30 (m, 1H), 3.72 – 3.59 (m, 3H), 2.17 – 2.08 (m, 1H), 1.56 – 1.49 (m, 1H), 1.46 (s, 9H), 1.44 (s, 9H). **$^{13}$C NMR** (101 MHz, $CDCl_3$) δ 172.2, 156.8, 82.4, 80.5, 58.4, 51.0, 36.7, 28.4 (3C), 28.1 (3C).

Spectroscopic data was consistent with literature reports.[12]

*tert*-Butyl N-(*tert*-butoxycarbonyl)-O-tosyl-L-homoserinate (Boc-L-hSe(OTs)-O*t*Bu)

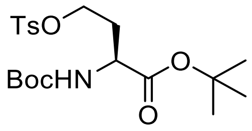


The following procedure was adapted from a known procedure.[12]

Under argon, Boc-L-Hse-O*t*Bu (825 mg, 3.00 mmol) was dissolved in dry DCM (9 mL) and the solution was cooled to 0 °C. $NEt_3$ (2.09 mL, 15.0 mmol), TsCl (1.14 g, 5.99 mmol) and DMAP (36.6 mg, 0.300 mmol) were added sequentially and the resulting mixture was allowed to stir at 0 °C for 15 min before warming to RT and stirring for a further 16 h. The mixture was then diluted with DCM (12 mL) and the organic phase was washed with $H_2O$ (2 x 20 mL) and brine (10 mL) then dried over $MgSO_4$, filtered and concentrated under reduced pressure. Combiflash purification by silica gel column chromatography (5 to 15% EtOAc in pet. ether over 15 min) gave Boc-L-Hse(OTs)-O*t*Bu as a white solid (921 mg, 72%).

$C_{20}H_{31}NO_7S$ (429.5 g/mol):
**$^1$H NMR** (400 MHz, $CDCl_3$) δ 7.78 (d, *J* = 8.4 Hz, 2H), 7.34 (d, *J* = 8.2 Hz, 2H), 5.01 (d, *J* = 5.9 Hz, 1H), 4.17 – 4.15 (m, 1H), 4.08 (td, *J* = 6.5, 2.1 Hz, 2H), 2.44 (s, 3H), 2.23 – 2.18 (m, 1H), 2.08 – 1.99 (m, 1H), 1.44 (s, 9H), 1.40 (s, 9H). **$^{13}$C NMR** (101 MHz, $CDCl_3$) δ 170.7, 155.3, 145.0, 132.9, 130.0 (2C), 128.2 (2C), 82.8, 80.0, 66.7, 51.2, 31.8, 28.4 (3C), 28.0 (3C), 21.8. **MS$^+$** 430.1 $[M + H]^+$, 452.2 $[M + Na]^+$ **Alpha D** $[\alpha]_D^{25}$ = -16.2 (c = 1.0, MeOH)

Spectroscopic data was consistent with literature reports.[12]

##### *tert*-Butyl (S)-2-(*tert*-butoxycarbonyl)amino)-4-fluorobutanoate (Boc-L-Fha-O*t*Bu)

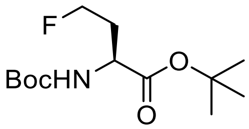


The following procedure was adapted from a known procedure.[12]

Under argon, Boc-L-Hse(OTs)-O*t*Bu (847 mg, 1.97 mmol) was dissolved in anhydrous *t*BuOH (20 mL). TBAF·3$H_2$O (1.87 g, 5.92 mmol) was added and the resulting mixture was allowed to stir at 70 °C for 3 h before quenching with sat. aq. $NaHCO_3$ (20 mL). The mixture was extracted with DCM (3 x 30 mL) and the combined organic phases was dried over $MgSO_4$, filtered and concentrated under reduced pressure. Combiflash purification by silica gel column chromatography (0 to 20% EtOAc in pet. ether over 13 min) gave Boc-L-Fha-O*t*Bu as a white solid (397 mg, 73%).

$C_{13}H_{24}FNO_4$ (277.3 g/mol):
**$^1$H NMR** (400 MHz, $CDCl_3$) δ 5.21 (d, *J* = 6.2 Hz, 1H), 4.64 – 4.55 (m, 1H), 4.52 – 4.43 (m, 1H), 4.34 – 4.24 (m, 1H), 2.31 – 1.96 (m, 2H), 1.47 (s, 9H), 1.44 (s, 9H). **$^{19}$F NMR** (376 MHz, $CDCl_3$) δ -219.4 (m). **$^{13}$C NMR** (101 MHz, $CDCl_3$) δ 171.1, 155.5, 82.4, 80.6 (d, *J* = 166 Hz), 80.0, 51.3, 33.5 (d, *J* = 19.8 Hz), 28.4 (3C), 28.1 (3C). **MS$^+$** 278.4 $[M + H]^+$

Spectroscopic data was consistent with literature reports.[12]

#### (S)-2-amino-4-fluorobutanoic acid trifluoroacetate (L-Fha•$CF_3CO_2H$)

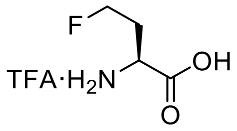


A solution of Boc L-Fha-O*t*Bu (386 mg, 1.39 mmol) in $H_2O$ (1.4 mL) was cooled to 0 °C. TFA (12.5 mL) was added and the reaction was allowed to stir at RT for 4 h. The solution was concentrated under a stream of nitrogen and dried azeotropically under reduced pressure with toluene to give the TFA salt of L-Fha-OH as a colourless solid (quant. yield). The unprotected amino acid was used directly in the next step.

##### (S)-2-((((9H-fluoren-9-yl)methoxy)carbonyl)amino)-4-fluorobutanoic acid (Fmoc-L-Fha)

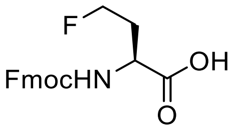


FmocOSu (493 mg, 1.46 mmol) was added to a mixture of crude L-Fha-OH (1.39 mmol) in THF (10 mL) and sat. aq. $NaHCO_3$ (5 mL). The resulting mixture was stirred at RT for 16 h and then diluted with $H_2O$ (10 mL). The aqueous layer was washed with $Et_2O$ (2 x 25 mL) and acidified with 1 M HCl until pH 1-2. The aqueous phased was extracted with EtOAc (3 x 30 mL). The combined organic layers was dried over $MgSO_4$, filtered and concentrated under reduced pressure to give Fmoc-L-Fha-OH as a white solid (414 mg, 82% by $^1$H NMR) without further purification.

$C_{19}H_{18}FNO_4$ (343.35 g/mol):
**$^1$H NMR** (600 MHz, Acetone-d$_6$) δ 7.86 (d, *J* = 7.6 Hz, 2H), 7.72 (t, *J* = 6.6 Hz, 2H), 7.41 (t, *J* = 7.5 Hz, 2H), 7.32 (td, *J* = 7.4, 1.2 Hz, 2H), 6.83 (d, *J* = 8.5 Hz, 1H), 6.83 (d, *J* = 8.6, 1H), 4.68 – 4.62 (m, 1H), 4.60 – 4.54 (m, 1H), 4.43 – 4.38 (m, 1H), 4.37 – 4.33 (m, 2H), 4.25 (t, *J* = 7.2 Hz, 1H), 2.39 – 2.30 (m, 1H), 2.17 – 2.09 (m, 1H).
**$^{19}$F NMR** (565 MHz, Acetone-d$_6$) δ -221.5 (tdd, *J* = 47.3, 29.6, 21.0 Hz). **$^{13}$C NMR** (151 MHz, Acetone-d$_6$) δ 173.5, 157.1, 145.1 (2C), 142.1 (2C), 128.5 (2C), 127.9 (2C), 126.2 (2C), 120.8 (2C), 81.2 (d, *J* = 164.0 Hz), 67.2, 51.3 (d, *J* = 4.9 Hz), 48.0, 33.2 (d, *J* = 20.3 Hz). **MS$^-$** 342.1 [M - H]$^-$ **Alpha D** $[\alpha]_D^{25}$ = -21.4 (c = 1.0, MeOH)

Spectroscopic data was consistent with literature reports.[12]

##### *tert*-Butyl N-(*tert*-butoxycarbonyl)-D-homoserinate (Boc-D-hSe-O*t*Bu)

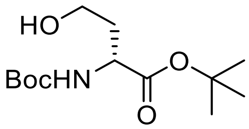

Under argon, Boc-D-Asp-O*t*Bu (2.03 g, 7.02 mmol) was dissolved in dry THF (70 mL) and the solution was cooled to 0 °C. Isobutyl chloroformate (2.73 mL, 21.1 mmol) and DIPEA (1.83 mL, 10.5 mmol) were added and the resulting mixture was allowed to stir at 0 °C for 1 h before warming to RT and stirring for a further 30 min. Sodium borohydride (1.86 g, 49.1 mmol) was added slowly at 0 °C followed by $H_2O$ (15 mL) under a stream of nitrogen. The mixture was left to stir overnight at RT and then acidified with 1 M HCl until pH 2. The organic later was separated and the aqueous phase was extracted with EtOAc (3 x 50 mL). The combined organic layers was washed with sat. aq. $NaHCO_3$ (2 x 50 mL) and brine (50 mL) then dried over $MgSO_4$, filtered and concentrated under reduced pressure. Combiflash purification by silica gel column chromatography (0 to 15% EtOAc in pet. ether over 7 min then 15 to 50% EtOAc in pet. ether over 11 min) gave Boc-D-hSe(homoserine)-O*t*Bu as a colourless oil (1.83 g, 95%).

$C_{13}H_{25}NO_5$ (275.4 g/mol):
**$^1$H NMR** (600 MHz, $CDCl_3$) δ 5.36 (d, *J* = 7.9 Hz, 1H), 4.31 (ddd, *J* = 11.1, 7.9, 3.7 Hz, 1H), 3.69 – 3.58 (m, 2H), 3.48 (d, *J* = 9.0, 4.8 Hz, 1H), 2.10 (ddt, *J* = 14.4, 9.5, 3.9 Hz, 1H), 1.55 – 1.49 (m, 1H), 1.44 (s, 9H), 1.41 (s, 9H). **$^{13}$C NMR** (151 MHz, $CDCl_3$) δ 172.1, 156.7, 82.3, 80.4, 58.3, 51.0, 36.6, 28.3 (3C), 28.1 (3C). **MS$^+$** 276.2 $[M + H]^+$ **IR** (ATR): $\tilde{\upsilon}/cm^{-1}$ = 3379, 2978, 2933, 1693, 1505, 1456, 1392, 1366, 1247, 1150, 1050, 996, 961, 903, 866, 846, 781. **Alpha D** $[\alpha]_D^{25}$ = 36.0 (c = 1.0, EtOH)

##### *tert*-Butyl N-(*tert*-butoxycarbonyl)-O-tosyl-D-homoserinate (Boc-D-hSe(OTs)-O*t*Bu)

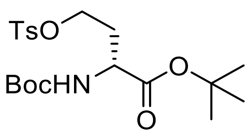


The following procedure was adapted from a known procedure.[12]

Under argon, Boc-D-Hse-O*t*Bu (825 mg, 3.00 mmol) was dissolved in dry DCM (9 mL) and the solution was cooled to 0 °C. $NEt_3$ (2.09 mL, 15.0 mmol), TsCl (1.14 g, 5.99 mmol) and DMAP (36.6 mg, 0.300 mmol) were added sequentially and the resulting mixture was allowed to stir at 0 °C for 15 min before warming to RT and stirring for a further 16 h. The mixture was then diluted with DCM (12 mL) and the organic phase was washed with $H_2O$ (2 x 20 mL) and brine (10 mL) then dried over $MgSO_4$, filtered and concentrated under reduced pressure. Combiflash purification by silica gel column chromatography (5 to 15% EtOAc in pet. ether over 15 min) gave Boc-D-Hse(OTs)-O*t*Bu as a white solid (1.11 mg, 86%).

$C_{20}H_{31}NO_7S$ (429.5 g/mol):
**$^1$H NMR** (600 MHz, $CDCl_3$) δ 7.72 (d, *J* = 8.4 Hz, 2H), 7.34 (d, *J* = 7.9 Hz, 2H), 4.95 (d, *J* = 7.6 Hz, 1H), 4.12 – 4.08 (m, 1H), 4.04 – 3.98 (m, 2H), 2.38 (s, 3H), 2.17 – 2.11 (m, 1H), 2.01 – 1.95 (m, 1H), 1.38 (s, 9H), 1.34 (s, 9H). **$^{13}$C NMR** (151 MHz, $CDCl_3$) δ 170.7, 155.3, 145.0, 132.8, 130.0 (2C), 128.2 (2C), 82.8, 80.0, 66.7, 51.2, 31.8, 28.4 (3C), 28.0 (3C), 21.8. **MS$^+$** 430.2 $[M + H]^+$, 452.3 $[M + Na]^+$ **IR** (ATR): $\tilde{\upsilon}$/cm$^{-1}$ = 3380, 2982, 2361, 2341, 1726, 1599, 1500, 1460, 1448, 1393, 1366, 1347, 1335, 1290, 1256, 1230, 1190, 1171, 1147, 1098, 1076, 1049, 1026, 988, 960, 935, 867, 848, 812, 797, 759, 729, 706. **Alpha D** $[\alpha]_D^{25}$ = 16.4 (c = 1.0, MeOH)

*tert*-Butyl (R)-2-(*tert*-butoxycarbonyl)amino)-4-fluorobutanoate (Boc-D-Fha-O*t*Bu)

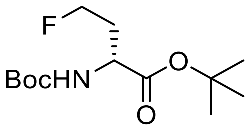


The following procedure was adapted from a known procedure.[12]

Under argon, Boc-D-Hse(OTs)-O*t*Bu (850 mg, 1.98 mmol) was dissolved in anhydrous *t*BuOH (20 mL). TBAF·3$H_2O$ (1.87 g, 5.94 mmol) was added and the resulting mixture was allowed to stir at 70 °C for 3 h before quenching with sat. aq. $NaHCO_3$ (20 mL). The mixture was extracted with DCM (3 x 30 mL) and the combined organic phases was dried over $MgSO_4$, filtered and concentrated under reduced pressure. Combiflash purification by silica gel column chromatography (0 to 20% EtOAc in pet. ether over 13 min) gave Boc-D-Fha-O*t*Bu as a white solid (437 mg, 80%).

$C_{13}H_{24}FNO_4$ (277.3 g/mol):
**TLC:** $R_f$ = 0.6 (20% EtOAc/pet. ether) **$^1$H NMR** (600 MHz, $CDCl_3$) δ 5.21 (d, *J* = 5.8 Hz, 1H), 4.62 – 4.55 (m, 1H), 4.54 – 4.47 (m, 1H), 4.30 (q, *J* = 6.6 Hz, 1H), 2.26 – 2.17 (m, 1H), 2.14 – 2.04 (m, 1H), 1.47 (s, 9H), 1.44 (s, 9H). **$^{19}$F NMR** (565 MHz, $CDCl_3$) δ -219.4 (tt, *J* = 47.1, 25.9 Hz). **$^{13}$C NMR** (151 MHz, $CDCl_3$) δ 171.3, 155.5, 82.4, 80.7 (d, *J* = 165.3 Hz), 80.7, 51.3, 33.5 (d, *J* = 19.8 Hz), 28.5 (3C), 28.1 (3C). **MS$^+$** 278.1 $[M + H]^+$

##### (R)-2-amino-4-fluorobutanoic acid trifluoroacetate (D-Fha•$CF_3CO_2H$)

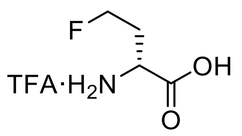


A solution of Boc-D-Fha-O*t*Bu (386 mg, 1.39 mmol) in $H_2O$ (1.4 mL) was cooled to 0 °C. TFA (12.5 mL) was added and the reaction was allowed to stir at RT for 4 h. The solution was concentrated under a stream of nitrogen and dried azeotropically under reduced pressure with toluene to give the TFA salt of D-Fha-OH as a colourless solid (quant. yield). The unprotected amino acid was used directly in the next step.

(R)-2-((((9H-fluoren-9-yl)methoxy)carbonyl)amino)-4-fluorobutanoic acid (Fmoc-D-Fha)

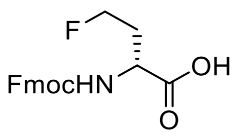


FmocOSu (493 mg, 1.46 mmol) was added to a mixture of crude D-Fha-OH (1.39 mmol) in THF (10 mL) and sat. aq. $NaHCO_3$ (5 mL). The resulting mixture was stirred at RT for 16 h and then diluted with $H_2O$ (10 mL). The aqueous layer was washed with $Et_2O$ (2 x 25 mL) and acidified with 1 M HCl until pH 1-2. The aqueous phased was extracted with EtOAc (3 x 30 mL). The combined organic layers was dried over $MgSO_4$, filtered and concentrated under reduced pressure to give Fmoc-D-Fha-OH as a white solid (404 mg, 81% by $^1$H NMR) without further purification.

$C_{19}H_{18}FNO_4$ (343.35 g/mol):
**$^1$H NMR** (600 MHz, Acetone-$d_6$) δ 7.86 (d, *J* = 7.6 Hz, 2H), 7.72 (t, *J* = 6.7 Hz, 2H), 7.41 (t, *J* = 7.5 Hz, 2H), 7.32 (td, *J* = 7.5, 1.2 Hz, 2H), 6.84 (d, *J* = 8.5 Hz, 1H), 6.83 (d, *J* = 8.6, 1H), 4.68 – 4.62 (m, 1H), 4.60 – 4.54 (m, 1H), 4.43 – 4.38 (m, 1H), 4.37 – 4.33 (m, 2H), 4.25 (t, *J* = 7.2 Hz, 1H), 2.39 – 2.30 (m, 1H), 2.17 – 2.09 (m, 1H). **$^{19}$F NMR** (600 MHz, Acetone-$d_6$) δ -221.5 (m). **$^{13}$C NMR** (151 MHz, Acetone-$d_6$) δ 173.5, 157.1, 145.1 (2C), 142.1 (2C), 128.5 (2C), 127.9 (2C), 126.1 (2C), 120.8 (2C), 81.2 (d, *J* = 164.0 Hz), 67.2, 51.3 (d, *J* = 4.9 Hz), 48.0, 33.2 (d, *J* = 20.4 Hz). **IR** (ATR): $\tilde{\upsilon}$/cm$^{-1}$ = 3728, 2710, 3628, 3594, 3313, 2360, 2341, 1692, 1533, 1449, 1430, 1393, 1275, 1248, 1105, 1084, 1063, 1043, 995, 901, 448, 757, 739. **MS$^-$** 342.3 [M - H]$^-$ **Alpha D** $[\alpha]_D^{25}$ = 19.4 (c = 1.0, MeOH)

#### Diethyl 2-acetamido-2-(2,2-difluoroethyl)malonate

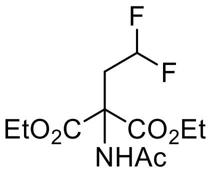


The following procedure was adapted from a known procedure.[13]

Under argon, diethyl acetamidomalonate (1.09 g, 5.02 mmol) was dissolved in anhydrous THF (10 mL). KO*t*Bu (563 mg, 5.02 mmol) was added under vigorous stirring and the resulting mixture was refluxed at 75 °C for 1.5 h. Difluoroethyl triflate (0.8 mL, 6.00 mmol) was added dropwise to the refluxing suspension and the resulting solution was allowed to stir for a further 3 h. The solution was concentrated and the residue was dissolved in EtOAc (25 mL), washed with 0.5 M HCl (2 x 10 mL), $H_2O$ (2 x 10 mL), sat. aq. $NaHCO_3$ (2 x 10 mL) and brine (10 mL). The organic phase was dried over $MgSO_4$, filtered and concentrated under reduced pressure. The oil was dissolved in $Et_2O$ (10 mL) and kept at -20 °C overnight. The mixture was filtered to give a white solid (386 mg, 27%).

$C_{11}H_{17}F_2NO_5$ (281.3 g/mol):
**$^1$H NMR** (400 MHz, $CDCl_3$) δ 6.89 (br s, 1H), 5.84 (tt, *J* = 55.8, 4.7 Hz, 1H), 4.26 (q, *J* = 7.1 Hz, 4H), 2.98 (td, *J* = 16.6, 4.7 Hz, 2H), 2.06 (s, 3H), 1.27 (t, *J* = 7.1 Hz, 6H). **$^{19}$F NMR** (376 MHz, $CDCl_3$) δ -116.4 (dt, *J* = 56, 17 Hz). **$^{13}$C NMR** (151 MHz, $CDCl_3$) δ 169.9, 167.2 (2C), 115.3 (t, *J* = 238.9 Hz), 63.3 (2C), 62.9 (t, *J* = 5.5 Hz), 37.0 (t, *J* = 22.6 Hz), 23.1, 14.0 (2C). **MS$^+$** 281.7 $[M + H]^+$

Spectroscopic data was consistent with literature reports.[13]

#### 2-amino-4,4-difluorobutanoic acid hydrochloride (L/D-*b*Fha•HCl)

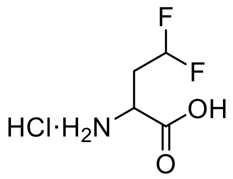

The following procedure was adapted from a known procedure.[13]

The malonate above (350 mg, 1.31 mmol) was dissolved in 6 M HCl (10 mL) and refluxed for 16 h. The aqueous phase was washed with $Et_2O$ (10 mL) and then dried azeotropically under reduced pressure with toluene to give the HCl salt of L/D-*b*Fha-OH as a colourless solid (quant. yield). The unprotected amino acid was used directly in the next step.

2-((((9H-fluoren-9-yl)methoxy)carbonyl)amino)-4,4-difluorobutanoic acid (Fmoc-L/D-*b*Fha)

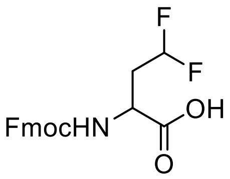


FmocOSu (464 mg, 1.38 mmol) was added to a mixture of crude L/D-*b*Fha-OH (1.31 mmol) in THF (10 mL) and sat. aq. $NaHCO_3$ (5 mL). The resulting mixture was stirred at RT for 16 h and then diluted with $H_2O$ (10 mL). The aqueous layer was washed with $Et_2O$ (2 x 25 mL) and acidified with 1 M HCl until pH 1-2. The aqueous phased was extracted with EtOAc (3 x 30 mL). The combined organic layers was dried over $MgSO_4$, filtered and concentrated under reduced pressure to give Fmoc-L/D-*b*Fha-OH as a white solid (351 mg, 74%) without further purification.

$C_{19}H_{17}F_2NO_4$ (361.3 g/mol):
**$^1$H NMR** (600 MHz, Acetone-$d_6$) δ 7.86 (dd, *J* = 7.7, 1.1 Hz, 2H), 7.71 (d, *J* = 7.5, 2.8 Hz, 2H), 7.41 (td, *J* = 7.5, 0.9 Hz, 2H), 7.32 (td, *J* = 7.4, 1.1 Hz, 2H), 6.98 (d, *J* = 8.6 Hz, 1H), 6.13 (tdd, *J* = 56.4, 5.5, 4.1 Hz, 1H), 4.46 (td, *J* = 9.0, 4.7 Hz, 1H), 4.37 (d, *J* = 7.2 Hz, 2H), 4.25 (t, *J* = 7.2 Hz, 1H), 2.54 – 2.45 (m, 1H), 2.43 – 2.31 (m, 1H). **$^{19}$F NMR** (565 MHz, Acetone-$d_6$) δ -116.1 (dt, *J* = 56, 17 Hz). **$^{13}$C NMR** (151 MHz, Acetone-$d_6$) δ 172.5, 157.0, 145.1 (2C), 142.1 (2C), 128.5 (2C), 127.9 (2C), 126.1 (2C), 120.8 (2C), 116.9 (t, *J* = 237.6 Hz), 67.3, 50.1 (t, *J* = 6.5 Hz), 48.0, 36.8 (t, *J* = 22.6 Hz). **HRMS** (ESI (-), MeOH): (*m/z*) calc. for $C_{19}H_{17}F_2NO_4$: 360.1053 $[M - H]^-$; found: 360.1041

**SPPS following Fmoc strategy**

*Preloading 2-chloro-trityl chloride resin:* 2-Chloro-trityl chloride resin (1-1.5 mmol/g loading) was swollen in dry DCM for 10 min and then preactivated with 2% $SOCl_2$ in DCM (2 x 4 mL). The resin was washed with DCM (7 × 4 mL), 5% DIPEA in DCM for 30 min then DCM (3 × 5 mL). A solution of Fmoc-aa-OH (1 equiv. relative to resin functionalisation) and DIPEA (2.0 equiv. relative to resin functionalisation) in DCM (final concentration 0.1 M of amino acid) was added and the resin shaken at rt for 1 h. The resin was washed with (5 × 4 mL). The resin was then treated with a solution of DCM/DIPEA/MeOH(17:2:1 v/v/v, 3 mL) for 30 min and washed with DCM (5 × 4 mL) and DMF (5 × 4 mL). The resin was subsequently submitted to iterative peptide assembly (Fmoc-SPPS).

*Deprotection:* The resin was treated with 20% piperidine/DMF (2 x 4 mL, 5 min) and washed with DMF (5 × 4 mL), $CH_2Cl_2$ (5 × 4 mL) and DMF (5 × 4 mL).

*General amino acid coupling:* A solution of protected amino acid (4 equiv.), PyBOP (4 equiv.) and NMM (8 equiv.) in DMF (final concentration 0.1 M) was added to the resin. After 1 h, the resin was washed with DMF (5 × 4 mL), DCM (5 × 4 mL) and DMF (5 × 4 mL).

*Capping:* Acetic anhydride/pyridine (1:9, v/v) was added to the resin (4 mL). After 10 min the resin was washed with $CH_2Cl_2$ (5 × 4 mL), DMF (5 × 4 mL), $CH_2Cl_2$ (5 × 4 mL) and DMF (5 × 4 mL).

*Cleavage:* A mixture of TFA, triisopropylsilane (TIS) and water (90:5:5, v/v/v) was added to the resin. After 2 h, the resin was washed with TFA (3 × 2 mL).

*Work-up:* The combined cleavage and TFA wash solutions were concentrated under a stream of nitrogen either to dryness or to < 2 mL. 40 mL of diethyl ether was added to precipitate the peptide, the suspension centrifuged at 4000 rpm for 10 min at 4 °C. The supernatant was removed and the pellet or dry residue was then dissolved in water containing 0.1% TFA and minimal amount of MeCN, filtered, purified by preparative HPLC and analyzed by LC-MS and ESI mass spectrometry.

Preparative HPLC purifications were performed on Shimadzu 2020 with a LCM20AD pump, SPD-20A UV/Vis operating at 220 nm at a flow rate of 7 mL/min) using Kinetex C18 100 Å column (5 μm, 21.2 x 250 mm).

A solvent system of $H_2O$ + 0.1% TFA/MeCN + 0.1% TFA was used with the gradient elution as follows:

| Step | Time / min | % $H_2O$ + 0.1% TFA | % MeCN + 0.1% TFA |
|---|---|---|---|
| 0 | 0 | 100 | 0 |
| 1 | 3 | 100 | 0 |
| 2 | 28 | 0 | 100 |
| 3 | 35 | 0 | 100 |
| 4 | 36 | 100 | 0 |
| 5 | 40 | 100 | 0 |

**Coupling conditions for L-Fmoc-Fha-OH, D-Fmoc-Fha-OH and L/D-Fmoc-*b*Fha-OH**

A solution of the Fmoc-aa-OH (2 equiv), HATU (1.8 equiv) and DIPEA (4 equiv) in DMF (final concentration 0.1 M) was added to the resin (1.0 equiv) and shaken. After 18 h, the resin was washed with DMF (5 x 4 mL), $CH_2Cl_2$ (5 x 4 mL), and DMF (5 x 4 mL). Synthesis of the desired peptide was completed using iterative Fmoc-SPPS.

H-A(L-Fha)ENLYFQ-OH (L-Fha reference peptide)

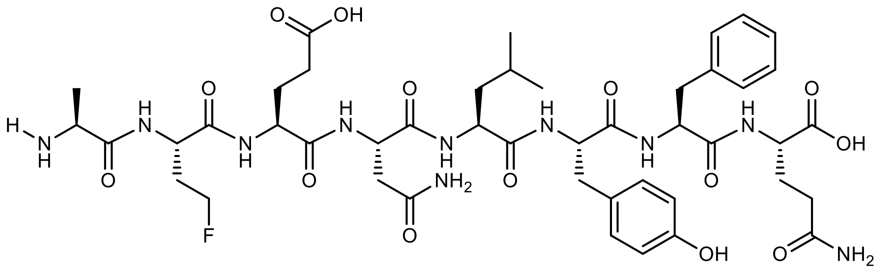

L-Fha peptide was prepared following the Fmoc-strategy SPPS and purified by preparative reverse phase HPLC as outlined in the general procedure.

HPLC chromatogram and MS analysis:

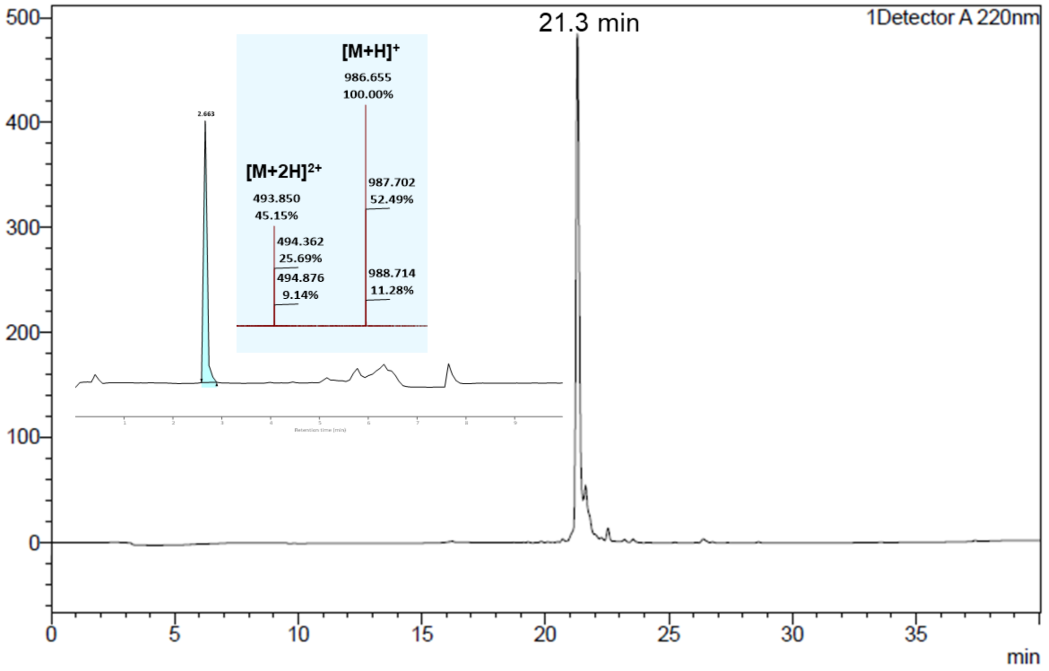


**MS+** 986.7 [M + H]$^+$, 493.9 [M + 2H]$^{2+}$

H-A(L/D-*b*Fha)ENLYFQ-OH (L/D-*b*Fha reference peptide)

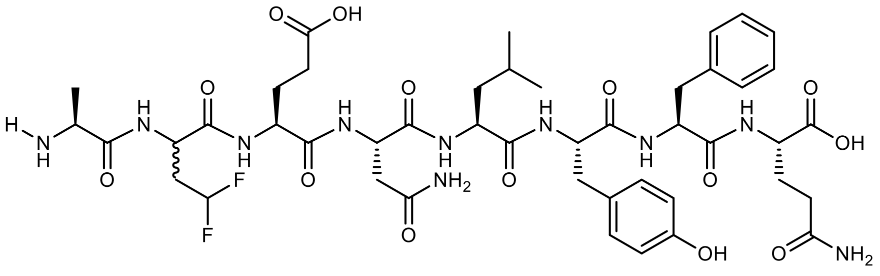


L/D-*b*Fha peptide was prepared following the Fmoc-strategy SPPS and purified by preparative reverse phase HPLC as outlined in the general procedure. Under these HPLC elution conditions, the peptide epimers were successfully separated.

HPLC chromatogram:

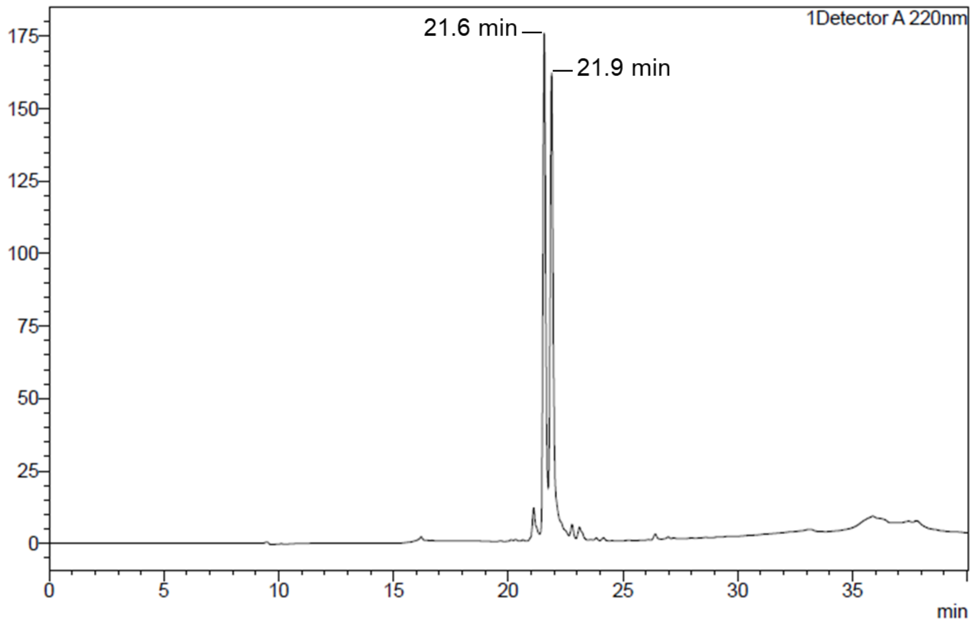


**MS⁺** 1004.6 $[M + H]^+$, 502.7 $[M + 2H]^{2+}$

MS analysis of peptide peak at 21.6 min:

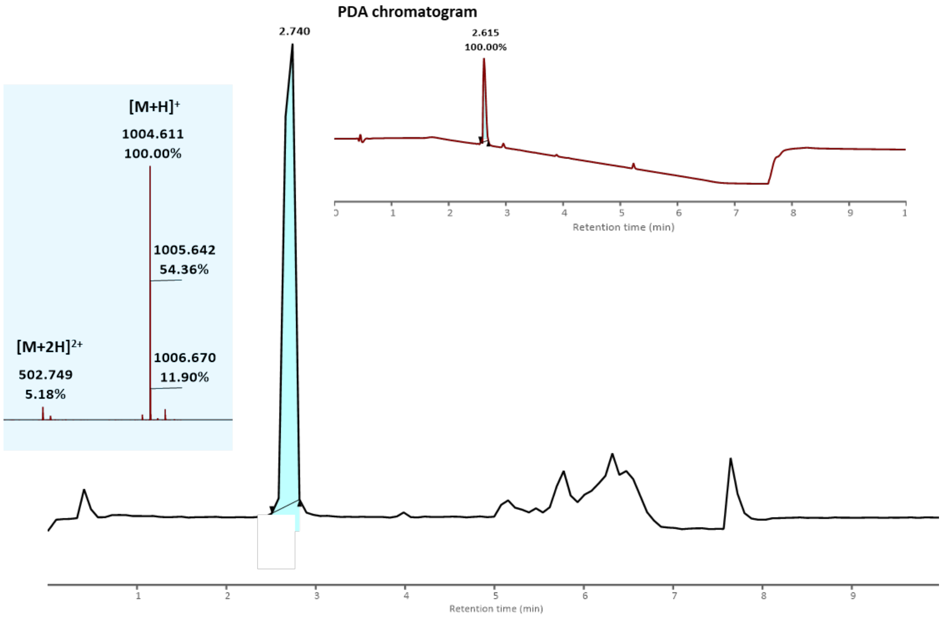


MS analysis of peptide peak at 21.9 min:

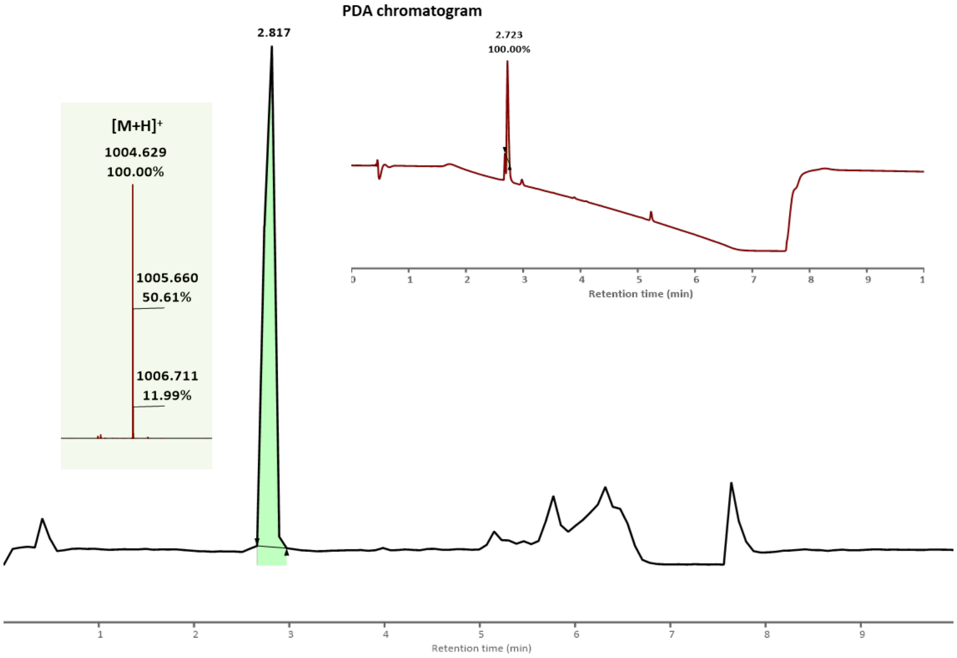

#### 2. Protein Production, Modification and Characterisation

##### 2.1 Intact Protein Mass Spectrometry General Methods and Data Analysis

Protein reactions were monitored by LC-MS analysis of the crude reaction mixtures. Intact protein mass spectrometry was performed on Waters QTof mass spectrometers (Xevo G2-XS or Xevo G2-S) coupled to Acquity UHPLC systems. A Thermo Proswift column (250 mm x 4.6 mm x 5 µm) with a flow rate of 0.300 mL $min^{-1}$ and a solvent system of water + 0.1% formic acid (solvent A) and acetonitrile + 0.1% formic acid (solvent B) were employed for a total run time of 10 min with gradient elution as follows:

| Step | Time / min | %Solvent A | %Solvent B |
|---|---|---|---|
| 0 | 0 | 95 | 5 |
| 1 | 1.0 | 95 | 5 |
| 2 | 7.0 | 5 | 95 |
| 3 | 8.0 | 5 | 95 |
| 4 | 8.1 | 95 | 5 |
| 5 | 10.0 | 95 | 5 |

Nitrogen was used as the desolvation (650 L/h) and cone (30 L/h) gas with the following instrument parameters: capillary voltage 3 kV, cone voltage 20 V, source temperature 100 ºC, desolvation temperature 400 ºC and collision energy 6.0 eV.

The LC-MS chromatogram is based on total ion count. As the protein modification is miniscule in comparison to the entire protein, the retention times of different protein products will not vary. Hence, a protein peak on the chromatogram accounts for all protein species in the reaction i.e., starting material, desired product and unwanted by-products. Furthermore, appreciable changes in ionisation between different protein products are not expected. Hence, their relative intensities are used to calculate % conversion.

The raw spectra of multiple charged ion series were deconvoluted using Waters MassLynx software (v4.1) and its maximum entropy (MaxEnt1) function with the following set-up: resolution 1.00 Da/channel, width at half height (protein-dependent i.e. 0.40 Da for histones, 0.75 Da for NFL), 12 minimum intensity ratios 33% left and right and an iteration to convergence. The deconvolution mass range is protein dependent i.e. 10000 – 20000 Da for *X.l.* Histone H3, 10000 – 25000 Da for human Histone eH3.1 and *X.l.* Histone H3.NTEV, 50000 – 75000 Da for NFL. Reaction conversions were calculated as the relative intensity ratio of the peak of interest against the sum for all peaks. On histones, ~10% baseline methionine oxidation +16 Da adducts are often present during preparation and storage. These adducts were combined with their corresponding starting material or product.

#### 2.2 Tandem Mass Spectrometry

LC-MS/MS was used to identify reaction products and confirm site of modifications on histones and map phosphorylation sites on NfL.

**Histones: In-solution proteolytic digest**

The crude protein reaction was desalted and purified from the reaction components with a PD MiniTrap G-25 column (pre-equilibrated with 50 mM Tris-HCl, 5 mM $CaCl_2$, pH 7.7 for ArgC and chymotrypsin digestion or 50 mM Tris-HCl, pH 8.0 for LysC digestion). After loading onto the desalting column, elution with the first 200 µL of buffer was discarded. Subsequent elution with 500 µL of buffer was collected (50-60% protein recovery). In preparation for ArgC and chymotrypsin digestion, EDTA was added to a final concentration of 2 mM.

For proteolysis using ArgC (Promega), the enzyme and an activation buffer were added following a 1:25 enzyme:protein ratio (*w/w*) in a solution with 50 mM Tris-HCl, 5 mM $CaCl_2$, 5 mM DTT final concentrations. For proteolysis using chymotrypsin (Roche) and LysC (Fujifilm Wako), an enzyme:protein ratio of 1:25 (*w/w*) were also used. All samples were incubated for 4 h at 37 °C, desalted by Oasis HLB cartridges (Waters) and dried in a speed-vac before resuspension in $H_2O$ with 5% FA and 5% DMSO.

Data acquisition

Samples were subjected to LC-MS/MS using an UltiMate 3000 nanoUHPLC system (Thermo Fisher Scientific) coupled to an Orbitrap Fusion Lumos (Thermo Fisher Scientific). The peptides were trapped on a C18 PepMap100 pre-column (300 µm i.d. x 5 mm, 100 Å, Thermo Fisher Scientific) using solvent A (0.1% formic acid in water), then separated on an in-house packed analytical column (75 µm i.d. x 50 cm in-house packed with ReproSil Gold 120 C18, 1.9 µm, Dr. Maisch GmbH) with a gradient of 8% to 38% B (0.1% formic acid in acetonitrile) over 15 minutes at a flow rate of 200 nL/min. Full scan MS spectra were acquired in the Orbitrap (scan range 350-1400 m/z, resolution 60000, AGC target 1200000). The 20 most intense peaks were selected for HCD fragmentation (30% of normalised collision energy, AGC target 50000, Orbitrap readout with resolution

7500), and ETciD (using calibrated charge-dependent parameters, AGC target 10000, 35% normalised collision energy of supplementary CID, rapid mode ion trap readout).

Data analysis

Spectra were searched using FragPipe (v18.0) MSFragger 3.5[14] with standard 'open' search settings using a bespoke .FASTA file containing sequences of pertinent and potential contaminant proteins. Data was filtered using the inbuilt tools within FragPipe to an FDR of below 1%. Modified peptides were discerned by filtering the resulting dataset using the expected changes in mass caused by each modification. Spectra were visualised using Proteomics Data Viewer[15].

**Neurofilament light chain: On-beads proteolytic digest**

Proteolytic digestion was performed using an SP3 protocol[16]. Briefly, NfL in PBS, 2 mM DTT, 4 M urea was reduced with 10 mM tris(2-carboxyethyl)phosphine and alkylated with 30 mM 2-chloroacetamide at room temperature for 30 min in the dark. The mixture was added to the bead solution (Speed Bead Magnetic Carboxylate Modified Particles, Cytiva 45152105050250), diluted using 100% ACN to the final concentration 77% ACN and incubated for 30 minutes at room temperature on a magnetic rack. The mixture was washed first with 80% ethanol five times and then with 100% ACN five times; the supernatant was discarded in each step. Trypsin/LysC solution in ammonium bicarbonate was added to match the 1:40 enzyme:protein ratio (w/w, for either enzyme), and the sample was incubated overnight at 37 °C. The supernatant containing products of proteolysis was aspirated and diluted in in $H_2O$ with 5% FA and 5% DMSO for LC-MS analysis.

Data acquisition

Samples were subjected to LC-MS/MS using an UltiMate 3000 nanoUHPLC system (Thermo Fisher Scientific) coupled to an Orbitrap Fusion Lumos (Thermo Fisher Scientific). The peptides were trapped on a C18 PepMap100 pre-column (300 µm i.d. x 5 mm, 100 Å, Thermo Fisher Scientific) using solvent A (0.1% formic acid in water), then separated on an in-house packed analytical column (50 µm i.d. x 50 cm in-house packed with ReproSil Gold 120 C18, 1.9 µm, Dr. Maisch GmbH) with a gradient of 5% to 40% B (0.1% formic acid in acetonitrile) over 15 minutes at a flow rate of 100 nL/min. Full scan MS spectra were acquired in the Orbitrap (scan range 350-2000 m/z, resolution 60000, AGC target 1200000). The 20 most intense peaks were selected for HCD fragmentation (30% of normalised collision energy, AGC target 50000, Orbitrap readout with resolution 7500), and ETciD (using calibrated charge-dependent parameters, AGC target 10000, 35% normalised collision energy of supplementary CID, rapid mode ion trap readout).

Data analysis

Spectra were searched using Andromeda search engine implemented in MaxQuant (v2.3.0)[17] and a bespoke .FASTA file containing sequences of pertinent and potential contaminant proteins, with phosphorylation at STY set as variable modification and otherwise standard settings. Data was filtered using the inbuilt tools within MaxQuant to an FDR of below 1%. Spectra were visualised using Proteomics Data Viewer[15].

#### 2.3 Biological instruments

Proteins were purified using an Äkta FPLC System UPC-900 (GE Healthcare, UK). Gel electrophoresis was performed using Invitrogen NuPAGE 10%, 12% or 4-12% Bis-Tris gels, Novex MiniCell tanks, and a BioRad PowerPac controller. Western blotting was performed using an iBlot gel transfer device from Thermo-Fisher. Colorimetric method (for alkaline phosphatase-conjugated antibody) was performed using Sigma BCIP/NBT Liquid Substrate System (catalogue number B1911). Chemiluminescent analysis (for horseradish peroxidase (HRP)-coupled antibody) was carried out using Thermo Scientific SuperSignal West Pico Plus Chemiluminescent substrate (catalogue number 34580). Nucleotide sequences were confirmed by the Source Bioscience DNA Sanger sequencing services based at Oxford University.

#### 2.4 Protein expression and purification

**Histone H3**

Expression and purification:

All human histones eH3.1 (e refers to the C-terminal FLAG-HA dual epitope tag; native residues Cys96 and Cys110 have been mutated to Ala) and *Xenopus laevis* histone variants (native residue Cys110 is mutated to Ala) were expressed and isolated as previously described.[4] The histone encoding plasmid was transformed into BL21 (DE3) pLysS *E.coli* cells (1 µL of plasmid per 20 - 50 µL of Competent Cells). The mixture was incubated in ice for 10 - 20 min. The cells were then heat-shocked for 45 - 50 s in a water bath at exactly 42 °C without shaking before returning back to ice for a further 2 min. Cold SOC medium (4 °C, 500 µl) was then added and the suspension was incubated for 60 min at 37 °C with shaking (approximately 250 rpm). 200 µL of the cells were added onto pre-warmed agar plates containing chloramphenicol and ampicillin to begin incubation at 37 °C for 12 - 14 h. Successful colonies were selected after overnight growth and used to inoculate 4 x 10 mL LB medium with the same antibiotics. This culture was left to grow at 37 °C overnight. 10 mL of starter culture was then added to 4 x 1 L of 2xYT media containing the same antibiotics and grown at 37 °C until $OD_{600}$ = 0.4 - 0.6 (around 2.5 h, measured against 1 mL of blank medium). IPTG was added to a final concentration of 0.5 mM and the flask was shaken at 37 °C for 2 h. Cells expressing histones are prone to lysis and longer expression time is detrimental on the yield. Cells were harvested by centrifugation for 15 min at 8k rpm at 4 °C and the pellet was resuspended into a 5-fold v/w wash buffer (50 mM Tris pH 7.5, 100 mM NaCl with protease inhibitors). Suspensions were flash-frozen and stored at -80 °C until lysis. Frozen cells were thawed in a water bath. When the solution was viscous, DNase was added, sonicated 5 × at 40% amplitude in 30 s bursts and then centrifuged at 13k rpm for 20 min at 4 °C. The supernatant was discarded, and the pellet was resuspended in 40 mL wash buffer. Sonication was repeated twice at 40% amplitude for 30 s, and the suspension centrifuged at 13k rpm for 20 min after each wash. After addition of 1 mL DMSO, the pellet was mixed with a spatula and left at RT for 10 min. 10 mL unfolding buffer (7M Gdn·HCl, 20 mM Tris pH 7.5, 10 mM DTT) was then added, and shaken for 1 h at RT, after which it was centrifuged for 20 min at 13k rpm at RT. The supernatant was concentrated to 1.5 mL (from a 2 L culture) and then filtered before loading onto an S200 size-exclusion column pre-equilibrated with 1 CV SAU-100 (7 M urea, 20 mM NaOAc, pH 5.2, 100 mM NaCl, 1 mM EDTA, 10 mM DTT). The histone was eluted with SAU-100 buffer and analysed by SDS-PAGE. Fractions containing the desired histones were pooled. Further purification by cation exchange chromatography with HiTrap SP 5 mL may be required. This was performed using a linear gradient of 0-100% SAU-1000 buffer (SAU-100 with 1000 mM NaCl final concentration). Pure fractions were pooled, dialysed thrice against water containing 2 mM βME and lyophilised to produce a fluffy white solid.

Protein sequences:

*Xenopus laevis* (*X.l.*) Histone H3.NTEV-Cys2

Mutations: R2C, C110A, TEV sequence after R2C, before T3

```
        10         20         30         40         50         60
ACENLYFQGT KQTARKSTGG KAPRKQLATK AARKSAPATG GVKKPHRYRP GTVALREIRR

        70         80         90        100        110        120
YQKSTELLIR KLPFQRLVRE IAQDFKTDLR FQSSAVMALQ EASEAYLVAL FEDTNLAAIH

       130        140
AKRVTIMPKD IQLARRIRGE RA
```

Calculated molecular weight: 16038 g $mol^{-1}$

Extinction coefficient ε: 5960 $M^{-1}$ $cm^{-1}$

*Xenopus laevis* (*X.l.*) Histone H3-Cys9

Mutations: K9C, C110A

```
        10         20         30         40         50         60
ARTKQTARCS TGGKAPRKQL ATKAARKSAP ATGGVKKPHR YRPGTVALRE IRRYQKSTEL

        70         80         90        100        110        120
LIRKLPFQRL VREIAQDFKT DLRFQSSAVM ALQEASEAYL VALFEDTNLA AIHAKRVTIM

       130
PKDIQLARRI RGERA
```

Calculated molecular weight: 15214 g $mol^{-1}$

Extinction coefficient ε: 4470 $M^{-1}$ $cm^{-1}$

All other *X.l.* histone lysine to cysteine mutants used for this paper (i.e., Cys18, Cys27, Cys36) share the same molecular weight and extinction coefficient.

*Homo sapiens* (Human) Histone eH3.1-Cys4

Mutations: K4C, C96A, C110A, C-terminal FLAG-HA epitope tags (e)

```
        10         20         30         40         50         60
ARTCQTARKS TGGKAPRKQL ATKAARKSAP ATGGVKKPHR YRPGTVALRE IRRYQKSTEL

        70         80         90        100        110        120
LIRKLPFQRL VREIAQDFKT DLRFQSSAVM ALQEAAEAYL VGLFEDTNLA AIHAKRVTIM

       130        140        150
PKDIQLARRI RGERAGGDYK DDDDKSAAGG YPYDVPDYA
```

Calculated molecular weight: 17720 g $mol^{-1}$

Extinction coefficient ε: 10430 $M^{-1}$ $cm^{-1}$

All other human histone lysine to cysteine mutants used for this paper (i.e., Cys9, Cys18) share the same molecular weight and extinction coefficient.

**Mouse Neurofilament Light Chain**

Plasmid construction:

Original mouse NfL plasmid was purchased from Addgene (pmNfL; catalog No: 83127). The mNfL gene was inserted into pEGFP vector with a *pCMV* promoter for protein expression. 6x Histidine tag was inserted onto the C-terminus by site-directed mutagenesis method (using primers mNfLHis-F1: 5-GGTGGCTAAGAAGAAAGATGGCTCTCATCATCACCACCACCATTGAGCCCTATTCC CAACTATTC-3; mNfLHis-R1: 5-GAATAGTTGGGAATAG GGCTCAATGGTGGTGGTGATGATGAGAGCCATCTTTCTTCTTAGCCACC-3) to aid downstream protein purification. The sequence of the plasmid obtained pEGFP-mNfL-HisTag was confirmed by Sanger Sequencing. Then the target gene *mNfL-HisTag* was amplified from the pEGFP-NfL-HisTag plasmid by infusion primers (mNfLHisInsert-F1: 5-GGAGACCCAAGCTTGGTACCATGAGTTCGTTCGGCTACG-3; mNfLHisInsert-R1: 5-CATGCT CGAGCGGCCGCTCAATGGTGGTGGTGATGATGAG-3) and inserted into linearized pCDNA3.1 vector by ligation based on KpnI/NotI cleavage sites. The sequence of the resulting plasmid pCDNA3-mNfL-HisTag was verified by Sanger Sequencing. This plasmid was used for mNfL protein expression in HEK293T cells.

Protein (*top*) and DNA sequence (*bottom*) of mouse neurofilament light chain (mNfL) (NCBI Reference Sequence: NP_035040.1) are shown below.

**Protein Sequence of mouse NFL (NCBI Reference Sequence: NP_035040.1):**

MSSFGYDPYFSTSYKRRYVETPRVHISSVRSGYSTARSAYSSYSAPVSSSLSVRRSYSSSSGSLMPSLENLDLSQ
VAAISNDLKSIRTQEKAQLQDLNDRFASFIERVHELEQQNKVLEAELLVLRQKHSEPSRFRALYEQEIRDLRLAA
EDATNEKQALQGEREGLEETLRNLQARYEEEVLSREDAEGRLMEARKGADEAALARAELEKRIDSLMDEIAFLKK
VHEEEIAELQAQIQYAQISVEMDVSSKPDLSAALKDIRAQYEKLAAKNMQNAEEWFKSRFTVLTESAAKNTDAVR
AAKDEVSESRRLLKAKTLEIEA**C**RGMNEALEKQLQELEDKQNADISAMQDTINKLENELRSTKSEMARYLKEYQD
LLNVKMALDIEIAAYRKLLEGEETRLSFTSVGSITSGYSQSSQVFGRSAYSGLQSSSYLMSARSFPAYYTSHVQE
EQTEVEETIEATKAEEAKDEPPSEGEAEEEEKEKEEGEEEEGAEEEEAAKDESEDTKEEEEGGEGEEEDTKESEE
EEKKEESAGEEQVAKKKD

**DNA Sequence of mouse NFL:**

ATGAGTTCGTTCGGCTACGATCCGTACTTTTCGACCTCCTACAAGCGGCGCTATGTGGAGACGCCCCGGGTGCAC
ATCTCCAGCGTGCGCAGCGGCTACAGCACGGCGCGCTCCGCGTACTCCAGCTACTCCGCGCCGGTCTCCTCCTCG
CTGTCCGTGCGCCGCAGCTACTCGTCCAGCTCTGGCTCTTTGATGCCCAGCCTGGAGAATCTCGATCTGAGCCAG
GTAGCCGCCATCAGCAACGACCTCAAGTCTATCCGCACACAAGAGAAGGCACAGCTGCAGGACCTCAACGATCGC
TTCGCCAGCTTCATCGAGCGCGTGCACGAGCTGGAGCAGCAGAACAAGGTCCTGGAAGCCGAGCTGTTGGTGCTG
CGCCAGAAACACTCTGAGCCTTCCCGCTTCCGCGCCCTGTACGAGCAGGAGATCCGCGATCTGCGGCTGGCAGCG
GAAGACGCCACTAACGAGAAGCAGGCGCTGCAGGGCGAGCGCGAGGGGCTGGAGGAGACTCTGCGCAACCTGCAG
GCTCGCTATGAGGAAGAAGTGCTGAGCCGCGAGGACGCCGAGGGCCGGCTGATGGAAGCGCGCAAAGGTGCGGAT
GAGGCCGCGCTCGCCCGCGCCGAGCTGGAGAAGCGCATCGACAGCCTGATGGACGAGATAGCTTTCCTGAAGAAG
GTGCACGAGGAAGAGATCGCCGAGCTGCAGGCTCAGATCCAGTATGCTCAGATCTCCGTGGAGATGGACGTGTCC
TCCAAGCCCGACCTCTCCGCCGCTCTCAAGGACATCCGCGCTCAGTACGAGAAGCTGGCCGCCAAGAACATGCAG
AACGCCGAAGAGTGGTTCAAGAGCCGCTTCACCGTGCTAACCGAGAGCGCCGCCAAGAACACCGACGCTGTGCGC
GCTGCCAAGGACGAGGTGTCGGAAAGCCGCCGCCTGCTCAAGGCTAAGACCCTGGAGATCGAAGCC**TGC**CGGGGT
ATGAACGAAGCTCTGGAGAAGCAGCTGCAGGAGCTAGAGGACAAGCAGAATGCAGACATTAGCGCCATGCAGGAC
ACAATCAACAAACTGGAGAATGAGCTGAGAAGCACGAAGAGCGAGATGGCCAGGTACCTGAAGGAGTACCAGGAC
CTCCTCAATGTCAAGATGGCCTTGGACATCGAGATTGCAGCTTACAGAAAACTCTTGGAAGGCGAAGAGACCAGG
CTCAGTTTCACCAGCGTGGGTAGCATAACCAGCGGCTACTCTCAGAGCTCGCAGGTCTTCGGCCGTTCTGCTTAC
AGTGGCTTGCAGAGCAGCTCCTACTTGATGTCTGCTCGCTCTTTCCCAGCCTACTATACCAGCCACGTCCAGGAA
GAGCAGACAGAGGTCGAGGAGACCATTGAGGCTACGAAAGCTGAGGAGGCCAAGGATGAGCCCCCCTCTGAAGGA
GAAGCAGAAGAGGAGGAGAAGGAGAAAGAGGAGGGAGAGGAAGAGGAAGGCGCTGAGGAGGAAGAAGCTGCCAAG
GATGAGTCTGAAGACACAAAAGAAGAAGAAGAAGGTGGTGAGGGTGAGGAGGAAGACACCAAAGAATCTGAAGAG
GAAGAGAAGAAAGAGGAGAGTGCTGGAGAGGAGCAGGTGGCTAAGAAGAAAGATTGA

Protein and DNA sequences of mNfL following mutagenesis experiments to insert HisTag onto the C-terminus are shown below.

**Protein Sequence of NFL-His:**

MSSFGYDPYFSTSYKRRYVETPRVHISSVRSGYSTARSAYSSYSAPVSSSLSVRRSYSSSSGSLMPSLENLDLSQV
AAISNDLKSIRTQEKAQLQDLNDRFASFIERVHELEQQNKVLEAELLVLRQKHSEPSRFRALYEQEIRDLRLAAED
ATNEKQALQGEREGLEETLRNLQARYEEEVLSREDAEGRLMEARKGADEAALARAELEKRIDSLMDEIAFLKKVHE
EEIAELQAQIQYAQISVEMDVSSKPDLSAALKDIRAQYEKLAAKNMQNAEEWFKSRFTVLTESAAKNTDAVRAAKD
EVSESRRLLKAKTLEIEA**C**RGMNEALEKQLQELEDKQNADISAMQDTINKLENELRSTKSEMARYLKEYQDLLNVK
MALDIEIAAYRKLLEGEETRLSFTSVGSITSGYSQSSQVFGRSAYSGLQSSSYLMSARSFPAYYTSHVQEEQTEVE
ETIEATKAEEAKDEPPSEGEAEEEEKEKEEGEEEEGAEEEEAAKDESEDTKEEEEGGEGEEEDTKESEEEEKKEES
AGEEQVAKKKDGSHHHHHH

**DNA Sequence of NFL-His:**

ATGAGTTCGTTCGGCTACGATCCGTACTTTTCGACCTCCTACAAGCGGCGCTATGTGGAGACGCCCCGGGTGCACA
TCTCCAGCGTGCGCAGCGGCTACAGCACGGCGCGCTCCGCGTACTCCAGCTACTCCGCGCCGGTCTCCTCCTCGCT
GTCCGTGCGCCGCAGCTACTCGTCCAGCTCTGGCTCTTTGATGCCCAGCCTGGAGAATCTCGATCTGAGCCAGGTA
GCCGCCATCAGCAACGACCTCAAGTCTATCCGCACACAAGAGAAGGCACAGCTGCAGGACCTCAACGATCGCTTCG
CCAGCTTCATCGAGCGCGTGCACGAGCTGGAGCAGCAGAACAAGGTCCTGGAAGCCGAGCTGTTGGTGCTGCGCCA
GAAACACTCTGAGCCTTCCCGCTTCCGCGCCCTGTACGAGCAGGAGATCCGCGATCTGCGGCTGGCAGCGGAAGAC
GCCACTAACGAGAAGCAGGCGCTGCAGGGCGAGCGCGAGGGGCTGGAGGAGACTCTGCGCAACCTGCAGGCTCGCT
ATGAGGAAGAAGTGCTGAGCCGCGAGGACGCCGAGGGCCGGCTGATGGAAGCGCGCAAAGGTGCGGATGAGGCCGC
GCTCGCCCGCGCCGAGCTGGAGAAGCGCATCGACAGCCTGATGGACGAGATAGCTTTCCTGAAGAAGGTGCACGAG
GAAGAGATCGCCGAGCTGCAGGCTCAGATCCAGTATGCTCAGATCTCCGTGGAGATGGACGTGTCCTCCAAGCCCG
ACCTCTCCGCCGCTCTCAAGGACATCCGCGCTCAGTACGAGAAGCTGGCCGCCAAGAACATGCAGAACGCCGAAGA
GTGGTTCAAGAGCCGCTTCACCGTGCTAACCGAGAGCGCCGCCAAGAACACCGACGCTGTGCGCGCTGCCAAGGAC
GAGGTGTCGGAAAGCCGCCGCCTGCTCAAGGCTAAGACCCTGGAGATCGAAGCC**TGC**CGGGGTATGAACGAAGCTC
TGGAGAAGCAGCTGCAGGAGCTAGAGGACAAGCAGAATGCAGACATTAGCGCCATGCAGGACACAATCAACAAACT
GGAGAATGAGCTGAGAAGCACGAAGAGCGAGATGGCCAGGTACCTGAAGGAGTACCAGGACCTCCTCAATGTCAAG
ATGGCCTTGGACATCGAGATTGCAGCTTACAGAAAACTCTTGGAAGGCGAAGAGACCAGGCTCAGTTTCACCAGCG
TGGGTAGCATAACCAGCGGCTACTCTCAGAGCTCGCAGGTCTTCGGCCGTTCTGCTTACAGTGGCTTGCAGAGCAG
CTCCTACTTGATGTCTGCTCGCTCTTTCCCAGCCTACTATACCAGCCACGTCCAGGAAGAGCAGACAGAGGTCGAG
GAGACCATTGAGGCTACGAAAGCTGAGGAGGCCAAGGATGAGCCCCCCTCTGAAGGAGAAGCAGAAGAGGAGGAGA
AGGAGAAAGAGGAGGGAGAGGAAGAGGAAGGCGCTGAGGAGGAAGAAGCTGCCAAGGATGAGTCTGAAGACACAAA
AGAAGAAGAAGAAGGTGGTGAGGGTGAGGAGGAAGACACCAAAGAATCTGAAGAGGAAGAGAAGAAAGAGGAGAGT
GCTGGAGAGGAGCAGGTGGCTAAGAAGAAAGATGGCTCTCATCATCACCACCACCATTGA

The PCR reaction to insert the HisTag was run under conditions as shown below and the sequence corresponding to pEGFP-NfL-HisTag was subsequently confirmed by Sanger Sequencing using the universal primer EGFP-CR.

.

| PCR components | PCR reaction | | |
|---|---|---|---|
| 5 µL 10× reaction buffer | 95 °C | 60 s | 1× |
| 1 µL dNTPmix solution | 95 °C | 50 s | 19× |
| 3 µL Quick solution | 60 °C | 50 s | |
| 5 µL pmNfL-3 (2 ng/ul) | 68 °C | 10 mins | |
| 5 µL NfLHis-F1 (25 ng/ul) | 68 °C | 7 mins | 1× |
| 5 µL NfLHis-R1 (25 ng/ul) | 10 °C | ∞ | 1× |
| 25 µL $ddH_2O$ | | | |
| 1 µL Ultra pfu enzyme | | | |

Insertion into pCDNA3 vector:

The NfL-HisTag gene was truncated from the pmNfL plasmid and inserted into pCDNA3 vector by ligation based on KpnI/NotI cleavage sites. Standard PCR conditions shown below were employed to amplify the gene, and the PCR product was analysed on a 1% agarose DNA gel. The linearized pCDNA3 vector was obtained following digestion using KpnI-HF and NotI-HF of the plasmid Syn4-pcDNA3. Purification of NfL-HisTag and the linearized pCDNA3 vector were performed using agarose gel electrophoresis, and their concentrations were measured by a NanoSpectrophotometer.

| PCR components | PCR reaction | | |
|---|---|---|---|
| 1 µL NfLHis (2 ng/µL) | 98 °C | 40 s | 1× |
| 1.5 µL NfLHis-F1 (10 µM) | 98 °C | 15 s | 35× |
| 1.5 µL NfLHis-R1 (10 µM) | 55 °C | 30 s | |
| 21 µL $ddH_2O$ | 72 °C | 2 mins | |
| 25 µL CloneAmp HiFi PCR Premix | 72 °C | 5 mins | 1× |

Next, a typical ligation reaction was carried out under conditions which included 100 ng of pCDNA3 vector, 61.53 ng of the NfL-HisTag insert, 1.0 µL of infusion ligation enzyme and 1.24 µL of $ddH_2O$.The ligation product was subsequently transferred into XL10-gold competent cells and plated on LB agar plates (with 100 µg/mL ampicillin) for overnight culture at 37 °C. The single colonies were then selected for overnight small-scale liquid culture (10 ml LB broth with 100 µg/mL ampicillin). The plasmid (pCDNA3-NfL-His) for NfL-HisTag was extracted, and its sequence was confirmed by Sanger Sequencing using the universal primers CMVF-pCDNA3 and bGHR.

Protein expression and purification:

HEK293T cells were grown to 80% confluency in 20 x 150 mm dishes in DMEM supplemented with 10% FBS and 4 mM L-glutamine. A transfection solution was pre-made (40.8 mL OMEM, 1008 µg of plasmid at 875 ng $µL^{-1}$, 3.06 mL of PEI max at 1 mg $mL^{-1}$) by dissolving plasmid in OMEM and then adding PEI Max, before mixing and leaving to incubate at RT for 30 min. Hence, final concentrations in OMEM were 22.46 µg $mL^{-1}$ of plasmid and 68 µg $mL^{-1}$ of PEI Max (i.e. 3x plasmid). At this point the media was removed by aspiration and swapped to OMEM (10 mL) and then transfection solution (2.1 mL) was added. The cells were the incubated at 37 °C, 5% $CO_2$ for 6 h before the media was removed and replaced with 25 mL DMEM (10% FBS, 4 mM L-glutamine) and left to incubate for 4 days. The media was then removed and the cells were harvested by scrapping and washed with PBS and spun down and the cell pellet was stored -80 °C before purification.

The cell pellet from 38 x 150 mm dishes was suspended in 33 mL of lysis buffer (6 M Urea, 20 mM Tris·HCl, 500 mM NaCl, 0.5 mM TCEP, pH 7.8) and then lysed using a Dounce Homogeniser (type A, 7 mL at a time, last round 5 mL + 2 mL for wash) with 40 strokes. This was then centrifuged (13,000 rpm, 4 °C, 50 min) and then the supernatant was removed and added to pre-washed $Ni^{2+}$ NTA resin (10 mL, 50% by volume in EtOH, i.e. 5 mL total resin) and incubated with inversion at 4 °C for 3 h. The resin was then split between 4 columns and the protein was then eluted using an imidazole gradient (50 to 300 mM, in the same lysis buffer), and analysed by SDS-PAGE. Clean fractions were combined, concentrated with an Amicon 10 kDa MWCO centrifugal filter and then buffer exchanged into 4 M Gdn·HCl, 50 mM Tris·HCl, pH 8.0 using a pre-equilibrated PD MiniTrap G-25 column and concentrated to 15.8 mg $mL^{-1}$ for long-term storage in -80 °C. Below are the volumes and concentrations of imidazole used for the purification of WT mNfL and SDS-PAGE analysis of each fraction.

Stepwise elution gradient:

| Fraction | Imidazole (mM) | Volume per column (mL) | Total volume (mL) |
|---|---|---|---|
| FT | | | |
| W1 | 0 | 8.5 | 34 |
| W2 | 0 | 3.5 | 14 |
| W3 | 10 | 3.5 | 14 |
| W4 | 20 | 7 | 28 |
| W5 | 20 | 7 | 28 |
| E1 | 50 | 1.5 | 6 |
| E2 | 50 | 2 | 8 |
| E3 | 100 | 2 | 8 |
| E4 | 150 | 2 | 8 |
| E5 | 200 | 2 | 8 |
| E6 | 300 | 3 | 12 |

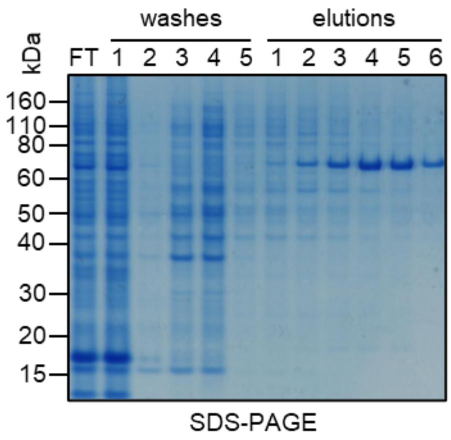

Protein sequence:

Mouse Neurofilament Light Chain (mNfL)

Mutations: C-terminal 6x Histidine tag

```
        10         20         30         40         50         60
MSSFGYDPYF STSYKRRYVE TPRVHISSVR SGYSTARSAY SSYSAPVSSS LSVRRSYSSS

        70         80         90        100        110        120
SGSLMPSLEN LDLSQVAAIS NDLKSIRTQE KAQLQDLNDR FASFIERVHE LEQQNKVLEA

       130        140        150        160        170        180
ELLVLRQKHS EPSRFRALYE QEIRDLRLAA EDATNEKQAL QGEREGLEET LRNLQARYEE

       190        200        210        220        230        240
EVLSREDAEG RLMEARKGAD EAALARAELE KRIDSLMDEI AFLKKVHEEE IAELQAQIQY

       250        260        270        280        290        300
AQISVEMDVS SKPDLSAALK DIRAQYEKLA AKNMQNAEEW FKSRFTVLTE SAAKNTDAVR

       310        320        330        340        350        360
AAKDEVSESR RLLKAKTLEI EACRGMNEAL EKQLQELEDK QNADISAMQD TINKLENELR

       370        380        390        400        410        420
STKSEMARYL KEYQDLLNVK MALDIEIAAY RKLLEGEETR LSFTSVGSIT SGYSQSSQVF

       430        440        450        460        470        480
GRSAYSGLQS SSYLMSARSF PAYYTSHVQE EQTEVEETIE ATKAEEAKDE PPSEGEAEEE

       490        500        510        520        530        540
EKEKEEGEEE EGAEEEEAAK DESEDTKEEE EGGEGEEEDT KESEEEEKKE ESAGEEQVAK

       550
KKDGSHHHHH H
```

Calculated molecular weight: 62344 g mol$^{-1}$ (without Met)

Extinction coefficient ε: 35300 M$^{-1}$ cm$^{-1}$

ESI analysis of WT mNfL:

Observed mass: 62385 and 62422 g mol$^{-1}$

Calculated mass for 1x carbamylation: 62387 g mol$^{-1}$

Calculated mass for 2x carbamylation: 62430 g mol$^{-1}$

Observed mass: 62467 and 62507 g mol$^{-1}$

Calculated mass for 1x carbamylation and 1x phosphorylation: 62467 g mol$^{-1}$

Calculated mass for 2x carbamylation and 1x phosphorylation: 62510 g $mol^{-1}$

Observed mass: 62547 and 62588 g $mol^{-1}$

Calculated mass for 1x carbamylation and 2x phosphorylation: 62547 g $mol^{-1}$

Calculated mass for 2x carbamylation and 2x phosphorylation: 62590 g $mol^{-1}$

Observed mass: 62627 g $mol^{-1}$

Calculated mass for 1x carbamylation and 3x phosphorylation: 62627 g $mol^{-1}$

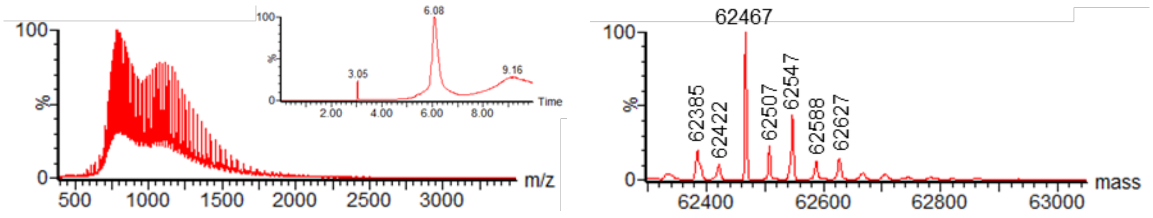


The levels and extent of phosphorylation differs between expression batches (see Supplementary Discussion).

**Human Neurofilament Light Chain**

Plasmid construction:

Plasmid containing *hNfL* gene was purchased from Addgene (phNfL; catalog No: 132589). 10x Histidine tag was inserted onto the N-terminus of *hNfL* by site-directed mutagenesis method using primers (HishNfL-F1: 5-GGAGCCACCGGCCGCCACCATGCATCATCACCACCACCATCACCACCATCAC AGTTCCTTCAGCTACGAGCCG-3; HishNfL-R1: 5-CGGCTCGTAGCTGAAGGAACTGTGATG GTGGTGATGGTGGTGGTGATGATGCATGGTGGCGGCCGGTGGCTCC-3). The sequence of the plasmid obtained pEGFP-HisTag-hNfL was confirmed by Sanger Sequencing. Then the target gene *HisTag-hNfL* was amplified from pEGFP-HisTag-hNfL plasmid by infusion primers (HishNfLinsert-F1: 5-GGAGACCCAAGCTTGGTACCA TGCATCATCACCACCACC-3; HishNfLinsert-R1: 5-CATGCTCGAGCGGCCGCTCAATCTTTC TTCTTAGCTGCTTG-3) and inserted into linearized pCDNA3.1 vector by infusion ligation method based on KpnI/NotI cleavage sites. The sequence of the resulting plasmid pCDNA3-HisTag-hNfL was verified by Sanger Sequencing. This plasmid will be used for hNfL protein expression in HEK293T cells.

Protein (*top*) and DNA sequence (*bottom*) of human neurofilament light chain (hNfL) (NCBI Reference Sequence: NP_006149.2) are shown below.

**Protein sequence of human NFL (NCBI Reference Sequence: NP_006149.2):**

MSSFSYEPYYSTSYKRRYVETPRVHISSVRSGYSTARSAYSSYSAPVSSSLSVRRSYSSSSGSLMPSLENLDLSQ
VAAISNDLKSIRTQEKAQLQDLNDRFASFIERVHELEQQNKVLEAELLVLRQKHSEPSRFRALYEQEIRDLRLAA
EDATNEKQALQGEREGLEETLRNLQARYEEEVLSREDAEGRLMEARKGADEAALARAELEKRIDSLMDEISFLKK
VHEEEIAELQAQIQYAQISVEMDVTKPDLSAALKDIRAQYEKLAAKNMQNAEEWFKSRFTVLTESAAKNTDAVRA
AKDEVSESRRLLKAKTLEIEA**C**RGMNEALEKQLQELEDKQNADISAMQDTINKLENELRTTKSEMARYLKEYQDL
LNVKMALDIEIAAYRKLLEGEETRLSFTSVGSITSGYSQSSQVFGRSAYGGLQTSSYLMSTRSFPSYYTSHVQEE
QIEVEETIEAAKAEEAKDEPPSEGEAEEEEKDKEEAEEEEAAEEEEAAKEESEEAKEEEEGGEGEEGEETKEAEE
EEKKVEGAGEEQAAKKKD

**DNA sequence of human NFL in plasmid phNFL (Addgene 132589):**

ATGAGTTCCTTCAGCTACGAGCCGTACTACTCGACCTCCTACAAGCGGCGCTACGTGGAGACGCCCCGGGTGCAC
ATCTCCAGCGTGCGCAGCGGCTACAGCACCGCACGCTCAGCTTACTCCAGCTACTCGGCGCCGGTGTCTTCCTCG
CTGTCCGTGCGCCGCAGCTACTCCTCCAGCTCTGGATCGTTGATGCCCAGTCTGGAGAACCTCGACCTGAGCCAG
GTAGCCGCCATCAGCAACGACCTCAAGTCCATCCGCACGCAGGAGAAGGCGCAGCTCCAGGACCTCAATGACCGC
TTCGCCAGCTTCATCGAGCGCGTGCACGAGCTGGAGCAGCAGAACAAGGTCCTGGAAGCCGAGCTGCTGGTGCTG
CGCCAGAAGCACTCCGAGCCATCCCGCTTCCGGGCGCTGTACGAGCAGGAGATCCGCGACCTGCGCCTGGCGGCG
GAAGATGCCACCAACGAGAAGCAGGCGCTCCAGGGCGAGCGCGAAGGGCTGGAGGAGACCCTGCGCAACCTGCAG
GCGCGCTATGAAGAGGAGGTGCTGAGCCGCGAGGACGCCGAGGGCCGGCTGATGGAAGCGCGCAAAGGCGCCGAC
GAGGCGGCGCTCGCTCGCGCCGAGCTCGAGAAGCGCATCGACAGCTTGATGGACGAAATCTCTTTTCTGAAGAAA
GTGCACGAAGAGGAGATCGCCGAACTGCAGGCGCAGATCCAGTACGCGCAGATCTCCGTGGAGATGGACGTGACC
AAGCCCGACCTTTCCGCCGCGCTCAAGGACATCCGCGCGCAGTACGAGAAGCTGGCCGCCAAGAACATGCAGAAC
GCTGAGGAATGGTTCAAGAGCCGCTTCACCGTGCTGACCGAGAGCGCCGCCAAGAACACCGACGCCGTGCGCGCC
GCCAAGGACGAGGTGTCCGAGAGCCGTCGTCTGCTCAAGGCCAAGACCCTGGAAATCGAAGCATGCCGGGGCATG
AATGAAGCGCTGGAGAAGCAGCTGCAGGAGCTGGAGGACAAGCAGAACGCCGACATCAGCGCTATGCAGGACACG
ATCAACAAATTAGAAAATGAATTGAGGACCACAAAGAGTGAAATGGCACGATACCTAAAAGAATACCAAGACCTC
CTCAACGTGAAGATGGCTTTGGATATTGAGATTGCAGCTTACAGGAAACTCTTGGAAGGCGAGGAGACCCGACTC
AGTTTCACCAGCGTGGGAAGCATAACCAGTGGCTACTCCCAGAGCTCCCAGGTCTTTGGCCGATCTGCCTACGGC
GGTTTACAGACCAGCTCCTATCTGATGTCCACCCGCTCCTTCCCGTCCTACTACACCAGCCATGTCCAAGAGGAG
CAGATCGAAGTGGAGGAAACCATTGAGGCTGCCAAGGCTGAGGAAGCCAAGGATGAGCCCCCCTCTGAAGGAGAA
GCCGAGGAGGAGGAGAAGGACAAGGAAGAGGCCGAGGAAGAGGAGGCAGCTGAAGAGGAAGAAGCTGCCAAGGAA
GAGTCTGAAGAAGCAAAAGAAGAAGAAGAAGGAGGTGAAGGTGAAGAAGGAGAGGAAACCAAAGAAGCTGAAGAG
GAGGAGAAGAAAGTTGAAGGTGCTGGGGAGGAACAAGCAGCTAAGAAGAAAGATTGA

Protein and DNA sequences of hNfL following mutagenesis experiments to insert HisTag onto the N-terminus are shown below.

**Protein sequence of recombinant human NFL:**

MHHHHHHHHHHSSFSYEPYYSTSYKRRYVETPRVHISSVRSGYSTARSAYSSYSAPVSSSLSVRRSYSSSSG
SLMPSLENLDLSQVAAISNDLKSIRTQEKAQLQDLNDRFASFIERVHELEQQNKVLEAELLVLRQKHSEPSR
FRALYEQEIRDLRLAAEDATNEKQALQGEREGLEETLRNLQARYEEEVLSREDAEGRLMEARKGADEAALAR
AELEKRIDSLMDEISFLKKVHEEEIAELQAQIQYAQISVEMDVTKPDLSAALKDIRAQYEKLAAKNMQNAEE
WFKSRFTVLTESAAKNTDAVRAAKDEVSESRRLLKAKTLEIEACRGMNEALEKQLQELEDKQNADISAMQDT
INKLENELRTTKSEMARYLKEYQDLLNVKMALDIEIAAYRKLLEGEETRLSFTSVGSITSGYSQSSQVFGRS
AYGGLQTSSYLMSTRSFPSYYTSHVQEEQIEVEETIEAAKAEEAKDEPPSEGEAEEEEKDKEEAEEEEAAEE
EEAAKEESEEAKEEEEGGEGEEGEETKEAEEEEKKVEGAGEEQAAKKKD

**DNA sequence of recombinant human NFL in pCDNA3 vector:**

ATGCATCATCACCACCACCATCACCACCATCACAGTTCCTTCAGCTACGAGCCGTACTACTCGACCTCCTAC
AAGCGGCGCTACGTGGAGACGCCCCGGGTGCACATCTCCAGCGTGCGCAGCGGCTACAGCACCGCACGCTCA
GCTTACTCCAGCTACTCGGCGCCGGTGTCTTCCTCGCTGTCCGTGCGCCGCAGCTACTCCTCCAGCTCTGGA
TCGTTGATGCCCAGTCTGGAGAACCTCGACCTGAGCCAGGTAGCCGCCATCAGCAACGACCTCAAGTCCATC
CGCACGCAGGAGAAGGCGCAGCTCCAGGACCTCAATGACCGCTTCGCCAGCTTCATCGAGCGCGTGCACGAG
CTGGAGCAGCAGAACAAGGTCCTGGAAGCCGAGCTGCTGGTGCTGCGCCAGAAGCACTCCGAGCCATCCCGC
TTCCGGGCGCTGTACGAGCAGGAGATCCGCGACCTGCGCCTGGCGGCGGAAGATGCCACCAACGAGAAGCAG
GCGCTCCAGGGCGAGCGCGAAGGGCTGGAGGAGACCCTGCGCAACCTGCAGGCGCGCTATGAAGAGGAGGTG
CTGAGCCGCGAGGACGCCGAGGGCCGGCTGATGGAAGCGCGCAAAGGCGCCGACGAGGCGGCGCTCGCTCGC
GCCGAGCTCGAGAAGCGCATCGACAGCTTGATGGACGAAATCTCTTTTCTGAAGAAAGTGCACGAAGAGGAG
ATCGCCGAACTGCAGGCGCAGATCCAGTACGCGCAGATCTCCGTGGAGATGGACGTGACCAAGCCCGACCTT
TCCGCCGCGCTCAAGGACATCCGCGCGCAGTACGAGAAGCTGGCCGCCAAGAACATGCAGAACGCTGAGGAA
TGGTTCAAGAGCCGCTTCACCGTGCTGACCGAGAGCGCCGCCAAGAACACCGACGCCGTGCGCGCCGCCAAG
GACGAGGTGTCCGAGAGCCGTCGTCTGCTCAAGGCCAAGACCCTGGAAATCGAAGCATGCCGGGGCATGAAT
GAAGCGCTGGAGAAGCAGCTGCAGGAGCTGGAGGACAAGCAGAACGCCGACATCAGCGCTATGCAGGACACG
ATCAACAAATTAGAAAATGAATTGAGGACCACAAAGAGTGAAATGGCACGATACCTAAAAGAATACCAAGAC
CTCCTCAACGTGAAGATGGCTTTGGATATTGAGATTGCAGCTTACAGGAAACTCTTGGAAGGCGAGGAGACC
CGACTCAGTTTCACCAGCGTGGGAAGCATAACCAGTGGCTACTCCCAGAGCTCCCAGGTCTTTGGCCGATCT
GCCTACGGCGGTTTACAGACCAGCTCCTATCTGATGTCCACCCGCTCCTTCCCGTCCTACTACACCAGCCAT
GTCCAAGAGGAGCAGATCGAAGTGGAGGAAACCATTGAGGCTGCCAAGGCTGAGGAAGCCAAGGATGAGCCC
CCCTCTGAAGGAGAAGCCGAGGAGGAGGAGAAGGACAAGGAAGAGGCCGAGGAAGAGGAGGCAGCTGAAGAG
GAAGAAGCTGCCAAGGAAGAGTCTGAAGAAGCAAAAGAAGAAGAAGAAGGAGGTGAAGGTGAAGAAGGAGAG
GAAACCAAAGAAGCTGAAGAGGAGGAGAAGAAAGTTGAAGGTGCTGGGGAGGAACAAGCAGCTAAGAAGAAA
GATTGA

The PCR reaction to insert the HisTag was run under conditions as shown below and the sequence was confirmed by Sanger Sequencing using the universal primers CMVF-pCDNA3 and EGFP-CR.

.

| PCR components | PCR reaction | | |
|---|---|---|---|
| 5 μL 10× reaction buffer | 95 °C | 30 s | 1× |
| 1 μL dNTPmix solution | 95 °C | 30 s | 18× |
| 3 μL Quick solution | 55 °C | 60 s | |
| 2 μL phNfL-1 (5 ng/ul) | 68 °C | 8 mins | |
| 5 μL HishNfL-F1 (25 ng/ul) | 68 °C | 10 mins | 1× |
| 5 μL HishNfL-R1 (25 ng/ul) | 10 °C | ∞ | 1× |
| 28 μL $ddH_2O$ | | | |
| 1 μL Ultra pfu enzyme | | | |

Insertion into pCDNA3 vector:

The hNfL-HisTag gene was truncated and inserted into pCDNA3 vector by ligation based on KpnI/NotI cleavage sites. Standard PCR conditions shown below were employed to amplify the gene, and the PCR product was analysed on a 1% agarose DNA gel. The linearized pCDNA3 vector was obtained following digestion using KpnI-HF and NotI-HF of the plasmid Syn4-pcDNA3. Purification of hNfL-HisTag and the linearized pCDNA3 vector were performed using agarose gel electrophoresis, and their concentrations were measured by a NanoSpectrophotometer.

| PCR components | PCR reaction | | |
|---|---|---|---|
| 1 μL HisinsertNfL (2 ng/μL) | 98 °C | 40 s | 1× |
| 1.5 μL HishNFLinsert-F1 (10 μM) | 98 °C | 15 s | 35× |
| 1.5 μL HishNFLinsert-R1 (10 μM) | 55 °C | 30 s | |
| 21 μL $ddH_2O$ | 72 °C | 2 mins | |
| 25 μL CloneAmp HiFi PCR Premix | 72 °C | 5 mins | 1× |

Next, a typical ligation reaction was carried out under conditions which included 100 ng of pCDNA3 vector, 63.3 ng of the NfL-HisTag insert, 1.0 µL of infusion ligation enzyme and 1.49 µL of $ddH_2O$.The ligation product was subsequently transferred into XL10-gold competent cells and plated on LB agar plates (with 100 µg/mL ampicillin) for overnight culture at 37 $^oC$. The single colonies were then selected for overnight small-scale liquid culture (10 ml LB broth with 100 µg/mL ampicillin). The plasmid (pCDNA3-HishNfL) for HisTag hNfL was extracted, and its sequence was confirmed by Sanger Sequencing using the universal primers CMVF-pCDNA3 and bGHR.

Protein expression and purification:

HEK293T cells were grown to 80% confluency in 10 x 150 mm dishes in DMEM supplemented with 10% FBS and 4 mM L-glutamine. A transfection solution was pre-made (20 mL OMEM, 500 µg of plasmid at 1042 ng $\mu L^{-1}$, 1.5 mL of PEI max at 1 mg $mL^{-1}$) by dissolving plasmid in OMEM and then adding PEI Max, before mixing and leaving to incubate at RT for 30 min. Hence, final concentrations in OMEM were 22.73 µg $mL^{-1}$ of plasmid and 68 µg $\mu L^{-1}$ of PEI Max (i.e. 3x plasmid). At this point the media was removed by aspiration and swapped to OMEM (10 mL) and then transfection solution (2.175 mL) was added. The cells were the incubated at 37 °C, 5% $CO_2$ for 6 h before the media was removed and replaced with 25 mL DMEM (10% FBS, 4 mM L-glutamine) and left to incubate for 4 days. The media was then removed and the cells were harvested by scrapping and washed with PBS and spun down and the cell pellet was stored -80 °C before purification.

The cell pellet from 10 x 150 mm dishes was suspended in 20 mL of lysis buffer (6 M Urea, 20 mM Tris·HCl, 500 mM NaCl, 0.5 mM TCEP, pH 7.8) and then lysed on ice using a Dounce Homogeniser (type A) with 40 strokes. This was then centrifuged (13,000 rpm, 4 °C, 50 min) and then the supernatant was removed and added to pre-washed $Ni^{2+}$ NTA resin (3 mL, 50% by volume in EtOH, i.e. 1.5 mL total resin) and incubated with inversion at 4 °C for 2 h. The resin was then transferred to an empty column and the protein was then eluted using an imidazole gradient (100 to 500 mM, in the same lysis buffer), and analysed by SDS-PAGE. Clean fractions were combined, concentrated with an Amicon

10 kDa MWCO centrifugal filter and then buffer exchanged into 3 M Gdn·HCl, 50 mM Tris·HCl, pH 8.0 using a pre-equilibrated PD MiniTrap G-25 column and concentrated to for long-term storage in -80 °C. Below are the volumes and concentrations of imidazole used for the purification of WT hNfL and SDS-PAGE analysis of each fraction.

Stepwise elution gradient:

| Fraction | Imidazole (mM) | Volume per column (mL) |
|---|---|---|
| FT | | |
| W1 | 15 | 10 |
| W2 | 30 | 1.9 |
| W3 | 30 | 1.9 |
| W4 | 50 | 1.9 |
| W5 | 50 | 1.9 |
| E1 | 100 | 1.5 |
| E2 | 150 | 1.5 |
| E3 | 200 | 1.5 |
| E4 | 300 | 1.5 |
| E5 | 500 | 1.5 |
| E6 | 500 | 1.9 |
| E7 | 500 | 1.9 |
| E8 | 500 | 1.9 |

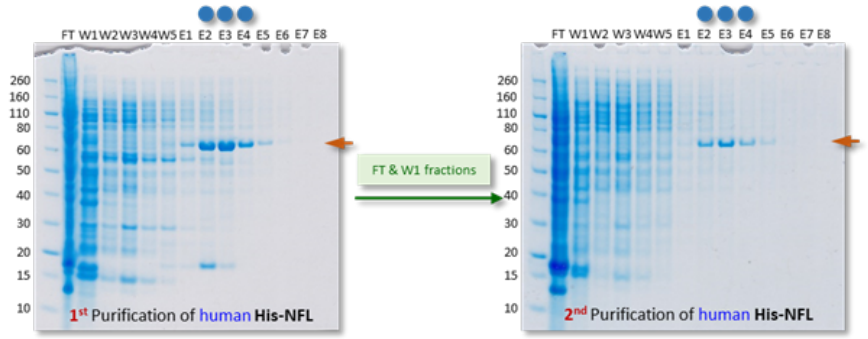

Protein sequence:

Human Neurofilament Light Chain (hNfL)

Mutations: N-terminal 10x Histidine tag

```
        10         20         30         40         50         60
MHHHHHHHHH HSSFSYEPYY STSYKRRYVE TPRVHISSVR SGYSTARSAY SSYSAPVSSS

        70         80         90        100        110        120
LSVRRSYSSS SGSLMPSLEN LDLSQVAAIS NDLKSIRTQE KAQLQDLNDR FASFIERVHE

       130        140        150        160        170        180
LEQQNKVLEA ELLVLRQKHS EPSRFRALYE QEIRDLRLAA EDATNEKQAL QGEREGLEET

       190        200        210        220        230        240
LRNLQARYEE EVLSREDAEG RLMEARKGAD EAALARAELE KRIDSLMDEI SFLKKVHEEE

       250        260        270        280        290        300
IAELQAQIQY AQISVEMDVT KPDLSAALKD IRAQYEKLAA KNMQNAEEWF KSRFTVLTES

       310        320        330        340        350        360
AAKNTDAVRA AKDEVSESRR LLKAKTLEIE ACRGMNEALE KQLQELEDKQ NADISAMQDT

       370        380        390        400        410        420
INKLENELRT TKSEMARYLK EYQDLLNVKM ALDIEIAAYR KLLEGEETRL SFTSVGSITS

       430        440        450        460        470        480
GYSQSSQVFG RSAYGGLQTS SYLMSTRSFP SYYTSHVQEE QIEVEETIEA AKAEEAKDEP

       490        500        510        520        530        540
PSEGEAEEEE KDKEEAEEEE AAEEEEAAKE ESEEAKEEEE GGEGEEGEET KEAEEEEKKV

       550
EGAGEEQAAK KKD
```

Calculated molecular weight: 62888 g mol$^{-1}$ (with Met)

Extinction coefficient ε: 36790 M$^{-1}$ cm$^{-1}$

ESI analysis of WT hNfL:

Observed mass: 62930 g mol$^{-1}$

Calculated mass for 1x carbamylation: 62931 g mol$^{-1}$

Observed mass: 63009 g mol$^{-1}$

Calculated mass for 1x carbamylation and 1x phosphorylation: 63011 g mol$^{-1}$

Observed mass: 63089 g mol$^{-1}$

Calculated mass for 1x carbamylation and 2x phosphorylation: 63091 g $mol^{-1}$

Observed mass: 63169 g $mol^{-1}$

Calculated mass for 1x carbamylation and 3x phosphorylation: 63171 g $mol^{-1}$

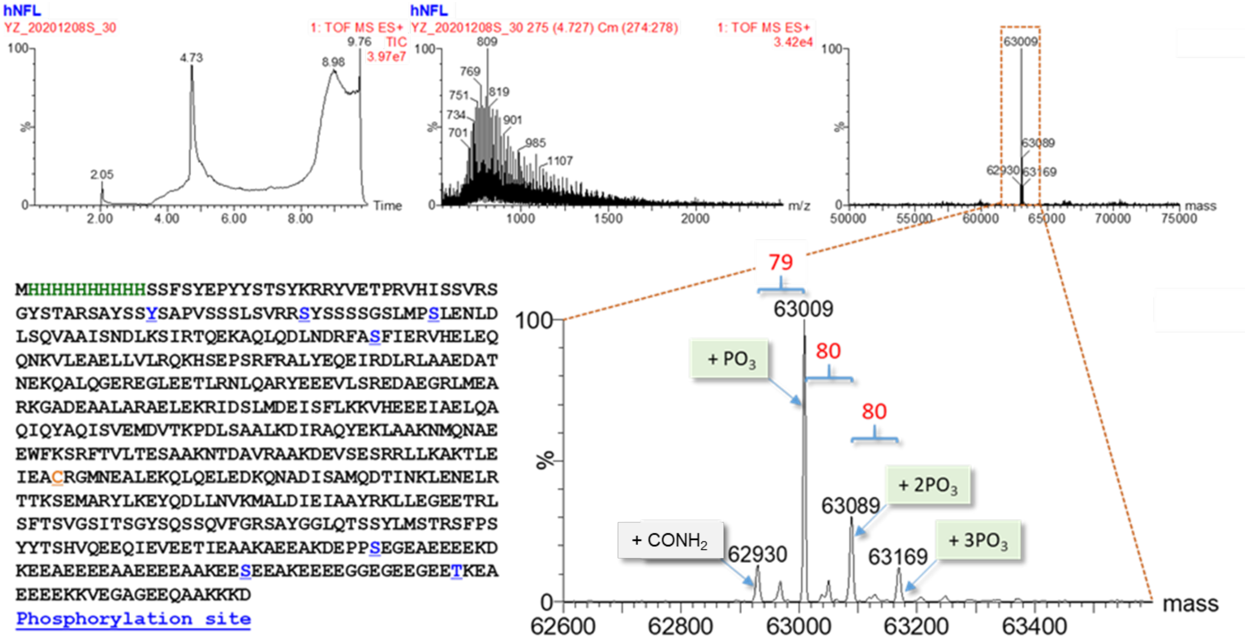

**Commercially available antibodies for NfL detection**

Antibodies were used largely as per the manufacturer's recommendations. Primary antibodies: UD1 antibody (Uman Diag 27016), UD3 antibody (Uman Diag 27018), DA2 antibody (Novus NB300-132) and *anti*-NFL antibody (GeneTex GTX134085). Secondary antibodies: Goat Anti-mouse IgG-Alkaline Phosphatase antibody produced in goat (Promega S3721), Goat Anti-Rabbit IgG-Alkaline Phosphatase antibody produced in goat (Sigma A3687).

General Procedure: SDS-PAGE gel of the samples was run on a Novex Bis-Tris NuPAGE before transferring to a nitrocellulose membrane with an iBlot gel transfer device. The membrane was then blocked (TBS with 5% BSA) for 1 h at rt, washed with TBS for 5 min then incubated with the primary antibody (dilution 1:2000 for all primary antibodies mentioned above) in TBS with 5% BSA overnight at 4 °C. The membrane was washed 3 times with TBS containing 0.1% Tween-20 prior to incubation with an Alkaline Phosphatase-coupled secondary antibody (dilution 1:5000) in TBS containing 0.1% Tween-20. The membrane was washed 3 times with TBS containing 0.1% Tween-20 and then once with TBS before detection by a colorimetric method (Sigma BCIP/NBT Liquid Substrate System, catalogue number B1911).

**LC-MS analysis of different NfL samples**

LC-MS analysis were performed on NfL protein samples produced in-house (mouse and human NfL) and on those purchased commercially (from Encor Biotechnology Inc.). As shown in the spectra below, NfL samples produced in-house (top and middle) generated reasonable ion series that deconvoluted to give the expected protein mass. In contrast, LC-MS analysis of the commercially supplied NfL sample (bottom) was problematic, despite using a similar protein concentration across all samples.

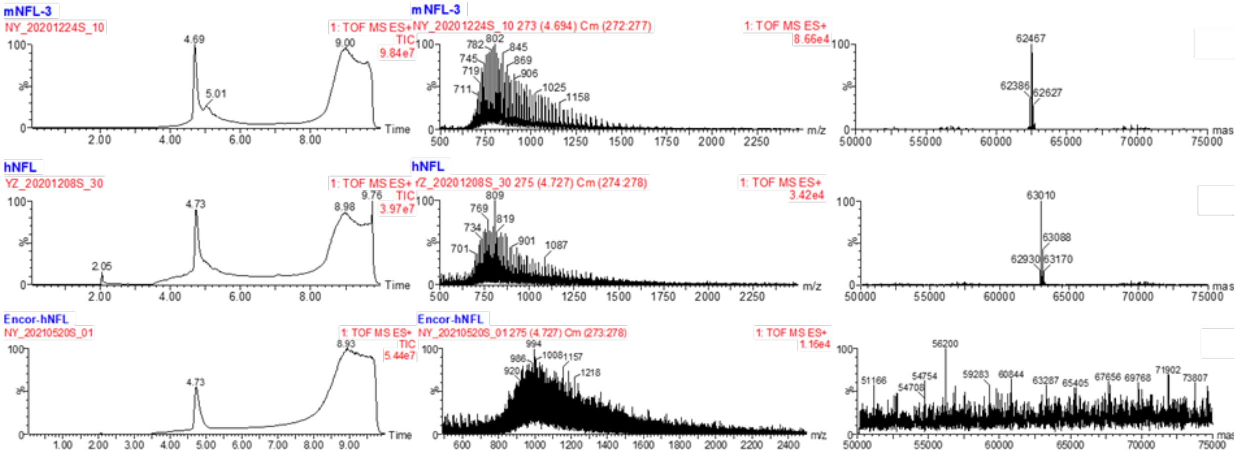

#### 2.5 Dha Formation

**Histones H3**

Lyophilised *X.l.* and human histones (7 mg) was dissolved in denaturing phosphate buffer (100 mM $NaP_i$, pH 8, 3 M Gdn·HCl, 500 µL). DTT was added (30 mg) and the reaction mixture was shaken for 30 min at RT (500 rpm) to reduce disulfide bonds, before desalting into 1 mL of the same buffer (PD Minitrap G25, GE Healthcare). The resulting protein concentration was determined using a Nanodrop and immediately followed by the addition of DBHDA (60 equiv from a freshly prepared 0.5 M DMSO stock) and subsequent shaking (500 rpm) at 25 °C for 45 min, then 37 °C for at least 2 h. The protein was desalted as before to remove the excess DBHDA and exchange into the desired buffer. Protein yield and concentration were determined by its UV absorbance using the Nanodrop and conversion was determined by LC-MS analysis.

Representative LC-MS analysis (ion series and magnification of the deconvoluted spectrum):

*Xenopus laevis* (*X.l.*) Histone H3.NTEV-Dha2

Calculated mass: 16004 Da

Observed mass: 16004 Da

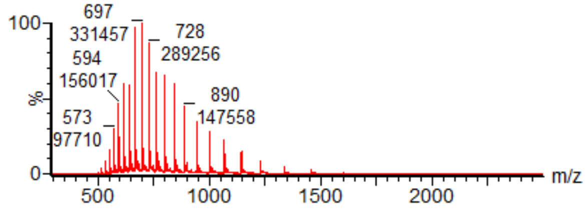


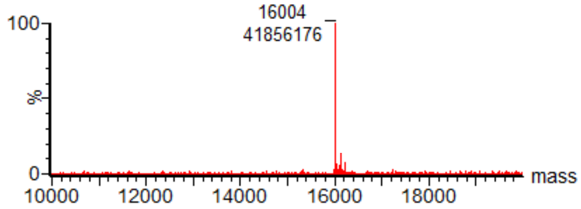

*Xenopus laevis* (*X.l.*) Histone H3-Dha9

Calculated mass: 15180 Da

Observed mass: 15180 Da

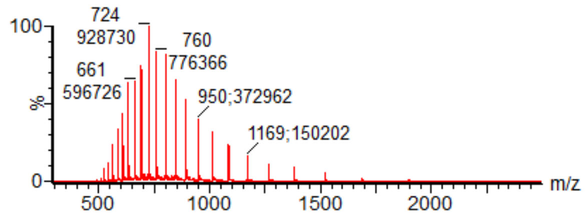


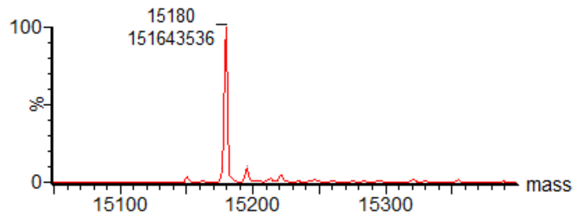


Human Histone eH3.1-Dha4

Calculated mass: 17686 Da

Observed mass: 17686 Da

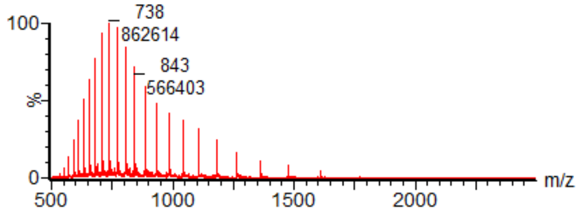


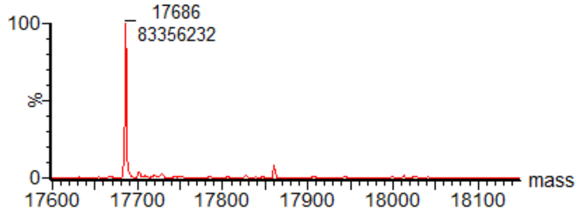

**Mouse Neurofilament Light Chain**

To a solution of WT mNfL in denaturing Tris buffer (50 mM Tris·HCl, pH 8, 4 M Gdn·HCl, 500 μL), TCEP was added to make a 0.5 mM TCEP-buffered solution in a 100 μL reaction volume and a final protein concentration of approximately 25 μM. Methyl 2,5-dibromopentanoate, often referred to as MDBP, was added (3000 equiv. from a freshly prepared 1.5 M DMSO stock) and the reaction mixture was shaken for 3 h at 37 °C (700 rpm). The protein was desalted to remove the excess MDBP and exchange into the desired buffer. Subsequent steps that involve the monofluoromethylation of mNfL-Dha323 require buffer exchange of the protein into 500 mM $NH_4OAc$ containing high concentration of denaturant (typically, 3 to 7 M Gdn·HCl, pH 6). Protein yield and concentration were determined by its UV absorbance using the Nanodrop and conversion was determined by LC-MS analysis.

ESI analysis of mNfL-Dha323:

Observed mass: 62353 and 62391 g $mol^{-1}$

Calculated mass for 1x carbamylation: 62353 g $mol^{-1}$

Calculated mass for 2x carbamylation: 62396 g $mol^{-1}$

Observed mass: 62434 and 62473 g $mol^{-1}$

Calculated mass for 1x carbamylation and 1x phosphorylation: 62433 g $mol^{-1}$

Calculated mass for 2x carbamylation and 1x phosphorylation: 62473 g $mol^{-1}$

Observed mass: 62513 and 62564 g $mol^{-1}$

Calculated mass for 1x carbamylation and 2x phosphorylation: 62513 g $mol^{-1}$

Calculated mass for 2x carbamylation and 2x phosphorylation: 62556 g $mol^{-1}$

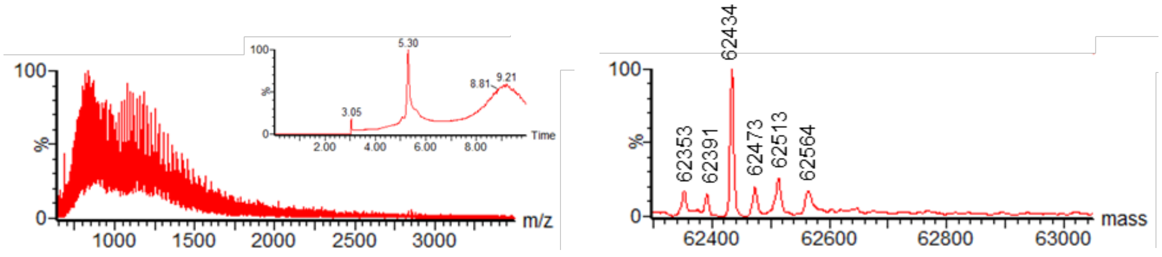

#### 2.6 'Cold' Protein Chemistry

**General protein reaction protocol**

All protein solutions and solvents were degassed for at least 8 h in a glovebox (<10 ppm $O_2$). All solid reagents were weighed and then transferred into the glovebox where stock solutions were prepared prior to protein reactions. Protein reactions were carried out in clear glass vials (ThermoFisher Scientific 9 mm clear glass screw thread vial with 300 μL fused insert, for reactions ≤50 μL, ThermoFisher Scientific 2.0 mL 9 mm clear glass screw thread vial, flat bottom, for reactions ≤100 μL) with gas-tight lids. Standard reactions were prepared by diluting the Dha-containing protein solution in the buffer of choice to reach the desired protein concentration, followed by sequential addition of photocatalyst, sulfone reagents and iron additives in the glovebox. The reactions were then mixed thoroughly with a pipette, capped and moved out of the glovebox for irradiation. A variable light intensity (10 – 50 W) photobox was used with blue LEDs arranged in series to allow up to 7 reactions at a time. Cooling fan is attached to the photobox for temperature control especially for reactions requiring >20 min. After irradiation, an aliquot of the crude reaction mixture was obtained and diluted into 0.1% formic acid (final protein concentration approx. 0.02 mg $mL^{-1}$) for LC-MS analysis. For epitope tagged human histone eH3.1 and His-tagged NfL, crude reaction mixtures were diluted with 15 mM and 50 mM β-mercaptoethanol in 0.1% formic acid respectively. For long-term storage of the modified protein product, the crude mixture should be purified using PD MiniTrap G-25 (GE Healthcare) desalting columns and protein UV absorbance monitored before storing in the freezer. Unless stated otherwise, conversions reported are to the desired single addition fluoroalkylated product.

#### Formation of *X.l.* Histone H3.NTEV-*b*Fha2 with btSOOF

Under non-denaturing conditions:

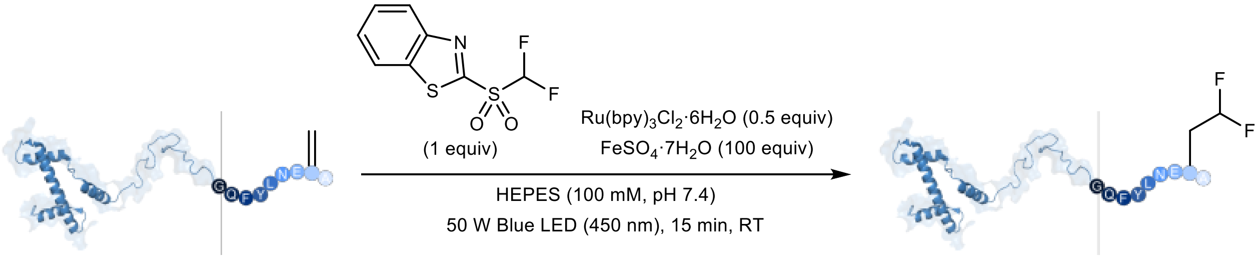


*X.l.* Histone H3.NTEV-Dha2 **90% conversion** *X.l.* Histone H3.NTEV-*b*Fha2

In the glovebox, degassed HEPES buffer (100 mM, pH 7.4) was added to a glass HPLC vial containing *X.l.* Histone H3.NTEV-Dha2 (100 µg) such that the final concentration is 1 mg mL$^{-1}$ (62 µM) i.e. total reaction volume is 100 µL. $Ru(bpy)_3Cl_2·6H_2O$ (1 µL of a 3.1 mM stock prepared fresh in degassed $H_2O$, 0.5 equiv), btSOOF radical precursor (1 µL of a 6.2 mM stock prepared fresh in degassed DMSO, 1 equiv) and $FeSO_4·7H_2O$ (2 µL of a 310 mM stock prepared fresh in degassed $H_2O$, 100 equiv) were added sequentially and the vial was sealed with a lid before transferring out of the glovebox and irradiating with blue LED light (50 W) for 15 min. A sample of the crude mixture was then analysed by LC-MS to determine reaction conversion. Keeping all other conditions constant, the same conversion was observed with 2 equiv of photocatalyst.

ESI-MS analysis of *X.l.* Histone H3.NTEV:

Calculated H3 *b*Fha mass: 16056 Da

Observed mass: 16055 Da (+16 Da is methionine oxidation)

Calculated H3 Dha mass: 16004 Da

Observed mass: 16003 Da

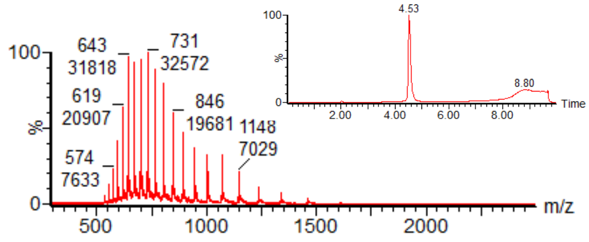


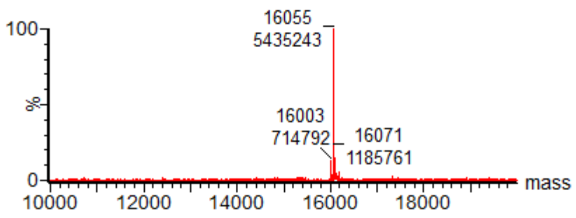

#### Formation of *X.l.* Histone H3-*b*Fha9 with btSOOF

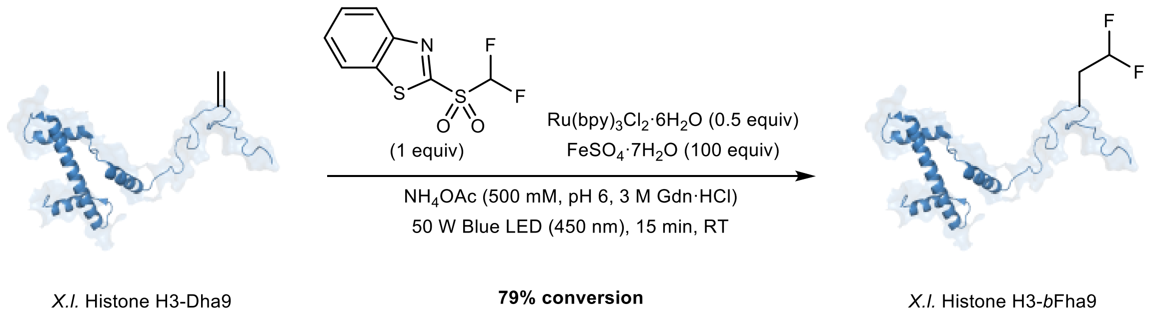


In the glovebox, degassed $NH_4OAc$ buffer (500 mM, pH 6, 3 M Gdn·HCl) was added to a glass HPLC vial containing *X.l.* Histone H3-Dha9 (100 µg) such that the final concentration is 1 mg $mL^{-1}$ (66 µM) i.e. total reaction volume is 100 µL. $Ru(bpy)_3Cl_2·6H_2O$ (1 µL of a 3.3 mM stock prepared fresh in degassed $H_2O$, 0.5 equiv), btSOOF radical precursor (0.825 µL of a 8 mM stock prepared fresh in degassed DMSO, 1 equiv) and $FeSO_4·7H_2O$ (2 µL of a 330 mM stock prepared fresh in degassed $H_2O$, 100 equiv) were added sequentially and the vial was sealed with a lid before transferring out of the glovebox and irradiating with blue LED light (50 W) for 15 min. A sample of the crude mixture was then analysed by LC-MS to determine reaction conversion.

ESI-MS analysis of *X.l.* Histone H3:
Calculated H3 *b*Fha mass: 15232 Da
Observed mass: 15231 Da
Calculated H3 Dha mass: 15180 Da
Observed mass: 15179 Da

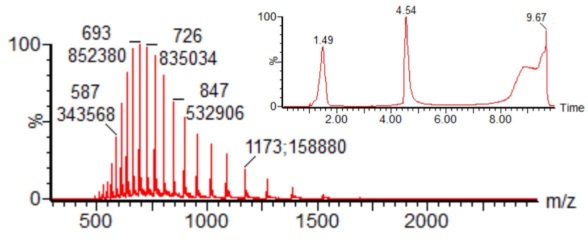


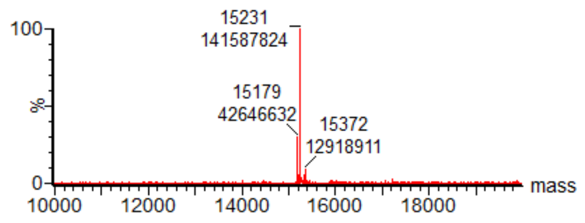

#### Formation of *X.l.* Histone H3.NTEV-Fha2 with mono-btSOOF

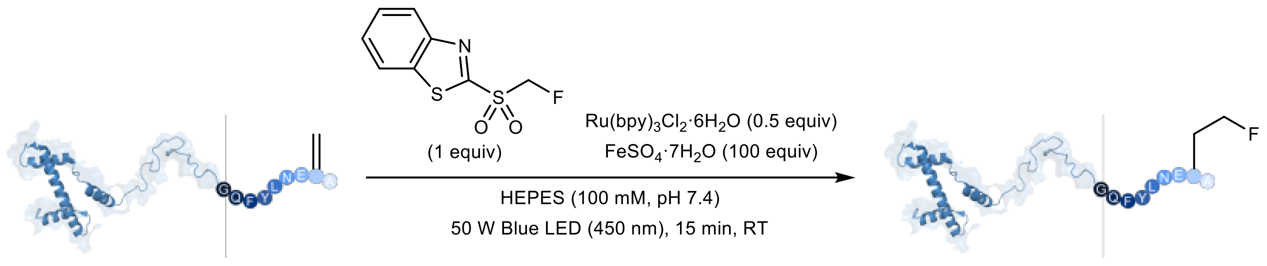


In the glovebox, degassed HEPES buffer (100 mM, pH 7.4) was added to a glass HPLC vial containing *X.l.* Histone H3.NTEV-Dha2 (100 µg) such that the final concentration is 1 mg mL$^{-1}$ (62 µM) i.e. total reaction volume is 100 µL. $Ru(bpy)_3Cl_2·6H_2O$ (1 µL of a 3.1 mM stock prepared fresh in degassed $H_2O$, 0.5 equiv), mono-btSOOF radical precursor (1 µL of a 6.2 mM stock prepared fresh in degassed DMSO, 1 equiv) and $FeSO_4·7H_2O$ (2 µL of a 310 mM stock prepared fresh in degassed $H_2O$, 100 equiv) were added sequentially and the vial was sealed with a lid before transferring out of the glovebox and irradiating with blue LED light (50 W) for 15 min. A sample of the crude mixture was then analysed by LC-MS to determine reaction conversion. Keeping all other conditions constant, similar conversion was observed with 2 equiv of photocatalyst (~90% without any double addition by-product).

ESI-MS analysis of *X.l.* Histone H3.NTEV:
Calculated H3 Fha mass: 16038 Da
Observed mass: 16037 Da (+16 Da is methionine oxidation)
Calculated H3 Dha mass: 16004 Da
Observed mass: 16003 Da

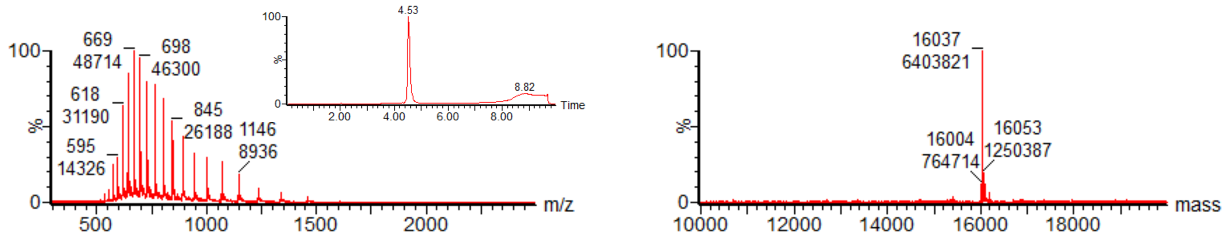

#### Formation of *X.l.* Histone H3-Fha27 with mono-btSOOF

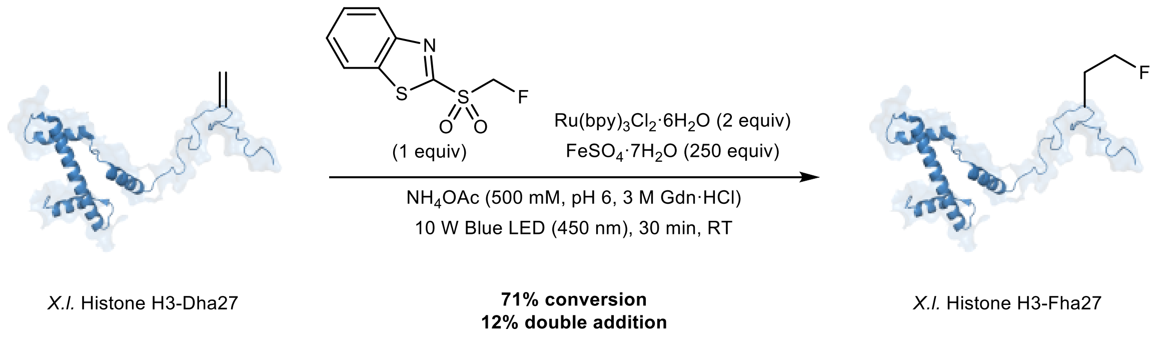


In the glovebox, degassed $NH_4OAc$ buffer (500 mM, pH 6, 3 M Gdn·HCl) was added to a glass HPLC vial containing *X.l.* Histone H3-Dha27 (100 µg) such that the final concentration is 1 mg $mL^{-1}$ (66 µM) i.e. total reaction volume is 100 µL. $Ru(bpy)_3Cl_2 \cdot 6H_2O$ (2 µL of a 6.6 mM stock prepared fresh in degassed $H_2O$, 2 equiv), mono-btSOOF radical precursor (0.825 µL of a 8 mM stock prepared fresh in degassed DMSO, 1 equiv) and $FeSO_4 \cdot 7H_2O$ (5 µL of a 330 mM stock prepared fresh in degassed $H_2O$, 250 equiv) were added sequentially and the vial was sealed with a lid before transferring out of the glovebox and irradiating with blue LED light (10 W) for 30 min. A sample of the crude mixture was then analysed by LC-MS to determine reaction conversion.

ESI-MS analysis of *X.l.* Histone H3:
Calculated H3 Fha mass: 15214 Da
Observed mass: 15213 Da
Calculated H3 $Fha_2$ double addition mass: 15246 Da
Observed mass: 15245 Da
Calculated H3 Dha mass: 15180 Da
Observed mass: 15179 Da

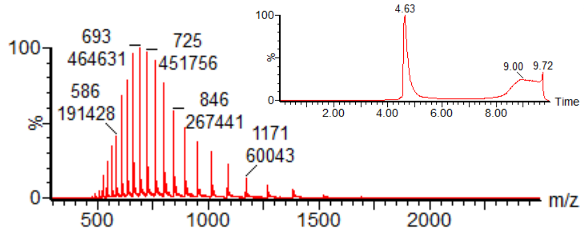


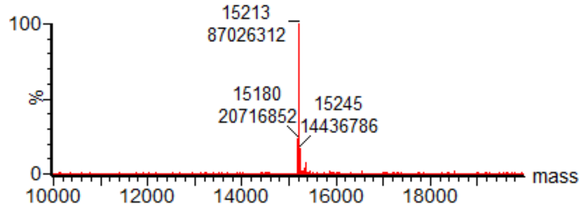

#### Formation of Human Histone eH3.1-Fha4 with mono-btSOOF

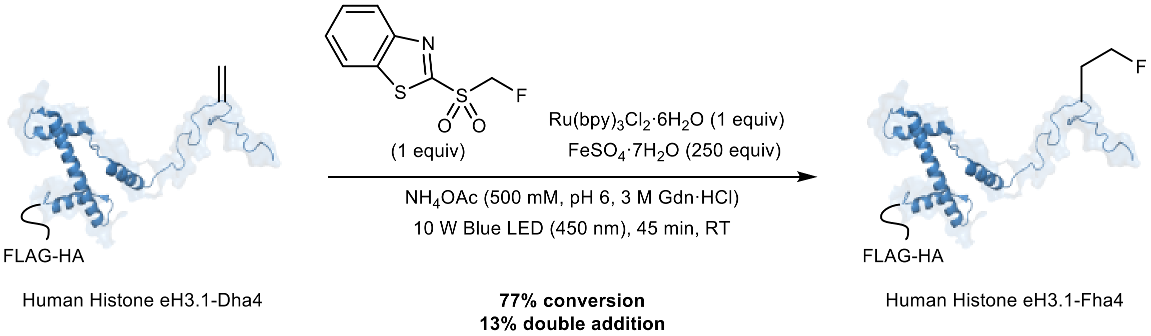


In the glovebox, degassed $NH_4OAc$ buffer (500 mM, pH 6, 3 M Gdn·HCl) was added to a glass HPLC vial containing Human Histone eH3.1-Dha4 (100 µg) such that the final concentration is 1 mg $mL^{-1}$ (57 µM) i.e. total reaction volume is 100 µL. $Ru(bpy)_3Cl_2·6H_2O$ (1 µL of a 5.7 mM stock prepared fresh in degassed $H_2O$, 1 equiv), mono-btSOOF radical precursor (1 µL of a 5.7 mM stock prepared fresh in degassed DMSO, 1 equiv) and $FeSO_4·7H_2O$ (5 µL of a 285 mM stock prepared fresh in degassed $H_2O$, 250 equiv) were added sequentially and the vial was sealed with a lid before transferring out of the glovebox and irradiating with blue LED light (10 W) for 45 min. A sample of the crude mixture was then analysed by LC-MS to determine reaction conversion.

ESI-MS analysis of Human Histone eH3.1:

Calculated H3.1 Fha mass: 17720 Da

Observed mass: 17719 Da (+16 Da is methionine oxidation)

Calculated H3.1 $Fha_2$ double addition mass: 17752 Da

Observed mass: 17751 Da

Calculated H3.1 Dha mass: 17686 Da

Observed mass: 17686 Da

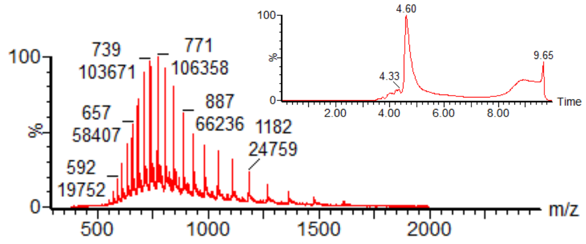


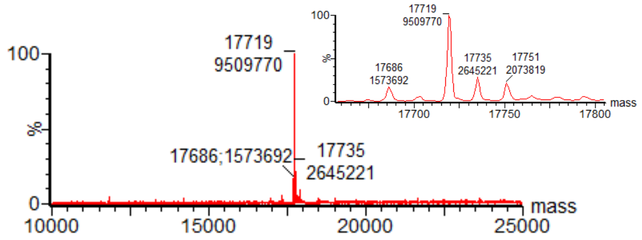

#### Formation of NfL-Fha323 with mono-btSOOF

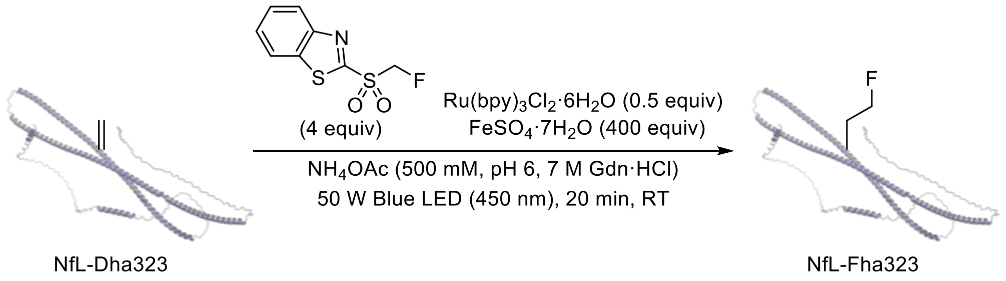


In the glovebox, degassed $NH_4OAc$ buffer (500 mM, pH 6, 7 M Gdn·HCl) was added to a glass HPLC vial containing mNfL-Dha323 (500 µg) such that the final concentration was 5.0 mg $mL^{-1}$ (80 µM) i.e. total reaction volume was 100 µL. $Ru(bpy)_3Cl_2{\cdot}6H_2O$ (0.4 µL of a 10 mM stock prepared fresh in degassed $H_2O$, 0.5 equiv), mono-btSOOF radical precursor (3.2 µL of a 10 mM stock prepared fresh in degassed DMSO, 4 equiv) and $FeSO_4{\cdot}7H_2O$ (12.8 µL of a 250 mM stock prepared fresh in degassed $H_2O$, 400 equiv) were added sequentially and the vial was sealed with a lid before transferring out of the glovebox and irradiating with blue LED light (50 W) for 20 min. At the end of the reaction, the reaction mixture was incubated with 1000 equiv. of EDTA (from a 0.5 M stock) for at least 15 min (400 rpm). A sample of the crude mixture was then analysed by LC-MS to determine reaction conversion.

ESI analysis of mNfL-Fha323:
Observed mass: 62429 g $mol^{-1}$
Calculated mass for 2x carbamylation: 62430 g $mol^{-1}$
Observed mass: 62468 and 62507 g $mol^{-1}$
Calculated mass for 1x carbamylation and 1x phosphorylation: 62467 g $mol^{-1}$
Calculated mass for 2x carbamylation and 1x phosphorylation: 62510 g $mol^{-1}$
Observed mass: 62547 and 62583 g $mol^{-1}$
Calculated mass for 1x carbamylation and 2x phosphorylation: 62547 g $mol^{-1}$
Calculated mass for 2x carbamylation and 2x phosphorylation: 62590 g $mol^{-1}$
Observed mass: 62624 g $mol^{-1}$
Calculated mass for 1x carbamylation and 3x phosphorylation: 62627 g $mol^{-1}$

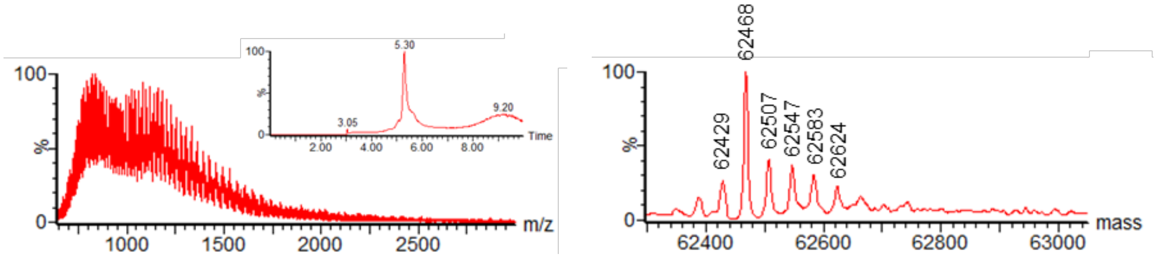

100
%
0
m/z
1000
1500
2000
2500
100
%
0
5.30
3.05
9.20
Time
2.00
4.00
6.00
8.00
62468
62429
62507
62547
62583
62624
100
%
0
mass
62400
62600
62800
63000

##### 2.7 Dephosphorylation of mNfL by λ-protein phosphatase

On-resin dephosphorylation:

70 µL of purified mNfL in Tris·HCl buffer (50 mM, pH 8.125, 3 M Gdn·HCl) was loaded onto 100 µL of freshly prepared Ni-NTA resin. The mixture was incubated for 1 h on ice and the supernatant was removed, The resin was washed with 'cleavage buffer' (50 mM Tris·HCl, pH 7.5, 100 mM NaCl) several times before adding 1.5 µL *λ*-protein phosphatase, 5 µL $MnCl_2$ (10 mM) in 45 µL cleavage buffer to the resin. The resin was incubated for a further 30 min with shaking (600 rpm) at 30 °C. The supernatant was removed and the resin was washed twice with 'washing buffer' (20 mM Tris·HCl, pH 8.125, 3 M Gdn·HCl). The supernatant was removed and the protein was eluted with 250 mM imidazole, pH 7.8, 3 M Gdn·HCl. Dephosphorylation of mNfL was confirmed by LC-MS analysis and Pro-Q Diamond staining.

Pro-Q Diamond staining:

SDS-PAGE was performed on mNfL before and after treatment with λ-protein phosphatase. The gel was fixed with 50% methanol, 20% acetic acid, 30% $H_2O$ at room temperature with gentle agitation (1 h initially followed by an overnight incubation) and then washed three times with water for 10 min each time. The gel was stained with the Pro-Q diamond phosphoprotein gel stain solution for 90 min in the dark and then destained with 20% MeCN, 50 mM NaOAc, pH 4.0 (3 x 30 min). After destaining, the gel was washed with water (2 x 5 min) prior to image analysis using a Bio-Rad Gel Documentation system. Finally, the gel was stained with Coomassie blue to asses total protein concentration.

#### 3. Radiochemistry

##### 3.1 General Methods for Radiochemistry

For radiosynthesis carried out on the Advion Nanotek® radiosynthesizer, [$^{18}$F]fluoride was produced by Allliance Medical (UK), Invicro (UK) or PETIC (UK) via the $^{18}$O(p,n)$^{18}$F reaction and delivered as [$^{18}$F]fluoride in $^{18}$O-enriched-water. Radiosynthesis and azeotropic drying were performed on a NanoTek microfluidic device. HPLC analysis was performed with a Dionex Ultimate 3000 dual channel HPLC system equipped with shared autosampler, parallel UV-detectors and LabLogic NaI/PMT-radiodetectors with Flowram analog output. The radio signal is delayed by 0.1-0.3 min from the UV signal. Radio-TLC was performed on Merck Keiselgel 60 F254 plates with DCM/MeOH (9:1) elution system. Analysis was performed using a plastic scintillator/PMT detector. Radiochemical yield (RCY) for a protein reaction is the radio-fluorinated histone product expressed as a percentage against the remaining $^{18}$F-reagent. The values were determined by radio-HPLC of the crude protein reaction mixture. RCYs for the $^{18}$F-labelling of small molecules are calculated based on the equation shown below taking into account TLC and LC purities. All isolated activity yields are non-decay corrected (n.d.c.). All molar activities are decay corrected until the end of synthesis (EOS) unless stated otherwise.

$$\text{RCY (\%)} = \frac{\text{LC purity (\%)} \times \text{TLC purity (\%)}}{100}$$

[$^{18}$F]/$K_{2.2.2}$ elution on Advion Nanotek® radiosynthesizer: [$^{18}$F]Fluoride was separated from $^{18}$O-enriched-water using anion exchange cartridges (Waters Sep-Pak Accell Plus QMA Carbonate Plus Light Cartridge, pre-activated with $H_2O$ (10.0 mL)) and subsequently released with a solution of $K_{2.2.2}$/$K_2CO_3$ (Kryptofix® (7.5 mg in 600 µL of MeCN) and $K_2CO_3$ (1.5-2 mg in 150-200 µL of $H_2O$)) into the concentrator. The solution was further dried by azeotropic drying using dry MeCN (3 x 500 µL) under a flow of $N_2$ at 105 °C.

For radiosynthesis carried out on the Trasis AllinOne radiosynthesizer, [$^{18}$F]fluoride was produced in-house (Turku PET Centre, Turku, Finland and PETIC, Cardiff, UK) by the $^{18}$O(p,n)$^{18}$F nuclear reaction and supplied as [$^{18}$F]fluoride in $^{18}$O-enriched-water.

**Small molecule HPLC analysis:**

Method A: Synergi Hydro-RP C18 80 Å column was used (4 µm, 4.6 x 150 mm) with a flow rate of 1 mL $min^{-1}$, temperature of 25 °C and UV-Vis wavelength of 220 nm. A solvent system of $H_2O$/MeCN was used with the gradient elution as follows:

| Step | Time / min | % $H_2O$ | % MeCN |
|---|---|---|---|
| 0 | 0 | 95 | 5 |
| 1 | 1 | 95 | 5 |
| 2 | 10 | 5 | 95 |
| 3 | 14 | 5 | 95 |
| 4 | 16 | 95 | 5 |
| 5 | 18 | 95 | 5 |

**Protein HPLC analysis:**

Method B: Phenomenex Jupiter C4 300 Å column (5 µm, 4.6 x 250 mm) column was used with a flow rate of 1 mL $min^{-1}$, temperature of 25 °C and UV-Vis wavelength of 220 nm. A solvent system of $H_2O$ + 0.1% TFA/MeCN + 0.1% TFA was used with the gradient elution as follows:

| Step | Time / min | % $H_2O$ + 0.1% TFA | % MeCN + 0.1% TFA |
|---|---|---|---|
| 0 | 0 | 95 | 5 |
| 1 | 2 | 95 | 5 |
| 2 | 22 | 5 | 95 |
| 3 | 24.5 | 5 | 95 |
| 4 | 25 | 95 | 5 |
| 5 | 30 | 95 | 5 |

Method C: Phenomenex Jupiter C4 300 Å column (5 μm, 4.6 x 250 mm) column was used with a flow rate of 1 mL $min^{-1}$, temperature of 25 °C and UV-Vis wavelength of 220 nm. A solvent system of $H_2O$ + 0.1% TFA/MeCN + 0.1% TFA was used with the gradient elution as follows:

| Step | Time / min | % $H_2O$ + 0.1% TFA | % MeCN + 0.1% TFA |
|---|---|---|---|
| 0 | 0 | 95 | 5 |
| 1 | 2 | 95 | 5 |
| 2 | 32 | 5 | 95 |
| 3 | 34.5 | 5 | 95 |
| 4 | 35 | 95 | 5 |
| 5 | 40 | 95 | 5 |

Method D: Phenomenex Jupiter C4 300 Å column (5 μm, 4.6 x 250 mm) column was used with a flow rate of 1 mL $min^{-1}$, temperature of 25 °C and UV-Vis wavelength of 220 nm. A solvent system of $H_2O$ + 0.1% TFA/MeCN + 0.1% TFA was used with the gradient elution as follows:

| Step | Time / min | % $H_2O$ + 0.1% TFA | % MeCN + 0.1% TFA |
|---|---|---|---|
| 0 | 0 | 95 | 5 |
| 1 | 2 | 95 | 5 |
| 2 | 42 | 5 | 95 |
| 3 | 44.5 | 5 | 95 |
| 4 | 45 | 95 | 5 |
| 5 | 50 | 95 | 5 |

##### 3.2 Manual radiosynthesis of [$^{18}$F]mono-btSOOF and [$^{18}$F]btSOOF using an Advion Nanotek® radiosynthesizer

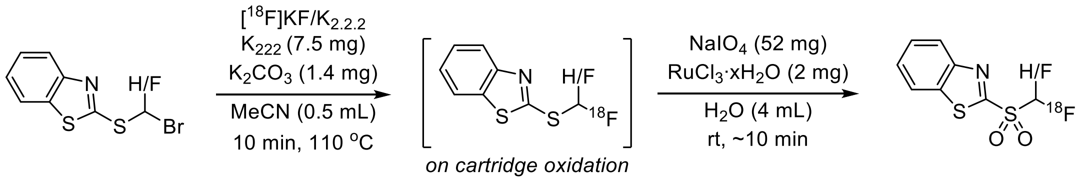


A solution of precursor (0.04 mmol in 0.5 mL MeCN) was added to a V-vial containing a stir bar and dry [$^{18}$F]KF/$K_{2.2.2}$ and the solution was left to stir at 110 °C for 10 min. The crude reaction containing the $^{18}$F-labelled compound was then allowed to cool prior to dilution with 4 mL of $H_2O$. The mixture was then filtered through a C18 plus cartridge (pre-conditioned with EtOH (10 mL) and $H_2O$ (10 mL)). A solution containing $NaIO_4$ (52 mg, 0.24 mmol) and $RuCl_3{\cdot}xH_2O$ (2 mg, 10 mol%) in $H_2O$ (4 mL) was passed through the C18 plus cartridge, passing 1 mL of oxidation solution every 1 min. After complete addition, the oxidation was left for 5 min at room temperature. The crude labelled sulfone reagent was then eluted from the cartridge with 1.2 mL of MeCN prior to semi-prep HPLC purification (flow rate = 4 mL min$^{-1}$, Phenomenex Synergi Hydro-RP 4 µm 80 Å LC column (250 x 10 mm)) with 53-55% MeCN in 25 mM ammonium formate buffer (isocratic)). [$^{18}$F]mono-btSOOF eluted at approximately 10-12 min under isocratic conditions with 53% MeCN in 25 mM ammonium formate buffer whereas [$^{18}$F]btSOOF eluted at approximately ~12-14 min with 55% MeCN in 25 mM ammonium formate buffer. The peak corresponding to the benzothiazole sulfone reagent was collected in a vial containing 20 mL of water. This solution was then passed over a C18 plus cartridge (pre-conditioned with EtOH (10 mL) and $H_2O$ (10 mL)). The reagent was then eluted from the cartridge into the corresponding reaction vial (3.0 mL V-vial) with $Et_2O$ (~1.2 mL total volume). This solution was then concentrated to dryness under a flow of $N_2$ at rt. Protein solution containing the photocatalyst and Fe(II) in degassed, buffered conditions was subsequently added under $N_2$.

| Reagent | Activity Yield | Molar Activity |
|---|---|---|
| [$^{18}$F]mono-btSOOF | 6 ± 2% (n = 5) | 3.6 GBq μmol$^{-1}$ |
| [$^{18}$F]btSOOF | 5 ± 0% (n = 2) | 0.3 GBq μmol$^{-1}$ |

[$^{18}$F]mono-btSOOF radio-HPLC trace: see section 3.3

[$^{18}$F]btSOOF radio-HPLC trace**:**

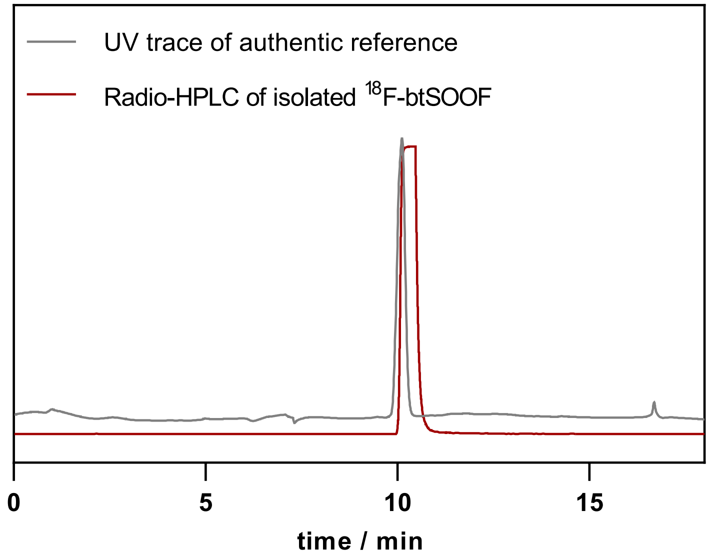


Molar activity measurements

A calibration curve for each authentic reference was recorded by preparing samples of a range of concentrations by serial dilution. These were injected onto an HPLC and the UV response was measured by integrating the peak of interest. The molar activity of $^{18}$F-reagents were determined by subjecting an aliquot of the purified product to HPLC analysis. The UV response corresponding to the reagent was then integrated, to give the amount of non-radioactive $^{19}$F analogue that was detected. Molar activity was then calculated.

Molar activity of [$^{18}$F]mono-btSOOF:

UV peak area = 14.1 mAu

Amount of 19F product = 14.1/11476 = 0.0012 μmol

Activity at 10.38 am (end of synthesis) = 4.36 MBq

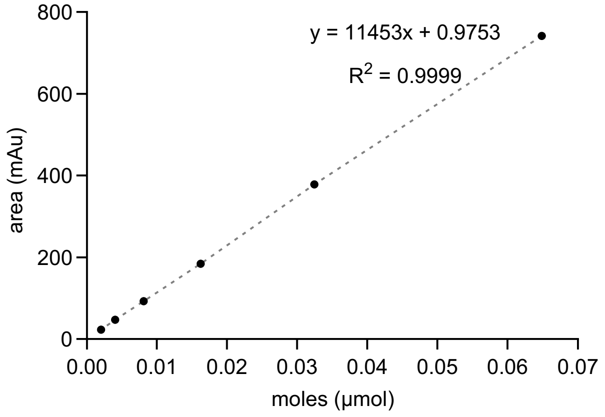


MA = (4.36/0.0012)/1000 = 3.55 GBq/μmol; decay corrected from end of synthesis

Molar activity of [$^{18}$F]btSOOF:

UV peak area = 194.57 mAu

Amount of $^{19}$F product = 194.57/10799 = 0.018 μmol

Activity at 10.21 am (end of synthesis) = 5.42 MBq

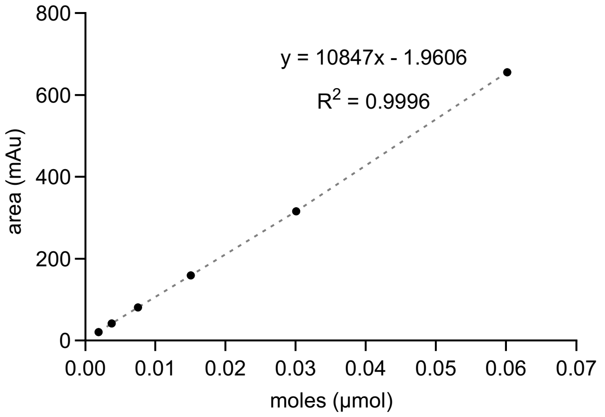


MA = (5.42/0.018)/1000 = 0.30 GBq/μmol; decay corrected from end of synthesis

#### 3.3 Automated radiosynthesis of [$^{18}$F]mono-btSOOF using a Trasis AllinOne radiosynthesizer

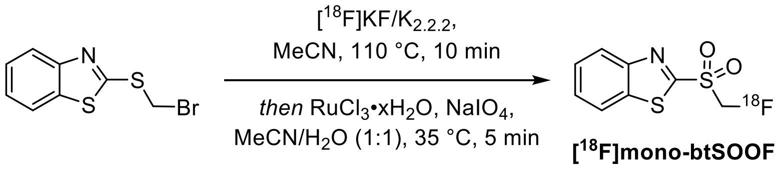


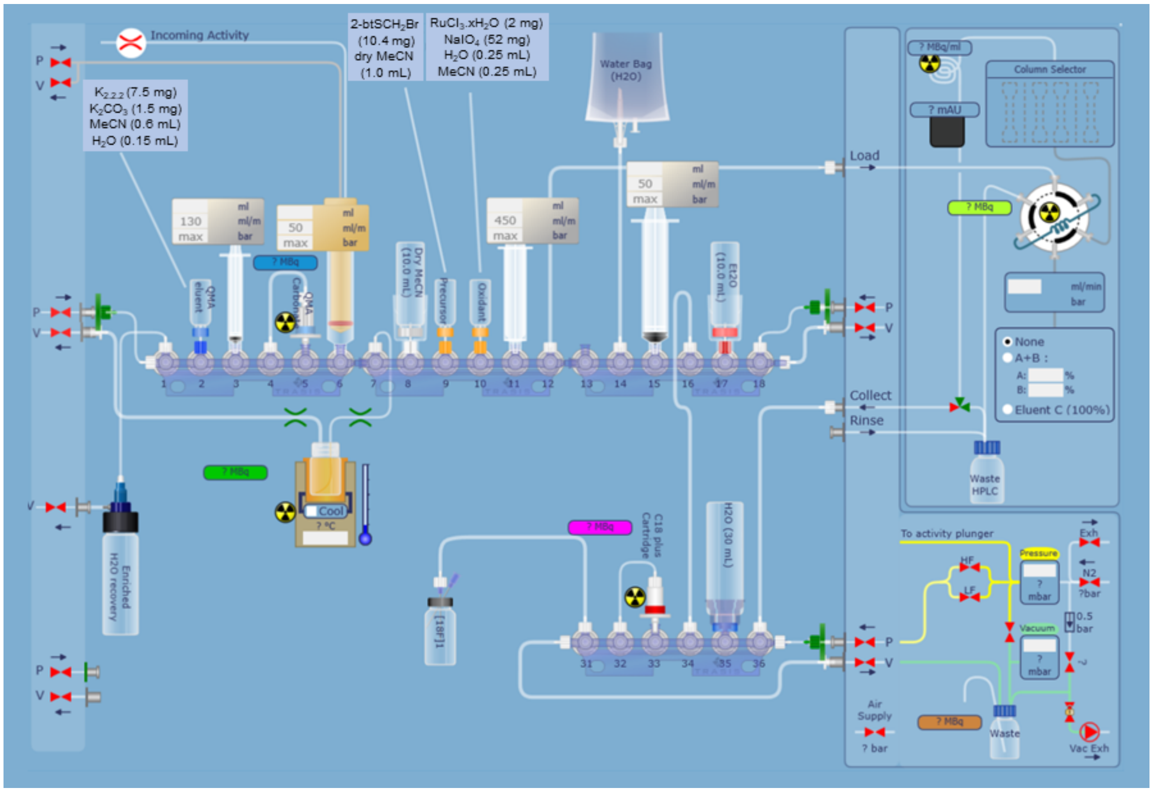


Cassette layout for the automated synthesis of [$^{18}$F]mono-btSOOF on the Trasis AllInOne platform.

The automated radiosynthesis of the [$^{18}$F]mono-btSOOF reagent was carried out using a Trasis AllinOne automated radiosynthesis platform, using a custom cassette set-up. A vial charged with $K_2CO_3$ (1.5 mg), Kryptofix® 222 (7.5 mg), MeCN (0.6 mL) and $H_2O$ (0.15 mL) was inserted at position 2. A vial charged with 2-((bromomethyl)thio)benzo[*d*]thiazole (10.4 mg, 0.04 mmol) and MeCN (1.0 mL) was inserted at position 9. A vial charged with $RuCl_3$ hydrate (2 mg), $NaIO_4$ (52 mg), $H_2O$ (0.25 mL) and MeCN (0.25 mL) was inserted at position 10. Two vials were filled with dry MeCN (approximately 10 mL) and placed in positions 8 and 17. A water bag was placed in position 14. The HPLC collection vial was filled with $H_2O$ (20 mL) and placed in position 34. A Waters Sep-Pak AccellPlus QMA Carbonate Plus Light Cartridge was pre-condtioned with $H_2O$ prior to use and placed in position 5. A Waters Sep-Pak C18 Plus Short cartridge was activated with EtOH (10 mL) and then $H_2O$ (10 mL) prior to use and was placed in position 33.

[$^{18}$F]Fluoride in [$^{18}$O]water was received directly from the cyclotron into the Trasis AllInOne radiosynthesis platform (Starting activity 100 GBq after 110 GBq from end of bombardment). The [$^{18}$F]fluoride was separated from the water by trapping on the QMA cartridge, followed by elution with the solution from the vial in position 2 into the reactor. The [$^{18}$F]fluoride was then dried, and once this was complete, the $^{18}$F-fluorination of((bromomethyl)thio)benzo[*d*]thiazole proceeded in the same reactor at 110 °C for 10 minutes. The reactor was then cooled and the contents of the vial at position 10 were added to the same reactor. The oxidation reaction proceeded at 35 °C for 5 minutes. The crude reaction mixture was diluted with $H_2O$ and MeCN and transferred to the HPLC injection loop for semi-preparative reverse phase HPLC purification (MeCN:$H_2O$ = 1:1, Q = 4 mL min$^{-1}$) with a Phenomenex Synergi Hydro-RP 4 µm 80 Å LC column (250 x 10 mm). The desired purified product ($t_R$ = 9.0-9.5 min) was collected in the vial at position 34. This solution was diluted further with $H_2O$ for reformulation using a C18 Plus cartridge. [$^{18}$F]mono-btSOOF was then eluted with MeCN (2.0 mL) into a vial. An aliquot of this product was taken for radio-HPLC analysis to determine the molar activity. Yield (from end of bombardment) and molar activity data are given in the table below. For use in a protein labelling reaction, [$^{18}$F]mono-btSOOF was eluted from the cartridge into the corresponding reaction vial (3.0 mL V-vial) with $Et_2O$ (~1.2 mL total volume) and concentrated to dryness under reduced pressure (supplemented with a flow of nitrogen).

Molar activity of [$^{18}$F]mono-btSOOF synthesised on a Trasis AllinOne platform:

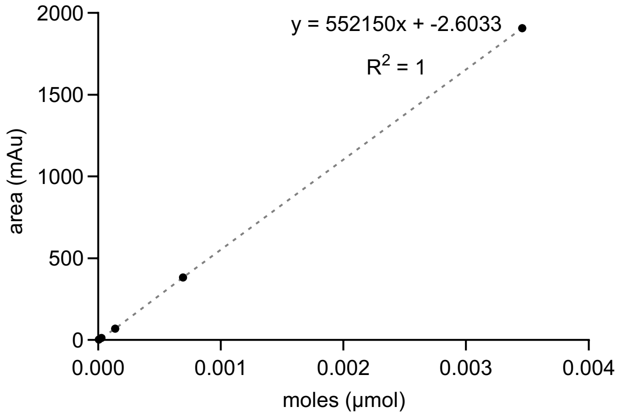


Run 1

| Measurement | Activity injected (MBq, d.c.) | Peak area (mAu*s) | S4 injected (μmol) | $A_m$ (GBq μmol$^{-1}$) |
|---|---|---|---|---|
| **1** | 1.22 | 4.3 | 7.8 x 10$^{-6}$ | 156 |
| **2** | 1.02 | 4.3 | 7.8 x 10$^{-6}$ | 131 |
| **3** | 1.00 | 3.8 | 6.9 x 10$^{-6}$ | 145 |
| | | | Average | 144 ± 10 |

Run 2

| Measurement | Activity injected (MBq, d.c.) | Peak area (mAu*s) | S4 injected (μmol) | $A_m$ (GBq μmol$^{-1}$) |
|---|---|---|---|---|
| **1** | 1.13 | 10.8 | 1.96 x 10$^{-5}$ | 57.7 |
| **2** | 1.17 | 10.4 | 1.88 x 10$^{-5}$ | 61.9 |
| **3** | 1.11 | 10.2 | 1.88 x 10$^{-5}$ | 60.0 |
| | | | Average | 59.9 ± 1.7 |

Hence, average molar activity is 102 ± 42 GBq μmol$^{-1}$ ($n$ = 2).

HPLC used a Agilent 1200 LC system with LabLogic gamma-RAM Model 4 detector, Flow rate = 1.0 mL/min, temperature = 25 °C HPLC gradient: water/MeCN, Phenomenex Synergi 4 μm Hydro RP 80Å 150 x 4.6 mm LC column.

Solvent gradient details:

0-1 min (25% MeCN) isocratic

1-10 min (25% MeCN to 95% MeCN) linear increase

10-16 min (95% MeCN) isocratic

16-18 min (95% MeCN to 25% MeCN) linear decrease

18-20 min (25% MeCN) isocratic.

#### 3.4 Protein Radiochemistry

**General protein reaction protocol**

Protein solutions and solvents were degassed either in a glovebox (<10 ppm $O_2$) for at least 8 hours or by purging with nitrogen for at least 15 min. For maintaining anaerobic conditions with a glovebox, the photocatalyst and iron additive were weighed and then transferred into the glovebox where stock solutions were prepared with degassed solvents. Protein reactions (≤500 µL) were prepared in 3.0 mL V-vials (Wheaton clear glass V-vials) by diluting the Dha-containing protein solution in the buffer of choice to reach the desired protein concentration, followed by sequential addition of photocatalyst, iron additives and DMSO (<10% of total reaction volume) in the glovebox. The reactions were then mixed thoroughly with a pipette, capped with 18 mm PTFE/silicone septa inserted in the screw caps and moved out of the glovebox. The 3.0 mL V-vial (sealed with a PTFE/silicone septum) containing the dried and purified sulfone reagent was purged with nitrogen for at least 1 min. A nitrogen balloon was inserted into the vial with the protein solution. Under nitrogen atmosphere, the protein solution was transferred into the V-vial containing the sulfone reagent and the vial was gently shaken before irradiation. For cases where a glovebox is not readily accessible, anaerobic conditions were achieving by purging the protein reactions (≤500 µL) in 7.0 mL V-vials with a nitrogen or argon balloon for at least 20 min after mixing with other reaction reagents (pre-dissolved in degassed $H_2O$) and degassed DMSO. The now degassed protein reactions were transferred into vials containing the sulfone reagent under nitrogen and then shaken before irradiation. A variable light intensity (10 – 50 W) photobox was used with blue LEDs arranged in series to allow up to 3 reactions (in 3.0 mL Wheaton V-vials) at a time. Cooling fan is attached to the photobox for temperature control especially for reactions requiring >20 min. After irradiation, an aliquot of the crude reaction mixture was obtained and diluted into $H_2O$ for radio-HPLC analysis to confirm $^{18}F$-labelling and calculate RCY. Before any animal studies, the crude mixture was purified following incubation with EDTA and then using PD MiniTrap G-25 (GE Healthcare) desalting columns (protein recovery ~50-60%). Aliquots can also be obtained immediately after irradiation for LC-MS analysis after isotopic decay.

**Formation of *X.l.* Histone H3.NTEV-[$^{18}$F]Fha2 with [$^{18}$F]mono-btSOOF**

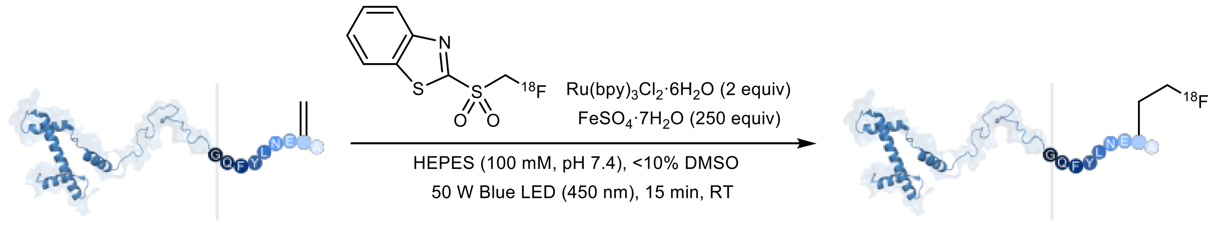


In the glovebox, degassed HEPES buffer (100 mM, pH 7.4) was added to a glass V-vial containing *X.l.* Histone H3.NTEV-Dha2 such that the final concentration is 125 μM in a total reaction volume of 500 μL. $Ru(bpy)_3Cl_2{\cdot}6H_2O$ (4 μL of a 31 mM stock prepared fresh in degassed $H_2O$, 2 equiv), $FeSO_4{\cdot}7H_2O$ (30 μL of a 520 mM stock prepared fresh in degassed $H_2O$, 250 equiv) and DMSO (<10% v/v) were added sequentially and the V-vial was sealed before transferring out of the glovebox. Under nitrogen, the protein solution was transferred into a nitrogen-filled V-vial containing the purified [$^{18}$F]mono-btSOOF, shaken and then irradiated with blue LED light (50 W) for 15 min. A sample of the crude mixture was then analysed by radio-HPLC to determine RCY as shown below. An aliquot was also taken for LC-MS analysis after decay to check for protein oxidation.

Crude HPLC analysis (Method B):

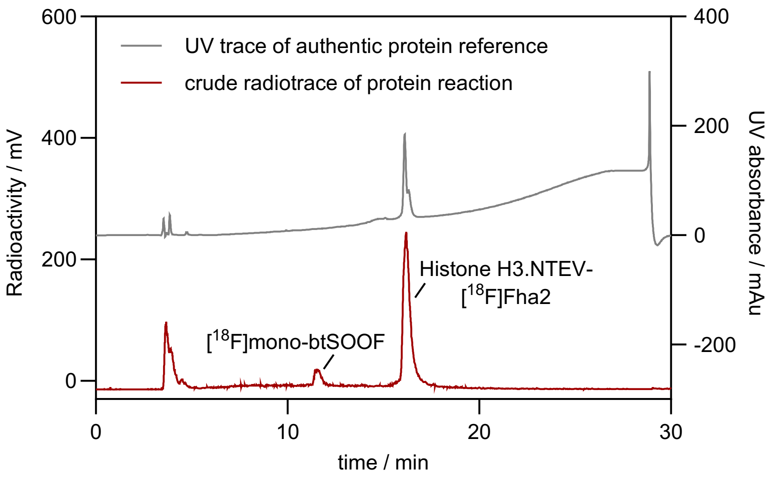

ESI-MS analysis of *X.l.* histone H3.NTEV:

Calculated H3 Dha mass: 16004 Da

Observed mass: 16003 Da (+16 Da is methionine oxidation)

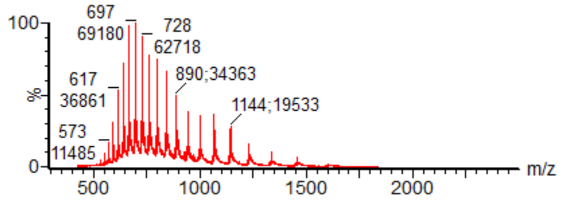


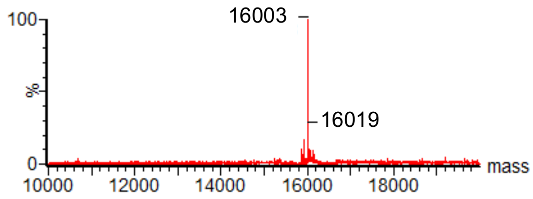

#### TEV cleavage on Histone H3.NTEV-[$^{18}$F]Fha2

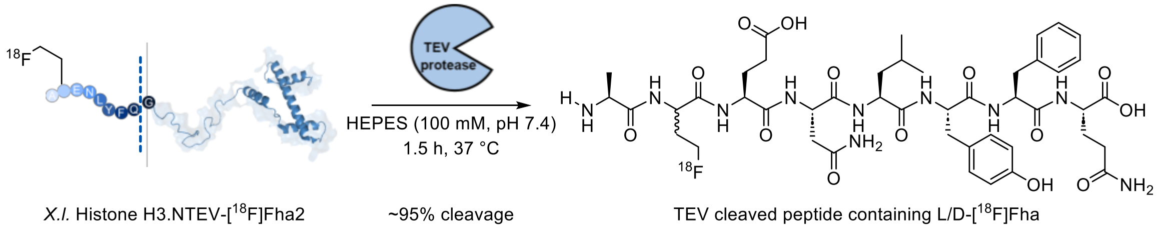


At the end of the reaction, the crude reaction mixture was loaded onto a PD-10 MiniTrap desalting column (pre-equilibrated with 100 mM HEPES, pH 7.4). $^{18}$F-labelled H3.NTEV was collected in the first 700 µL of buffered solution and 180 µL of the purified protein was incubated with 10 µL of TEV protease and 20 µL of TEV protease buffer (New England Biolabs) at 37 °C for 1.5 h. A sample was then analysed by radio-HPLC.

TEV cleavage HPLC analysis (Method D):

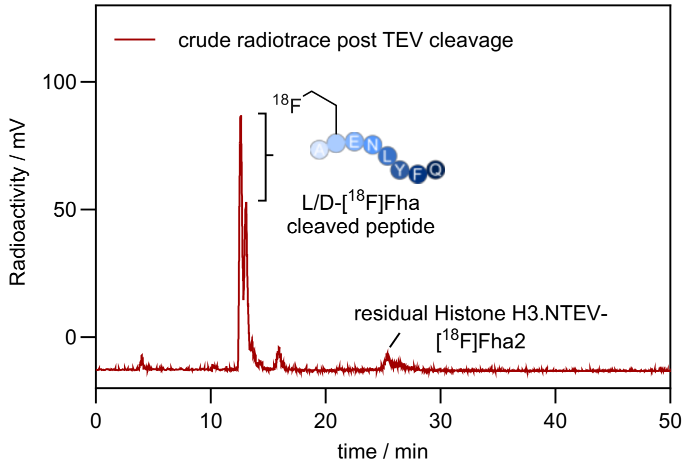


LC-MS analysis post-TEV cleavage:

ESI-MS analysis of TEV cleaved *X.l.* histone H3.NTEV:

Calculated TEV cleaved H3 mass: 15069 Da

Observed mass: 15068 Da (+16 Da is methionine oxidation)

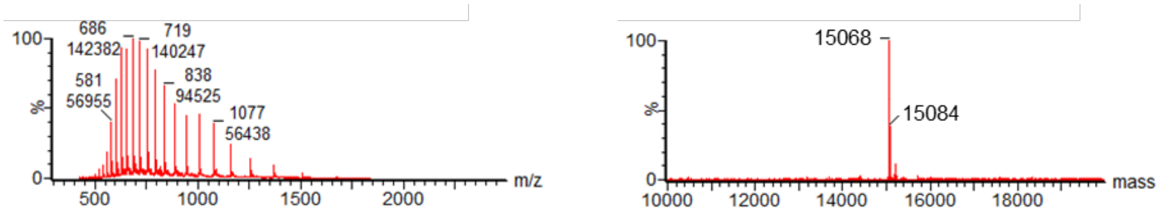

**Imine formation on Histone H3.NTEV-[$^{18}$F]Fha2 in the absence of iron**

The absence of any iron additive while keeping all other conditions outlined earlier constant resulted in the release of a single radiolabelled peptide product post-TEV cleavage which corresponds to imine formation (~69% cleavage following a 30 min incubation).

HPLC analysis post-TEV cleavage (Method B):

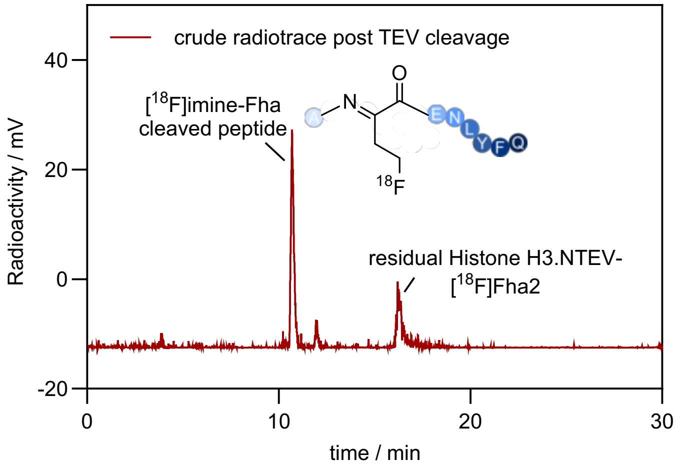

**Formation of *X.l.* Histone H3.NTEV-[$^{18}$F]*b*Fha2 with [$^{18}$F]btSOOF**

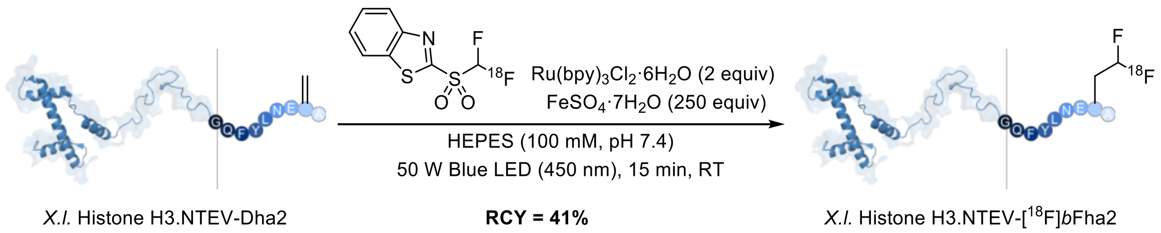


In the glovebox, degassed HEPES buffer (100 mM, pH 7.4) was added to a glass V-vial containing *X.l.* histone H3.NTEV-Dha2 such that the final concentration was 125 µM in a total reaction volume of 400 µL. $Ru(bpy)_3Cl_2·6H_2O$ (3.2 µL of a 31 mM stock prepared fresh in degassed $H_2O$, 2 equiv), $FeSO_4·7H_2O$ (24 µL of a 520 mM stock prepared fresh in degassed $H_2O$, 250 equiv) and DMSO (<10% v/v) were added sequentially and the V-vial was sealed before transferring out of the glovebox. Under nitrogen, the protein solution was transferred into a nitrogen-filled V-vial containing the purified [$^{18}$F]btSOOF, shaken and then irradiated with blue LED light (50 W) for 15 min. A sample of the crude mixture was then analysed by radio-HPLC to determine RCY.

Crude HPLC analysis (Method B):

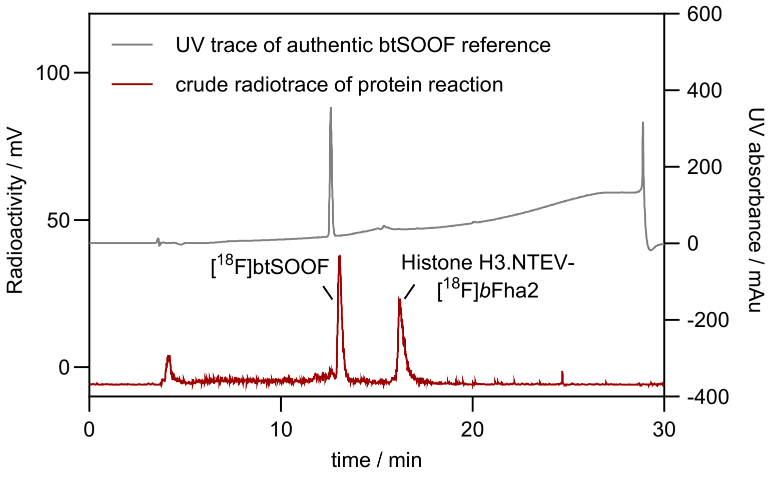

**TEV cleavage on Histone H3.NTEV-[$^{18}$F]*b*Fha2**

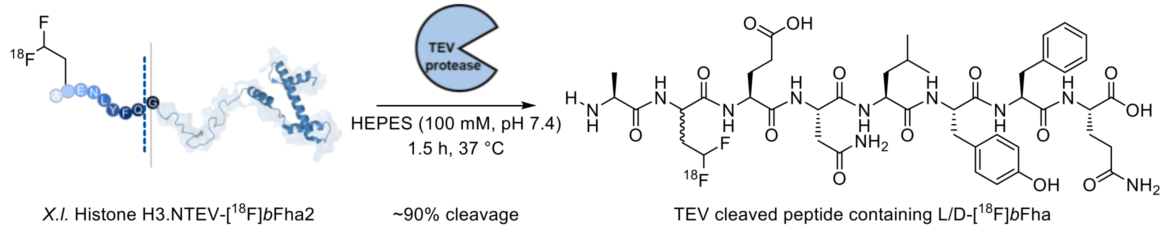


After subjecting the reaction to conditions outlined previously for the $^{18}$F-labelling of H3.NTEV-Dha2 with [$^{18}$F]btSOOF, the crude reaction mixture was loaded onto a PD-10 MiniTrap desalting column (pre-equilibrated with 100 mM HEPES, pH 7.4). $^{18}$F-labelled H3.NTEV was collected in the first 700 μL of buffered solution and 180 μL of the purified protein was incubated with 10 μL of TEV protease and 20 μL of TEV protease buffer (New England Biolabs) at 37 °C for 1.5 h. A sample was then analysed by radio-HPLC.

HPLC analysis post-TEV cleavage (Method C):

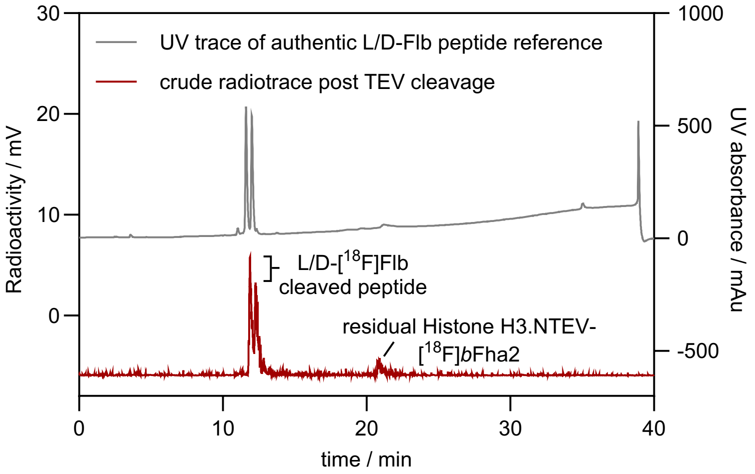

#### Formation of Human Histone eH3.1-[$^{18}$F]Fha4 with [$^{18}$F]mono-btSOOF

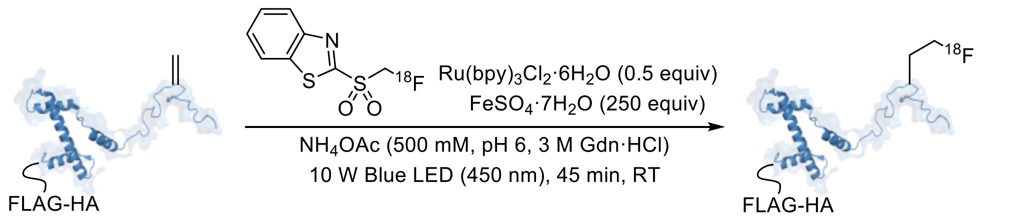


Human Histone eH3.1-Dha4 **RCY = 67 5% (*n* = 2)** Human Histone eH3.1-[$^{18}$F]Fha4

In the glovebox, degassed $NH_4OAc$ buffer (500 mM, pH 6, 3 M Gdn·HCl) was added to a glass V-vial containing human histone eH3.1-Dha4 such that the final concentration was 125 μM in a total reaction volume of 500 μL. $Ru(bpy)_3Cl_2·6H_2O$ (1 μL of a 31 mM stock prepared fresh in degassed $H_2O$, 0.5 equiv), $FeSO_4·7H_2O$ (30 μL of a 520 mM stock prepared fresh in degassed $H_2O$, 250 equiv) and DMSO (<10% v/v) were added sequentially and the V-vial was sealed before transferring out of the glovebox. Under nitrogen, the protein solution was transferred into a nitrogen-filled V-vial containing the purified [$^{18}$F]mono-btSOOF, shaken and then irradiated with blue LED light (10 W) for 45 min. A sample of the crude mixture was then analysed by radio-HPLC to determine RCY.

Crude HPLC analysis (Method B):

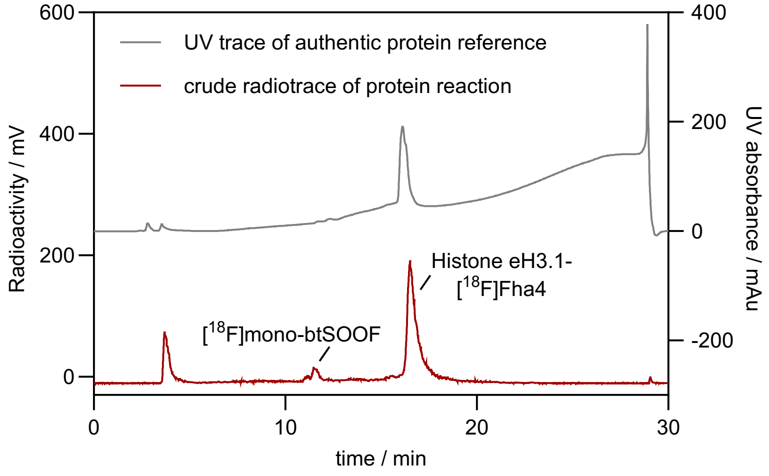

LC-MS analysis post-protein radiolabelling:

ESI-MS analysis of human histone eH3.1:

Calculated H3 Dha mass: 17686 Da

Observed mass: 17686 Da (+16 Da is methionine oxidation)

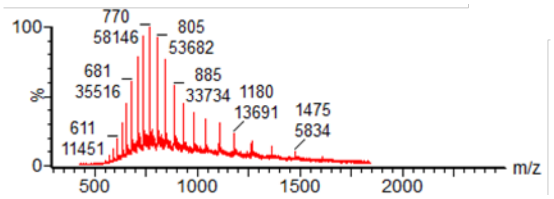


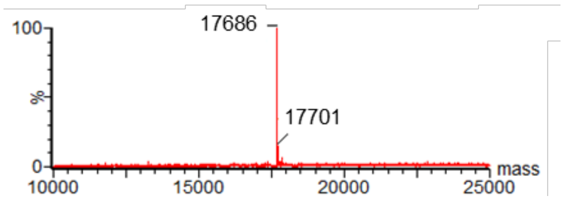


Effect of light flux and reaction times on the RCY of the protein $^{18}F$-labelling reaction:

| Human histone eH3.1-Dha4 | $Ru(bpy)_3^{2+}$ (equiv.) | $Fe^{2+}$ (equiv.) | Conditions | RCY |
|---|---|---|---|---|
| 125 µM | 2 | 250 | 50 W,<br>15 and 30 min | 57% (15 min)<br>58% (30 min) |
| 125 µM | 0.5 | 250 | 10 W,<br>30 and 45 min | 56% (30 min)<br>67 ± 5% (*n* = 2; 30 min) |

**Formation of purified mNfL-[$^{18}$F]Fha323 with [$^{18}$F]mono-btSOOF**

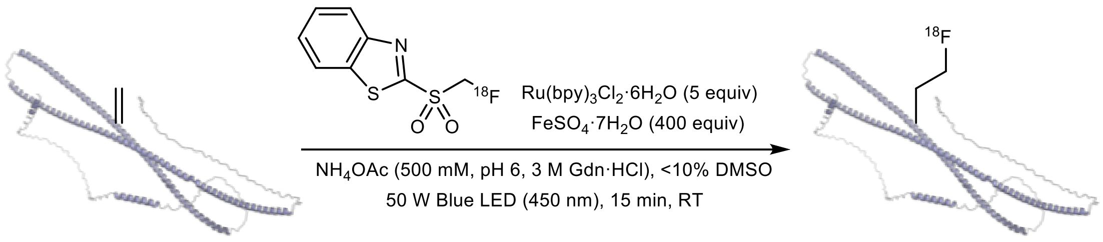


NfL-Dha323 **RCY = 39 6% (n = 2)** NfL-[$^{18}$F]Fha323

In the glovebox, degassed $NH_4OAc$ buffer (500 mM, pH 6, 3 M Gdn·HCl) was added to a glass V-vial containing mNfL-Dha323 such that the final concentration is 70 µM in a total reaction volume of 155 µL. $Ru(bpy)_3Cl_2·6H_2O$ (2 µL of a 27 mM stock prepared fresh in degassed $H_2O$, 5 equiv), $FeSO_4·7H_2O$ (10 µL of a 434 mM stock prepared fresh in degassed $H_2O$, 400 equiv) and DMSO (<10% v/v) were added sequentially and the V-vial was sealed before transferring out of the glovebox. Under nitrogen, the protein solution was transferred into a nitrogen-filled V-vial containing the purified [$^{18}$F]mono-btSOOF, shaken and then irradiated with blue LED light (50 W) for 15 min. A sample of the crude mixture was then analysed by radio-HPLC to determine RCY as shown below.

It is important to note that mNfL-Dha should not be kept in the freezer for more than one week at pH 6 before the $^{18}$F-protein labelling reaction. Instead, using freshly desalted mNfL-Dha in the $NH_4OAc$ reaction buffer at pH 6 will give higher RCY values. Increasing protein concentration and reaction volume will also give higher activity yield for pre-clinical applications.

Crude HPLC analysis (Method B):

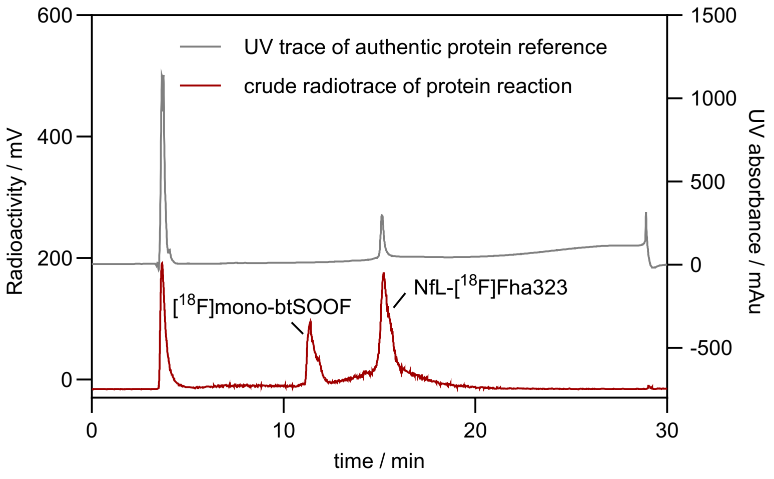


At the end of the reaction, 500 equiv. of EDTA (from a 0.5 M stock) was added and the crude reaction mixture was diluted up to a final volume of 500 µL before loading onto a PD MiniTrap G-25 desalting column (pre-equilibrated with PBS and 4 M urea). The first 200 µL elution volume was discarded. $^{18}$F-labelled NfL was collected in the next 400 µL of buffered solution. An aliquot was taken to check the radio- and UV purity.

Left: Radio-HPLC trace; Right: Corresponding UV trace (Method B)

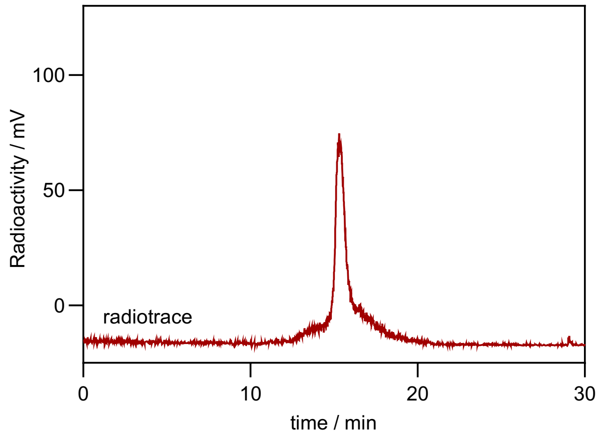


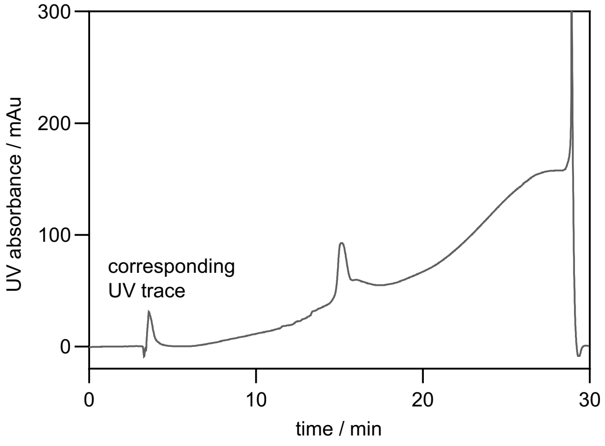

#### Formation of mNfL-[$^{18}$F]Fha323 for intravenous, intracerebral and intracerebroventricular administration

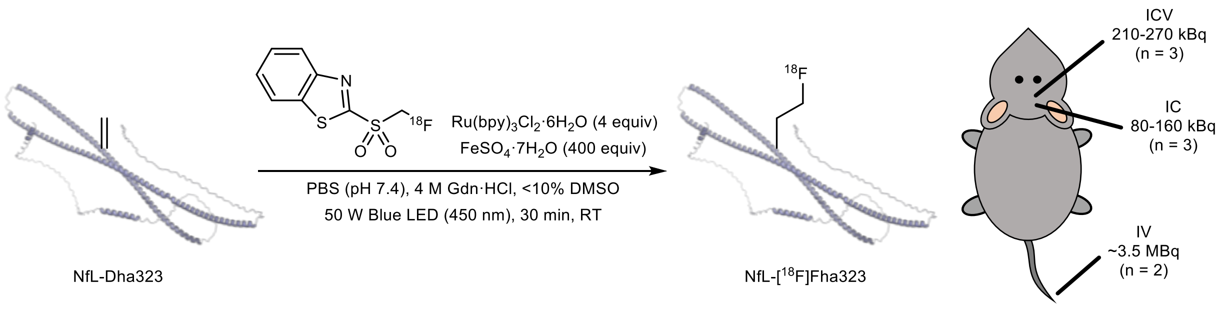


$^{18}$F-labelling on NfL-Dha323 for *in vivo* PET studies was performed with 90 µM mNfL-Dha323 in a total reaction volume of 300 µL, 4 equiv. of Ru(bpy)$_3$Cl$_2$·6H$_2$O, 400 equiv. of FeSO$_4$·7H$_2$O, DMSO (<10% v/v) and irradiation with blue LED at 50 W for 30 min. Degassed conditions was achieved using argon balloon in the absence of a glovebox. At the end of the reaction, 1000 equiv. of EDTA (from a 0.5 M stock) was added and the crude reaction mixture was diluted up to a final volume of 500 µL before loading onto a PD MiniTrap G-25 desalting column (pre-equilibrated with PBS and 4 M urea). The first 200 µL elution volume was discarded. $^{18}$F-labelled NfL was collected in the next 300 µL of buffered solution at an approximate concentration of 2.3 mg mL$^{-1}$ (for intravenous administration). Starting from dried [$^{18}$F]mono-btSOOF, an isolated activity yield of 10.2 ± 1.4% (n = 2) was obtained in 300 µL of the eluting buffer.

For $^{18}$F-labelled mNfL administered intracranially or intracerebroventricularly, the purified protein was concentrated to ~10 mg mL$^{-1}$ using an Amicon 10 kDa MWCO centrifugal filter to ensure sufficient activity concentration for PET imaging.

#### 3.5 Recombinant mNfL re-extraction from plasma

**mNFL re-extraction from plasma under *in vitro* condition**

Purified recombinant mNfL protein (300 µL) was added to isolated mouse plasma (200 µL) together with urea (urea final concentration reached 6 M) and fresh prepared Ni-NTA resin (200 µL). The mixture was incubated at 4 °C for 2 h then transferred into an empty column where the resin was washed with low concentration of imidazole buffer in 6 M urea. The target protein was subsequently eluted from the resin with high concentration of imidazole buffer in 6 M urea. Elution fractions were analyzed by SDS-PAGE and then Western blot using *anti*-HisTag antibody (Sigma, A5588). As shown in the figure below, clear protein bands corresponding to mNfL were observed in the elution fractions, suggesting that recombinant mNfL is relative stable and can be re-extracted from mouse plasma under *in vitro* condition.

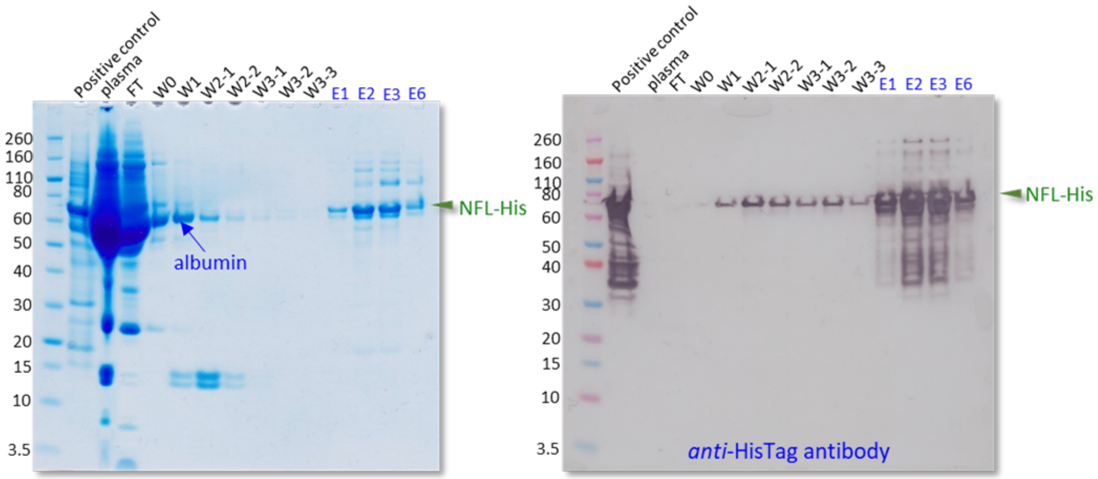

**mNFL re-extraction from plasma under *in vivo* condition**

Purified recombinant mNfL with a concentration of 23 µM was used for animal injection. Total 5 mice were sacrificed for the test: *1)* saline was injected and the mouse was killed immediately after the injection and the blood was extracted; *2)* mNfL protein 1.4 µg was injected and the mouse was killed after 30 min following blood extraction; *3)* mNfL protein 1.4 µg was injected and the mouse was killed after 60 min following blood extraction; *4)* mNfL protein 30 µg was injected and the mouse was killed immediately and the blood was extracted; *5)* mNfL protein 30 µg was injected and the mouse was killed after 60 min following blood extraction.

Protein mNfL re-extraction experiment: 200 µL of extracted mouse plasma 1, 2, 3, 4 and 5 was incubated with 200 µL fresh prepared Ni-NTA resin, respectively. Urea was added to each mixture to make the urea final concentration reached 6 M. Each mixture was then incubated at 4 °C for 2 h with shaking. The supernatant was removed and the resin was washed with 20 mM Tris, pH 7.8, 20 mM imidazole, 6 M urea several times. The target protein was eluted with 100 µL of elution buffer (20 mM Tris, pH 7.8, 500 mM imidazole, and 6 M urea). The eluted fractions were analysed by SDS-PAGE and Western blot using *anti*-HisTag (Sigma A5588) and *anti*-NFL (Genetex GTX134085) antibodies.

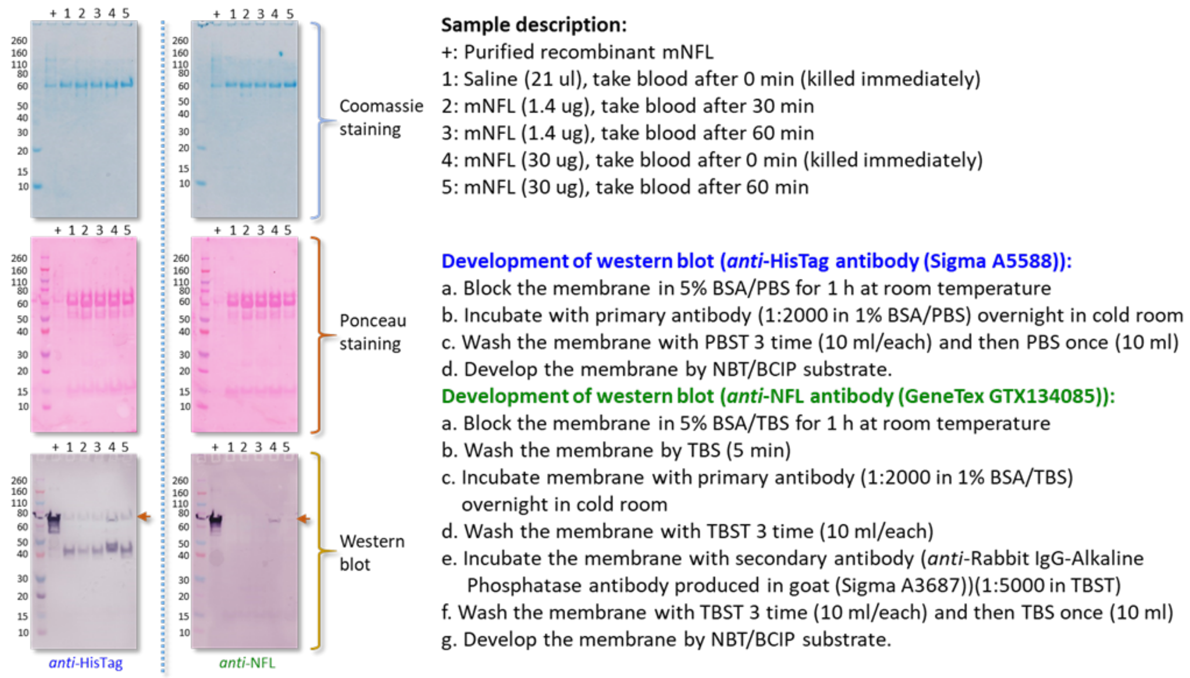


SDS-PAGE and western blot (using *anti*-HisTag antibody) to check the extraction of recombinant NFL-His from mouse plasma by Ni-NTA resin under *in vivo* condition.

##### 3.6 PET imaging of mNfL

All animal studies were approved by the national Project Authorization Board in Finland (license numbers: ESAVI-45421-2022 and ESAVI-12143-2022) and carried out in compliance with the European Union Directive 2010/EU/63 on the protection of animals used for scientific purposes.

Mice (C57BL/6JOlaHsd, Envigo) were anesthetized via inhalation of 3% isoflurane gas (0.5 L/min $O_2$) and placed on a warm heat mat in the prone position. For intravenous injection, the tail of the mouse was warmed with a heat lamp before the injection of $^{18}$F-NfL via lateral tail vein. For intracranial and intracerebroventricular injection, a capillary needle with a 50 µM lumen was used using a Stoelting Digital New Standard Stereotaxic Device and coordinates as follows: for i.c. injection, AP, +0.5 mm; ML, +2.0 mm; DV, -2.5 mm and for i.c.v., AP, +0.0 mm; ML, +0.8 mm; DV, -2.0 mm.

2 mice were used for i.v. injection and 3 mice each were for used i.c. and i.c.v. injections. Dynamic PET images (120 min) were acquired after $^{18}$F-NfL injections with Molecubes β-cube (PET) and X-cube (CT) (MOLECUBES NV, Ghent, Belgium). PET aquisition was performed in list-mode and reconstructed into time frames 18 × 10s, 17 × 60s, 20 × 300 s following i.v. injection and 30 × 10s, 15 × 60s, 10 × 300 s, 5 x 600 s following i.c.v. and i.c. injections. Image analysis was performed with the Carimas software (Turku PET Centre, Turku, Finland). Nonlinear regression analyses were performed using GraphPad Prism (GraphPad Software).

2 h after injection of $^{18}$F-NfL, mice were euthanized. Fluorine-18 uptake in selected tissue was determined and reported as a percentage of the injected dose per gram of tissue (%ID/g) decay-corrected to the time of injection using a Wizard 2480 automated gamma counter (Perkin Elmer). The %ID/g results are presented as mean ± standard deviation using GraphPad Prism 10 software. Values for ‘blood’ reflect unfractionated blood sample i.e., include plasma and erythrocytes.

To measure changes in %ID/mL over time in a given tissue, central parts of the organs were outlined on at least five CT slices in the coronal plane registered to the dynamic PET series in HERMIA Hybrid Viewer (v6.1, Hermes Medical Solutions AB, Stockholm, Sweden) for each mouse. These volumes were then transferred to the PET image series and mean activity concentrations within each volume were extracted for each time point. The %ID/mL were then calculated based on the injected radioactivity and presented as mean ± standard error of the mean (SEM) using GraphPad Prism 10 software.

Blood half-life measurement post-i.v. administration based on a 1-phase decay:

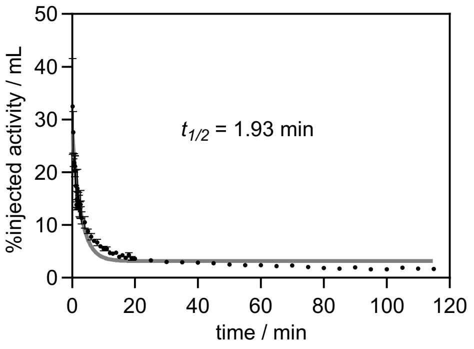


Time activity curve post-i.c. administration expressed as %ID/mL (*n* = 3):

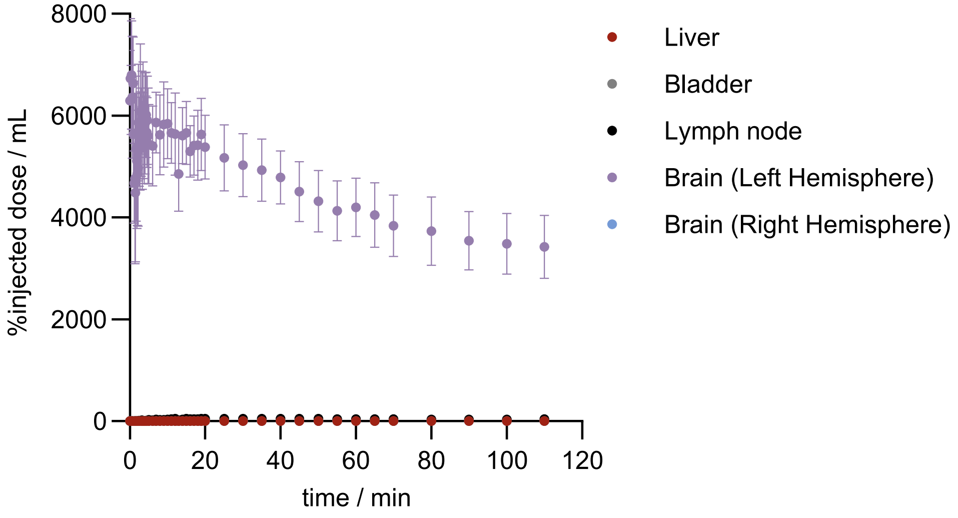

Time activity curve post-i.c.v. administration expressed as %ID/mL ($n$ = 3):

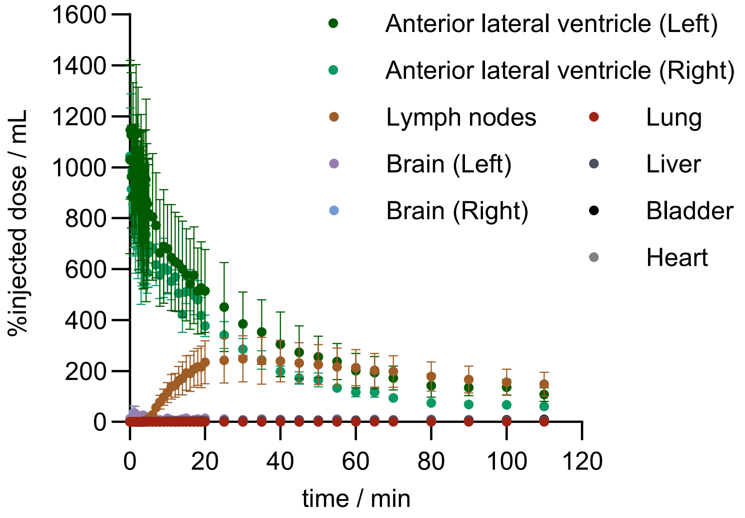


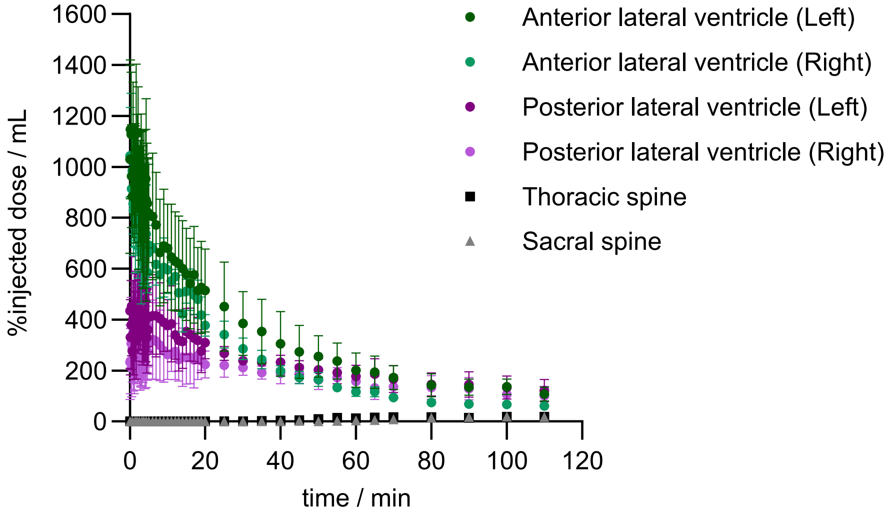

Determination of flow rate from thoracic to sacral spine ($n$ = 3) using time activity curves expressed as %ID/mL:

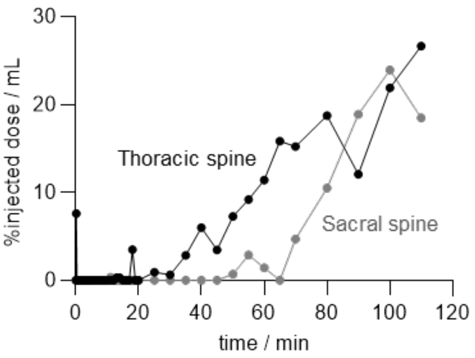

Flow rate: 25 mm / 20 min

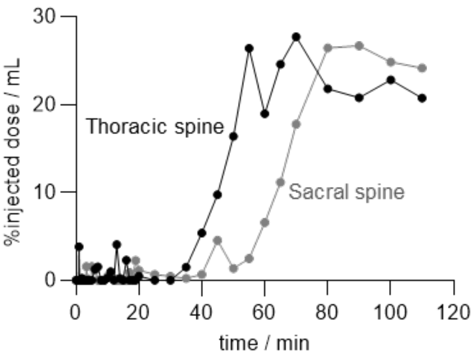

Flow rate: 21 mm / 25 min

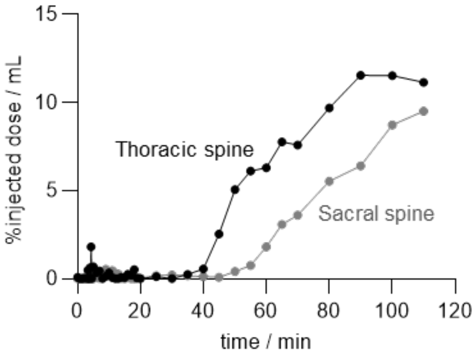

Flow rate: 23 mm / 15 min

**Relative Activity Uptake**

Biodistribution in the brain and CNS following i.c. and i.c.v. injections were expressed as relative activity uptake with respect to time, as seen in Figure 4. The relative activity uptake was calculated by normalising the %ID/mL of each organ to the organ with the highest $^{18}F$ uptake, which was set as 100% for each mice (typically near the start of a scan i.e., maximum activity concentration). The uptake in other organs with time were then calculated as a percentage relative to this maximum value. Statistical analysis were calculated using GraphPad Prism 10 software and expressed as mean ± standard error of the mean (SEM).

**mNfL re-extraction from spleen and liver after PET scanning and $^{18}$F decay**

Based on the organ counting data, the spleen and liver exhibited the highest accumulation of radioactivity/g after i.v. administration of [$^{18}$F]NfL. Hence, recombinant mNfL re-extraction experiments were performed on these harvested organs (after complete $^{18}$F decay) for gel electrophoresis. As shown in the figure below, the organs were homogenised using a Dounce homogenizer and the target protein was extracted by freshly prepared Ni-NTA resin (250 µL). Following a resin wash and protein elution, subsequent fractions were then analysed on SDS-PAGE and Western blot using *anti*-HisTag (SigmaAldrich A5588) and *anti*-NfL (GeneTex GTX134085) antibodies.

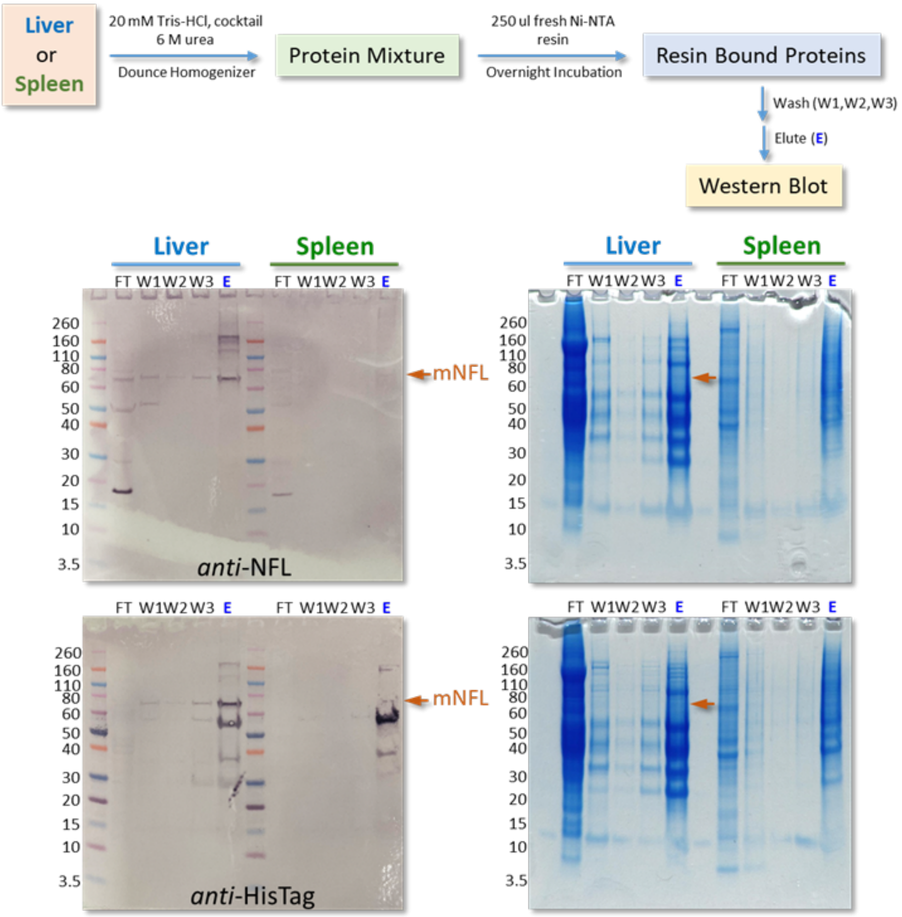


A clear protein band around the expected region of mNfL was observed by Western on the treated liver tissue, and to a lesser extent, the spleen corroborating the *in vivo* biodistribution studies after a 2 h incubation.

#### 3.7 Measuring aggregation kinetics of aggregation-prone proteins using absorbance measurements

WT and Fha-modified NfL proteins were buffer exchanged using a Amicon 10K MWCO centrifugal filter into labelling buffer (3 M Urea, 20 mM PIPES, 200 mM NaCl, 1 mM DTT, 2 mM $MgCl_2$, pH 8.0). The samples were then centrifuged at 100,000 g for 30 min, 4 °C and then ~95% of the solution was removed and the "pellet" discarded to give a solution with final concentration of 13.3 mg/mL for WT and 4.47 mg/mL for Fha-modified, both in 82 µL. WT mNfL was the diluted to the desired concentration of 4.47 mg/mL (75 µL stock taken and added to 148 µL buffer) and then stored on ice. Each reaction was carried out in 42 µL.

SFB-modified NfL samples with various levels of conjugation were generated following the protocol shown below. Two WT controls were also treated the same but with neat DMSO added, as well as the sample of Fha-modified NfL.

At this point the samples were loaded into a 96-well half volume UV-clear plate, 7.8 µL of protein sample in 5 x wells for each condition while the plate was kept on ice. Then, 92.2 µL of ice-cold buffer (0 M Urea, 20 mM PIPES, 200 mM NaCl, 1 mM DTT, 2 mM $MgCl_2$, pH 6.8) was added and then each well carefully mixed. Specifically, to each well used was added 92.2 µL to which 7.8 µL of the mNfL stock were then added to a final concentration of mNfL 0.35 mg/mL, urea 0.23 M at 4 °C (12.8x dilution). The plate was then added to the plate-reader set to record abs = 350 nm every 2 s, at 30 °C with shaking 2 s orbital shaking. The absorbance was recorded over 1 h. 5 wells were used for each condition (see figure below): WT mNfL in 3 M Urea, WT mNfL in the aggregation buffer, WT mNfL +2 equiv. SFB, WT mNfL +5 equiv. SFB, WT mNfL +10 equiv. SFB, mNfL-Fha323, buffer only, buffer + 10 equiv. SFB.

**Data analysis**

The data shown in Figure 5a were fitted to a simple aggregation model that assumes conversion of monomers to larger aggregates and leads to expected behaviour of A350 = A0 (1-exp(-(kt-t0)) where t0 is the dead time in the experiment. As all samples were initiated at the same time, t0 is expected to be constant for all samples. Empirically the globally fitted value was 15 minutes.

Absolute absorbance at 350 nm is presented in the figure below as mean ± standard deviation using GraphPad Prism 10 software. The Δabsorbance shown in Figure 5a was calculated by normalising the absorbance measurements for each NfL sample to the aggregation buffer, which was set to 0.

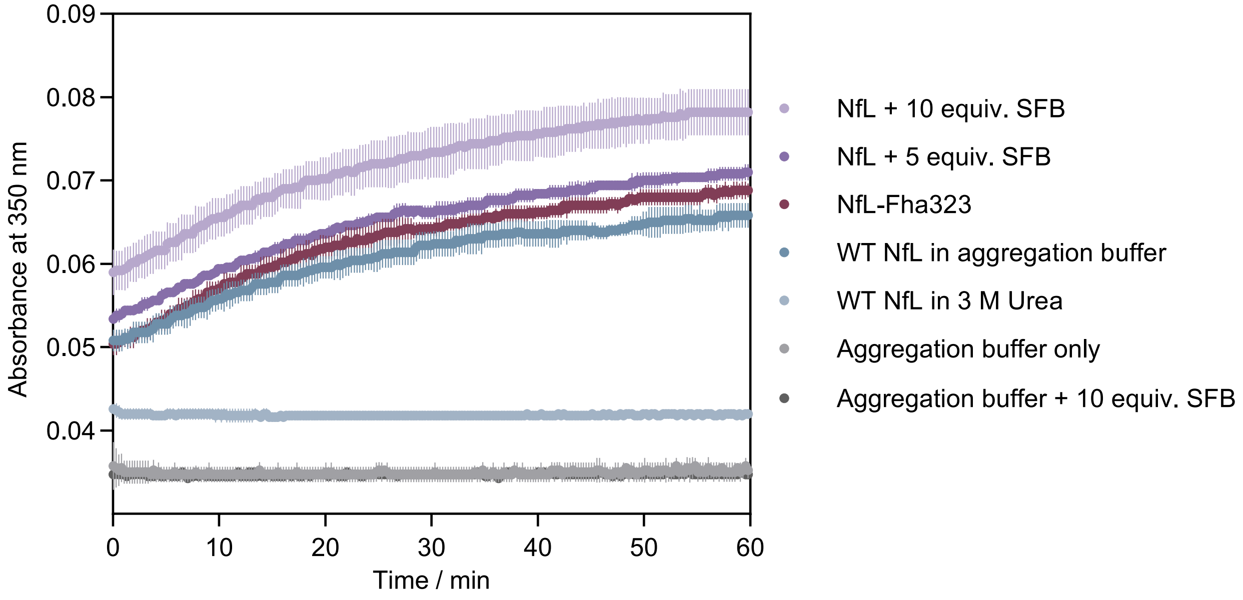


Employing 5 equiv. of SFB generated SFB-modified NfL with an average conjugation number of ~1.4. Increasing to 10 equiv. of SFB raised the average conjugation number to ~3.1.

Aggregation buffer: 0.23 M Urea, 20 mM PIPES, 200 mM NaCl, 1 mM DTT, 2 mM $MgCl_2$, pH 6.8.

**Generation of SFB-conjugated NfL for aggregation studies using absorbance measurements**

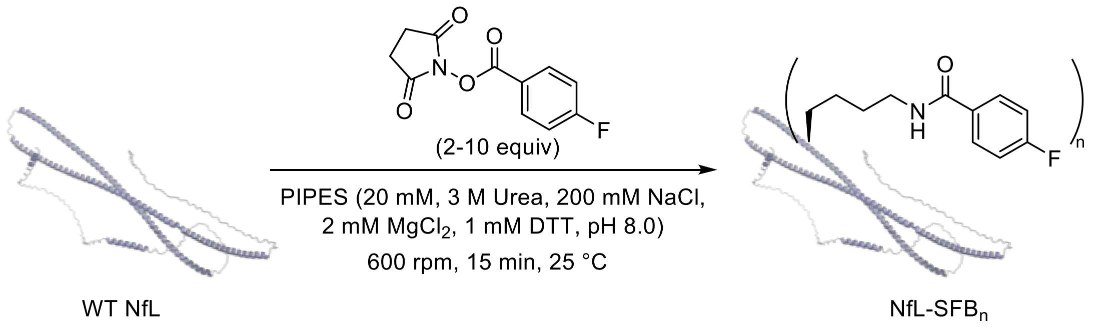


To generate SFB-modified NfL with various levels of conjugation, to the solution of WT sample (42 µL, 71 µM, 4.47 mg/mL) was added SFB (2, 5 or 10 equiv. in DMSO; 0.42 µL at 14.2 mM, 35.5 mM and 71.0 mM respectively) with shaking at 600 rpm at 25 °C for 15 min before being stored on ice.

Representative LC-MS analysis (ion series and magnification of the deconvoluted spectrum):

WT mNfL:

Observed mass: 62386 and 62424 g mol$^{-1}$

Calculated mass for 1x carbamylation: 62387 g mol$^{-1}$

Calculated mass for 2x carbamylation: 62430 g mol$^{-1}$

Observed mass: 62467 and 62506 g mol$^{-1}$

Calculated mass for 1x carbamylation and 1x phosphorylation: 62467 g mol$^{-1}$

Calculated mass for 2x carbamylation and 1x phosphorylation: 62510 g mol$^{-1}$

Observed mass: 62547 and 62587 g mol$^{-1}$

Calculated mass for 1x carbamylation and 2x phosphorylation: 62547 g mol$^{-1}$

Calculated mass for 2x carbamylation and 2x phosphorylation: 62590 g mol$^{-1}$

Observed mass: 62627 g mol$^{-1}$

Calculated mass for 1x carbamylation and 3x phosphorylation: 62627 g mol$^{-1}$

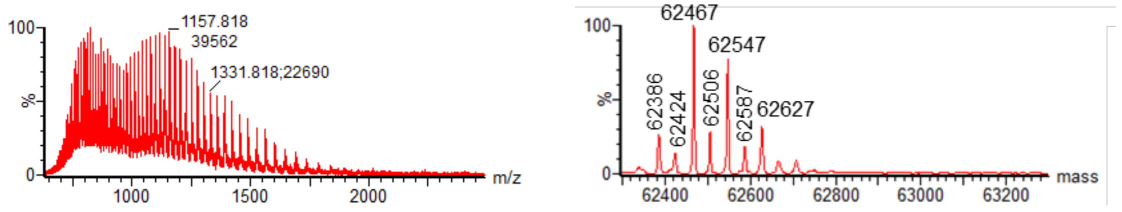


Incubation of WT mNfL with 5 equiv. of SFB resulted in an average conjugation number of ~1.4. ESI analysis showed extended of modification:

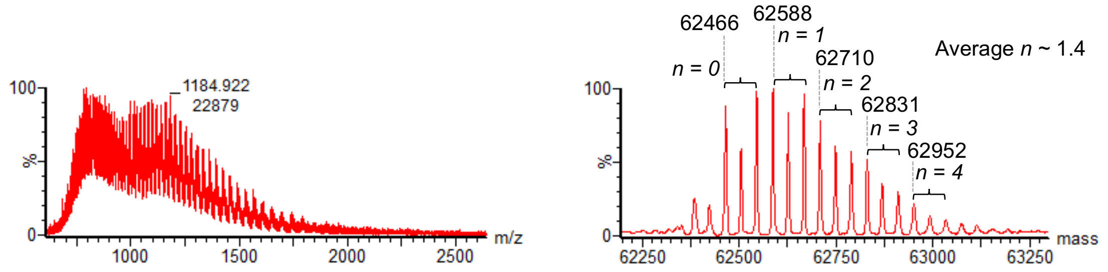


Incubation of WT mNfL with 10 equiv. of SFB resulted in an average conjugation number of ~3.1. ESI analysis showed extended of modification:

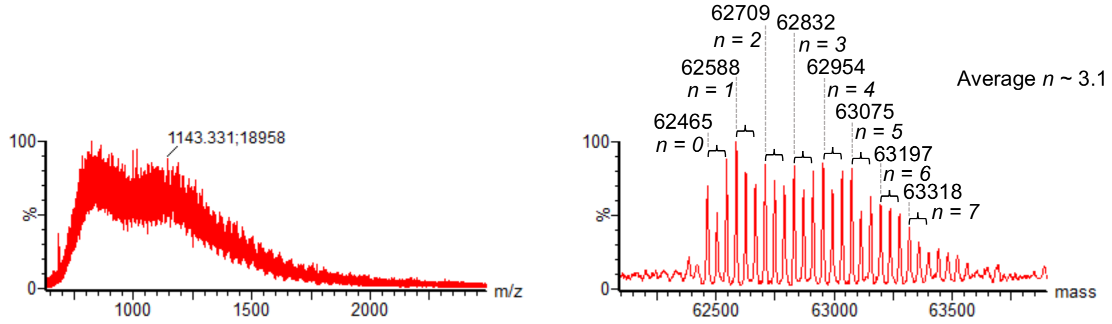

#### 3.8 Measuring aggregation kinetics of aggregation-prone proteins with NMR Spectroscopy

It is important to note that in order to compare aggregation kinetics, all variants of NfL (WT, Fha- and SFB-modified) must originate from the same protein expression batch.

**Generation of Fha-modified NfL**

NfL-Fha323 was generated following the protocol described previously. The reaction was repeated with a further 5 batch of samples. After incubation with EDTA, all reactions were combined and diluted with the reaction buffer to a final volume of 1 mL. The mixture was then loaded onto a PD MiniTrap G-25 desalting column (pre-equilibrated with PBS in $D_2O$ with 4 M urea and 2 mM DTT; 2 x 500 mL per column). The first 500 μL eluted from the desalting column was collected, protein concentration was measured using a NanoPhotometer® and then concentrated with an Amicon 10 kDa MWCO centrifugal filter (pre-washed with $D_2O$) for subsequent NMR studies.

ESI analysis of mNfL-Fha323:

Observed mass: 62389 and 62429 g $mol^{-1}$

Calculated mass for 1x carbamylation: 62387 g $mol^{-1}$

Calculated mass for 2x carbamylation: 62430 g $mol^{-1}$

Observed mass: 62469 and 62509 g $mol^{-1}$

Calculated mass for 1x carbamylation and 1x phosphorylation: 62467 g $mol^{-1}$

Calculated mass for 2x carbamylation and 1x phosphorylation: 62510 g $mol^{-1}$

Observed mass: 62548 and 62588 g $mol^{-1}$

Calculated mass for 1x carbamylation and 2x phosphorylation: 62547 g $mol^{-1}$

Calculated mass for 2x carbamylation and 2x phosphorylation: 62590 g $mol^{-1}$

Observed mass: 62627 and 62667 g $mol^{-1}$

Calculated mass for 1x carbamylation and 3x phosphorylation: 62627 g $mol^{-1}$

Calculated mass for 2x carbamylation and 3x phosphorylation: 62670 g $mol^{-1}$

Observed mass: 62706 g $mol^{-1}$

Calculated mass for 1x carbamylation and 4x phosphorylation: 62707 g $mol^{-1}$

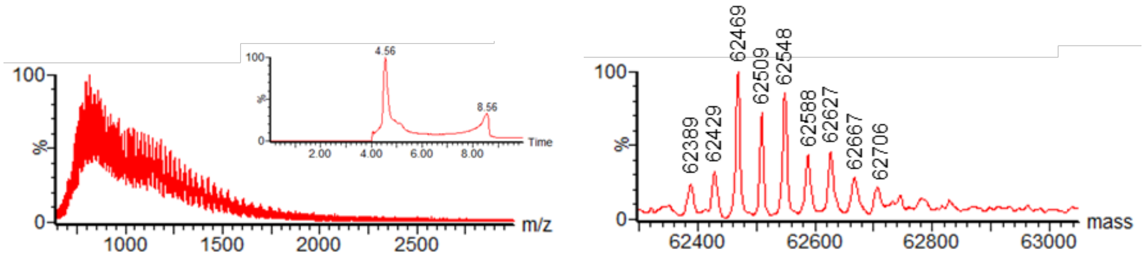

**Generation of SFB-conjugated NfL**

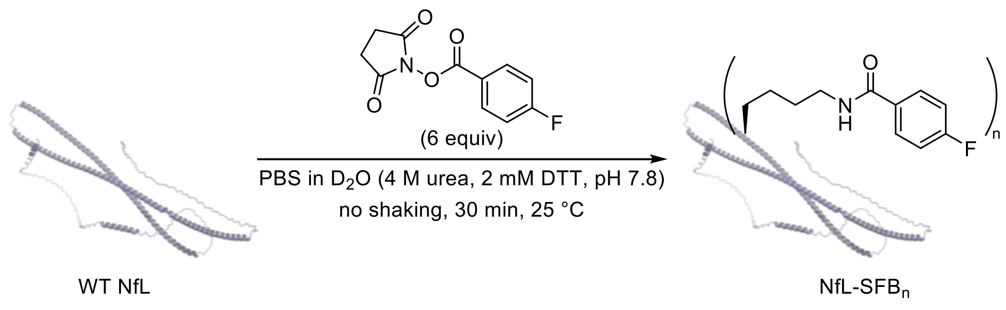


The reaction buffer (PBS in $D_2O$ with 4 M urea and 2 mM DTT) was added to a glass HPLC vial containing mNfL WT such that the final concentration was 5.6 mg $mL^{-1}$ (90 μM) in a total reaction volume of 50 μL. SFB (5.4 μL of a 5 mM stock prepared in DMSO, 6 equiv.) was added and the reaction was left at 25 ˚C for 30 min in a thermocycler (no shaking). The reaction was repeated with a further 9 batch of samples. A sample of the crude mixture was then analysed by LC-MS to determine reaction conversion. All reactions were combined and diluted with the reaction buffer to a final volume of 0.5 mL. The mixture was then loaded onto a PD MiniTrap G-25 desalting column (pre-equilibrated with PBS in $D_2O$ with 4 M urea and 2 mM DTT) and eluted with 1 mL of PBS in $D_2O$ with 4 M urea and 2 mM DTT. Protein concentration was measured using a combination of NanoPhotometer® and Bradford assay. A range of concentrations of WT mNfL were measured to establish a relationship between NanoPhotometer®-generated measurements and absorbance readings from the Bradford assay. For a range of concentrations of mNfL-SFB, absorbance measurements were taken and expected concentrations were calculated using the concentration-relationship graph of WT mNfL. The expected values for various concentrations of mNfL-SFB were averaged and compared with NanoPhotometer® readings to derive a correction value to be applied for future measurements based on NanoPhotometer®. The protein was then concentrated with an Amicon 10 kDa MWCO centrifugal filter (pre-washed with $D_2O$) for subsequent NMR studies.

Under the conditions shown in the reaction scheme, the calculated average number of conjugated SFB was ~1.2. The corresponding ion series is shown below.

Conditions: 90 μM NfL WT, pH 7.8, 6 equiv. SFB, 25 ˚C, 30 min, no shaking.

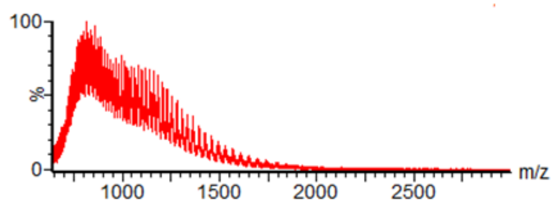


Under similar conditions with slightly higher starting protein concentrations and shaking (400 rpm), the corresponding ion series showed an overall shift to higher m/z values and could not be deconvoluted. This suggests possible aggregation event, even with reduced SFB loadings.

Conditions: 107 μM NfL WT, 5 equiv. SFB, 25 ˚C, 15 min, 400 rpm.

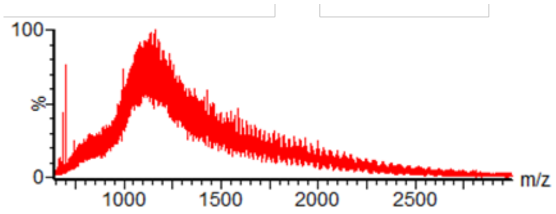


Conditions: 100 μM NfL WT, pH 7.8, 3 equiv. SFB, 25 ˚C, 30 min, 400 rpm.

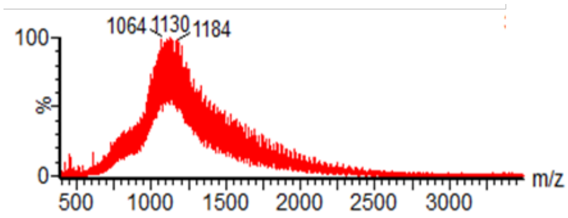

**NfL sample preparation for NMR studies**

Purified and concentrated mNfL samples (WT, SFB- and Fha-modified) stored in buffered $D_2O$ (PBS containing 137 mM NaCl, 2.7 mM KCl, 8.1 mM $Na_2HPO_4$ and 1.5 mM $KH_2PO_4$, with 4 M urea and 2 mM DTT, pH 7.8) were taken for ultracentrifugation (0 to 100k g in 10 min then 100k g for 20 min at 25 °C). The supernatant was then immediately separated and diluted accordingly with the same storage buffer to reach a target protein concentration of 206 µM (~12.9 ± 0.2 mg $mL^{-1}$) ± 2% across all NMR samples before initiating aggregating. Concentrations were measured using a NanoPhotometer®.

**Initiating NfL aggregation**

NfL variants (43.6 µL at 12.9 ± 0.2 mg $mL^{-1}$) were incubated at 25 °C for at least 10 min before diluting with 152.0 µL of PBS in $D_2O$ (containing 169 mM NaCl) to obtain a final protein concentration of 46 µM (~2.9 mg $mL^{-1}$) in PBS (with 0.89 mM urea, 0.44 M DTT, 2.7 mM KCl, 8.1 mM $Na_2HPO_4$ and 1.5 mM $KH_2PO_4$ and a final NaCl concentration of 162 mM in $D_2O$). The diluted mNfL samples were mixed by pipetting (40 cycles) and then immediately transferred (using a 1 mL syringe) into the NMR tube.

**NMR set-up and analysis**

NMR experiments to monitor NfL aggregation were carried out at 298 K using a 600 MHz NEO NMR Spectrometer equipped with a BBO Cryoprobe (CP2.1 BBO 600S3 BB-H&F-D-05-Z XT).

Aggregation was monitored using the PEW5 perfect echo Watergate sequence (available from the Manchester NMR Methodology Group website), with the following parameters: 64 scans, 4 dummy scans, a fixed receiver gain value of 101, an acquisition time of 1.7 seconds, and a relaxation delay (D1) of 1 second. This setup resulted in an experiment time of 3 minutes and 11 seconds per spectrum.
Samples were "multiplexed," meaning that the sample changer alternated between samples, effectively enabling the simultaneous recording of two overnight time courses. However, this approach does introduce delays between consecutive data points, as

loading, locking, tuning, matching, shimming, and other preparations took approximately 5 minutes per sample.

To achieve accurate multiplexing, precise timing is critical. The mixing of the second sample was timed to coincide precisely with the end of the recording of the first NMR spectrum of the first sample. A temperature-controlled sample changer is obviously essential. We used a SampleJet 99/5 with a Pre-Heating Unit. Notably, with this equipment, samples spent more time in the Pre-Heating Unit—located between the magnet bore and the carousel—than in the carousel racks. Consequently, setting the temperatures of both the Pre-Heating Unit and the carousel to 298 K was vital, and this temperature was equilibrated at least 30 minutes prior to the experiments.

To configure Bruker's ICON-NMR automation for alternating sample measurements, the system was set to execute experiments strictly in submission order. Experiments were submitted on alternate samples via spreadsheet import, which allowed rapid and accurate input of over a hundred individual experiments.

Data was analysed using the Mestrenova software package. A 5 Hz exponential window function was applied, followed by baseline correction using a first-order polynomial fit. Signal regions selected for analysis were chosen to avoid buffer resonances and any additional resonances introduced by the SFB and Fha labels. The integral within the spectral region 0.95 to 0.70ppm was quantified and its variation with time was reported. The error associated with Fha-labelled NfL (n = 1) was estimated by averaging the standard deviation obtained for wild-type (n = 2) and SFB-conjugated NfL (n = 2).

#### 4. Small molecule NMR spectra

$^1H$, $^{19}F$ and $^{13}C$ NMR Spectra of 2-((difluoromethyl)thio)pyridine in $CDCl_3$

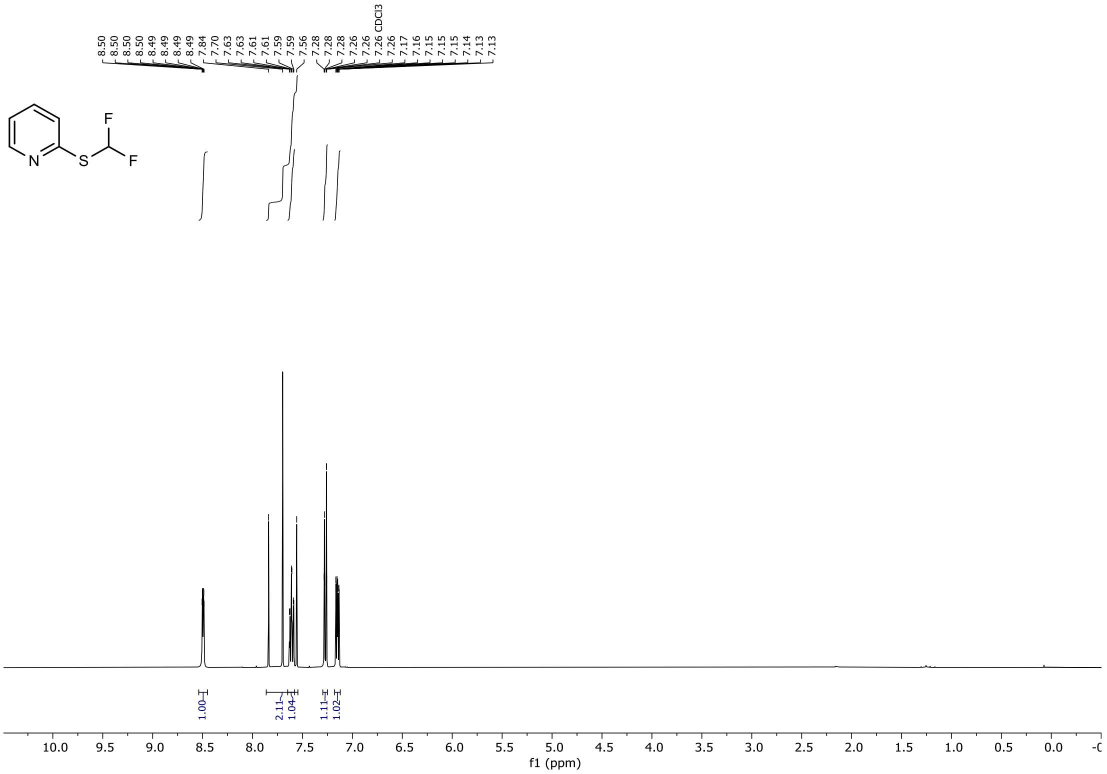

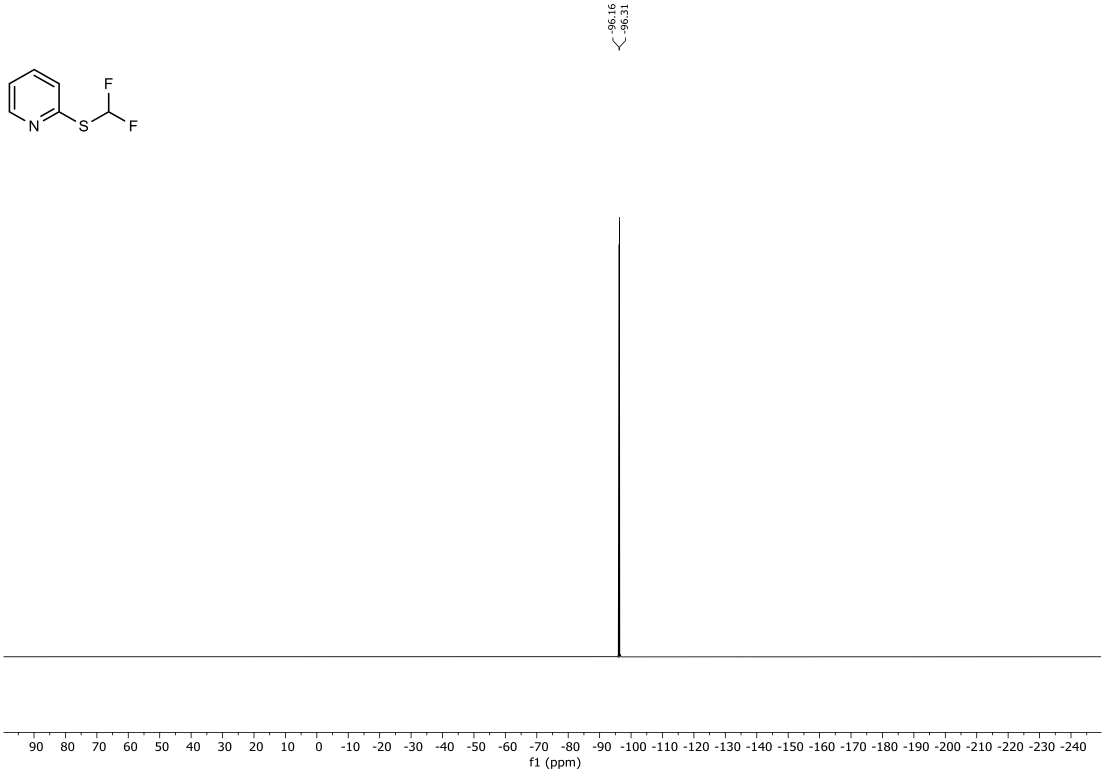

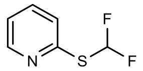

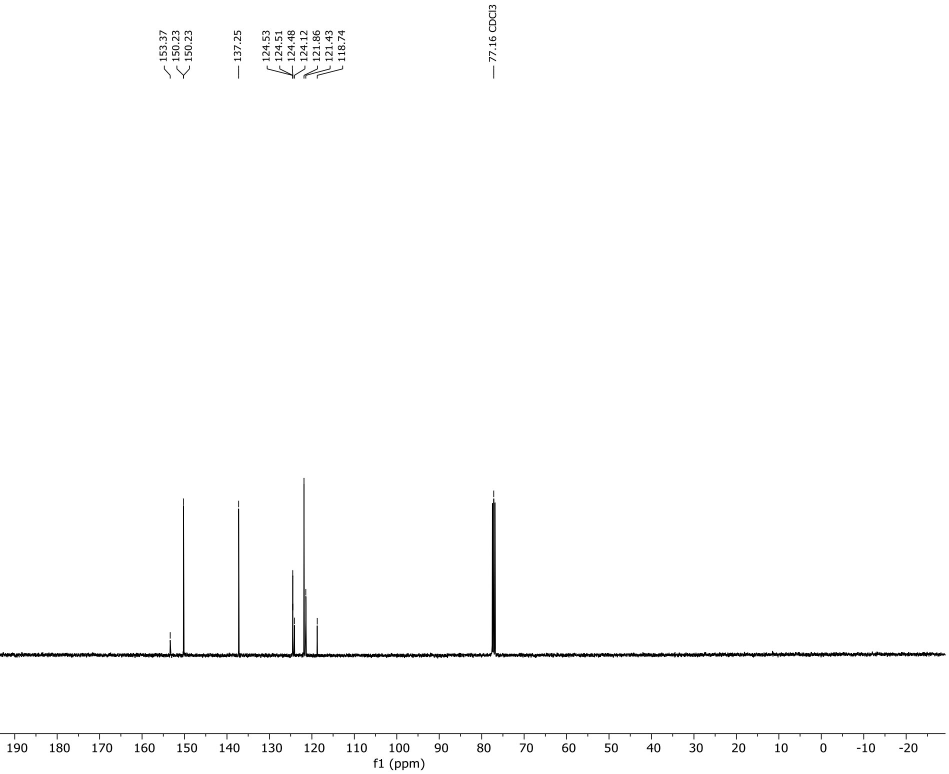

153.37
150.23
150.23
137.25
124.53
124.51
124.48
124.12
121.86
121.43
118.74
77.16 CDCl3
220 210 200 190 180 170 160 150 140 130 120 110 100 90 80 70 60 50 40 30 20 10 0 -10 -20
f1 (ppm)

$^{1}H$, $^{19}F$ and $^{13}C$ NMR Spectra of 2-((difluoromethyl)thio)pyrimidine in $CDCl_3$

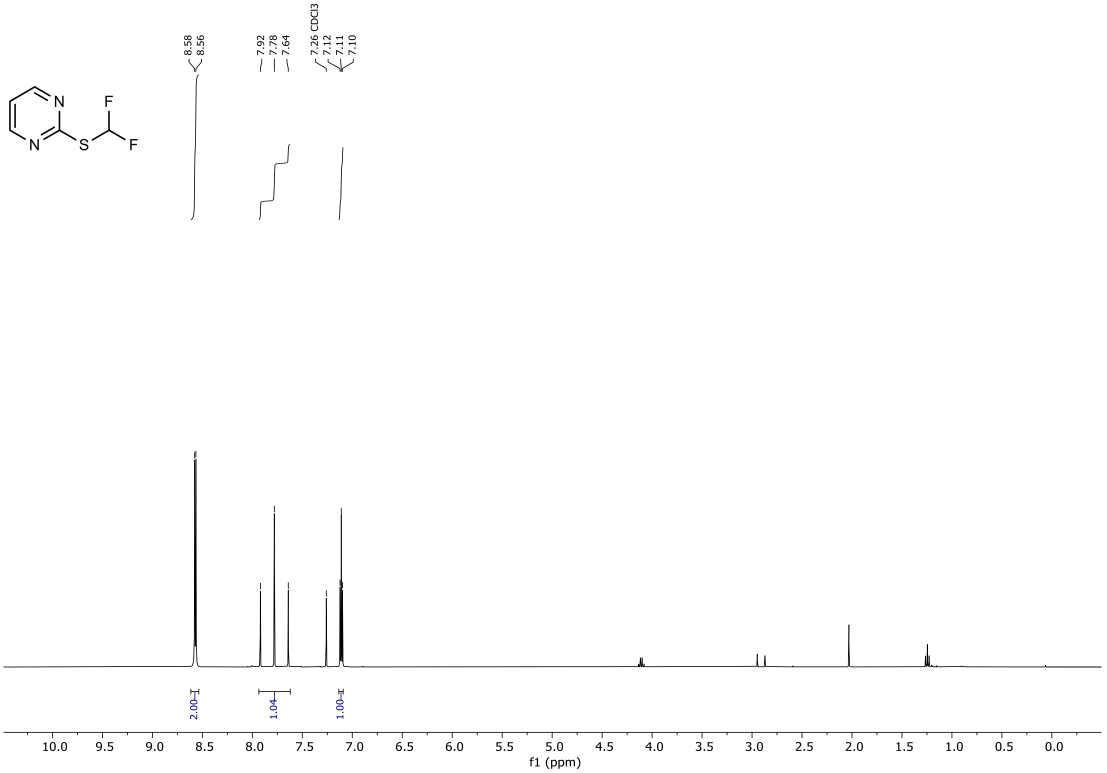


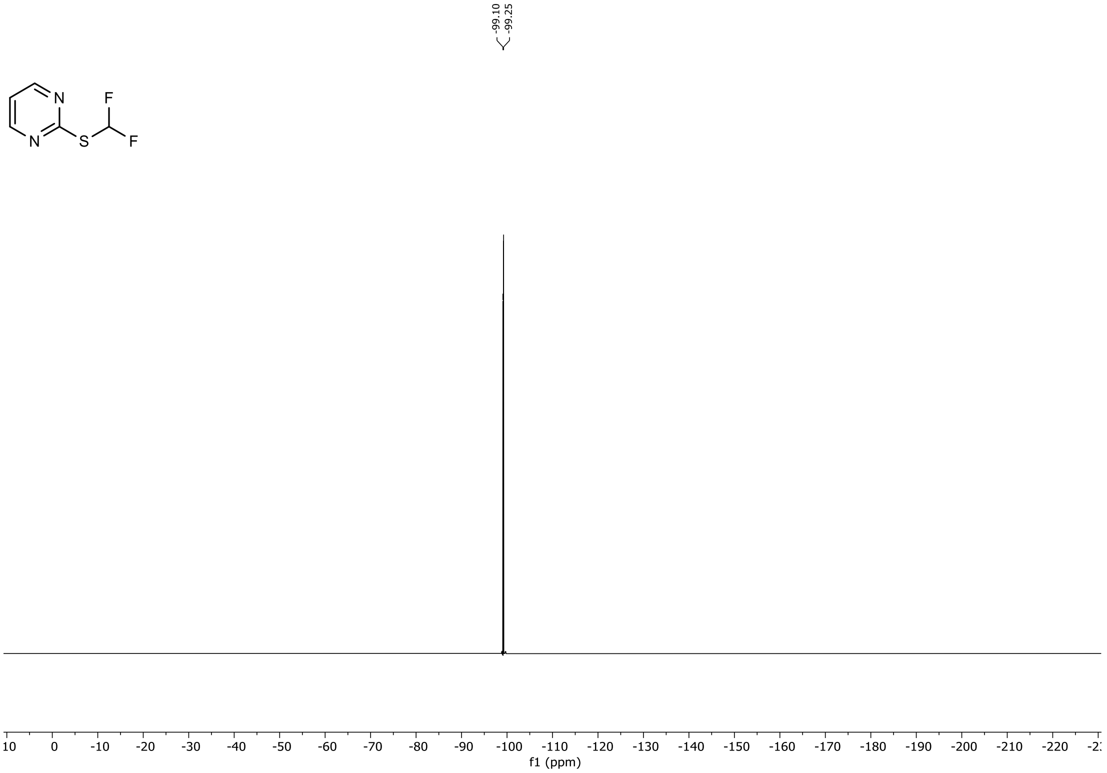

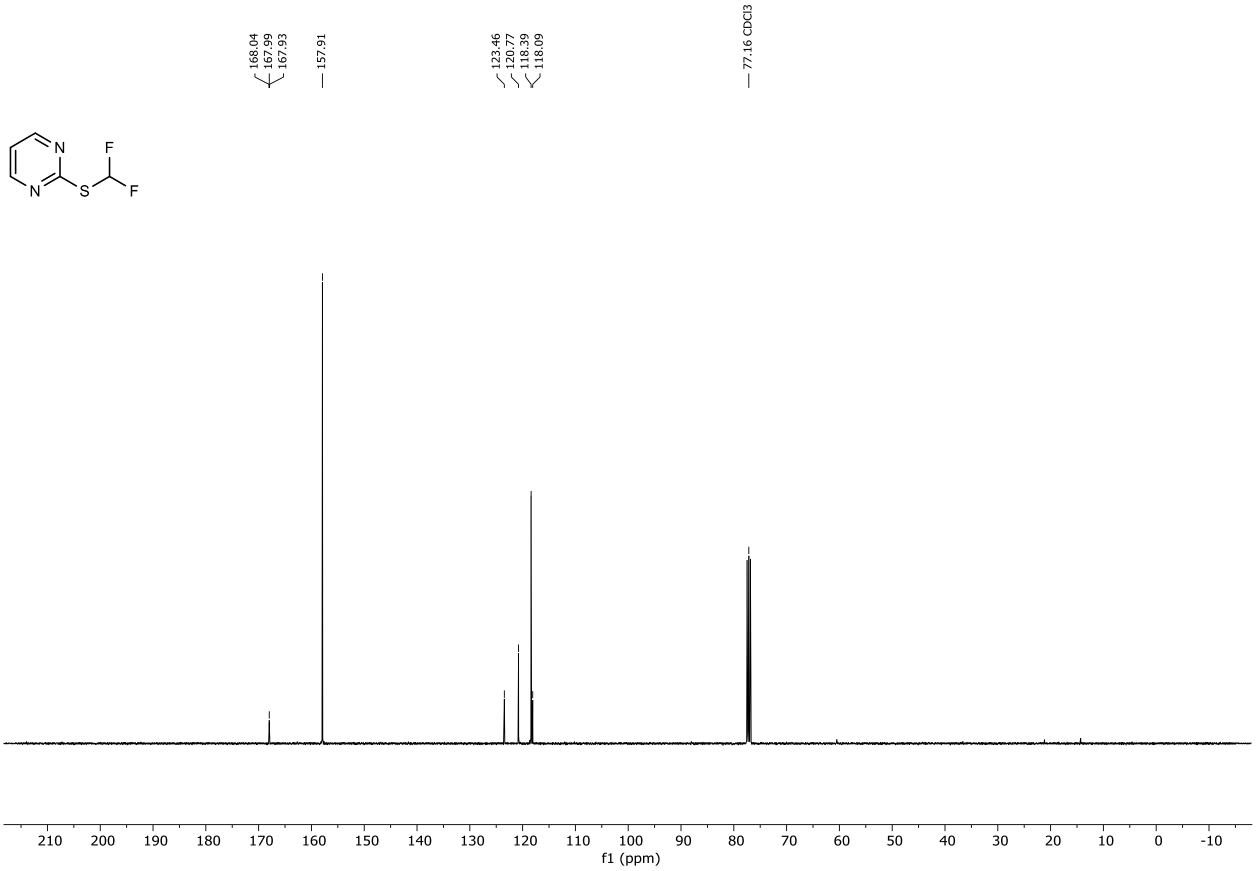

168.04
167.99
167.93
157.91
123.46
120.77
118.39
118.09
77.16 CDCl3
N
N
S
F
F
210
200
190
180
170
160
150
140
130
120
110
100
90
80
70
60
50
40
30
20
10
0
-10
f1 (ppm)

$^{1}H$, $^{19}F$ and $^{13}C$ NMR Spectra of 2-((difluoromethyl)thio)benzothiazole purified from N-difluoromethylated by-product in $CDCl_3$

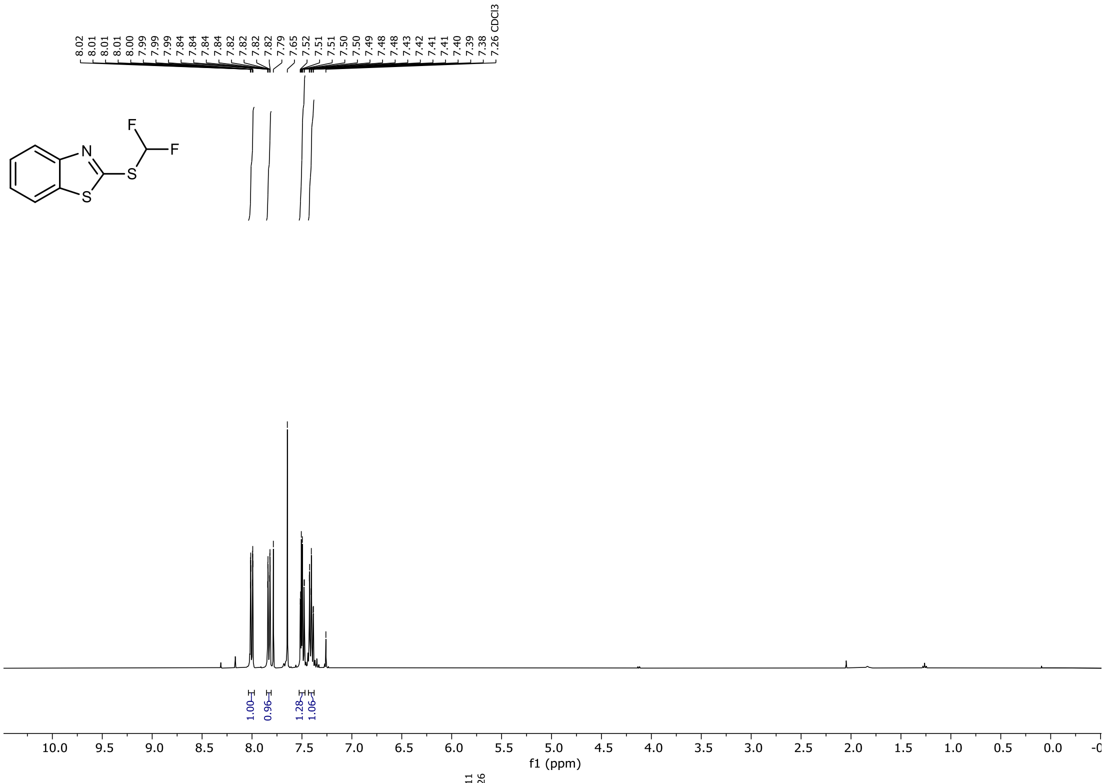

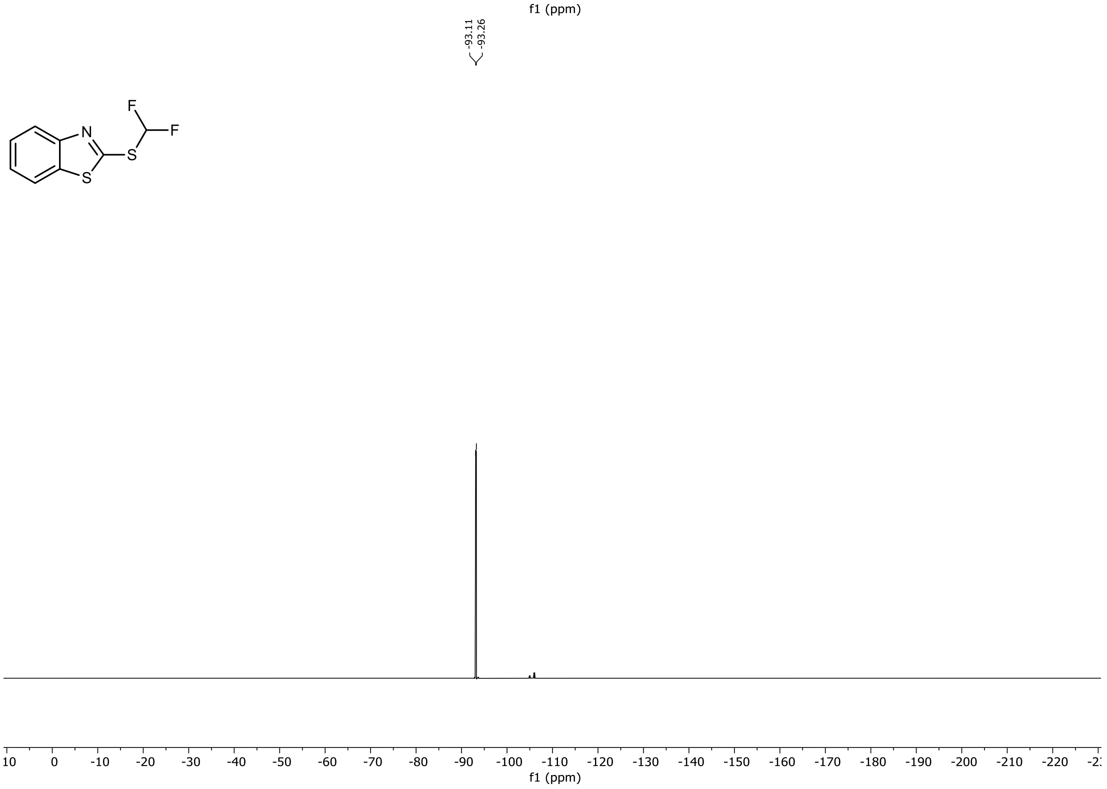

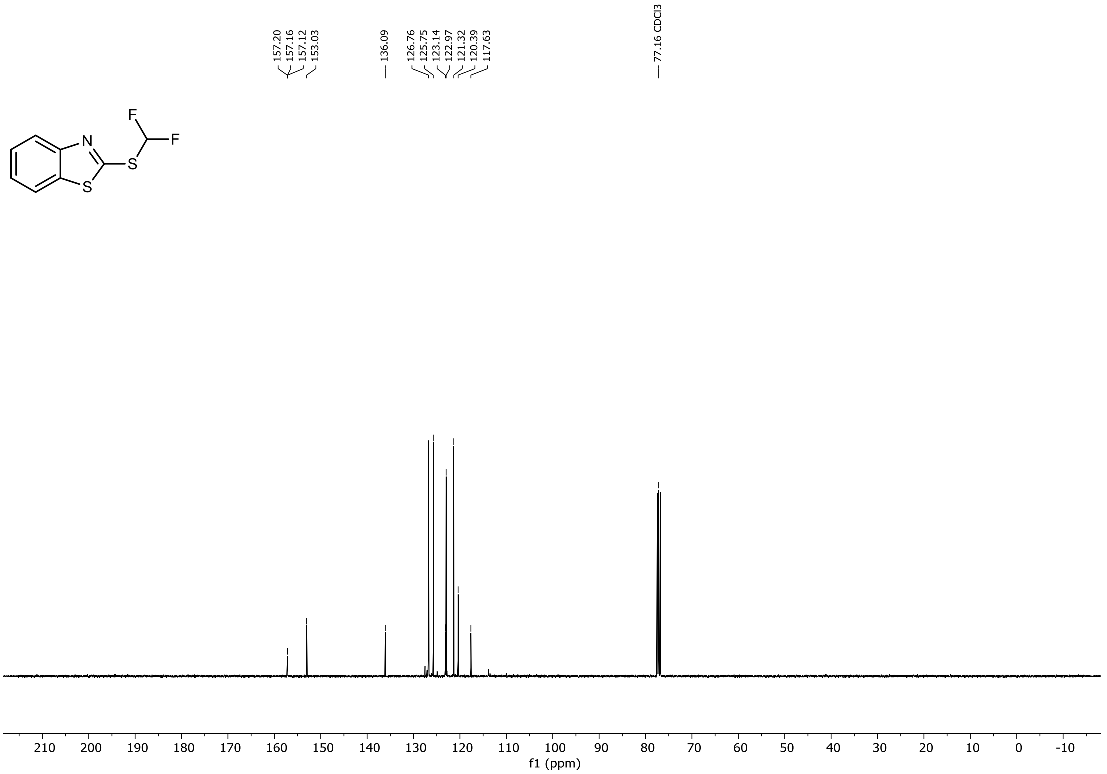

157.20
157.16
157.12
153.03
136.09
126.76
125.75
123.14
122.97
121.32
120.39
117.63
77.16 CDCl3
210 200 190 180 170 160 150 140 130 120 110 100 90 80 70 60 50 40 30 20 10 0 -10
f1 (ppm)

$^{1}H$, $^{19}F$ and $^{13}C$ NMR Spectra of 2-((fluoromethyl)thio)benzothiazole in $CDCl_3$

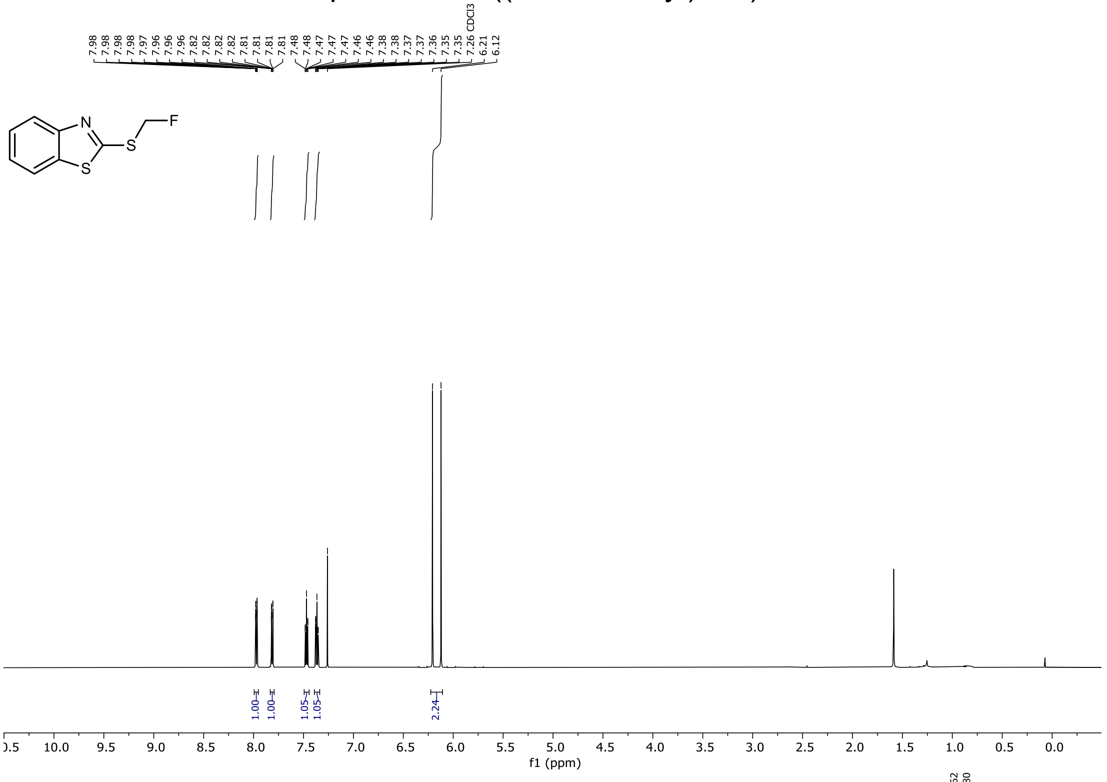


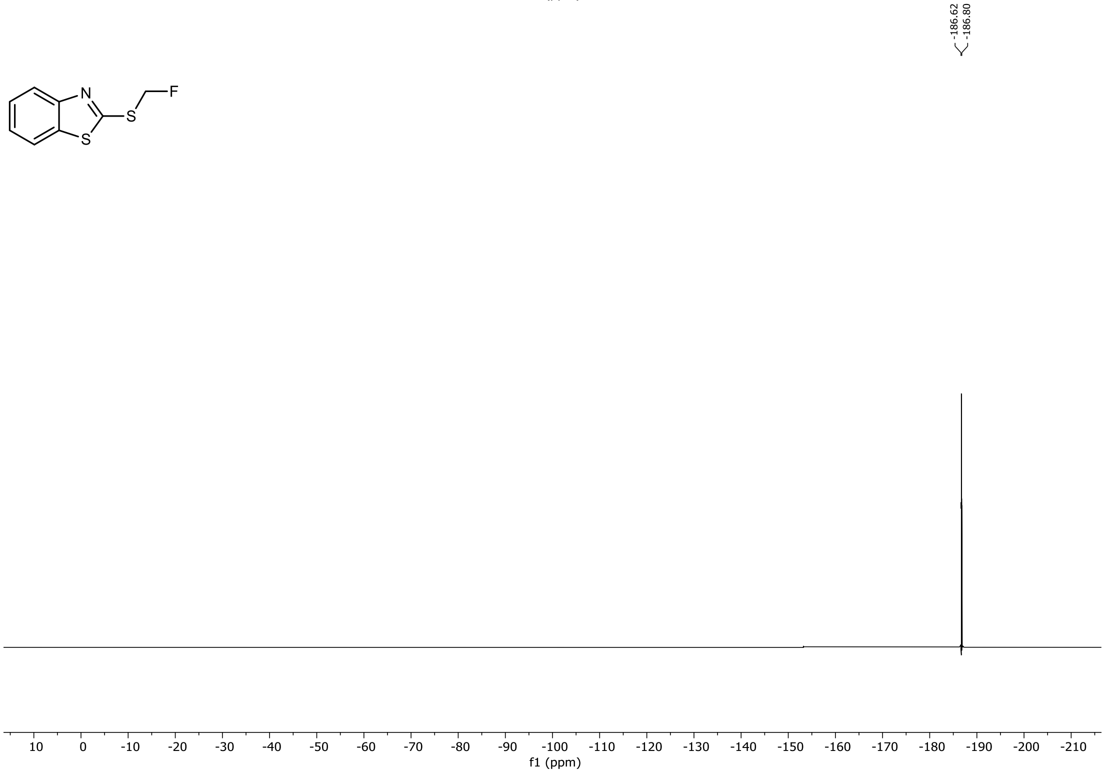

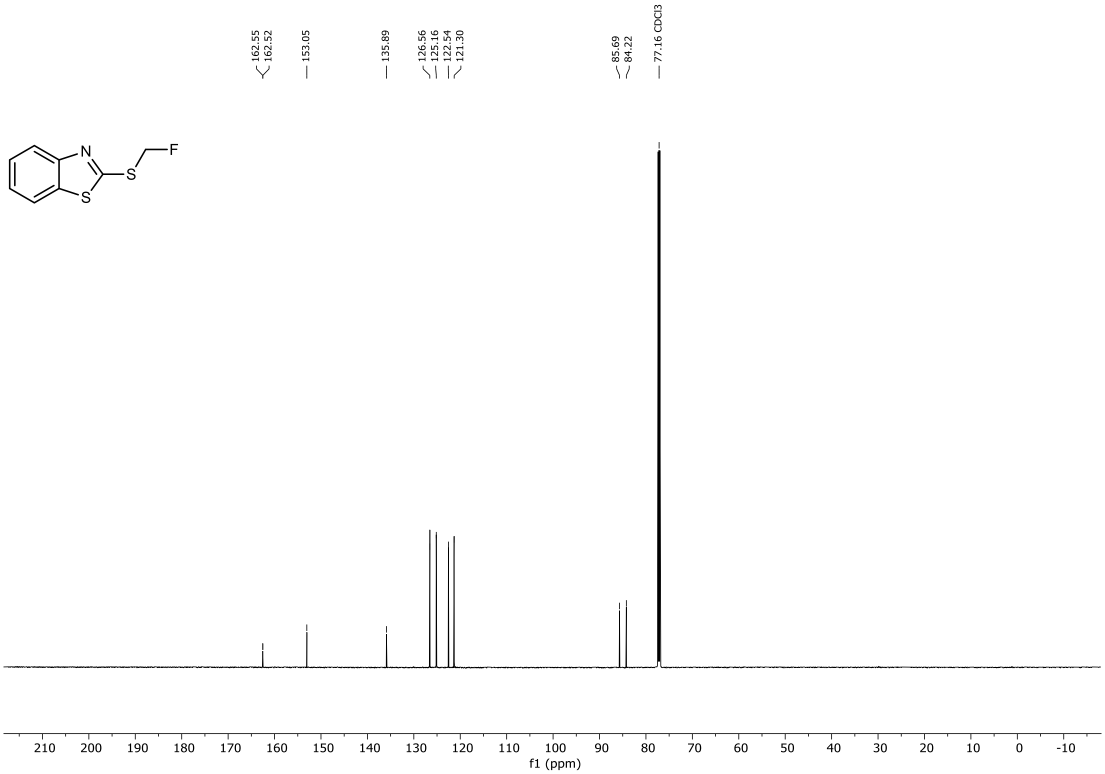
162.55
162.52
153.05
135.89
126.56
125.16
122.54
121.30
85.69
84.22
77.16 CDCl3
N
S
S
F
210 200 190 180 170 160 150 140 130 120 110 100 90 80 70 60 50 40 30 20 10 0 -10
f1 (ppm)

[1]H, [19]F and [13]C NMR Spectra of 2-((bromofluoromethyl)thio)pyridine in $CDCl_3$

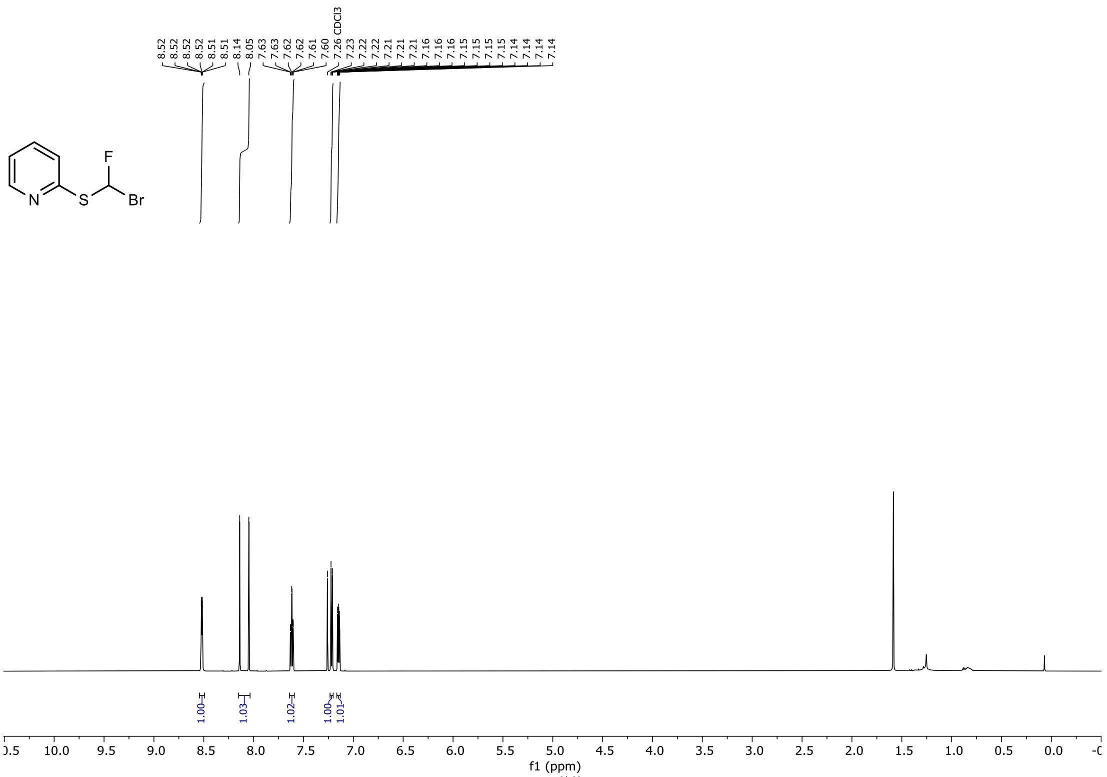


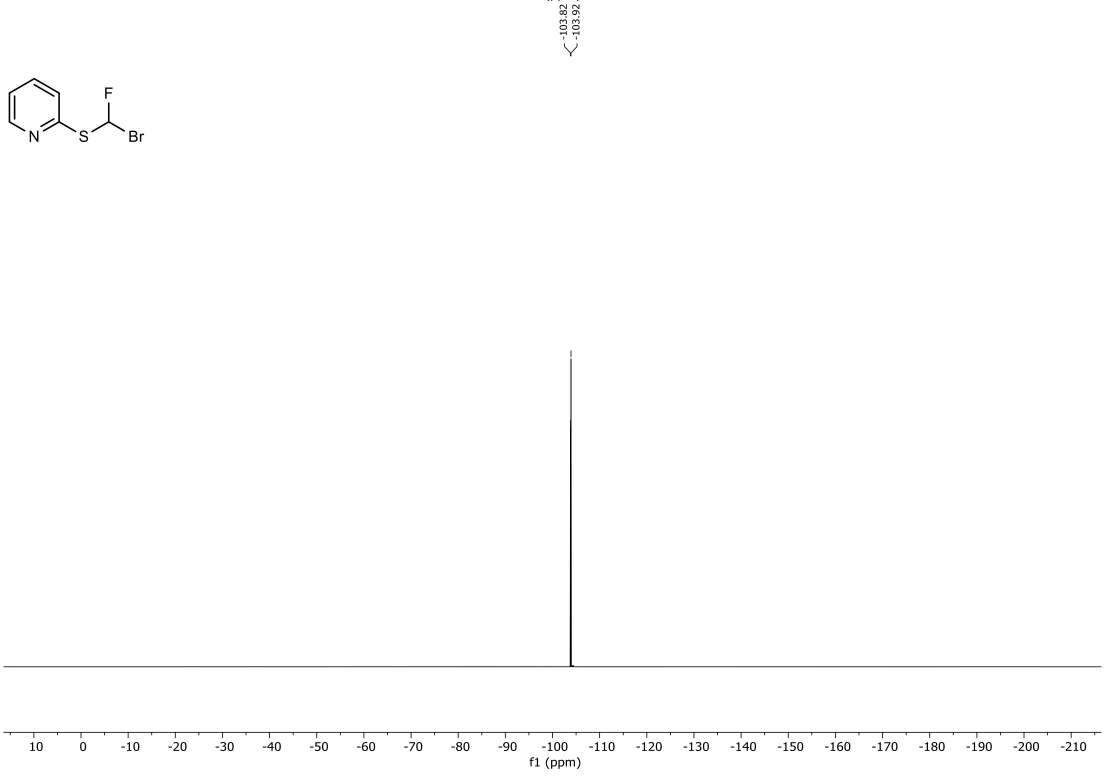

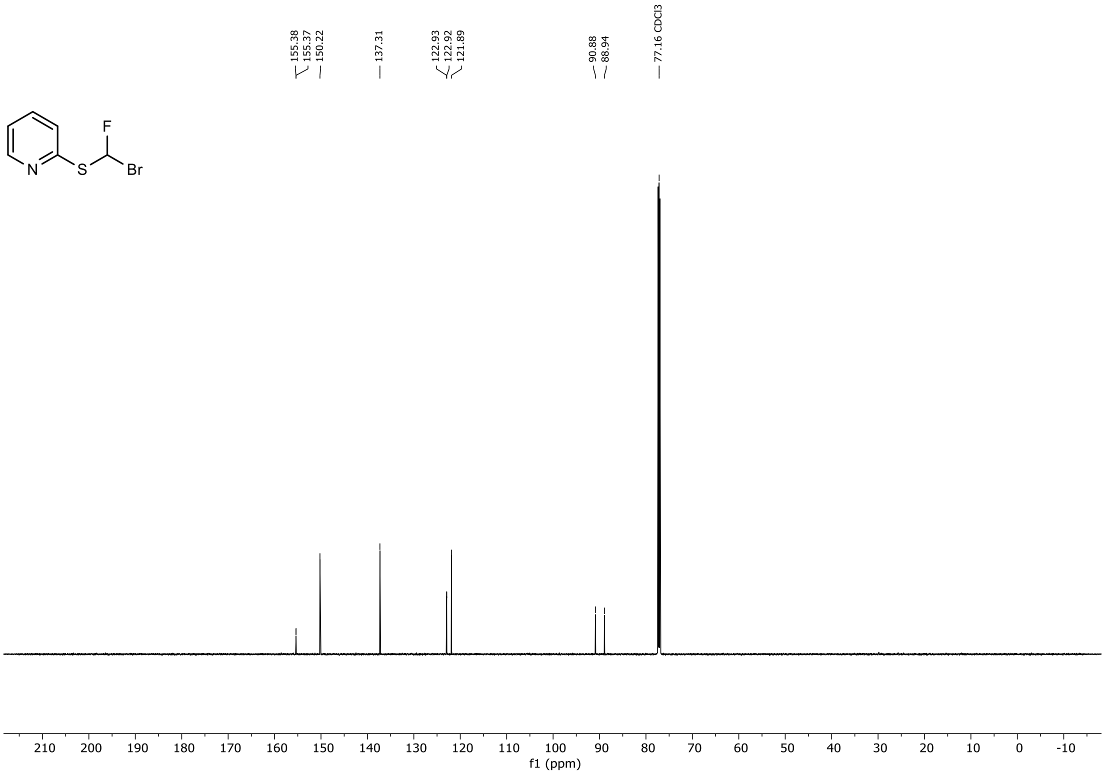
155.38
155.37
150.22
137.31
122.93
122.92
121.89
90.88
88.94
77.16 CDCl3
F
N
S
Br
210 200 190 180 170 160 150 140 130 120 110 100 90 80 70 60 50 40 30 20 10 0 -10
f1 (ppm)

$^{1}H$, $^{19}F$ and $^{13}C$ NMR Spectra of 2-((bromofluoromethyl)thio)pyrimidine in $CDCl_3$

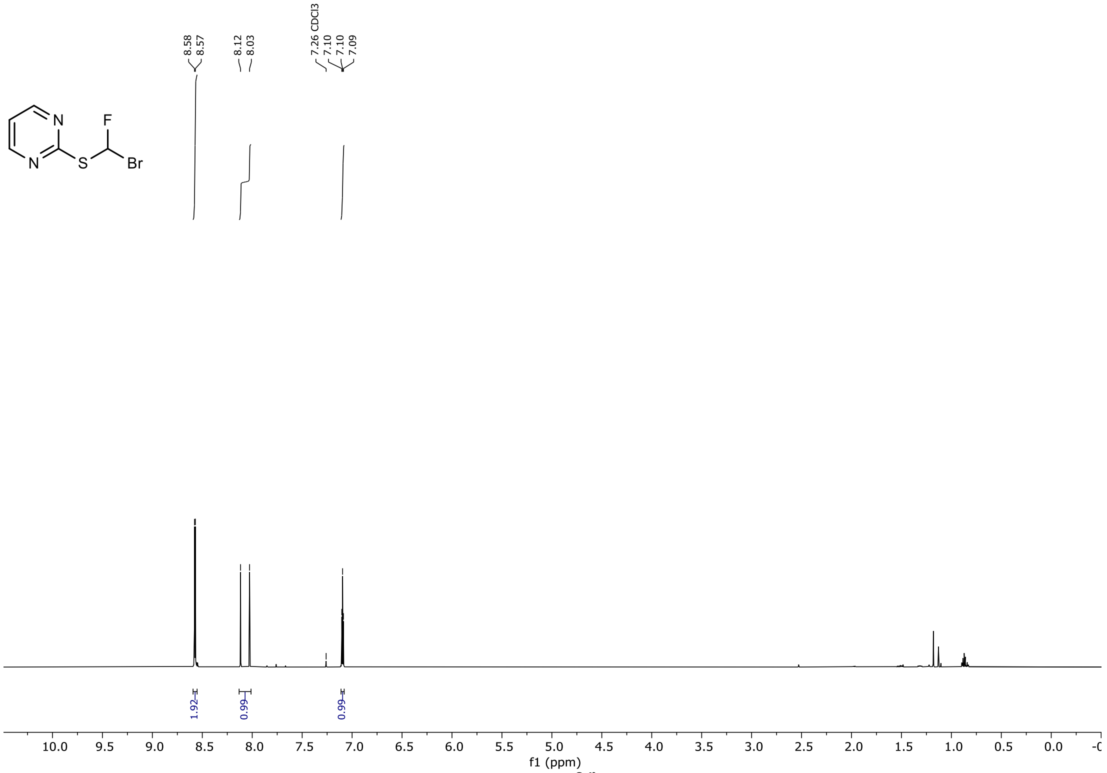


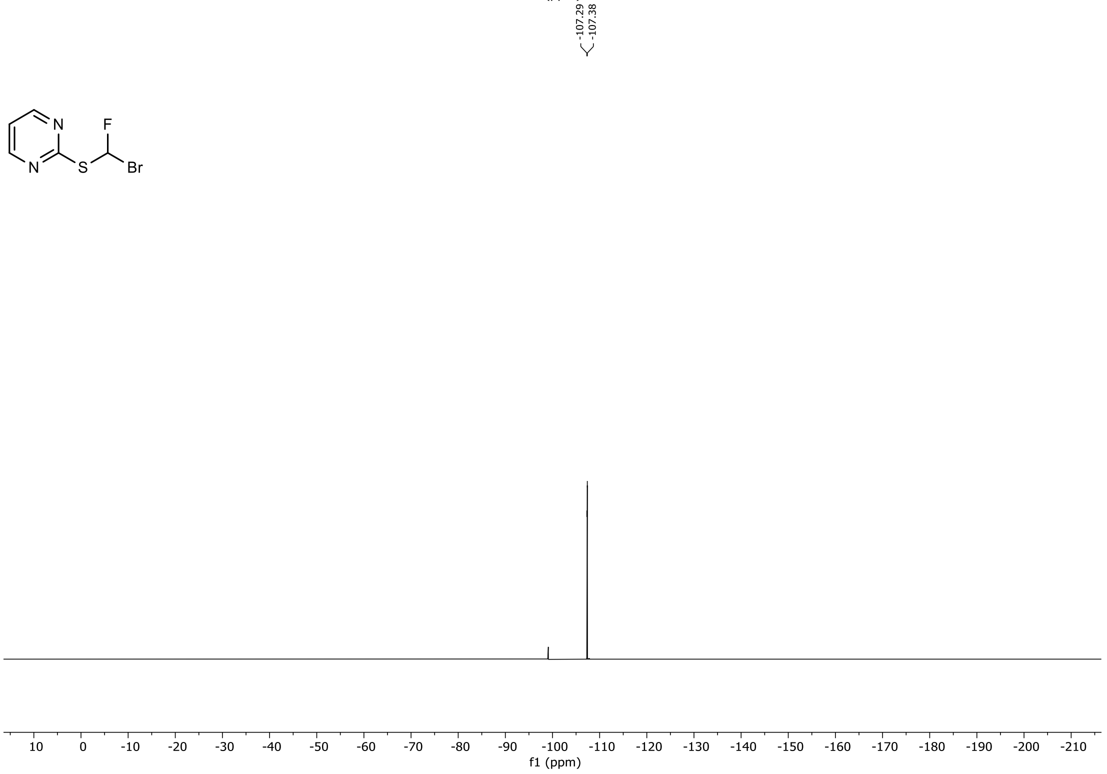

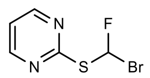

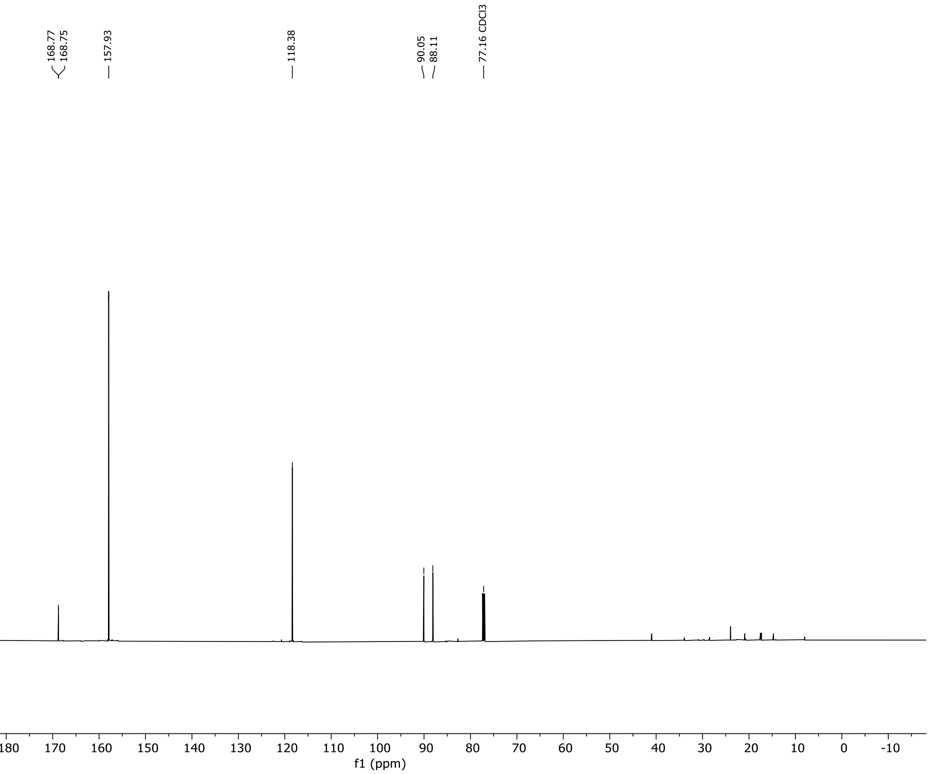
168.77
168.75
157.93
118.38
90.05
88.11
77.16 CDCl3
210 200 190 180 170 160 150 140 130 120 110 100 90 80 70 60 50 40 30 20 10 0 -10
f1 (ppm)

$^{1}H$, $^{19}F$ and $^{13}C$ NMR Spectra of 2-((bromofluoromethyl)thio)benzothiazole in $CDCl_3$

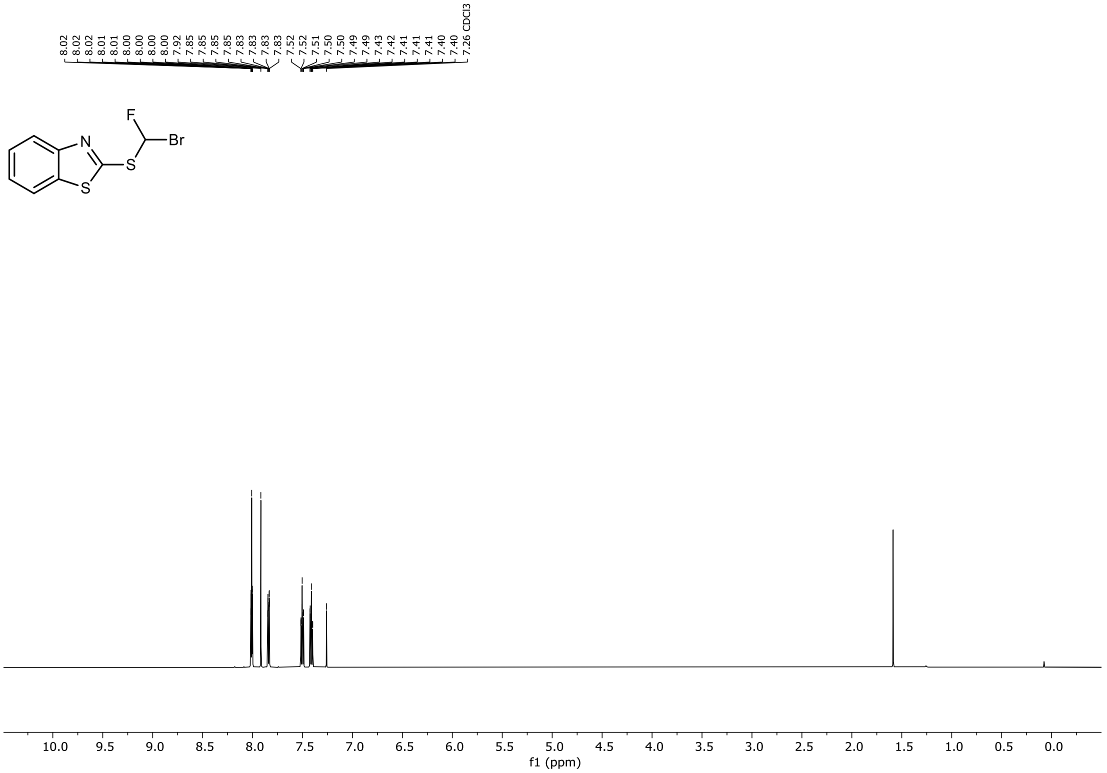

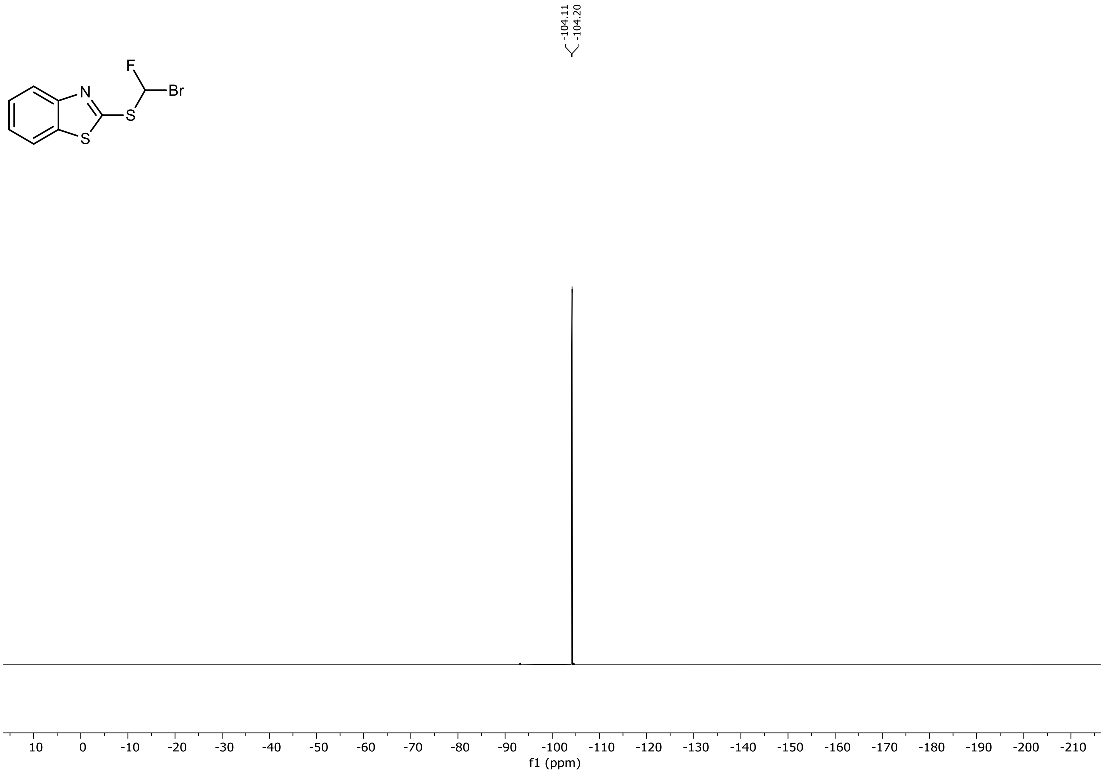

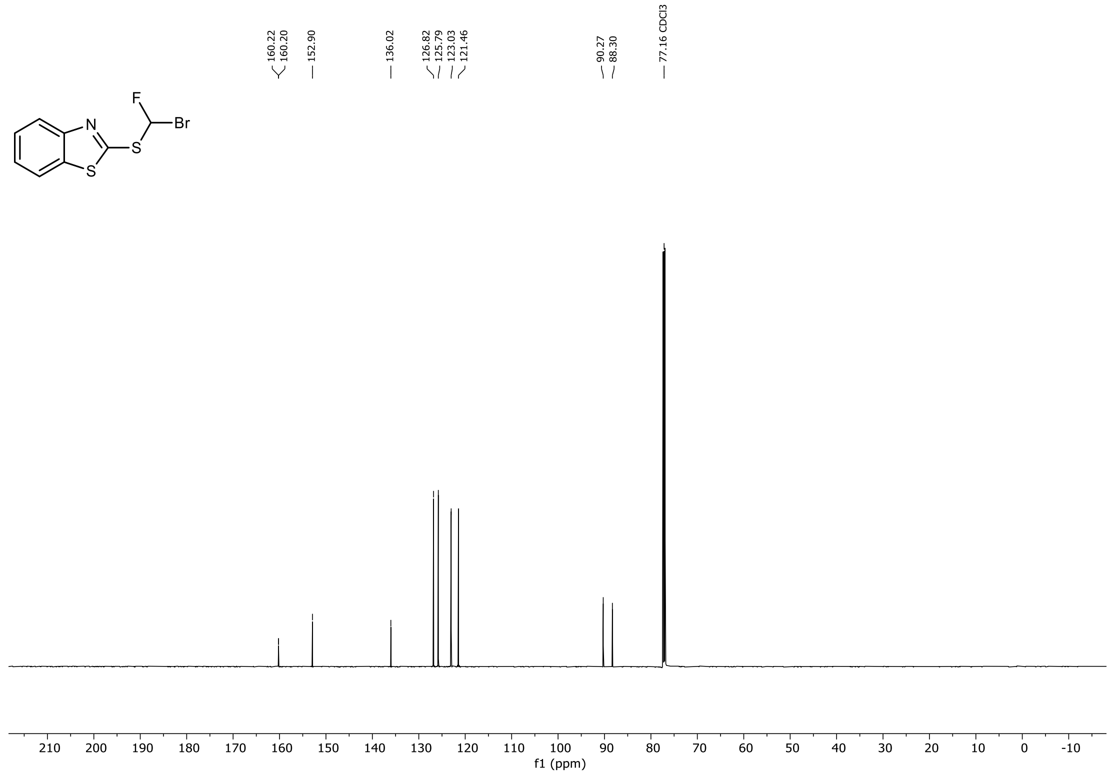

160.22
160.20
152.90
136.02
126.82
125.79
123.03
121.46
90.27
88.30
77.16 CDCl3
F
Br
N
S
S
210 200 190 180 170 160 150 140 130 120 110 100 90 80 70 60 50 40 30 20 10 0 -10
f1 (ppm)

### [1]H and [13]C NMR Spectra of 2-((bromomethyl)thio)benzothiazole in $CDCl_3$

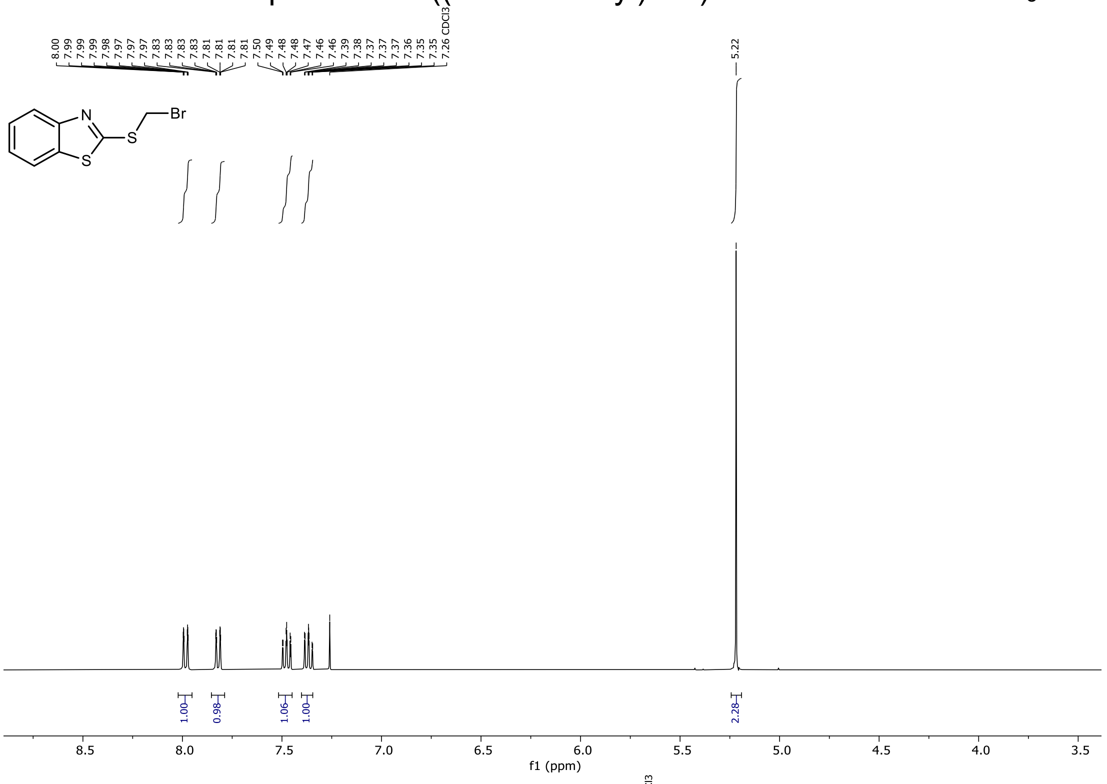


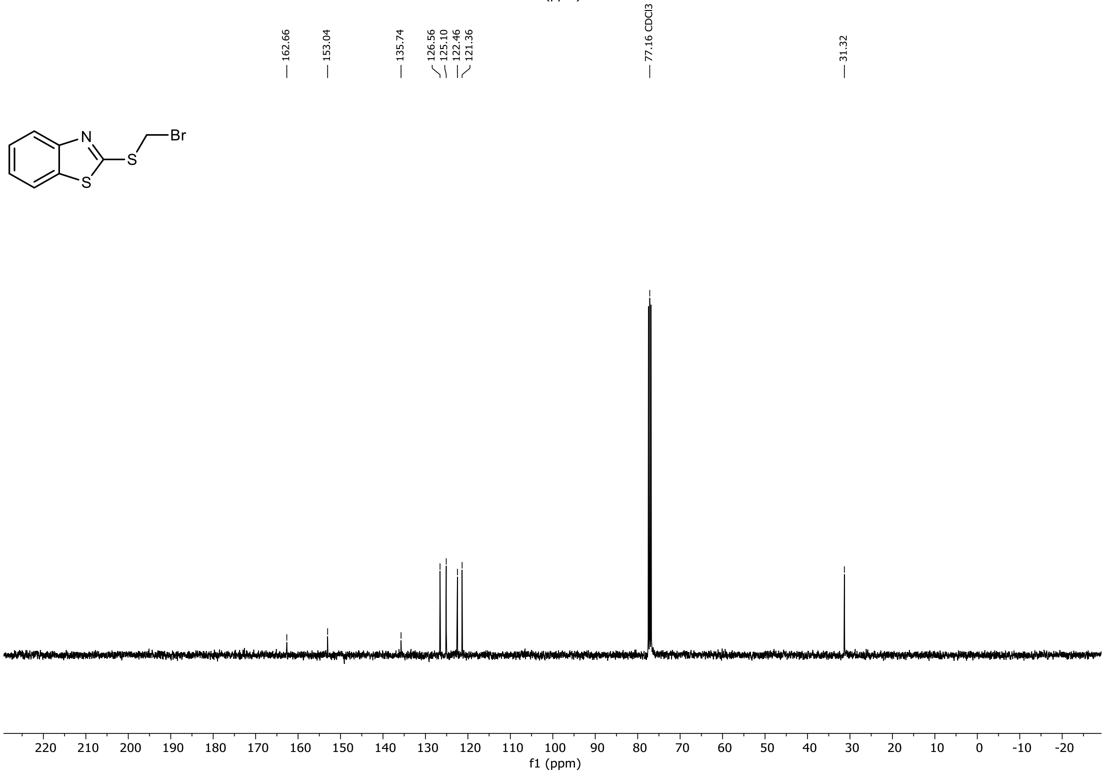

$^{1}H$, $^{19}F$ and $^{13}C$ NMR Spectra of 2-((difluoromethyl)sulfonyl)pyrimidine in $CDCl_3$

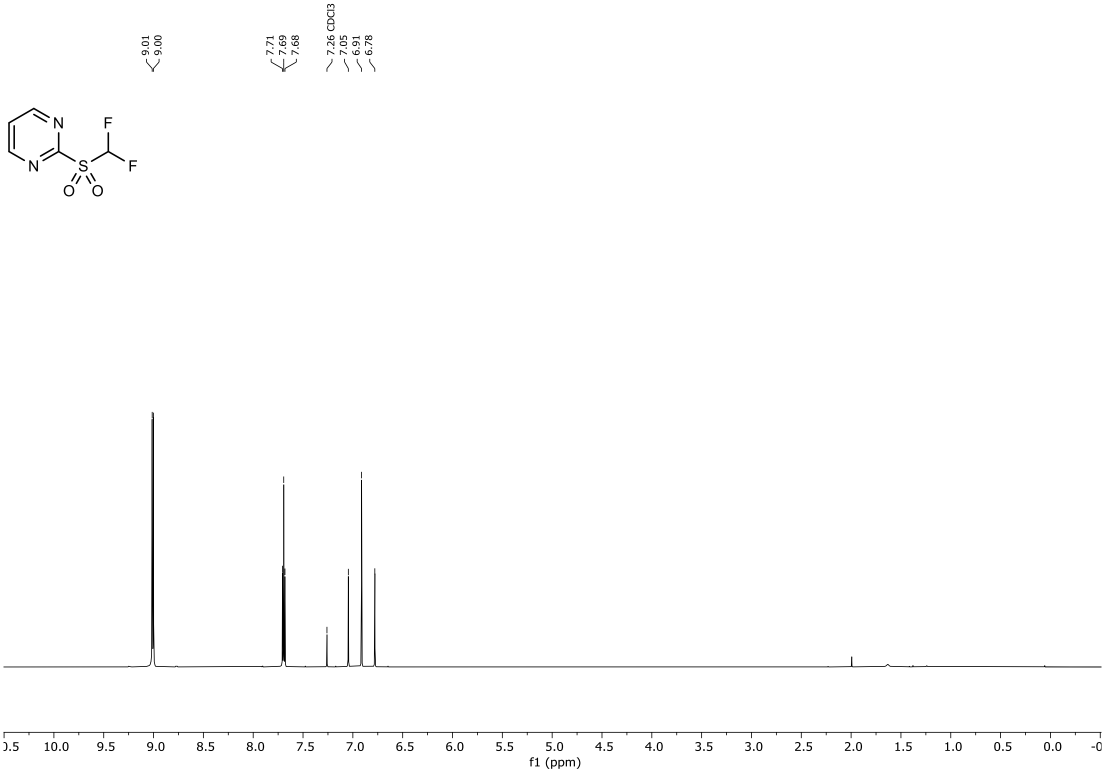


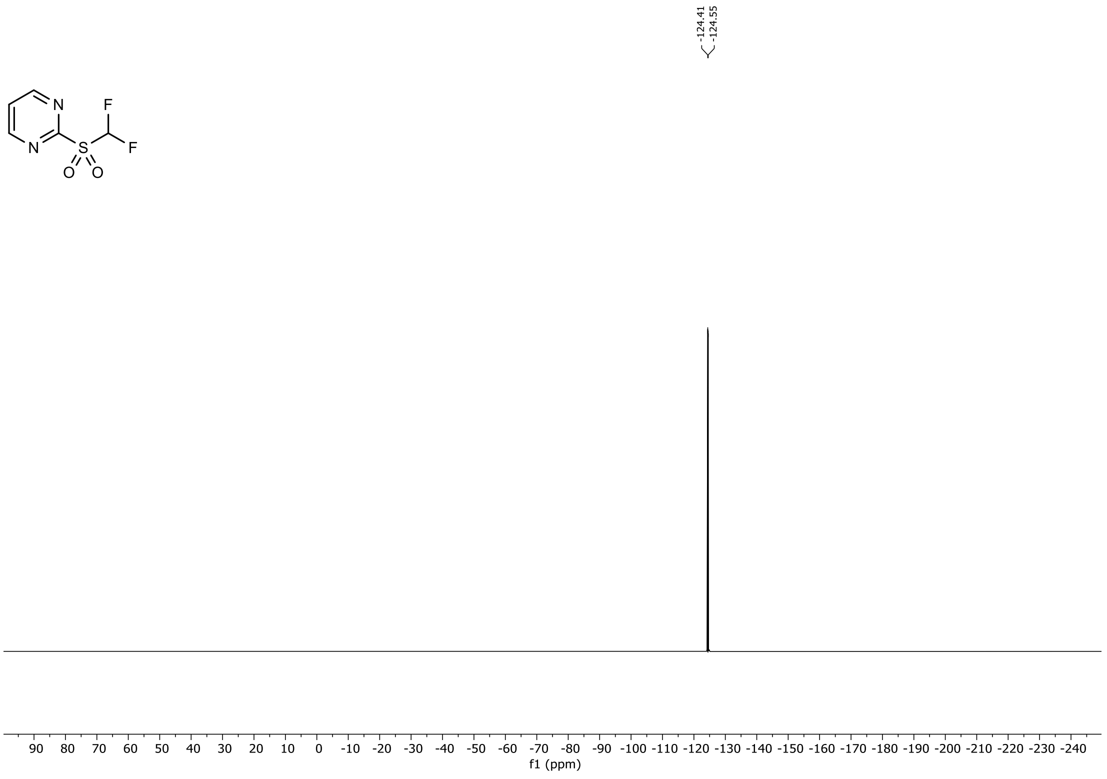

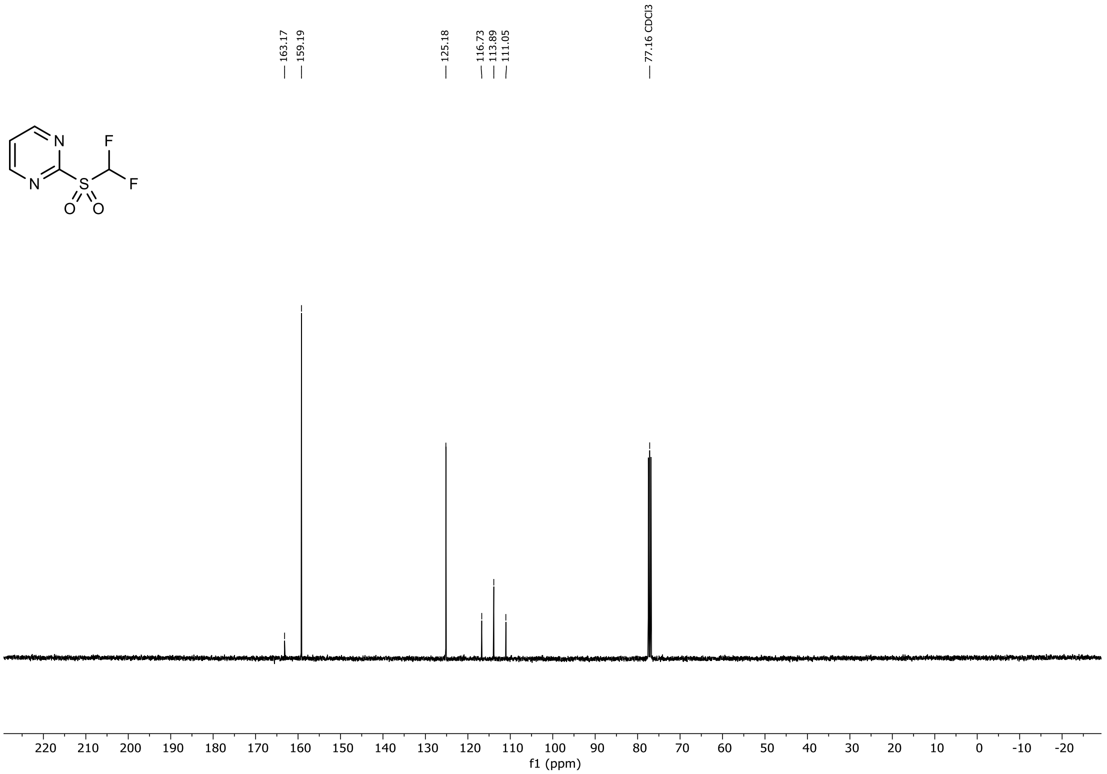
163.17
159.19
125.18
116.73
113.89
111.05
77.16 CDCl3
220 210 200 190 180 170 160 150 140 130 120 110 100 90 80 70 60 50 40 30 20 10 0 -10 -20
f1 (ppm)

$^{1}H$, $^{19}F$ and $^{13}C$ NMR Spectra of 2-((difluoromethyl)sulfonyl)benzothiazole in DMSO-$d_6$

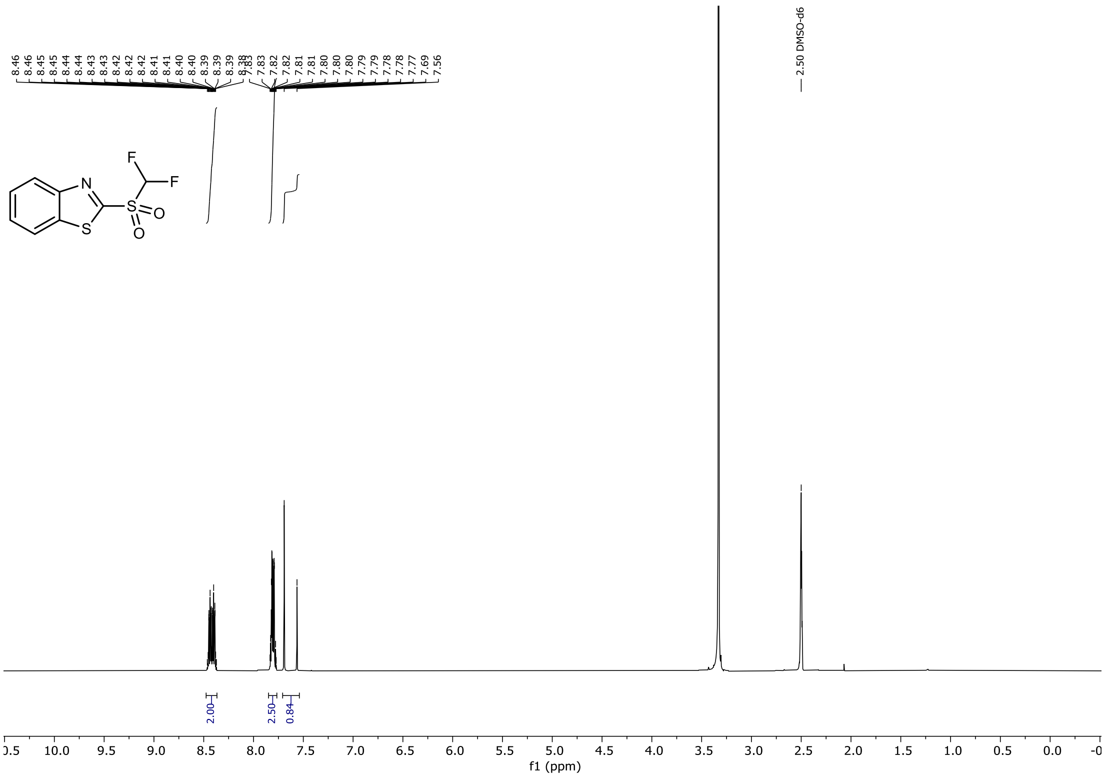


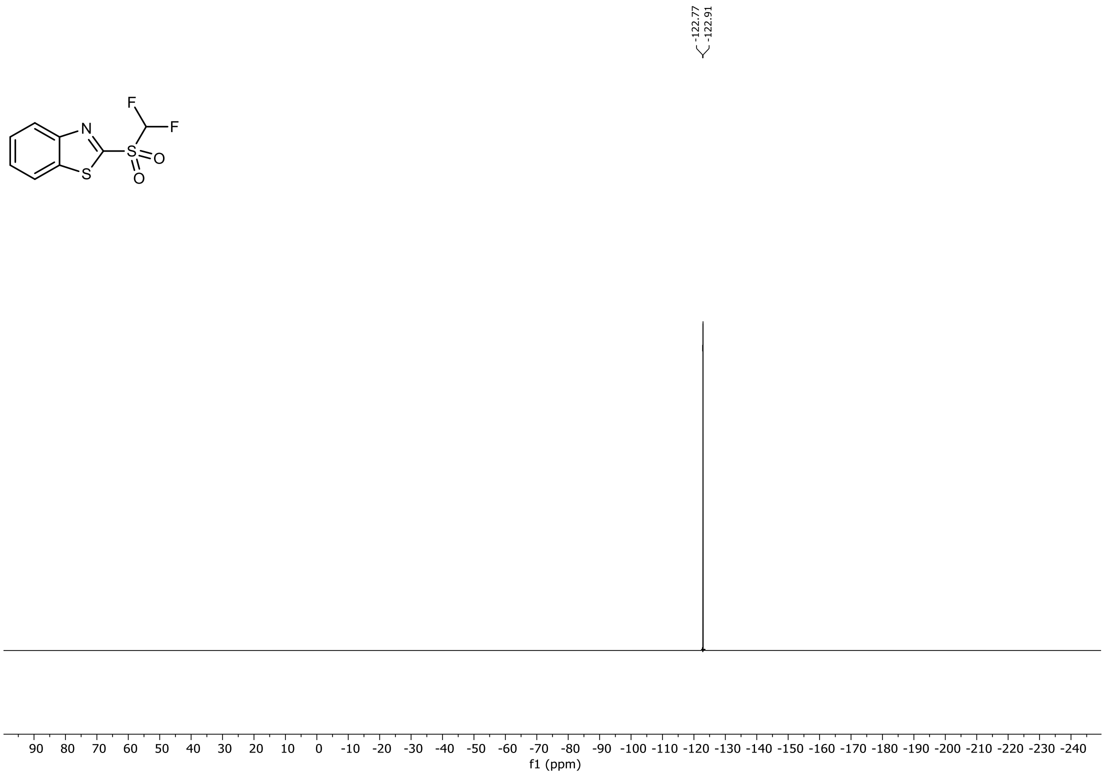

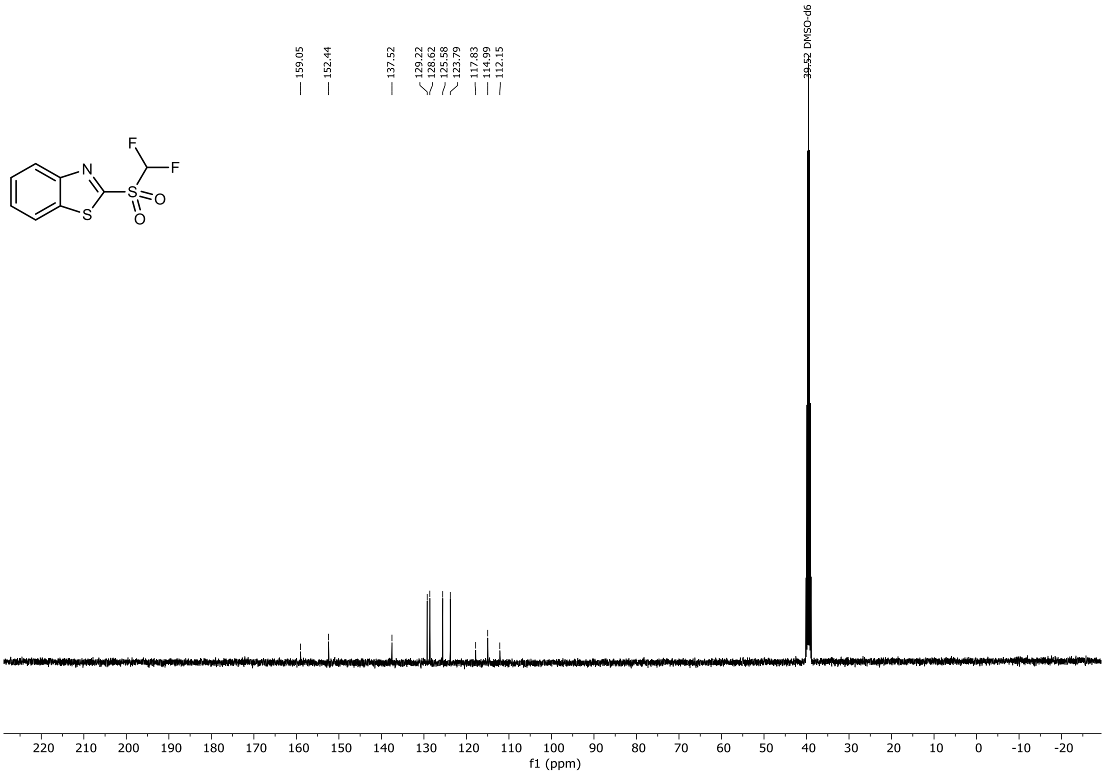
159.05
152.44
137.52
129.22
128.62
125.58
123.79
117.83
114.99
112.15
39.52 DMSO-d6
F
F
N
S
S
O
O
220 210 200 190 180 170 160 150 140 130 120 110 100 90 80 70 60 50 40 30 20 10 0 -10 -20
f1 (ppm)

$^{1}H$, $^{19}F$ and $^{13}C$ NMR Spectra of 2-((bromofluoromethyl)sulfonyl)pyridine in $CDCl_3$

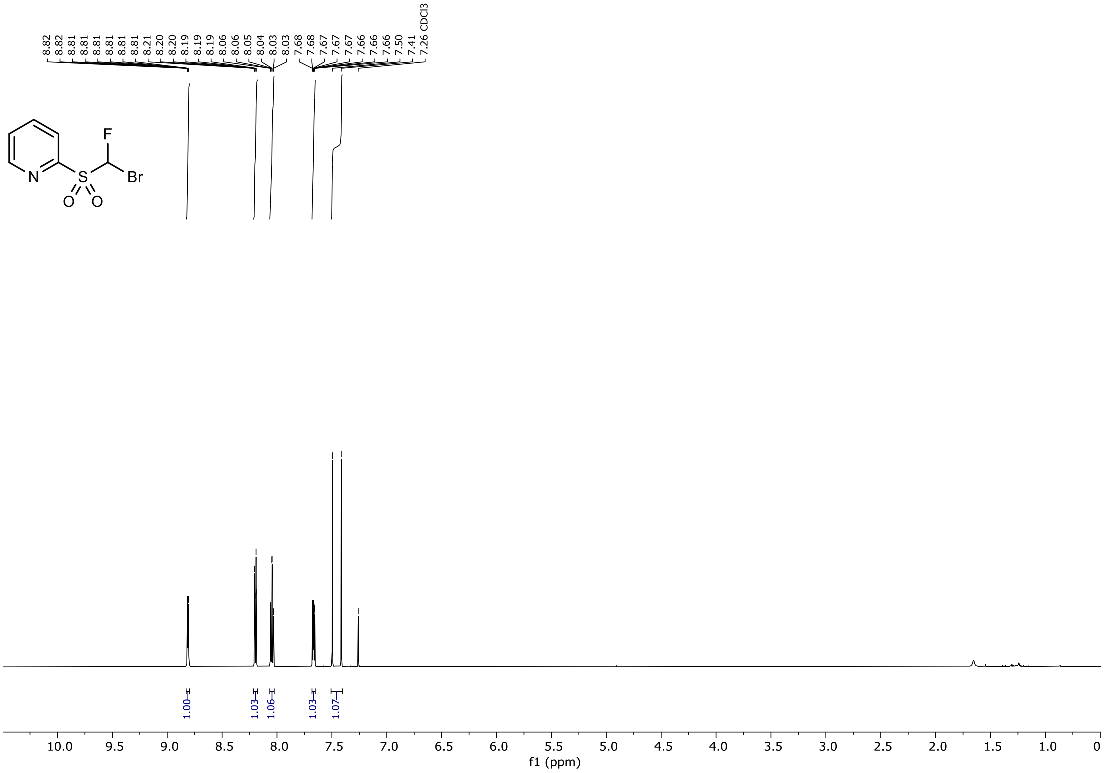


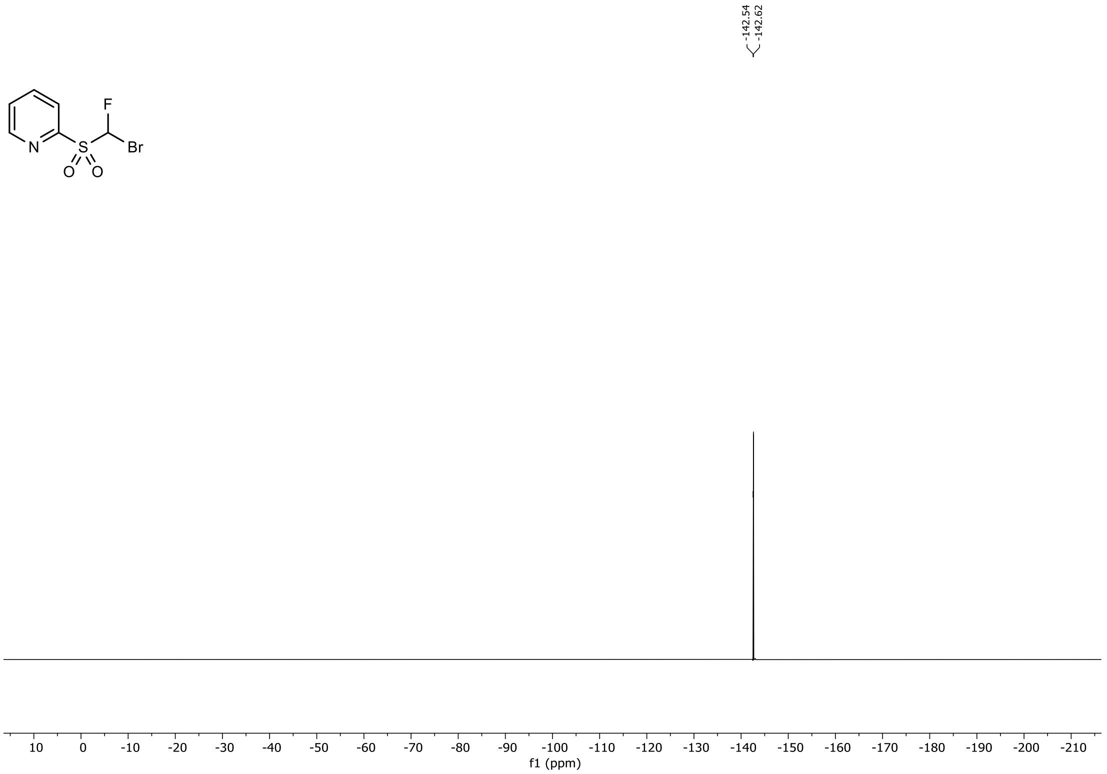

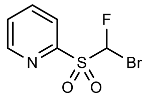

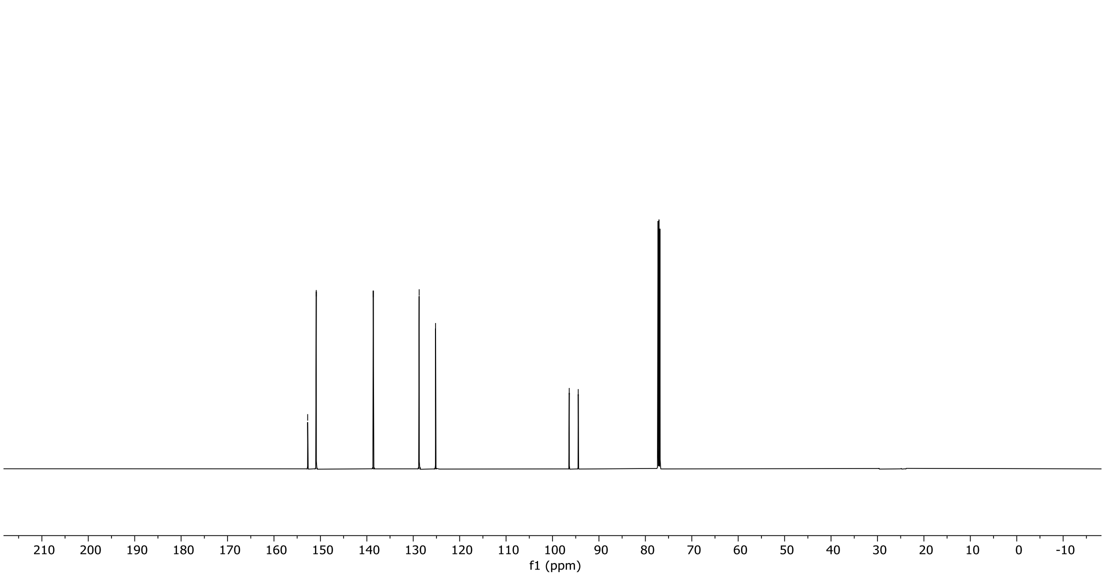
152.74
150.89
138.61
128.72
125.19
96.40
94.46
210
200
190
180
170
160
150
140
130
120
110
100
90
80
70
60
50
40
30
20
10
0
-10
f1 (ppm)

$^1H$, $^{19}F$ and $^{13}C$ NMR Spectra of 2-((bromofluoromethyl)sulfonyl)pyrimidine in $CDCl_3$

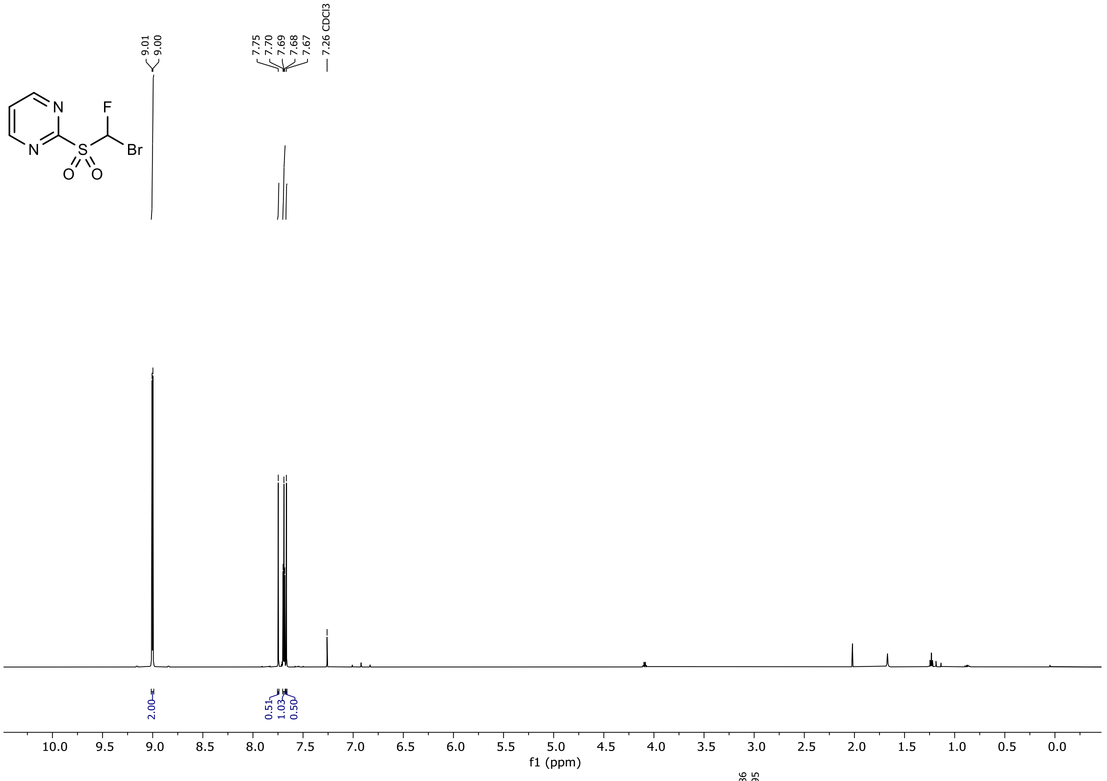


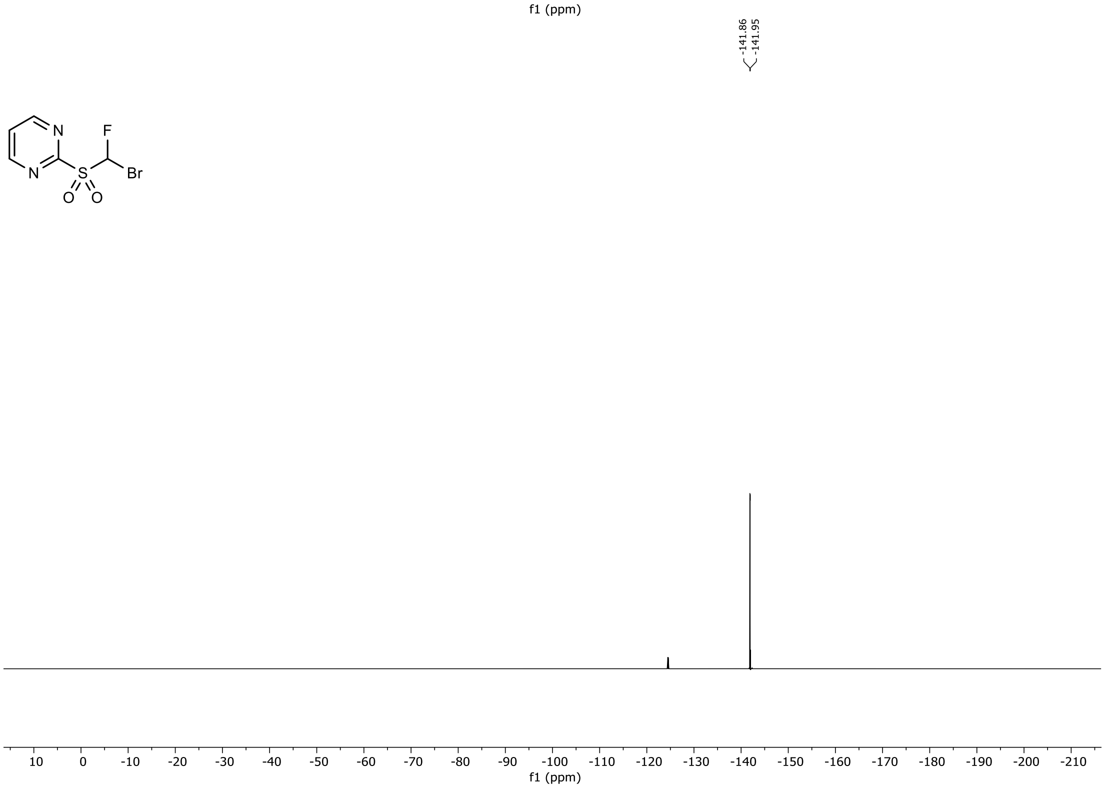

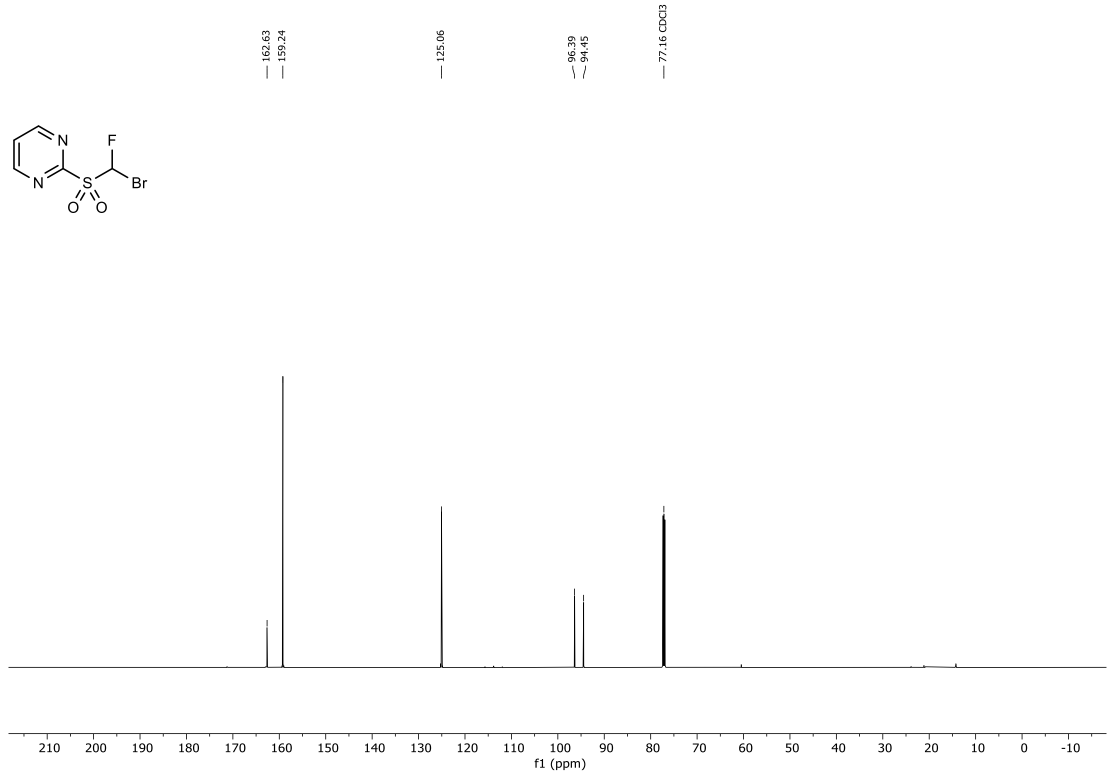
162.63
159.24
125.06
96.39
94.45
77.16 CDCl3
210 200 190 180 170 160 150 140 130 120 110 100 90 80 70 60 50 40 30 20 10 0 -10
f1 (ppm)

$^{1}H$, $^{19}F$ and $^{13}C$ NMR Spectra of 2-((bromofluoromethyl)sulfonyl)benzothiazole in $CDCl_3$

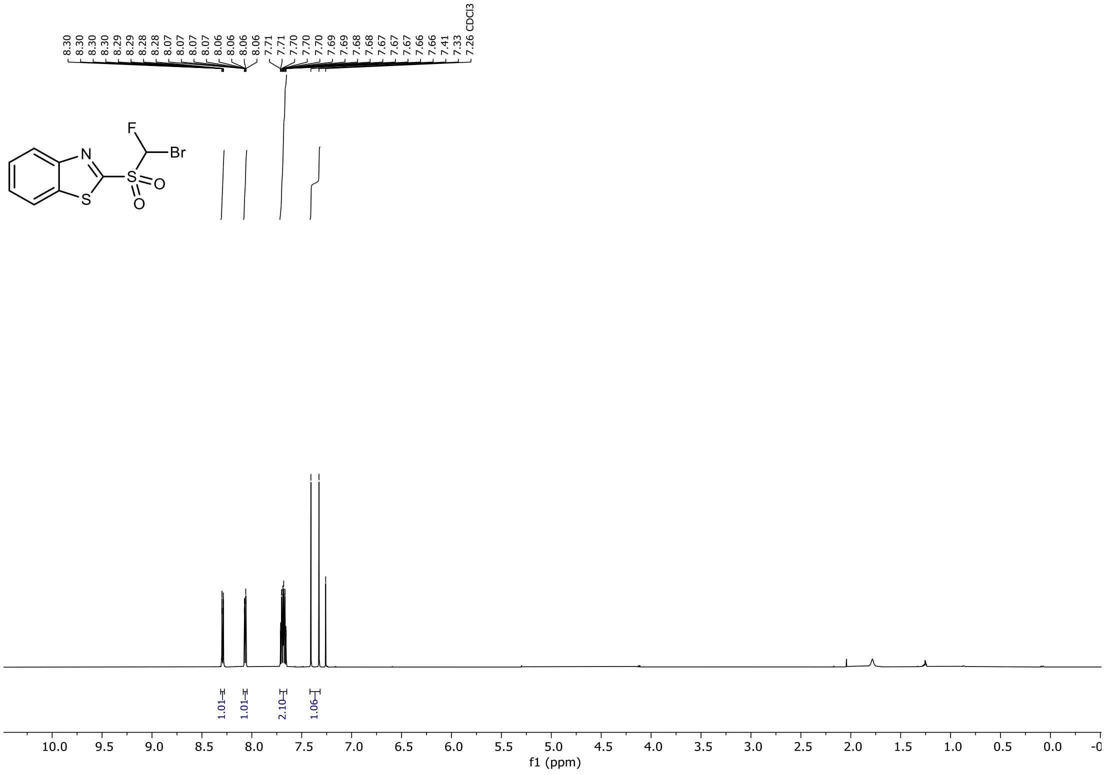


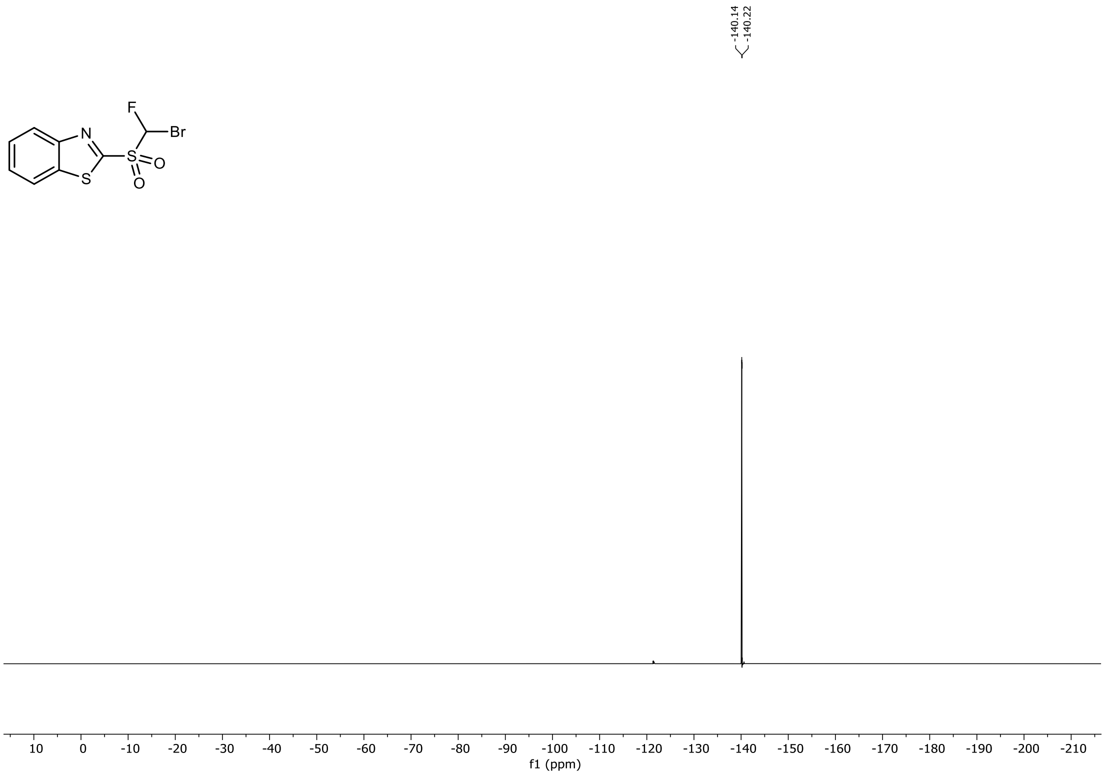

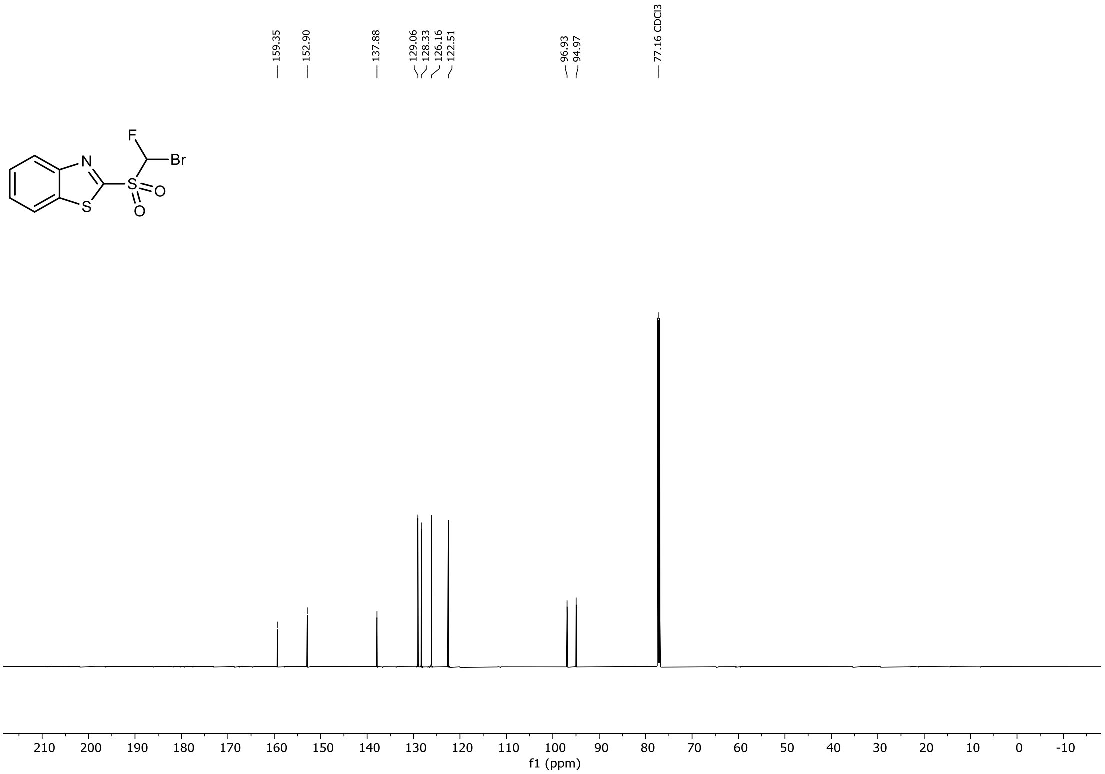

159.35
152.90
137.88
129.06
128.33
126.16
122.51
96.93
94.97
77.16 CDCl3
F
Br
N
S
S
O
O
210 200 190 180 170 160 150 140 130 120 110 100 90 80 70 60 50 40 30 20 10 0 -10
f1 (ppm)

$^{1}H$ and $^{19}F$ NMR Spectra of ethyl 2-(benzothiazole-2-ylthio)-2-fluoroacetate in $CDCl_3$

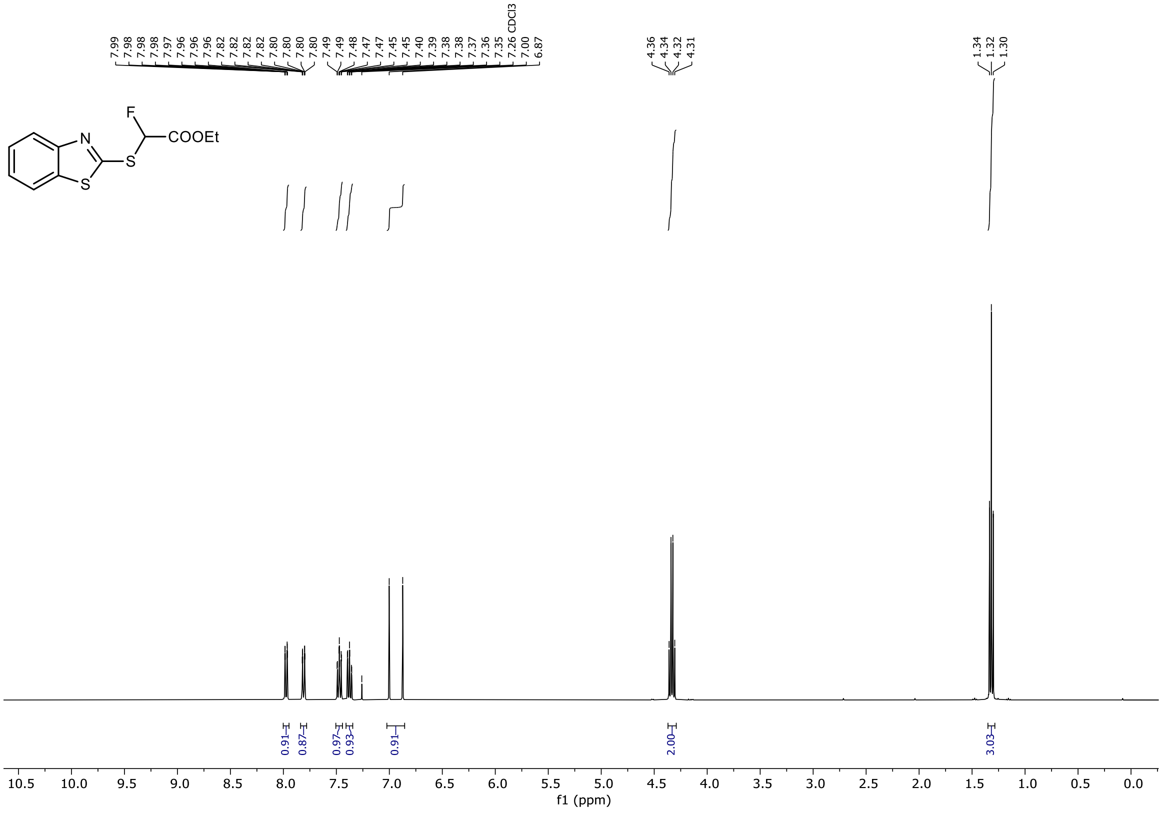

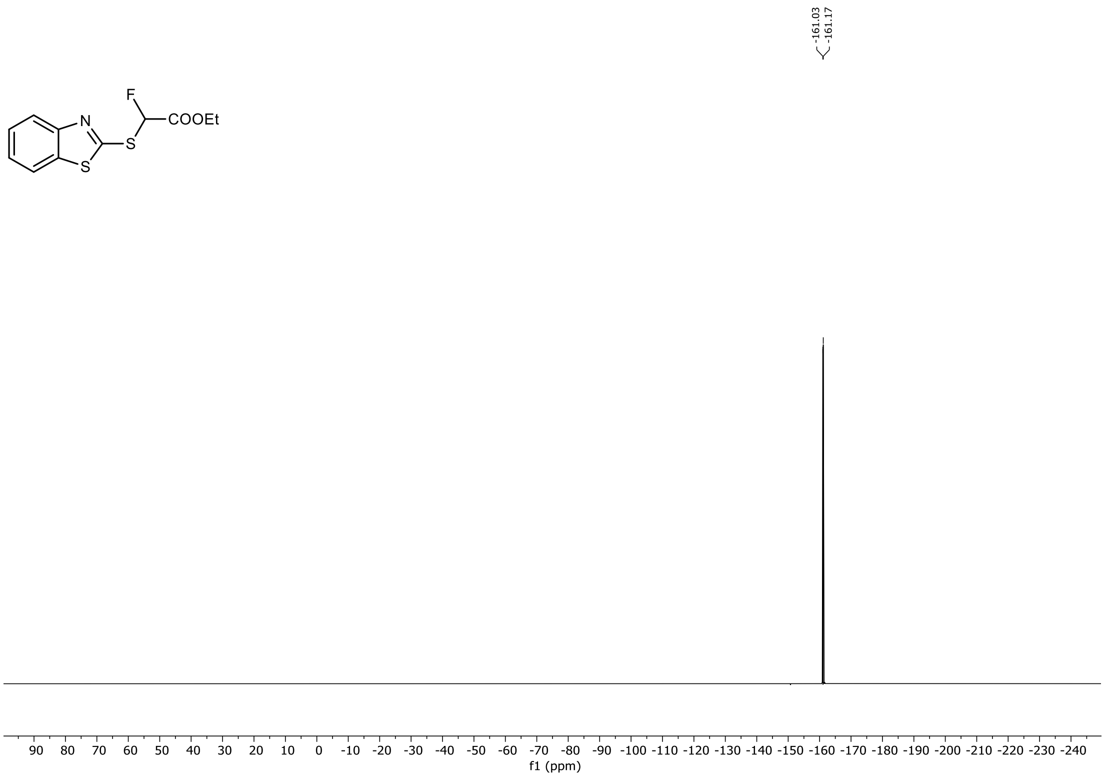

$^{1}H$ and $^{19}F$ NMR Spectra of ethyl 2-(benzothiazole-2-ylsulfonyl)-2-fluoroacetate in $CDCl_3$

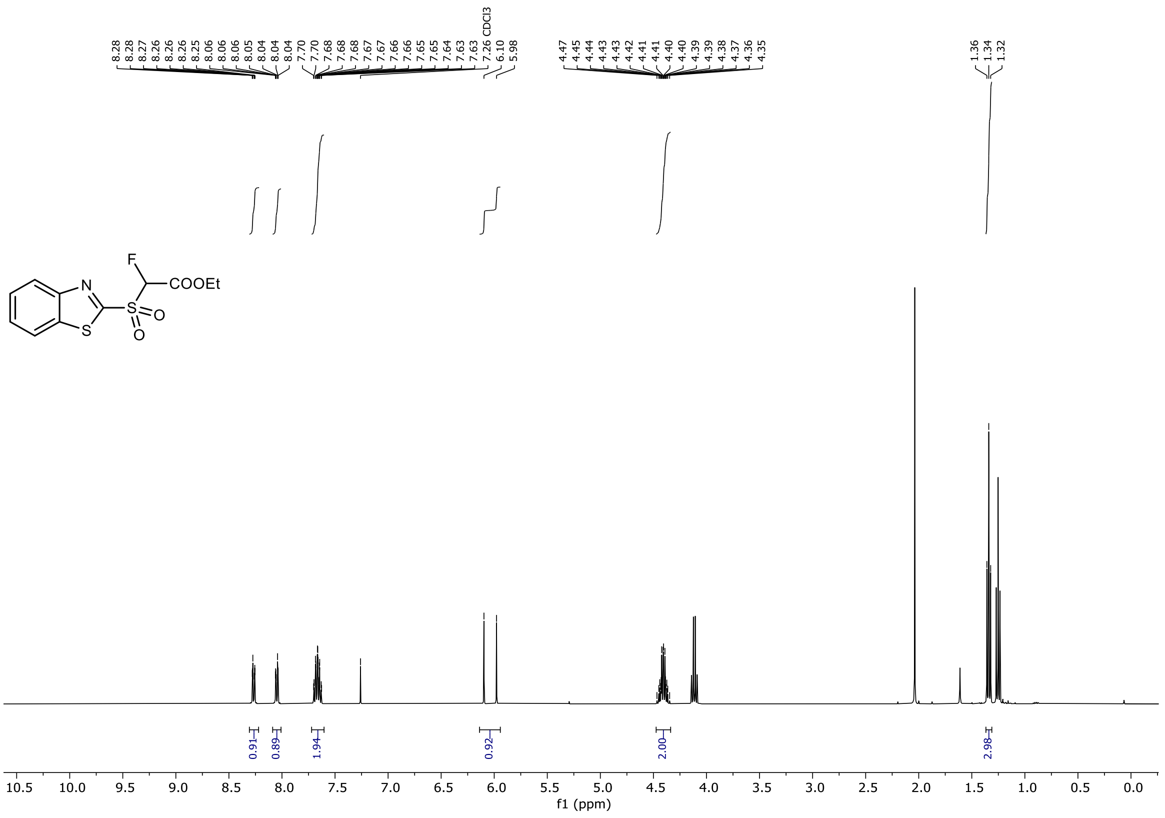

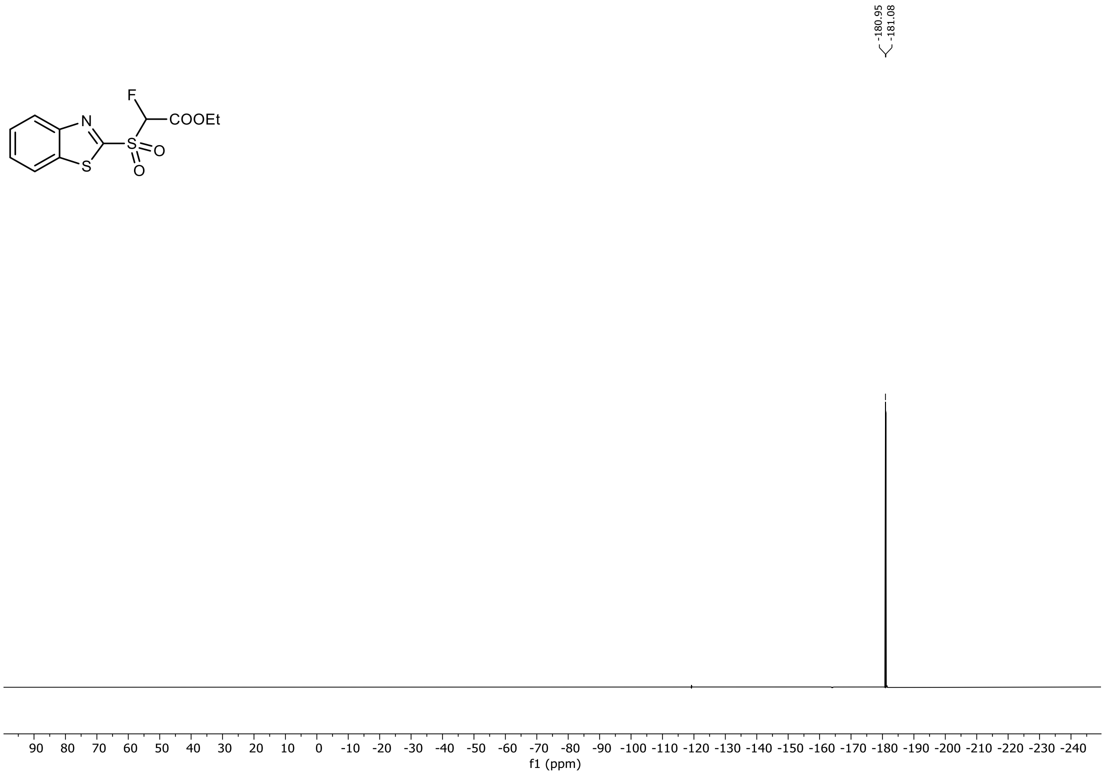

$^{1}H$ and $^{19}F$ NMR Spectra of 2-((fluoromethyl)sulfonyl)benzothiazole in DMSO-$d_6$

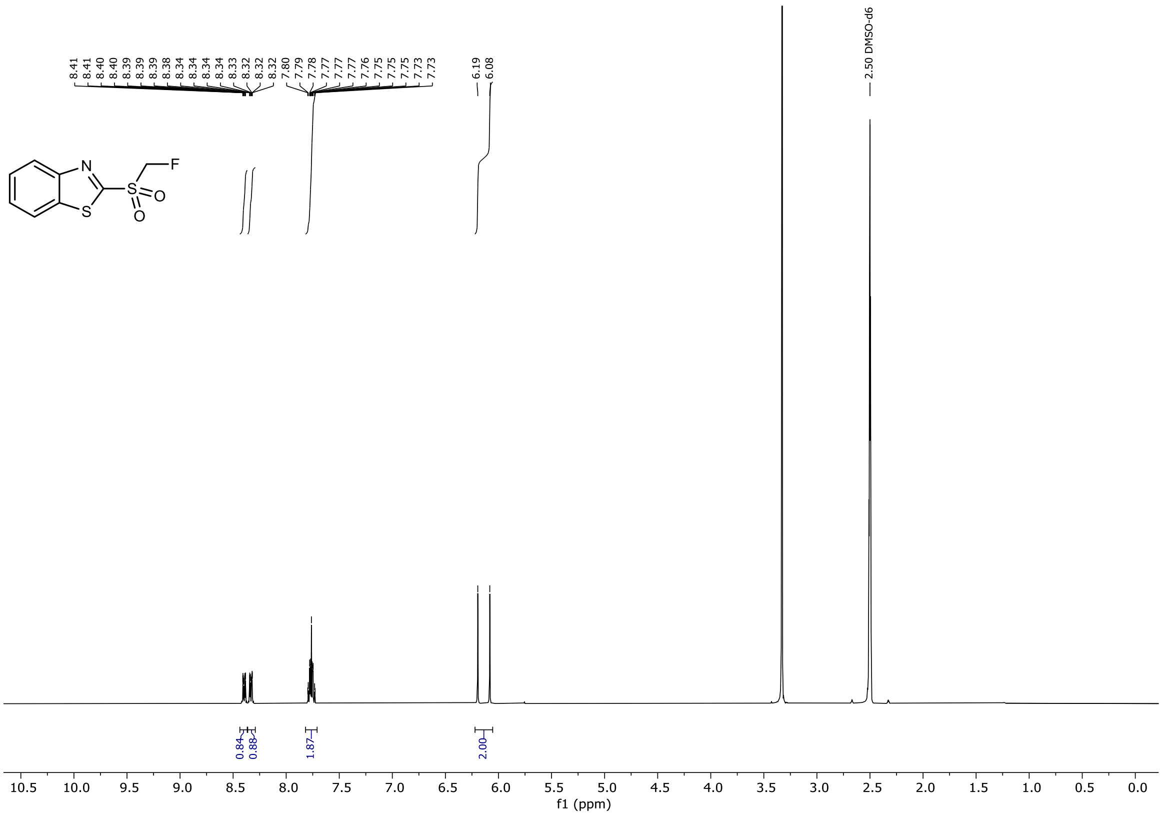

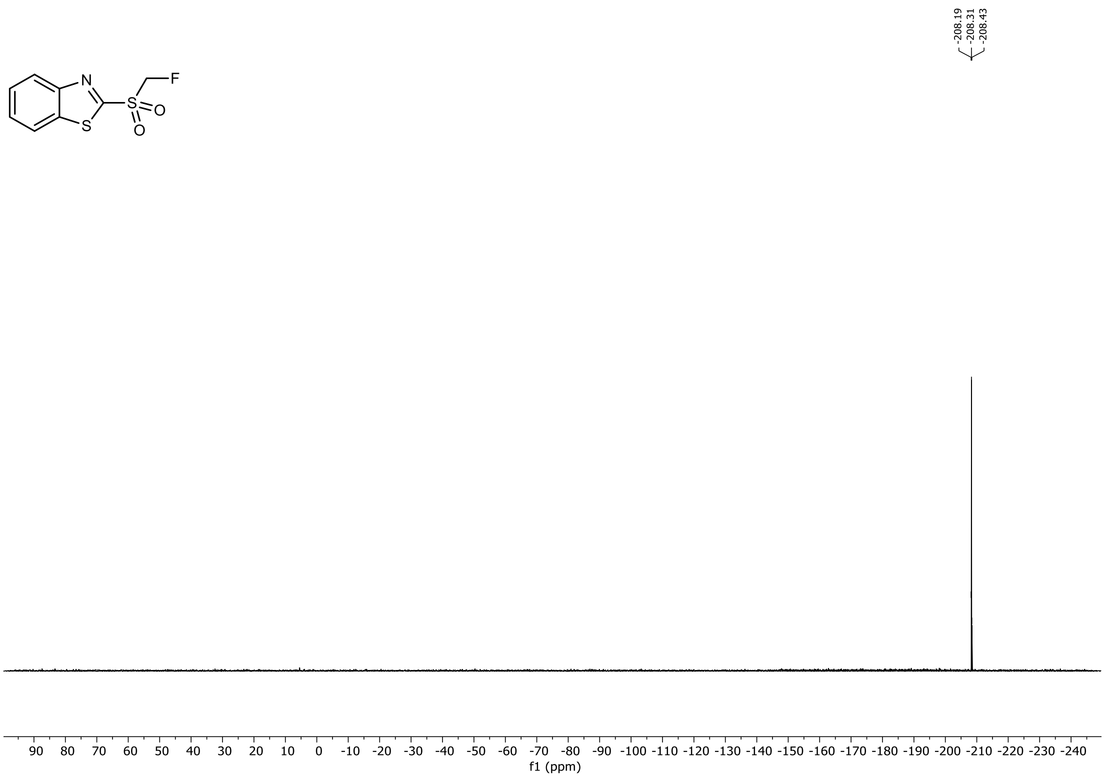

$^{1}H$ and $^{13}C$ NMR Spectra of *tert*-Butyl N-(*tert*-butoxycarbonyl)-L-homoserinate in $CDCl_3$

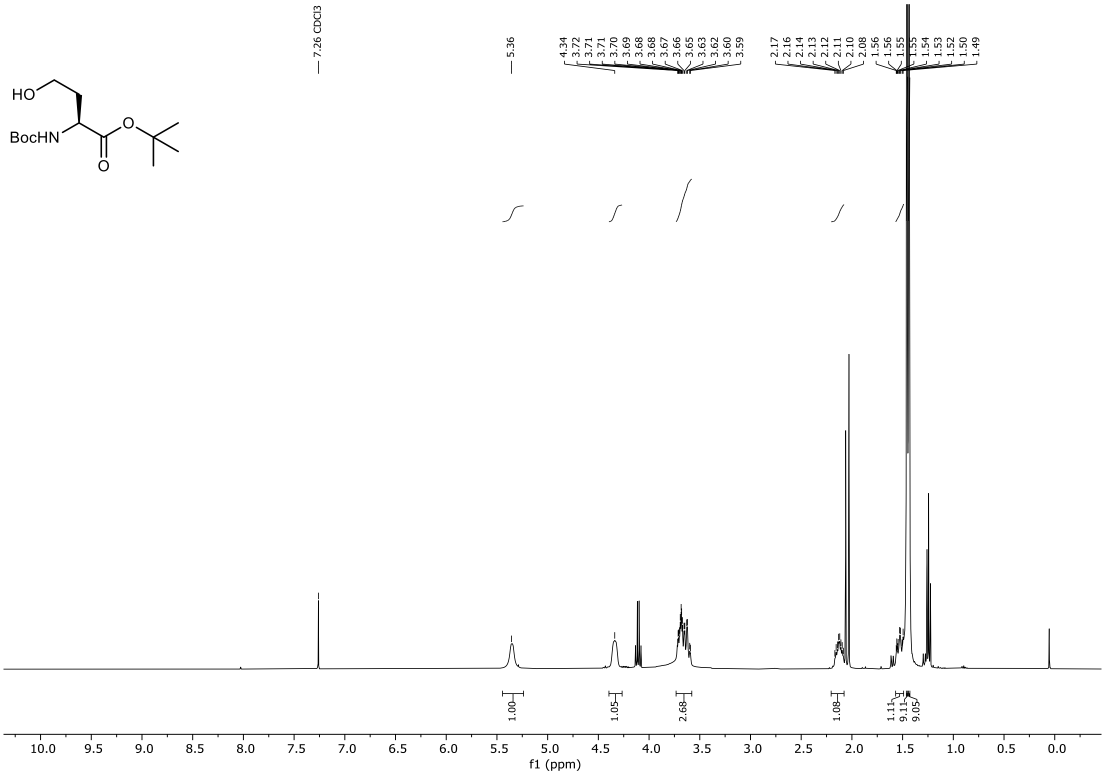


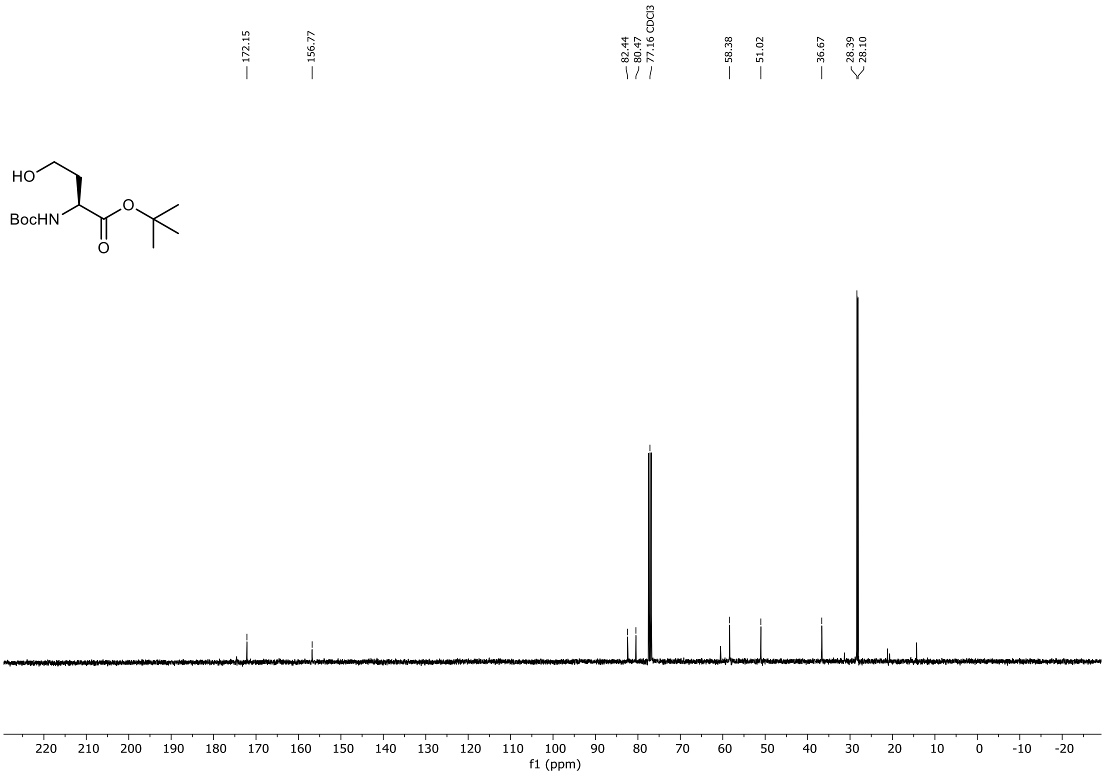

$^{1}H$ and $^{13}C$ NMR Spectra of *tert*-Butyl N-(*tert*-butoxycarbonyl)-O-tosyl-L-homoserinate in $CDCl_3$

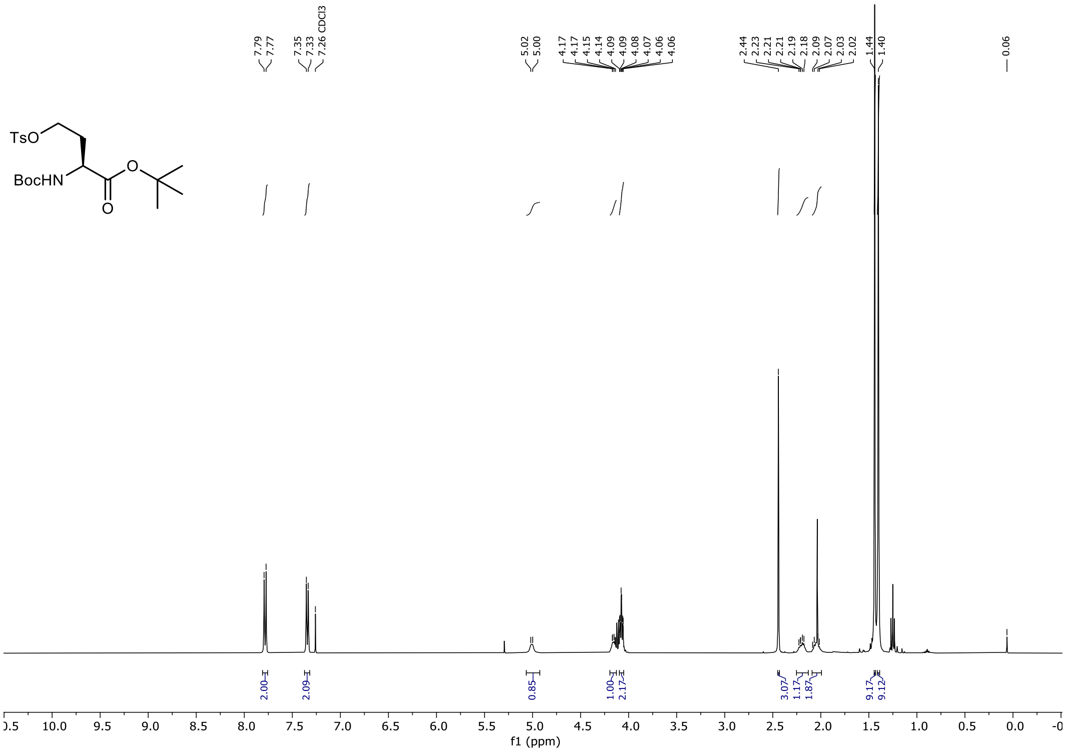


*C*H* peak at 2.02 – 2.09 ppm overlap with solvent peak at 2.04 ppm.

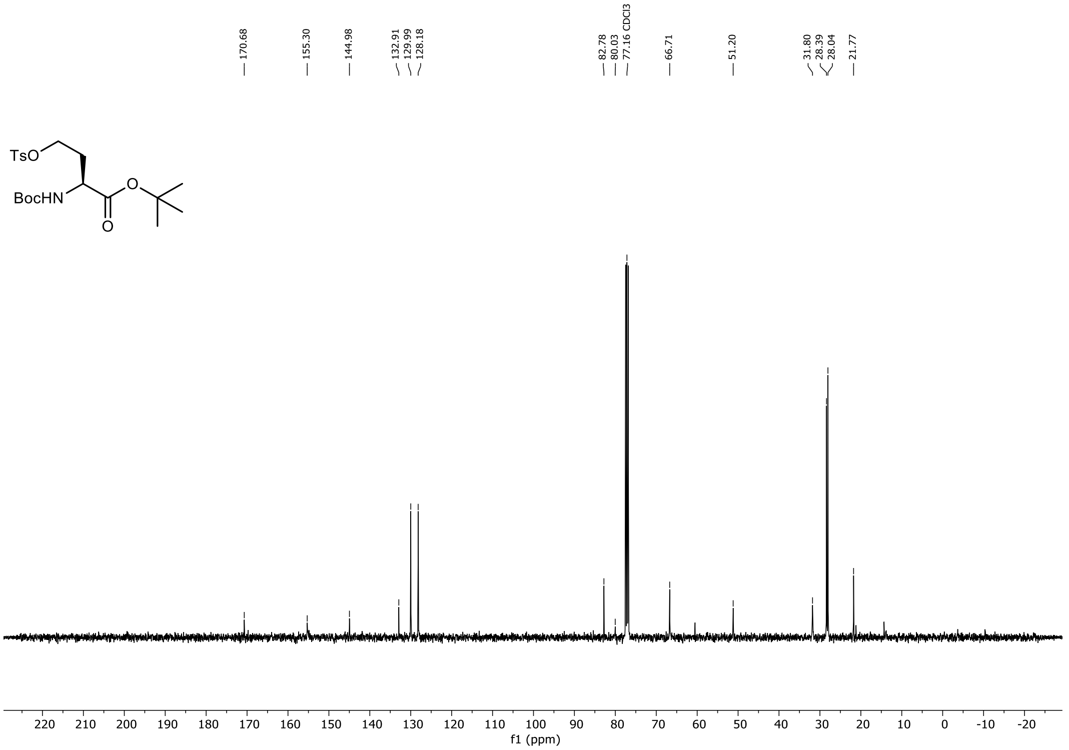

$^{1}H$, $^{19}F$ and $^{13}C$ NMR Spectra of *tert*-Butyl (S)-2-((tert-butoxycarbonyl)amino)-4-fluorobutanoate in $CDCl_3$

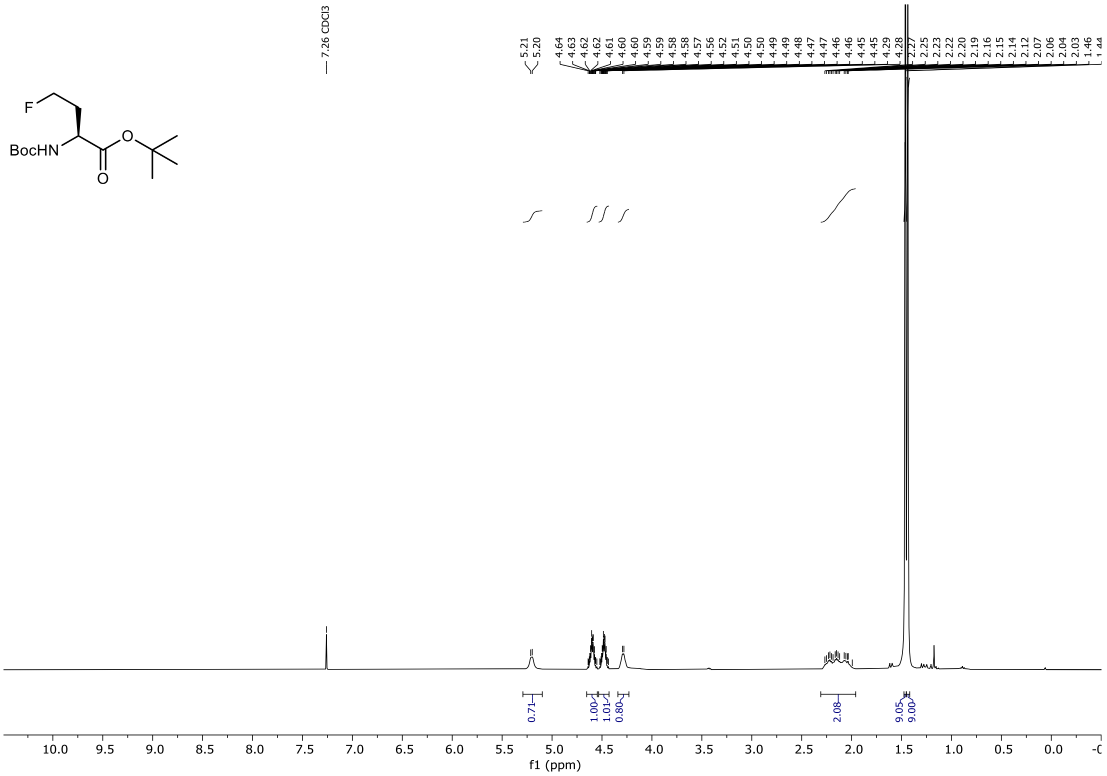


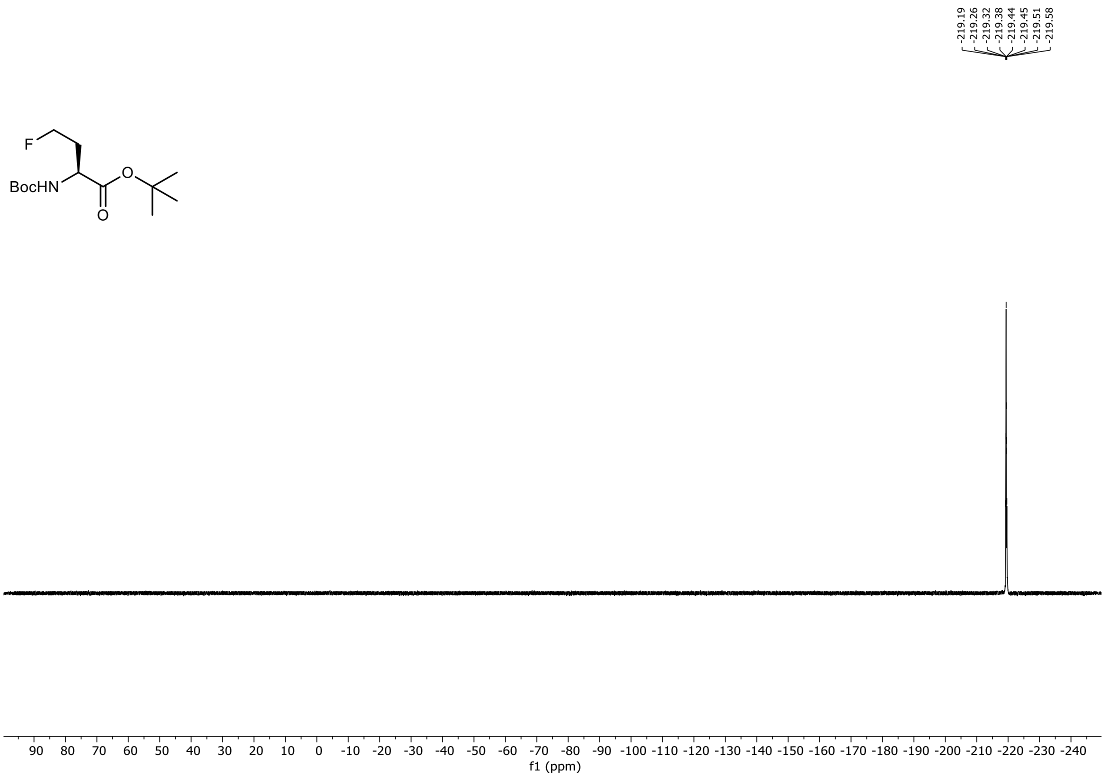

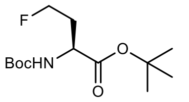
F
BocHN
O
O

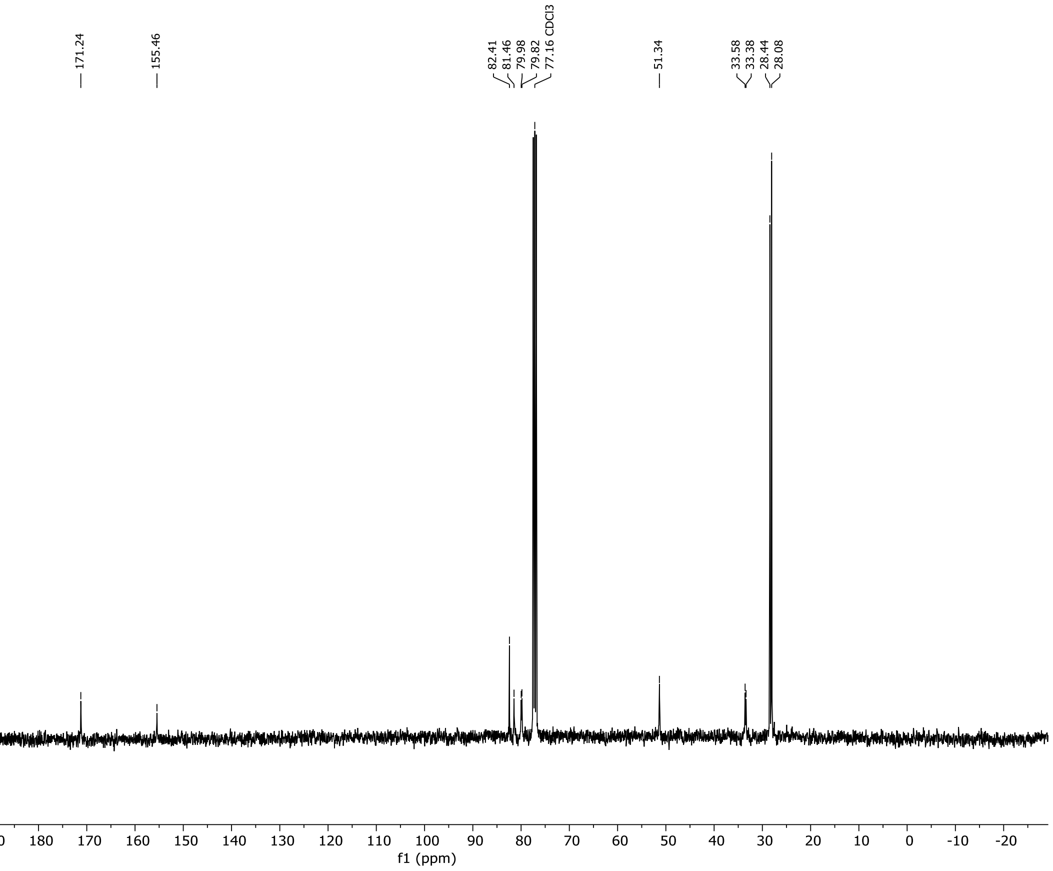
171.24
155.46
82.41
81.46
79.98
79.82
77.16 CDCl3
51.34
33.58
33.38
28.44
28.08
220 210 200 190 180 170 160 150 140 130 120 110 100 90 80 70 60 50 40 30 20 10 0 -10 -20
f1 (ppm)

[1]H, [19]F and [13]C NMR Spectra of (S)-2-((((9H-fluoren-9-yl)methoxy)carbonyl)amino)-4-fluorobutanoic acid in $(CD_3)_2CO$

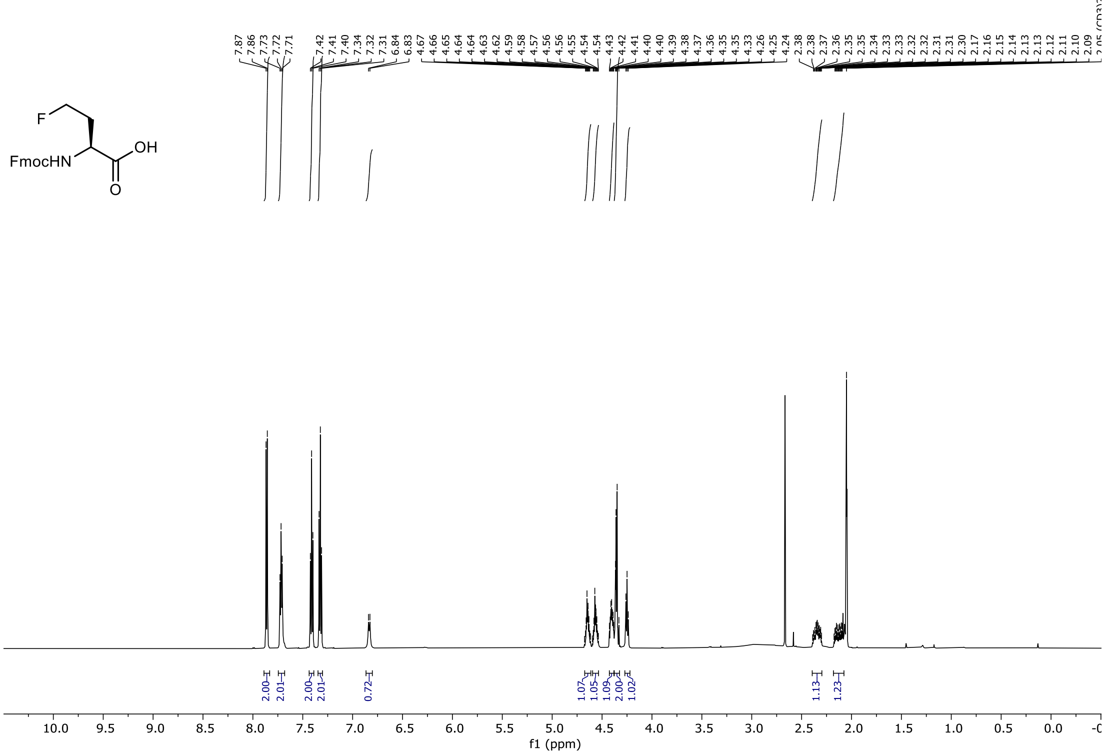


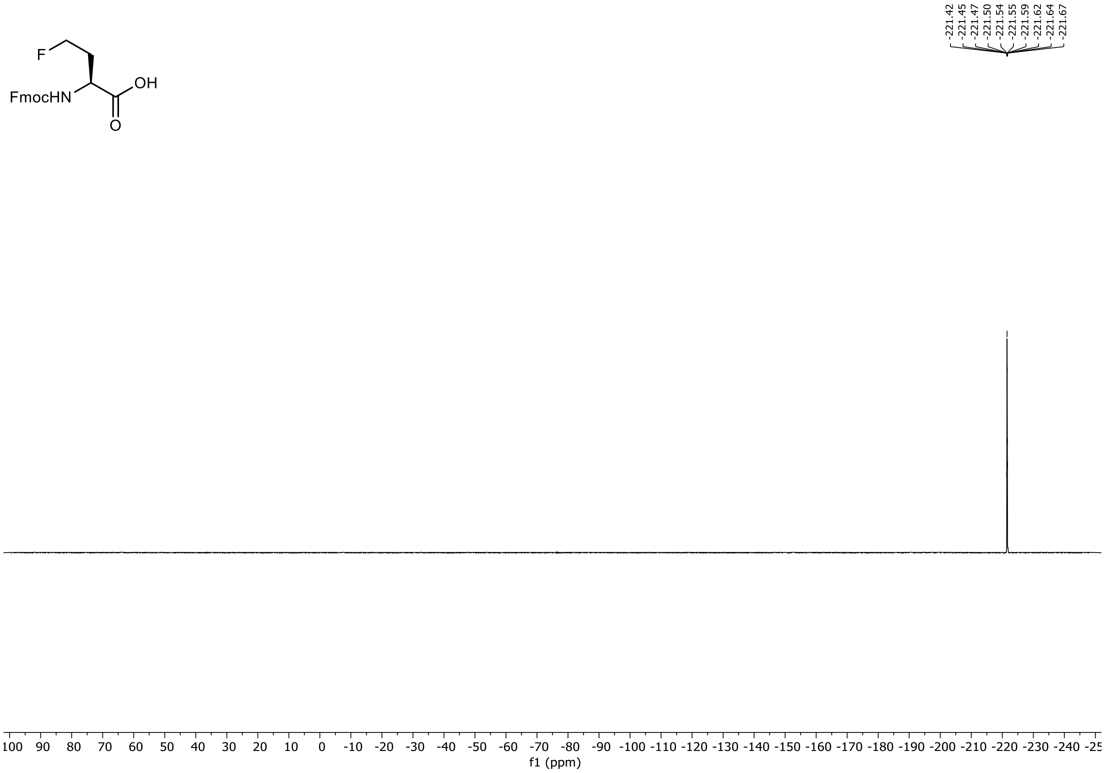

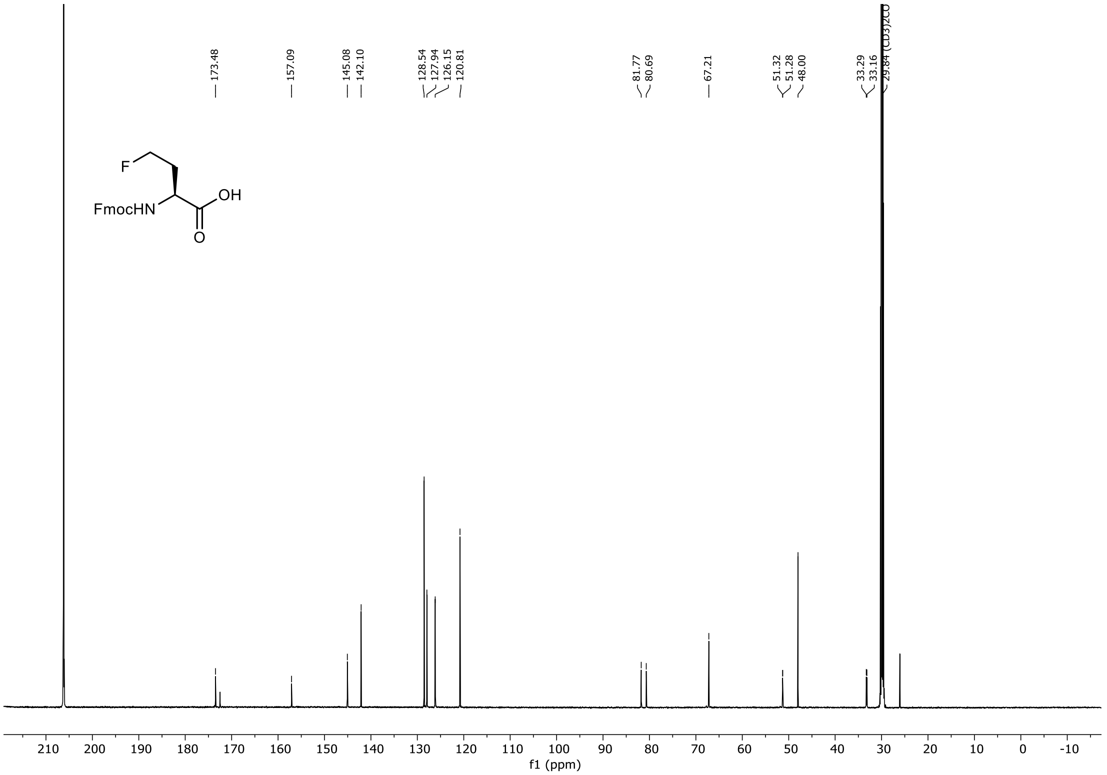
173.48
157.09
145.08
142.10
128.54
127.94
126.15
120.81
81.77
80.69
67.21
51.32
51.28
48.00
33.29
33.16
29.84 (CD3)2CO
F
FmocHN
OH
O
210 200 190 180 170 160 150 140 130 120 110 100 90 80 70 60 50 40 30 20 10 0 -10
f1 (ppm)

[1]H and [13]C NMR Spectra of *tert*-Butyl N-(*tert*-butoxycarbonyl)-D-homoserinate in $CDCl_3$

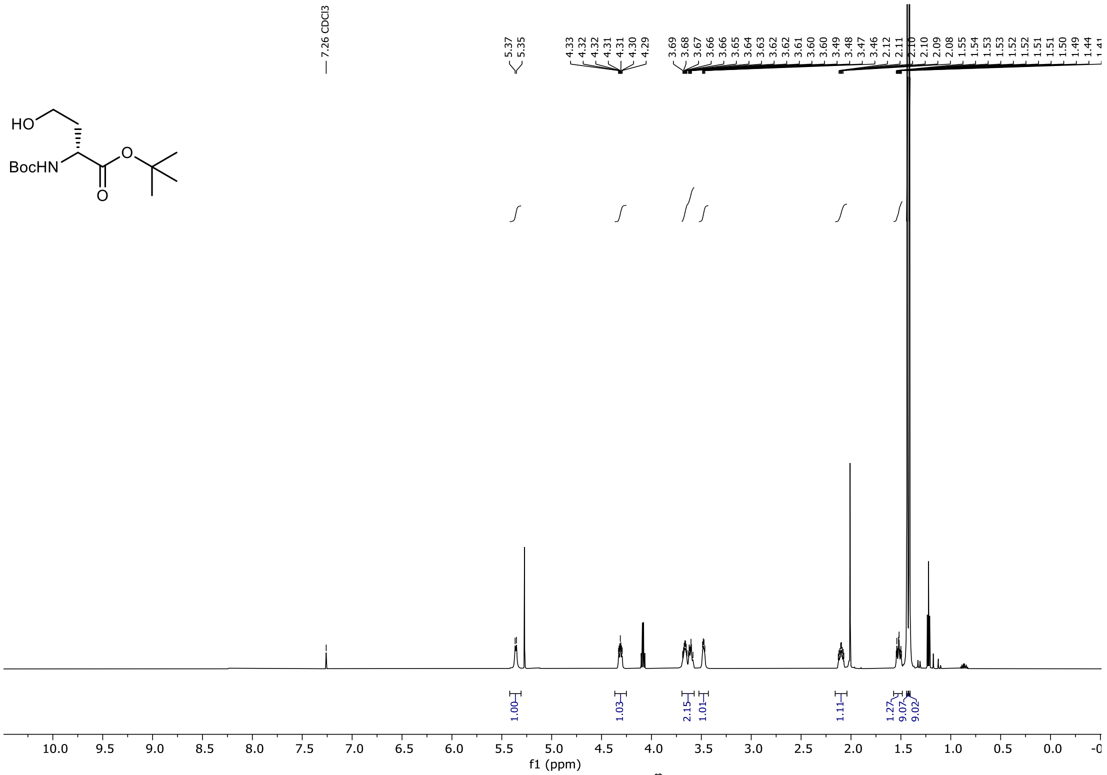


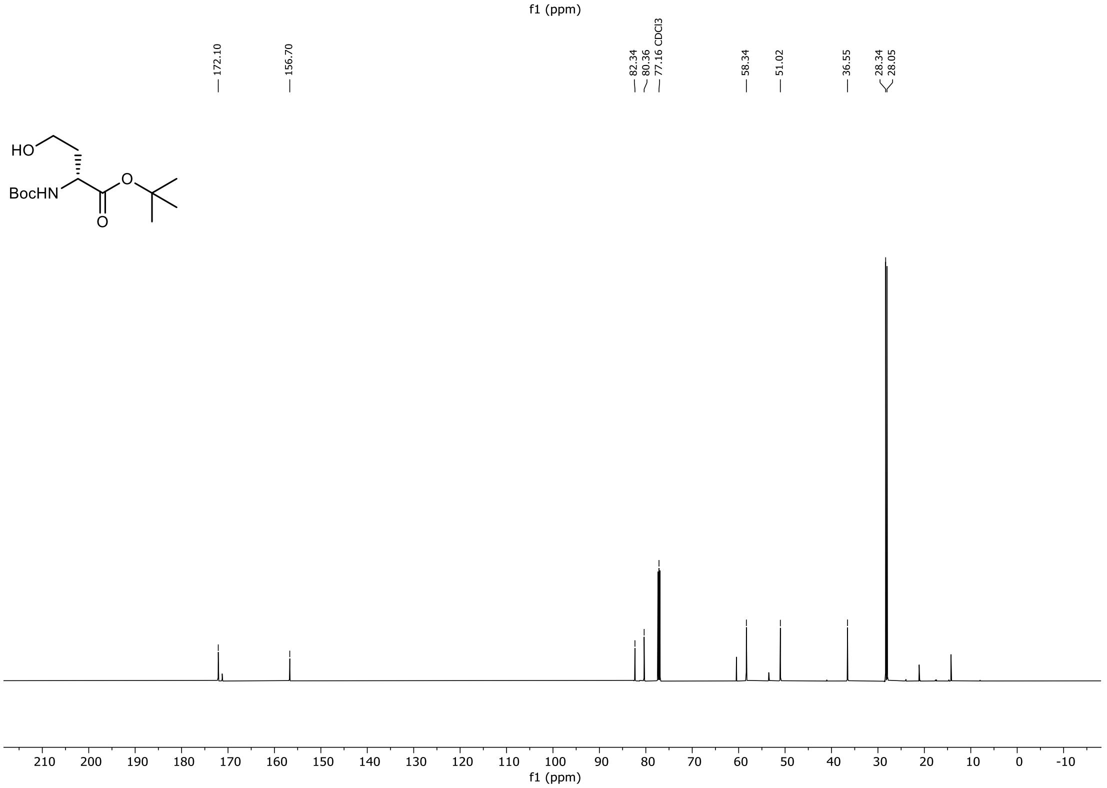

[1]H and [13]C NMR Spectra of *tert*-Butyl N-(*tert*-butoxycarbonyl)-O-tosyl-D-homoserinate in $CDCl_3$

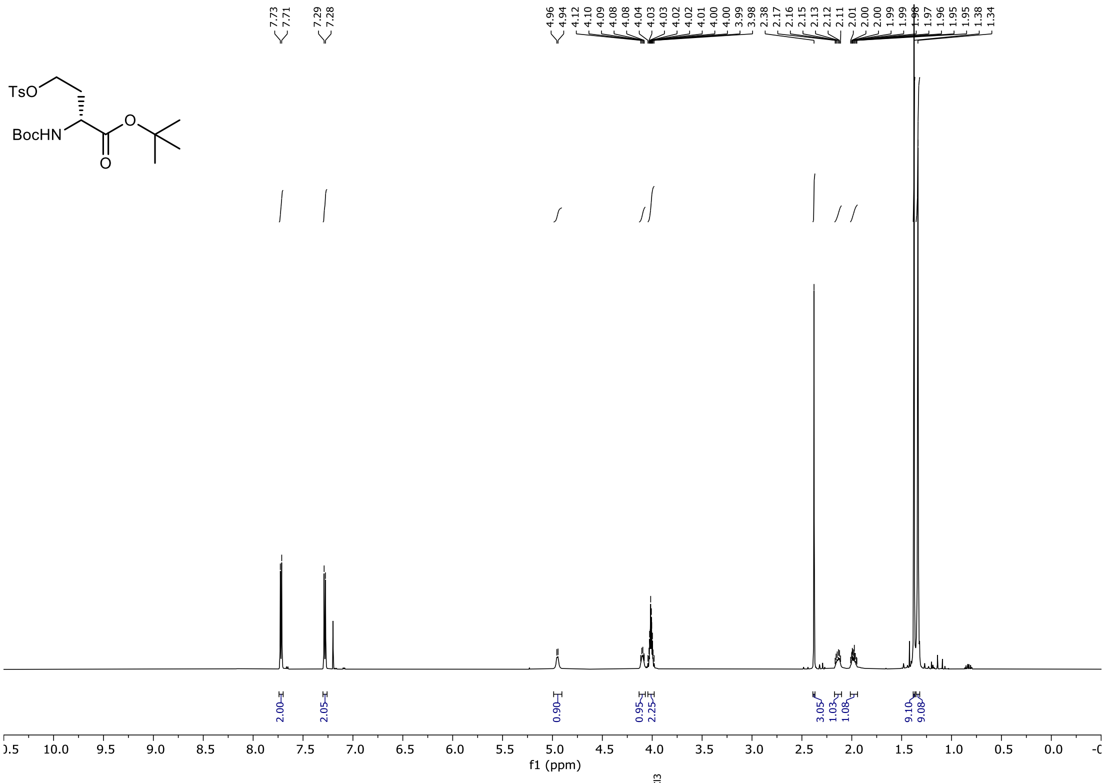


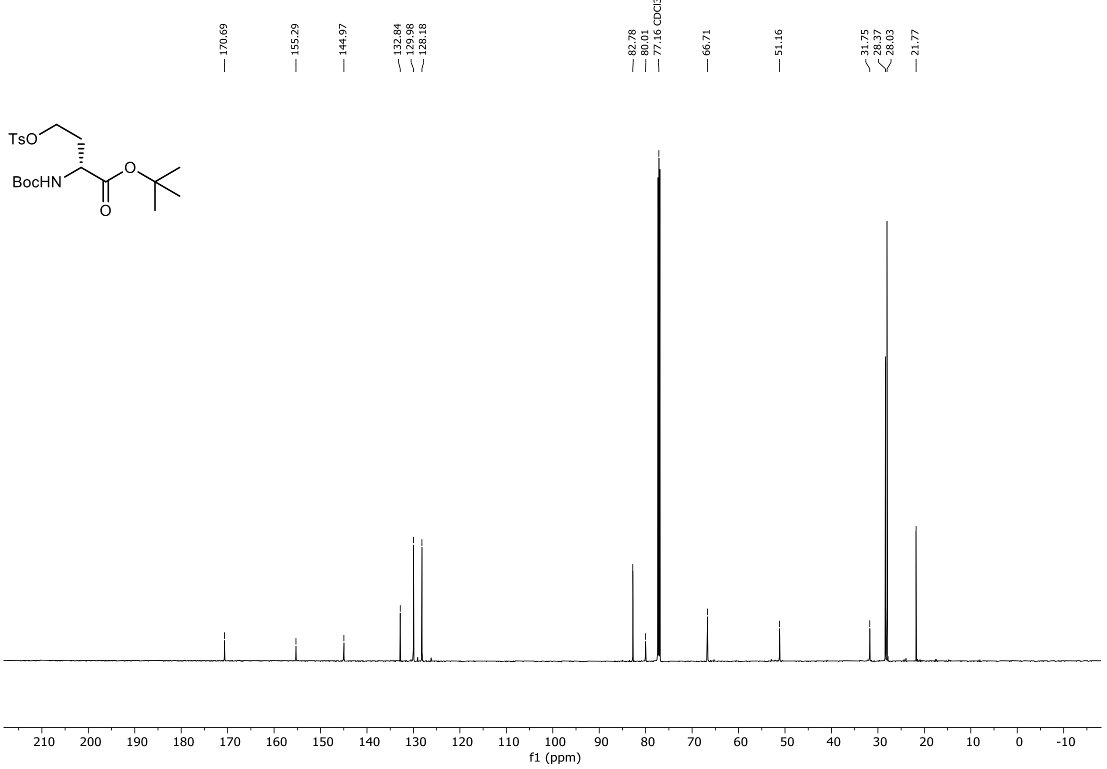

[1]H, [19]F and [13]C NMR Spectra of *tert*-Butyl (S)-2-((tert-butoxycarbonyl)amino)-4-fluorobutanoate in $CDCl_3$

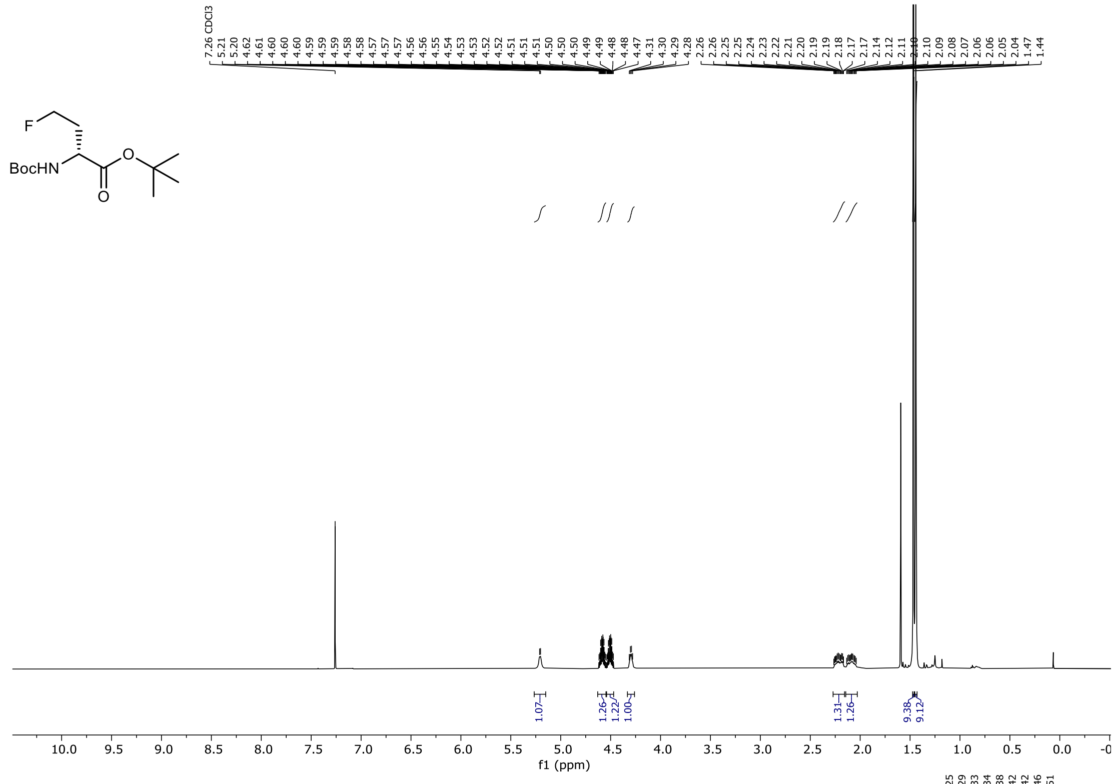


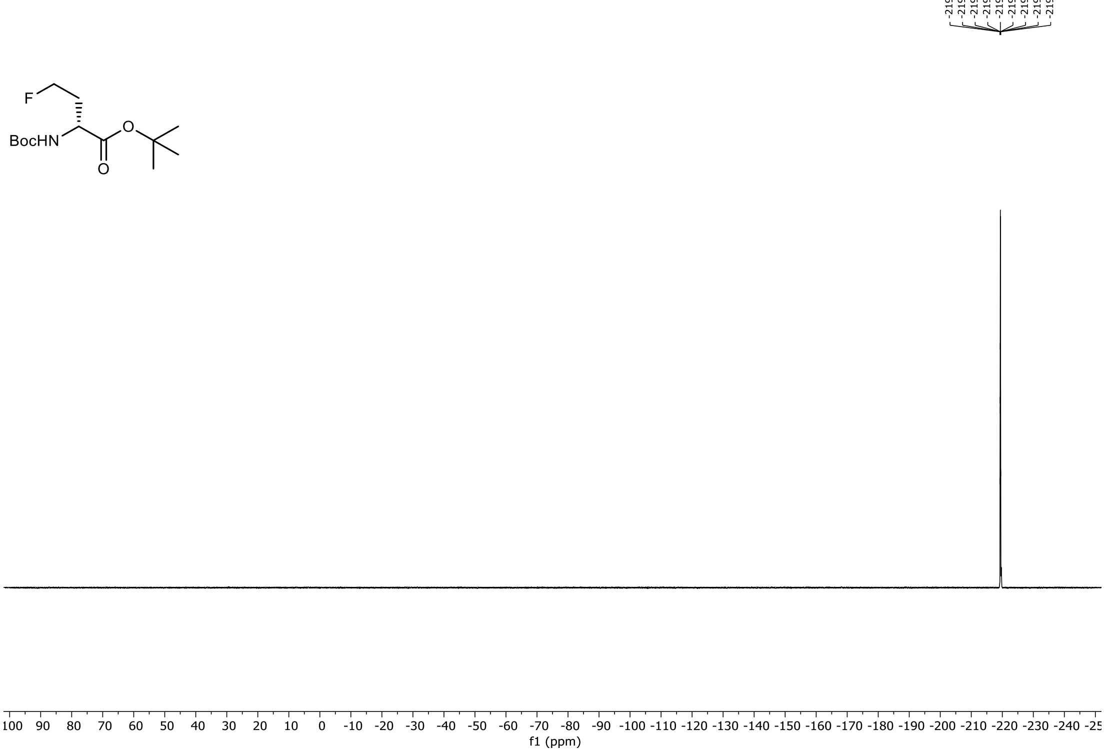

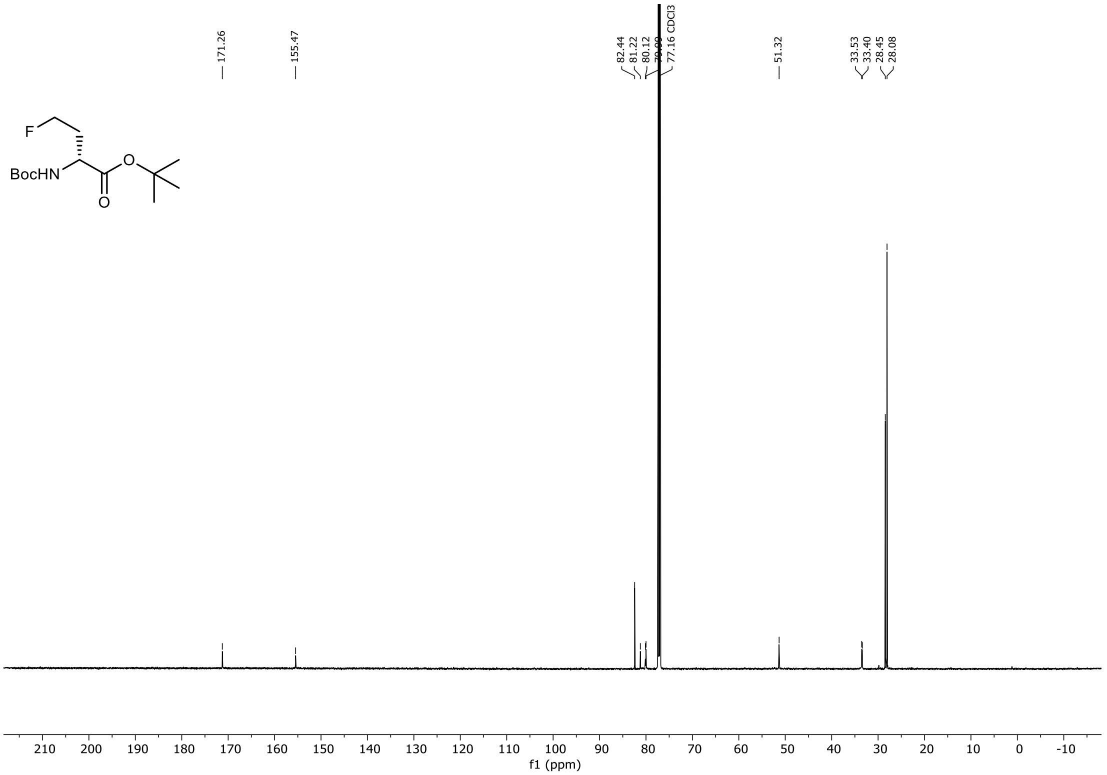

171.26
155.47
82.44
81.22
80.12
79.09
77.16 CDCl3
51.32
33.53
33.40
28.45
28.08
F
BocHN
O
O
210
200
190
180
170
160
150
140
130
120
110
100
90
80
70
60
50
40
30
20
10
0
-10
f1 (ppm)

[1]H, [19]F and [13]C NMR Spectra of (S)-2-((((9H-fluoren-9-yl)methoxy)carbonyl)amino)-4-fluorobutanoic acid in $(CD_3)_2CO$

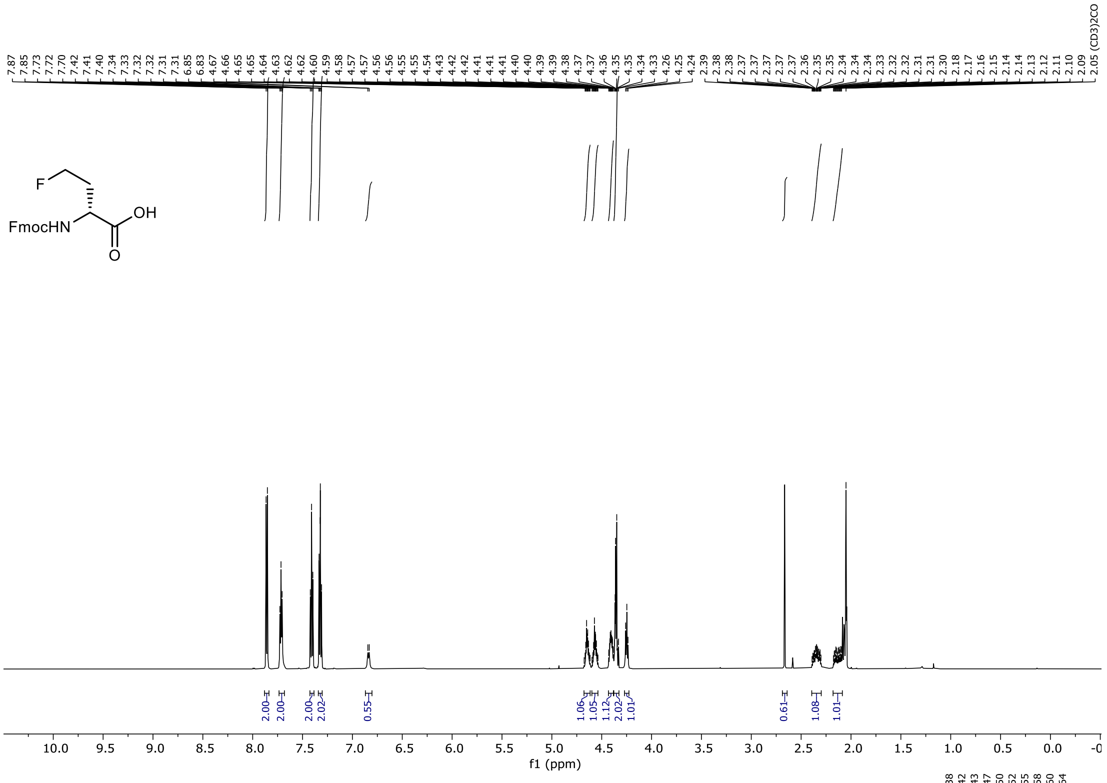

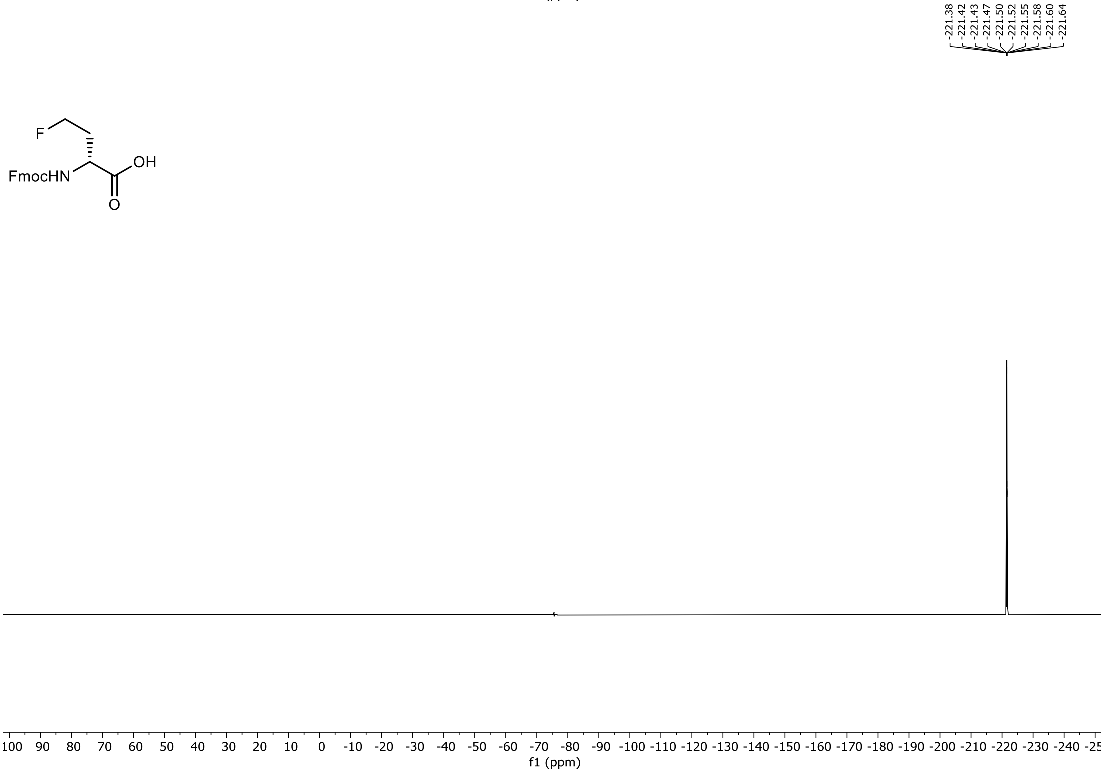

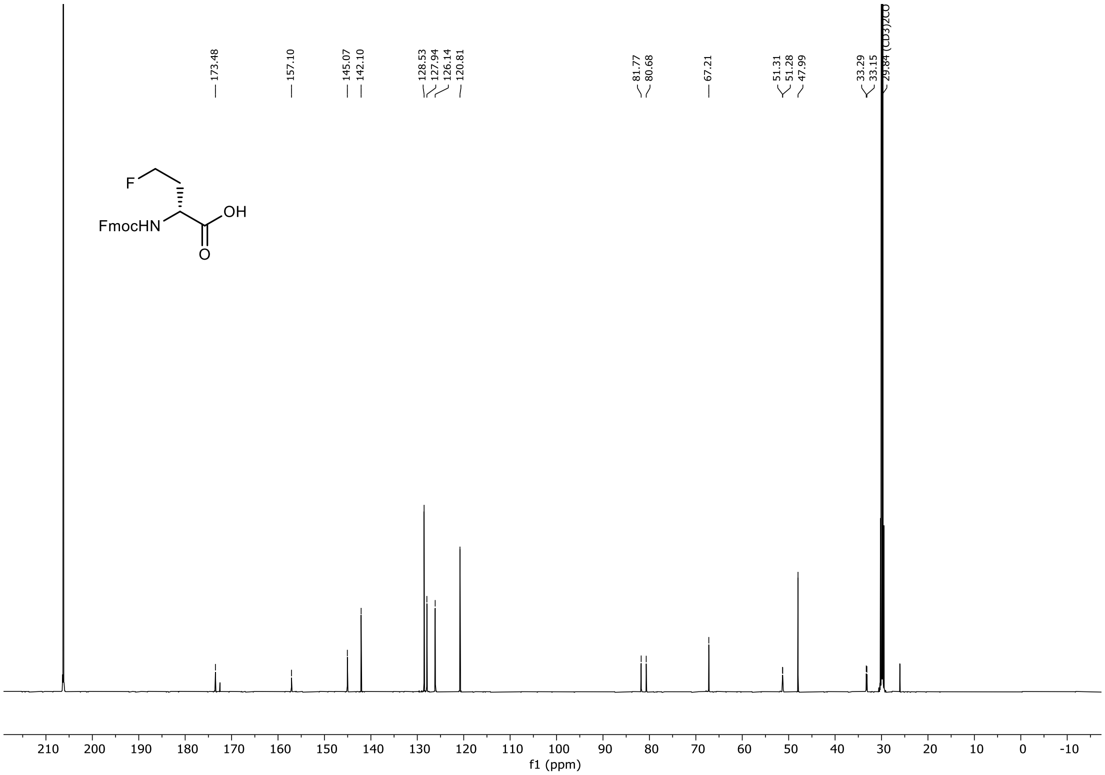

173.48
157.10
145.07
142.10
128.53
127.94
126.14
120.81
81.77
80.68
67.21
51.31
51.28
47.99
33.29
33.15
29.84 (CD3)2CO
F
OH
FmocHN
O
210 200 190 180 170 160 150 140 130 120 110 100 90 80 70 60 50 40 30 20 10 0 -10
f1 (ppm)

[1]H, [19]F and [13]C NMR Spectra of Diethyl 2-acetamido-2-(2,2-difluoroethyl)malonate in $CDCl_3$

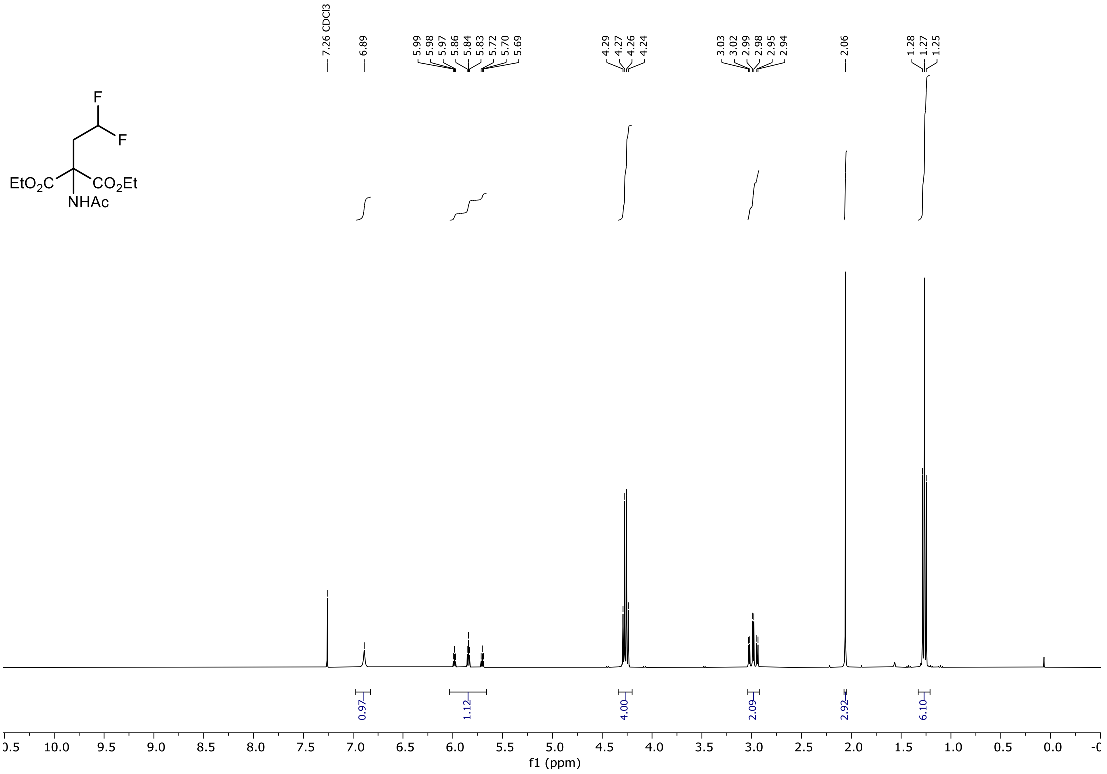

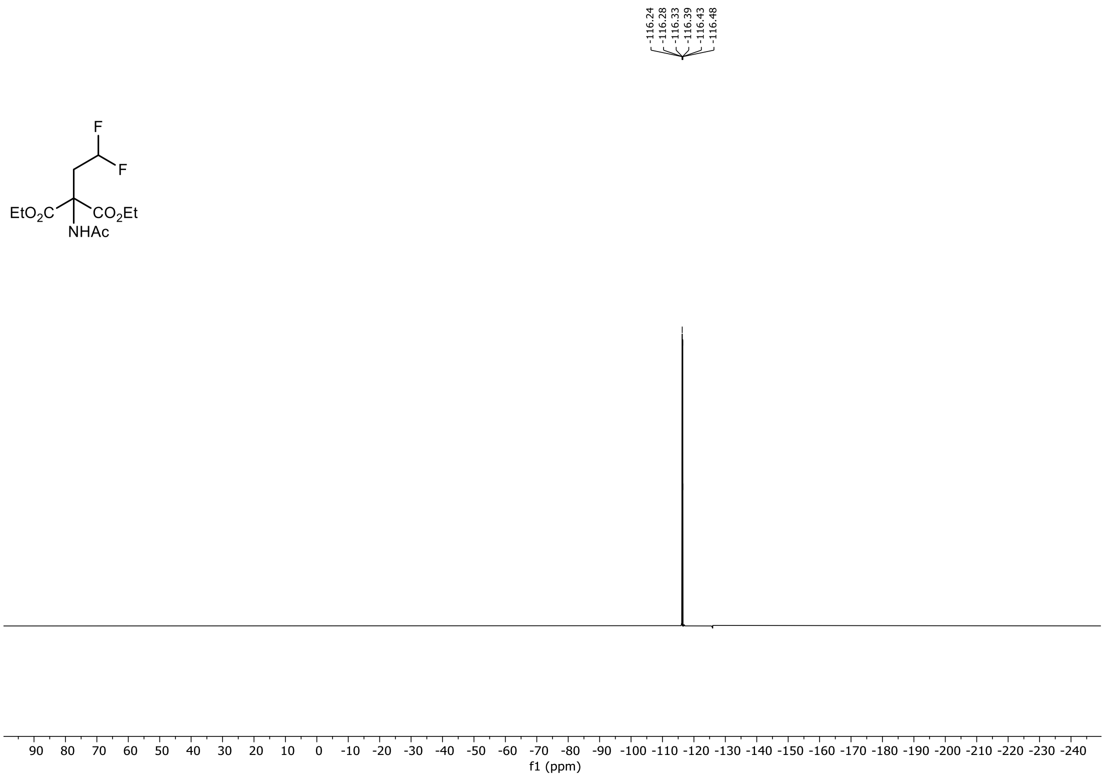

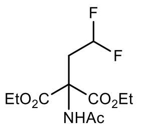
F
F
EtO2C
CO2Et
NHAc

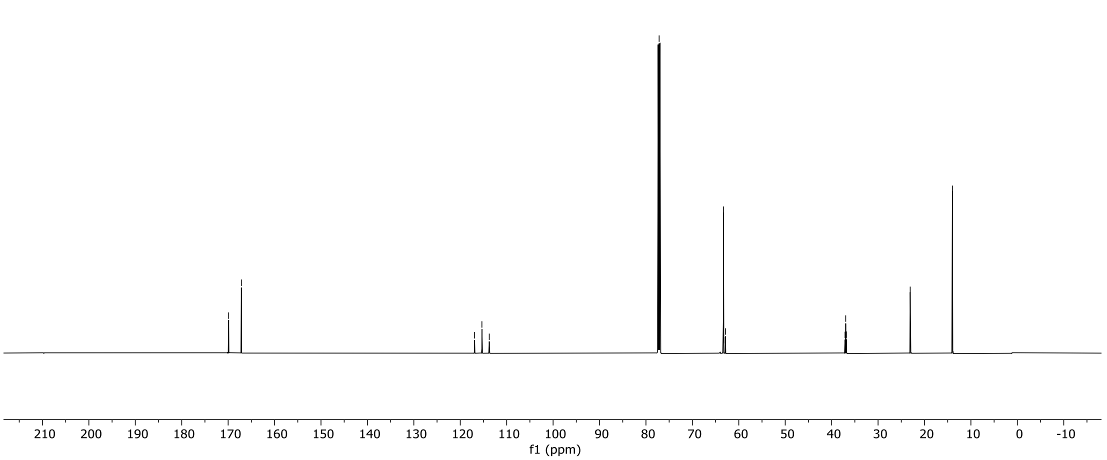
169.89
167.15
116.92
115.34
113.75
77.16 CDCl3
63.31
62.92
62.88
62.85
37.12
36.97
36.82
23.08
13.99
210 200 190 180 170 160 150 140 130 120 110 100 90 80 70 60 50 40 30 20 10 0 -10
f1 (ppm)

$^{1}H$, $^{19}F$ and $^{13}C$ NMR Spectra 2-((((9H-fluoren-9-yl)methoxy)carbonyl)amino)-4,4-difluorobutanoic acid in $(CD_3)_2CO$

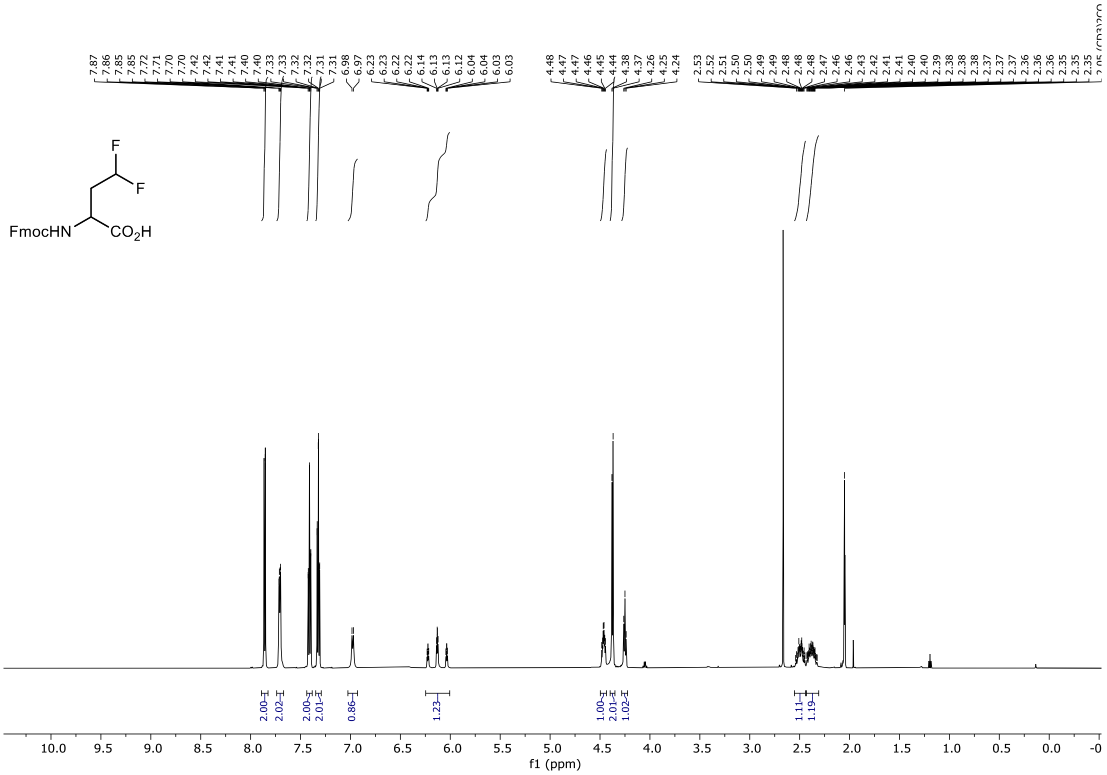


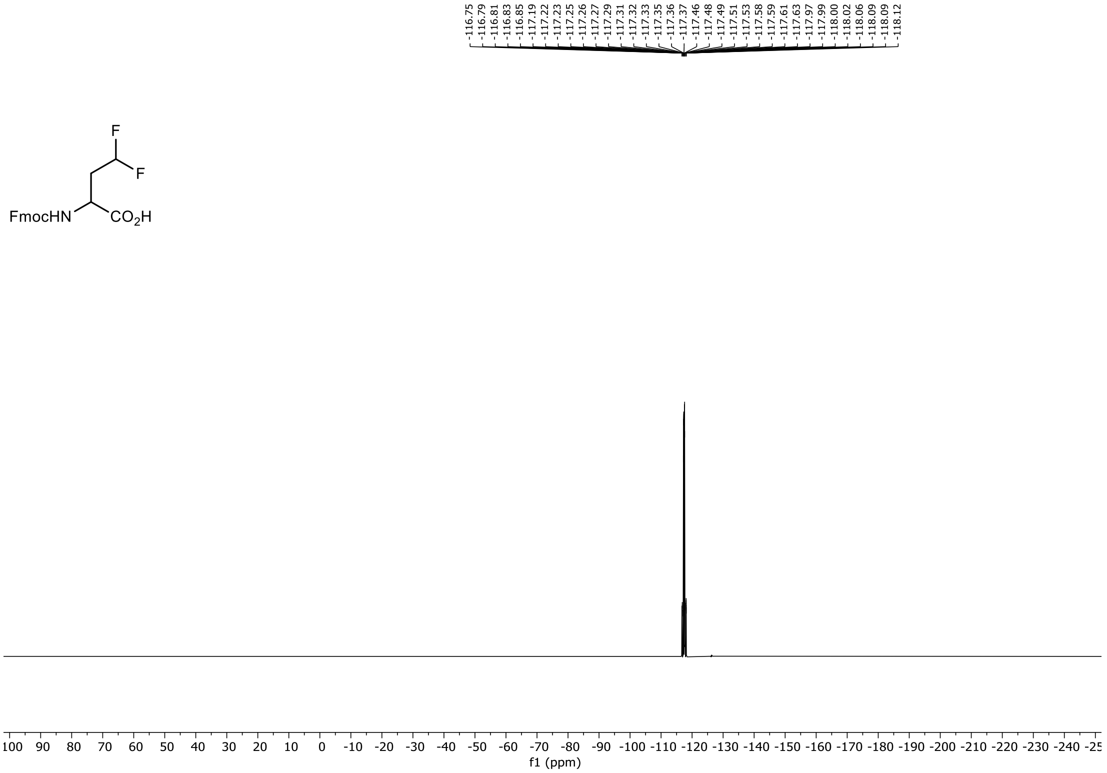

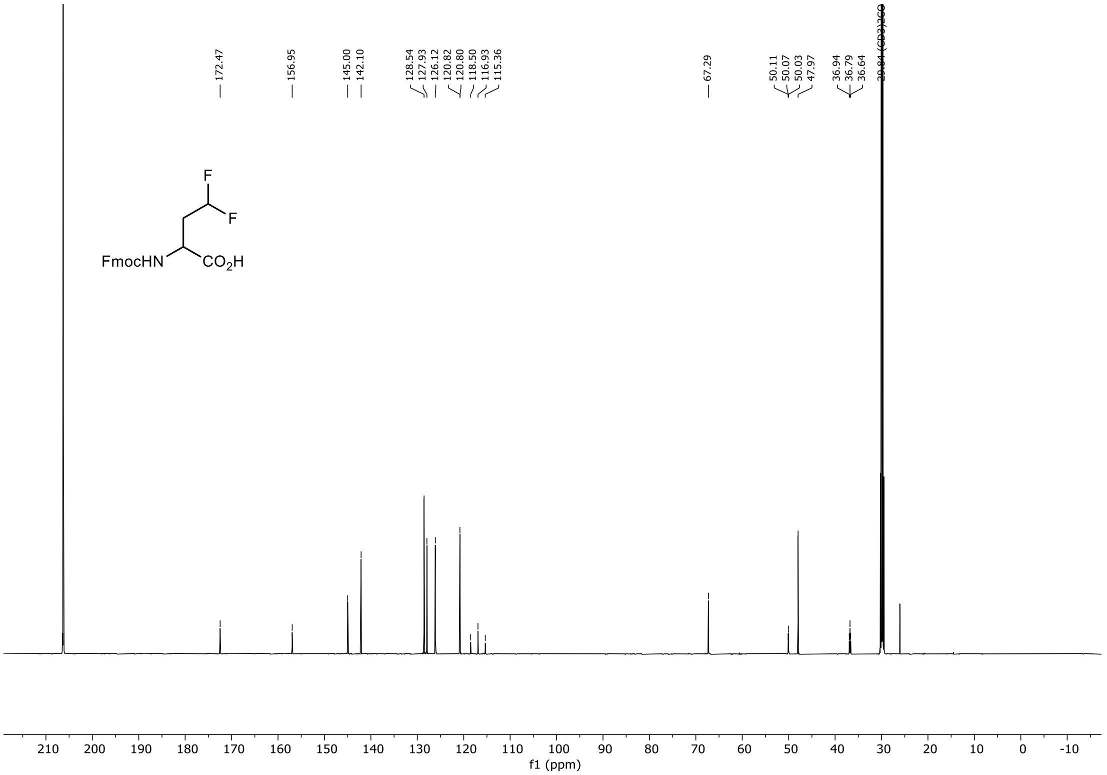

172.47
156.95
145.00
142.10
128.54
127.93
126.12
120.82
120.80
118.50
116.93
115.36
67.29
50.11
50.07
50.03
47.97
36.94
36.79
36.64
29.84 (CD3)2CO
F
F
FmocHN
CO2H
210
200
190
180
170
160
150
140
130
120
110
100
90
80
70
60
50
40
30
20
10
0
-10
f1 (ppm)
